# The *Hydractinia* cell atlas reveals cellular and molecular principles of cnidarian coloniality

**DOI:** 10.1101/2024.06.18.599157

**Authors:** David A. Salamanca-Díaz, Helen R. Horkan, Helena García-Castro, Elena Emili, Miguel Salinas-Saavedra, Maria Eleonora Rossi, Marta Álvarez-Presas, Rowan Mac Gabhann, Febrimarsa, Alberto Pérez-Posada, Nathan J. Kenny, Jordi Paps, Uri Frank, Jordi Solana

## Abstract

Coloniality is a widespread growth form in cnidarians, tunicates, and bryozoans, among others. Despite being modular, composed of multiple zooids and supporting tissues, colonies function as a single physiological unit. A major question in the biology of colonies is the cellular mechanism of generating structurally and functionally distinct colony parts. The cnidarian *Hydractinia* establishes colonies with different types of zooids (polyps), interconnected by a gastrovascular system that is attached to the substrate and known as stolons. We obtained single cell transcriptomic profiles of ∼200K *Hydractinia* cells, including isolated stolons and two polyp types. We characterised the major *Hydractinia* cell types and quantified their abundance across colony parts. Overall, we find that distinct colony parts are characterised primarily by distinct combinations of shared cell types and to a lesser extent by part-specific cell types. Therefore, we propose that both cell type combinations, as well as rarer cell type innovations, have been the main mechanism in the evolution of coloniality in cnidarians. We identified cell type-specific transcription factors (TFs) and gene networks expressed within these cell types. Notably, we discovered a previously unidentified, stolon-specific cell type, which expresses enzymes related to biomineralization and chitin synthesis, reminiscent of molluscan shell matrix proteins that may represent a crucial adaptation to the animal’s habitat. In summary, the *Hydractinia* cell atlas elucidates the fundamental cellular and molecular mechanisms underlying coloniality.

## Introduction

Coloniality ^1^ is a growth form that occurs in diverse animal groups including cnidarians ^2^, tunicates ^3^, and bryozoans ^4^, among others. Contrary to solitary animals, colonial organisms are modular, consisting of individual, clonal zooids that are connected by live tissue, sharing a vascular network, nervous system, and migratory cells ^5–8^. Growth and reproduction are mostly integrated in colonial animals; therefore, a single colony functions as a meta-individual, being a single physiological unit. All colonial animals exhibit robust regenerative capacities that are based, at least in some species, on adult pluripotent stem cells ^9^. Zooids within a colony may be polymorphic, i.e. they become specialised in one or more functions. To protect their genetic uniformity, many colonial animals possess a genetically based allorecognition system, allowing them to discriminate between self and non-self with a high degree of precision. It is thought that allorecognition evolved to prevent germ cell parasitism between conspecifics ^10^. Allorecognition genes have been identified in cnidarians and ascidians, and shown to be phylogenetically unrelated to each other and to vertebrate MHC/T-cell-mediated allorecognition ^11,12^. Finally, colonial growth has evolved and been lost multiple times, even within a single phylum ^2^. Despite the many interesting features of animal colonies, the cellular and molecular mechanisms that govern them remain unknown.

Members of the genus *Hydractinia* ^13^ form colonies that grow on gastropod shells inhabited by hermit crabs. A single, sexually produced larva colonises the shell and metamorphoses to generate a primary zooid (called polyp) and gastrovascular tubes that are attached to the substratum and known as stolons. Stolons develop into a dense network that continuously buds new, genetically identical but polymorphic polyps. A colony typically possesses several polyp types, each specialised in specific function such as prey capture (feeding polyp or gastrozooid), production of gametes (sexual polyps or gonozooid), or defence (dactylozooids). This growth process relies on a lineage of adult, migratory pluripotent stem cells known as i-cells ^5,9^. *Hydractinia symbiolongicarpus* (hereafter referred to as *Hydractinia*) is an established model organism. It is amenable to genetic manipulation, has a sequenced genome ^14^, multiple transcriptomic resources ^15^, and is easy to culture in the lab on glass slides that contain gastrozooids, gonozooids, and a stolonal network. Despite all cells being continuously derived from i-cells, it is unknown if different colony parts are made of the same cell types or if specific types are the building blocks of each specific colony component.

Recently, single cell transcriptomics (scRNA-seq) has provided an opportunity to classify individual cells into distinct cell types by analysing their unique gene expression profiles and clustering them based on shared transcriptional patterns ^16^. This approach offers unprecedented insights into cellular heterogeneity and complex biological systems ^17,18^. Using single cell transcriptomics, the cell type atlases of numerous organisms, including several solitary and colonial cnidarians ^19–29^ have been profiled. Single cell transcriptomics can reveal the cellular composition of the different parts of a colony by using part-specific libraries. However, previous single cell studies on colonial cnidarians have been constructed from mixed-cell libraries and could not profile colony parts individually.

Here, we characterised a single cell atlas of the individual colony parts of *Hydractinia*. We generated ∼200K cellular profiles from distinct samples including feeding- and sexual polyps, and stolons. We characterised all major cell types and the transcription factors (TFs) and gene networks that are expressed in them. Furthermore, we quantified their differential abundances in colony parts, revealing that most cell types are shared, albeit in different proportions, between distinct colony parts. Part-specific cell types were rare, but might have also played a role in the evolution of coloniality. Among the part-specific cell types, we uncovered one that is enriched in stolons and expresses biomineralization and chitin synthesis enzymes. We studied the genetic profile of that cell type, revealing a cluster of repeat-containing *Shematrin-like* genes that resemble molluscan shell matrix proteins, raising the possibility that *Hydractinia* has co-opted a biomineralization gene programme to attach to the gastropod shells inhabited by hermit crabs, a key adaptation to their environment. Altogether, the *Hydractinia* cell atlas reveals the cellular and molecular principles of their colonial growth.

### A cell type atlas of *Hydractinia*

We obtained cell suspensions of laboratory-grown *Hydractinia* colonies using ACME ^30^ with modifications (see Methods). We included mixed polyp samples, selected feeding polyps (gastrozooids) and reproductive polyps (gonozooids), as well as stolons (Figure 1A). We then performed a total of 4 SPLiT-seq experiments, comprising 13 sublibraries, and sequenced these libraries using Illumina NovaSeq 6000 technology at 150bp paired-end read length (Figure 1B, Supplementary File 1). We obtained a total of 6.9 billion reads.

**Figure 1:**
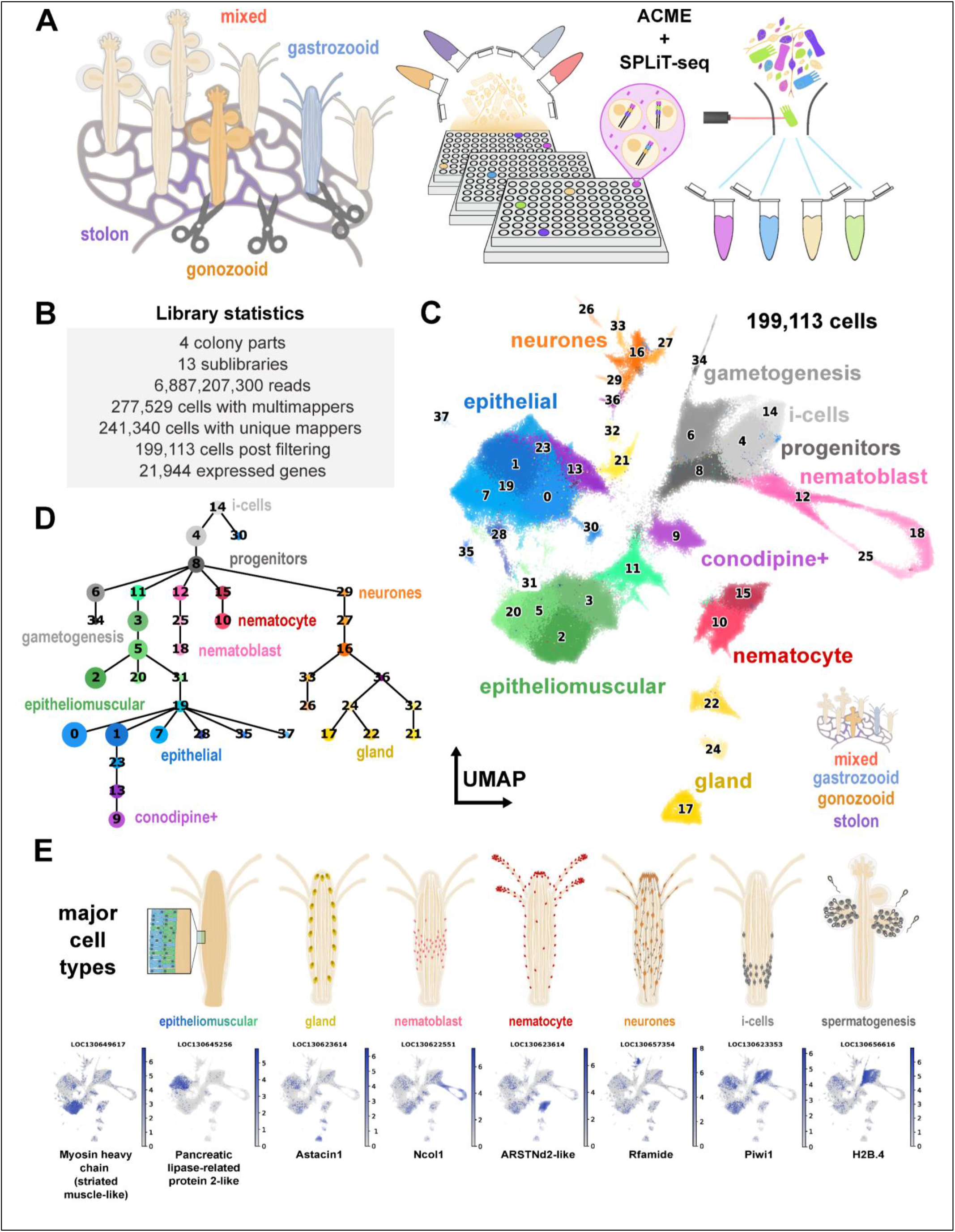
Single cell atlas of the colonial cnidarian *Hydractinia symbiolongicarpus*. **A**: Scheme of experimental design to collect and process the samples. **B**: UMAP projection of the annotated atlas with clusters coloured according to their cell type identity. **C**: General library statistics of the whole dataset. **D**: Abstracted graph of lineage reconstruction (PAGA) showing most probable relationships between clusters. Each node corresponds to an annotated cluster. Size of the nodes is representative of the amount of cells in the cluster, thickness of the edges is proportional to the connectivity probabilities.

We then used the *Hydractinia symbiologicarpus* chromosome-level genome assembly ^14^ to map these reads using our SPLiT-seq read processing pipeline ^30^. We further annotated the gene models using DIAMOND ^31^ (Supplementary Dataset). We used Muon ^32^ to build a multimodal matrix with uniquely mapped reads and with reads mapping to many loci, but used the matrix containing only singly mapping reads for the subsequent single cell analysis. After further filtering steps, we obtained a matrix containing 199,113 cells. We used Harmony ^33^ to leverage batch effects from the different experiments, and built a kNN graph using 40 neighbours and 75 principal components.

As previously reported, SPLiT-seq delivers low numbers of reads and genes per cell ^30,34–37^ (Supplementary Figure 1A-B). In spite of this, and thanks to the high number of cell profiles that can be obtained using SPLiT-seq, the Leiden algorithm for cell clustering at a resolution of 1.5 yielded a robust identification of 53 distinct cell clusters (Figure 1C, Supplementary Figure 1B-C). We generated marker genes for each cluster using both the Wilcoxon and Logistic Regression methods (Supplementary Files 2-4). We used PAGA to group these clusters in broad types (Figure 1D). Additionally, we performed a co-occurrence analysis on cell type clusters ^25^. This analysis corroborated our clustering results (Supplementary Figure 1F), and helped with cluster annotation together with gene marker examination of previously published *Hydractinia* and cnidarian cell type literature (Supplementary Note 1).

The *Hydractinia* cell type repertoire included i-cells, the pluripotent stem cells that drive colonial growth and regeneration, committed progenitors, germ cells, as well as a panoply of differentiated cell types (Figure 1E, Supplementary Note 1). Similar to other cnidarian cell atlases ^22–26,28,29^, the most abundant cell types were neurones, nematoblasts, nematocytes, epithelial and epitheliomuscular cells. We also identified a set of mucosal and digestive gland cells, as well as a novel cluster of *Conodipine*+ cells, marked by the expression of venomous proteins. Altogether, these results show that our single cell data resolves the major cell types of *Hydractinia* polyps and stolons.

### Cellular composition of *Hydractinia* colony parts

We then aimed to ascertain the cellular composition of each individual colony part. We examined the contribution of each sample to our integrated analysis, revealing key differences in their cellular composition (Figure 2A). In stolons, we observed markedly fewer gland cells, neurones, and germ cells; the latter were abundant in sexual polyps as expected. Furthermore, sexual polyps exhibited very low counts of nematocytes and gland cells, which were prevalent in the feeding polyps. These observations too were consistent with our initial expectations.

**Figure 2:**
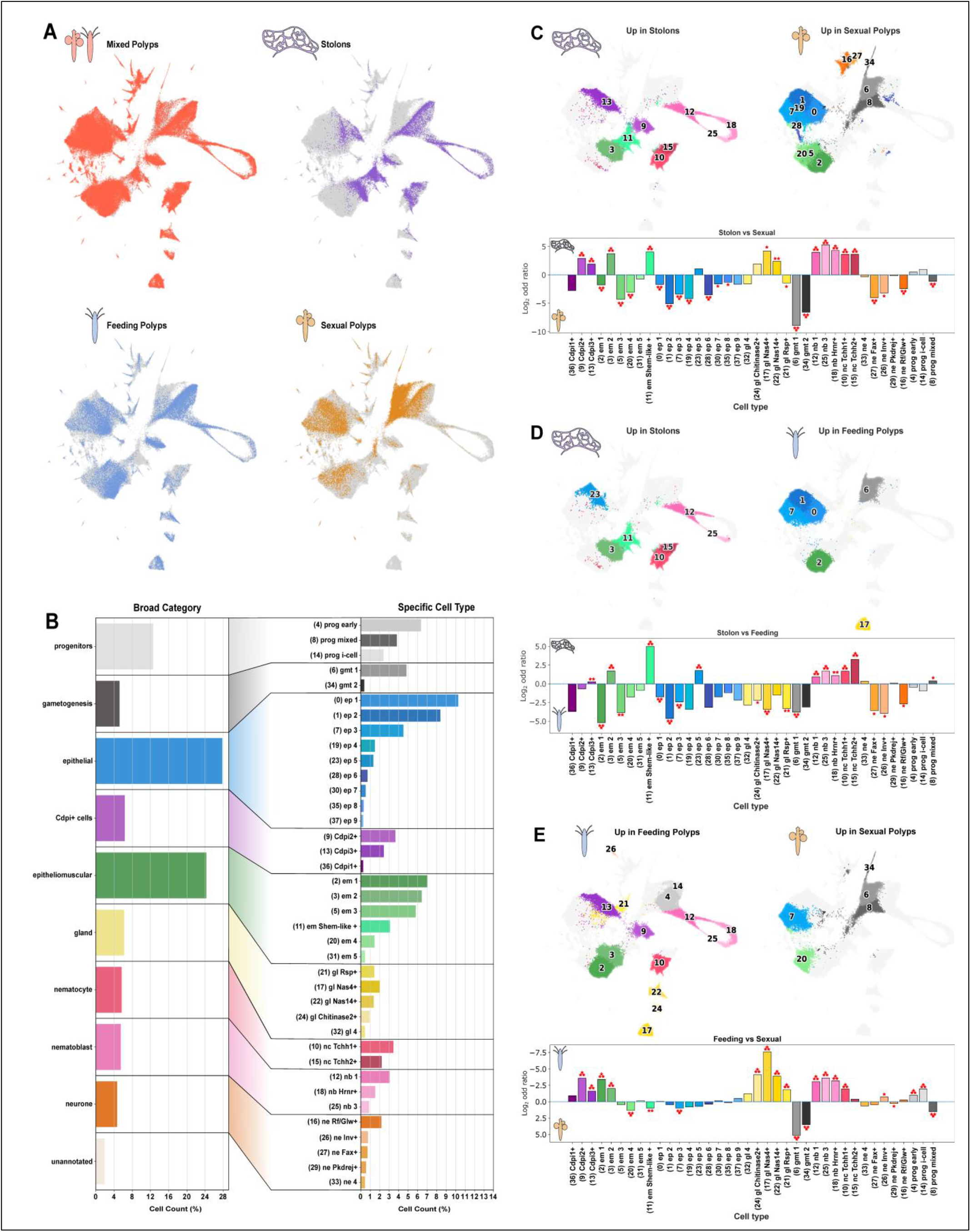
Cell proportions of colony parts across the single cell atlas. **A**: UMAPs showing presence of the different colony parts in the dataset (Mixed Polyps, Stolons, Feeding and Sexual Polyps). **B**: Barplots showing the cell count percentage at the broad cell category and specific cell cluster levels. **C**: Compositional comparison between stolon cells and cells coming from sexual polyps. **D**: Compositional comparison between cells coming from stolon samples and feeding polyp cells. **E**: Compositional comparison between cells sampled from sexual and feeding polyps. On the top half area of each sub item are the clusters enriched when the FDR was higher than 0.0005 in that respective condition (left or right). On the bottom half of each sub item are the calculated logarithms of the odd ratios of enrichment output from the fisher test for each cluster. Each asterisk present on top of the barplot translates to a level of significance under different FDR (*: 0.05, **: 0.005, ***: 0.0005).

To statistically analyse these observations, we first calculated the aggregated cell counts for each broad cell type category and individual cell type (Figure 2B, Supplementary File 5). We then calculated log2 odd ratios comparing feeding, sexual polyps and stolons (Figure 2C-E). Finally, we used scCODA, a Bayesian model, to analyse compositional changes in single cell data (Figure 2C-E) to detect credible changes. We leveraged distinct False Discovery Rates (FDRs,*: 0.05,**: 0.005,***: 0.0005) to gain an understanding of the different credible intervals within our results.

We first compared stolons to sexual polyps (Figure 2C). Top credible effects (FDR 0.0005: ***) with high log2 odd ratios in stolons included nematocytes (10, 15) and nematoblasts (12, 18, 25), consistent with a lower number of nematocytes in sexual polyps. These top credible effects also contained two epitheliomuscular clusters (3, 11) and two *Conodipine*+ clusters (9, 13). Conversely, top credible effects in sexual polyps included the germ cell clusters (6, 34), a mixed progenitor cluster (8), a range of epithelial (0,1,7,19,28), epitheliomuscular (2, 5, 20), and neuronal (16, 27) cell clusters.

We then compared stolons to feeding polyps (Figure 2D). Top credible effects with high log2 odd ratios in stolons included nematocytes (10, 15) and nematoblasts (12), indicating a higher frequency of nematocytes in stolons when compared to feeding polyps. These top credible effects also included an epithelial cluster (23) and two epitheliomuscular clusters (3, 11). Cluster 11 had the highest log2 odd ratio of this comparison, and was also highly enriched in stolons vs sexual polyps, indicating its high specificity to stolons. We termed this cluster *Shematrin-like*+ cells due to the high expression of LOC130630016, which was annotated as a *Shematrin-like* protein (Supplementary Files 2-4). In addition, another marker of this cluster was LOC130630014, annotated as *Prisilkin-39-like*. Interestingly, *Shematrins* and *Prisilkins* are glycine-rich repeat-containing proteins found in molluscan shells ^38,39^. We then examined the top credible effects with high odd ratios in feeding polyps. These included mixed progenitors (6), epithelial clusters (0, 1, 7), and an epitheliomuscular cluster (2). These epithelial and epitheliomuscular clusters were also credibly enriched in sexual polyps vs stolons and therefore are highly specific of polyps.

Finally, we compared feeding to sexual polyps (Figure 2E). Top credible effects with high log2 odd ratios in feeding polyps included a range of gland clusters (17,21,22, 24), nematocytes (10), nematoblasts (12, 18, 25), *Conodipine*+ cells (9, 13), and epitheliomuscular clusters (2, 3). Conversely, top credible effects in sexual polyps included the germ cell cluster (6, 34), a mixed progenitor cluster (8), an epithelial cluster (7), and an epitheliomuscular cluster (20).

Altogether, our results revealed the cellular composition of the main *Hydractinia* colony part. Sexual polyps are made of epithelial and epitheliomuscular cell types, neurones, and germ cells, and lower in gland cells and nematocytes. Feeding polyps contain abundant epithelial, epitheliomuscular, nematocyte, gland, *Conodipine*+, and neuronal cells. Stolons possess fewer neuronal and gland cells, but have abundant nematocytes and *Conodipine*+ cells. Finally, our data show that an epitheliomuscular cell type is highly specific to stolons. In summary, we find that colony parts seem to be defined by a combination of specific cell types and by different proportions of shared cell types.

### The transcriptional landscape of *Hydractinia* cell types

We then investigated the gene modules that underlie cell type differentiation per cell type and colony part. We used Weighted Gene Coexpression Network Analysis (WGCNA) ^40^ to identify genes with correlated expression patterns. WGCNA calculates an Adjacency and Topological Overlap Matrix (TOM) that captures the interconnectedness and co-expression patterns among genes, providing insights into modular structures and regulatory relationships within biological networks. Unlike marker finding algorithms, this approach can detect gene sets that are expressed in one single cell type or coordinately expressed in several. We identified 7,547 genes distributed over 38 modules of gene coexpression, corresponding to individual cell clusters and broad groups (Figure 3A, Supplementary Dataset, Supplementary Figure 2A-B). We used Gene Ontology (GO) analysis to extract biologically relevant terms for each cell type (Figure 3B, Supplementary File 6). Among broadly expressed gene modules we found a module expressed broadly in epithelial cells, enriched in GO terms such as “cell junction assembly” (Figure 3B). We also found a module expressed broadly in epitheliomuscular cells, enriched in GO terms such as “smooth muscle cell differentiation” and “digestive tract morphogenesis” (Figure 3B), consistent with the dual muscular and digestive epithelium nature of these cells. In addition, we found gene modules enriched in i-cells and mixed progenitors, enriched in GO terms related to DNA synthesis and replication as well as chromatin organisation (Figure 3B). Interestingly, a module expressed in *Conodipine*+ cells was enriched in biosynthetic and metabolic genes, consistent with a role in venom synthesis. Finally, we found a module expressed in *Shematrin-like*+ cells, enriched in GO terms such as “chitin metabolic process”. The genes annotated with these GOs include chitin synthase (LOC130613296) as well as chitinase (LOC130641097) and chitin deacetylase enzymes (LOC130614146), consistent with the chitinous nature of *Hydractinia* stolons ^41–43^. Our data shows that *Shematrin-like+* cells express chitin biosynthetic genes and are therefore key players in stolon chitinization.

**Figure 3:**
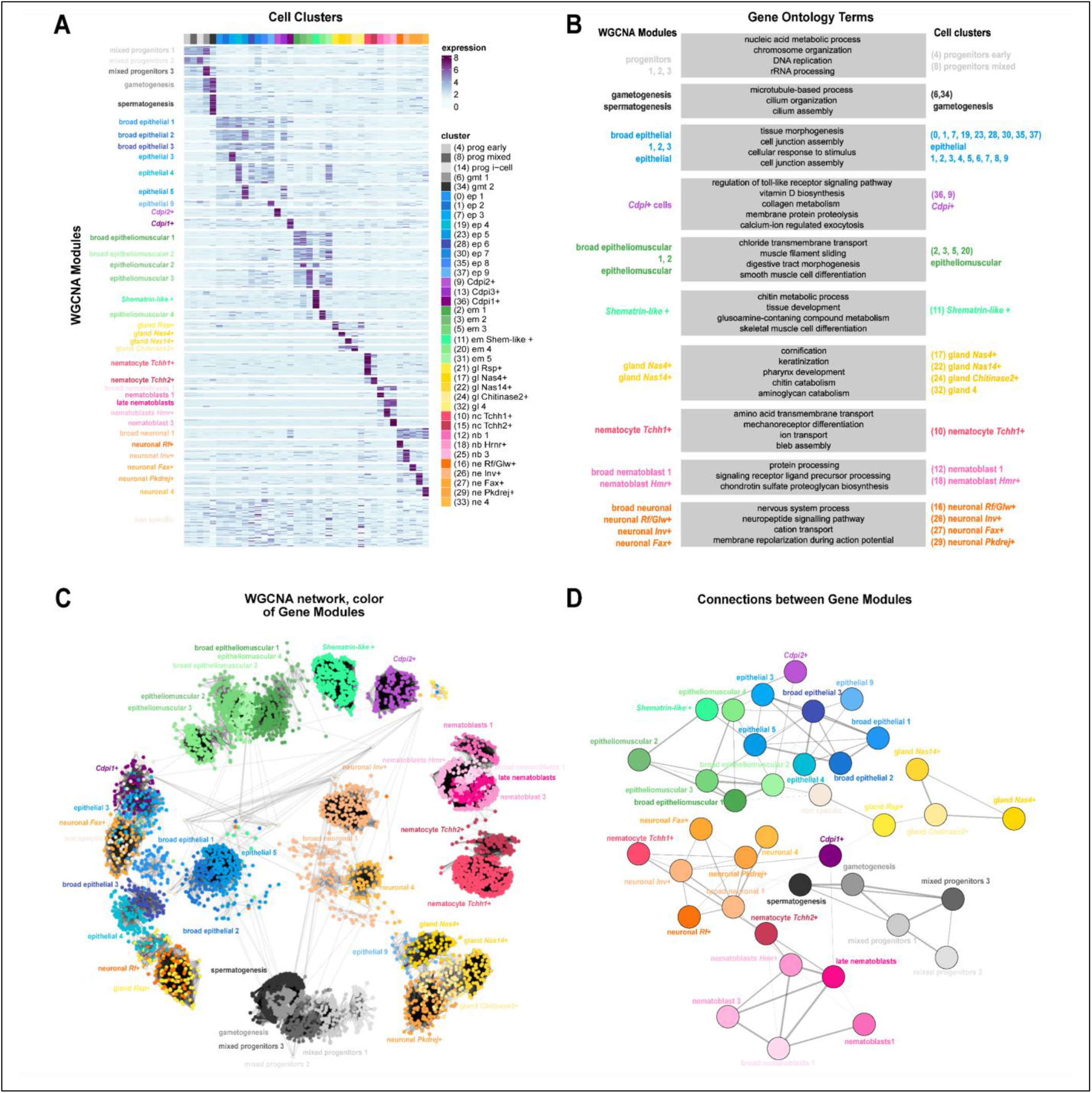
Regulatory landscape of whole colonies in *Hydractinia*. **A**: Expression heatmap of 7547 genes over 38 modules of coexpression (rows) sorted by annotated cell type (columns). Colour intensity represents normalised expression. **B**: Summarised gene ontology terms enriched in each module (left) associated with the respective cell cluster (right). **C**: Network visualisation of WGCNA modules using the Kamida-Kawai layout algorithm. Each dot represents a gene and the respective colours are the cluster of highest expression of a determined gene. Edges are a representation of co-expression values. **D**: Module visualisation of the gene network using the large connected graphs layout algorithm, where co-expression values are summarised between different modules, showing the associations between them.

We then aimed to elucidate the similarities between cell type expression modules to detect more subtle patterns of gene co-regulation between different cell types. One key advantage of WGCNA over marker finding algorithms is that connections between modules can be derived from the structure of the underlying graph. To do this, we visualised the WGCNA TOM matrix as a network, and identified several graph connected components that match the WGCNA modules (Figure 3C). This analysis shows that connected components and WGCNA modules largely overlap, i.e. genes within a module are mostly connected amongst themselves but also have connections with other modules that reveal the similarities between these genetic programmes. To visualise this, we plotted a network where the nodes represent each gene module and the edges represent the number of co-expressed genes from different pairs of modules (Figure 3D). This analysis recovered known relationships such as the connection between nematoblasts, nematocytes, and neurones ^20,44–47^, or the similarity between germ cells, mixed progenitors and i-cells ^48^. Furthermore, this analysis shows that *Conodipine*+ cells have mixed patterns, with the *Cdpi1*+ module connected to neurones, and the *Cdpi2*+ module connected to both epithelial and epitheliomuscular cell modules. Importantly, this analysis showed that the stolon-enriched *Shematrin-like*+ module was connected to other epitheliomuscular clusters, corroborating their epitheliomuscular identity, but also revealing its connection with the *Cdpi2+* module. This analysis shows that these cell types have genes that are co-expressed and suggests a functional or spatial relationship between them. Altogether, these analyses reveal the transcriptional patterns of each *Hydractinia* cell type and their similarities, providing a foundation for future functional studies.

### Transcription factor regulatory profiles in *Hydractinia*

We then aimed to identify TFs that may be driving the differentiation of *Hydractinia* cell type diversity. We annotated 801 TFs and identified a set of 69 of them with cell type-specific expression (Figure 4A, Supplementary Dataset). We then exploited our WGCNA TOM graph approach to obtain centrality values for each TF in each module. In essence, TFs with a high centrality within a module are connected to many genes within the module and therefore are highly correlated with them and candidate regulators. We plotted the top central TFs for each module (Figure 4B-C), which included several TFs previously described. Among the top central TFs we found *Sox9-like* (broad epithelial), ETS-related (epitheliomuscular and gland cells), *Forkhead box protein N4-like* (nematoblast and nematocytes), and Photoreceptor-specific nuclear receptor-like (neurones).

**Figure 4:**
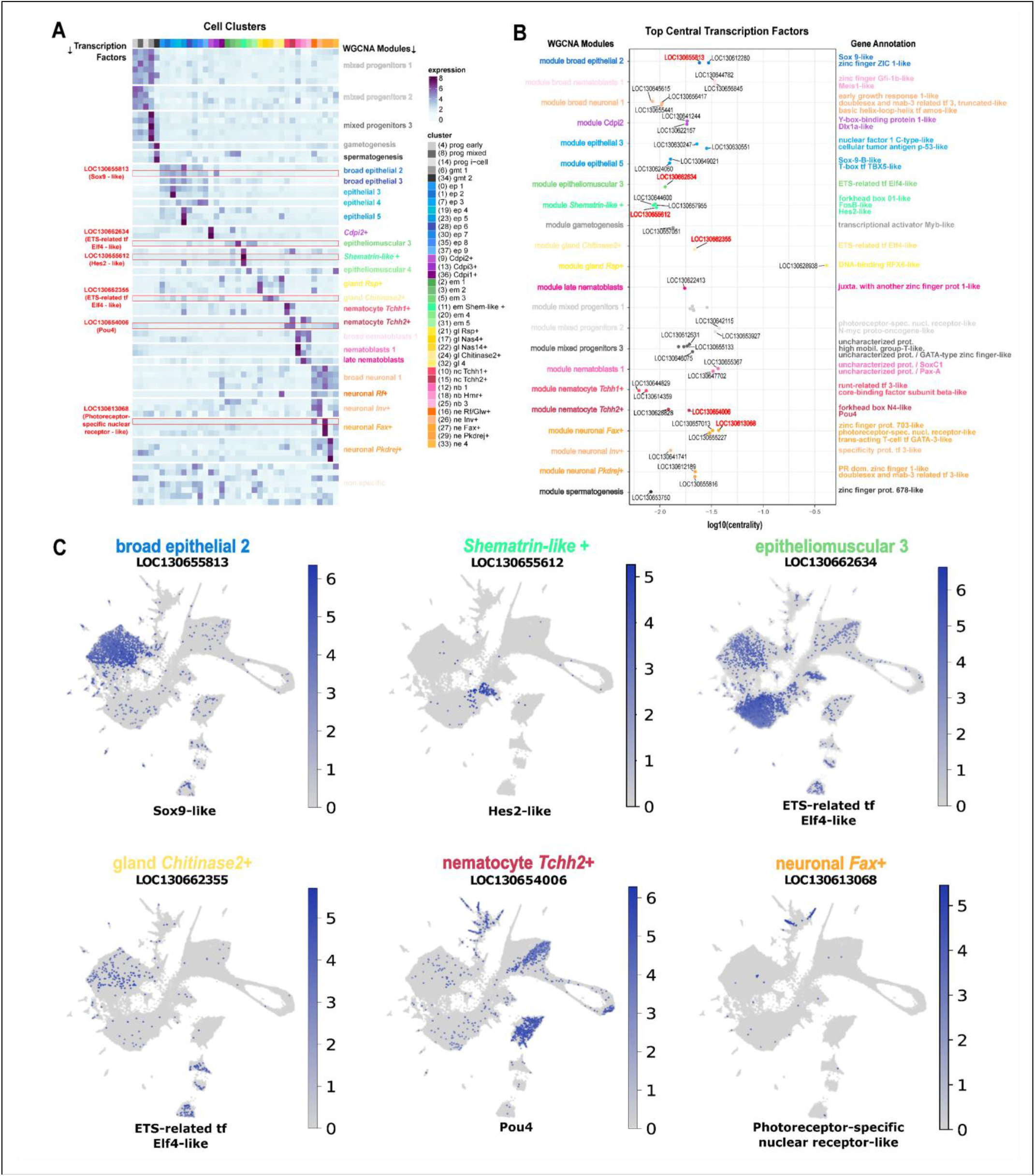
Transcription factor expression landscape across whole colonies of *Hydractinia*. **A**: Expression heatmap of the transcription factors found in the output of WGCNA over 38 modules of coexpression (rows) sorted by annotated cell type (columns), highlighting in red the factors selected shown in C. **B**: Stripplot graphing the obtained top central transcription factors in WGCNA modules with their respective annotations, highlighting the selected factors in red. **C**: Feature expression plots of selected top central transcription factors.

One transcription factor identified as top central, to the 26_nematocyte_Tchh2+ module, was a POU domain, class 4 transcription factor (Supplementary Figure 3). In mammals, POU4 is vital to proper development of mechanosensory cells and neurones in the inner ear and surrounding structures ^49^. POU4’s role in the development of mechanosensory neurones is consistent across metazoans ^50^. In *Nematostella*, *NvPOU4* RNA *in situ* hybridization predominantly marks late differentiating stinging cells (nematocytes) at the distal end of the tentacles, along with some differentiating neurones in the body column. *NvPOU4* -/- homozygote knockout mutants fail to generate terminally differentiated nematocytes ^51^. They produce nematoblasts devoid of a capsule or stinging apparatus (harpoon), but that are still marked by *NvNCol3* (a marker of early nematogenesis). *NvPOU4* has been proposed to have an ancestral function in mechanosensory cell development and appears in cnidarians to have a dual role, acting also in production of mechanosensory machinery in the cnidarian-specific stinging cell type, cnidocytes ^51,52^. The UMAP expression plot of *Hydractinia Pou4* is consistent with this dual role, with expression in both the neural and nematocyte lineages (Figure 4C). The co-expression of *Pou4* and *Trichohyalin* in the *Tchh1* and *2*+ nematocyte types suggests a potential co-opting of gene modules traditionally associated with mechanosensory cells and follicle development for the production of the unique mechanosensory machinery found in nematocysts.

Our analysis also unveiled TFs that are expressed in other cell types characterised in this study. For instance, top central factors in the *Shematrin-like*+ module include a *Forkhead box protein O1-like*, a *FosB-like* and a *Hes2-like* TFs that are highly specific to this stolon enriched cell type. Collectively, these results reveal the TFs that are specific to *Hydractinia* cell types and will open numerous research avenues to investigate their role in cell type differentiation and evolution.

### *Shematrin-like*+ cells express a cluster of molluscan-like repeat-containing genes and a biomineralization gene programme

We then aimed to further investigate the nature of *Shematrin-like*+ cells. Our previous results show that these epitheliomuscular cells are highly enriched in stolons (Figure 2), express genes similar to molluscan shell matrix protein coding genes *Shematrin* and *Prisilkin* (Supplementary Files 3-4) as well as chitin synthesis genes (Figure 3). To gain further insight into *Shematrin-*like genes we analysed their genomic locations, taking advantage of the newly assembled chromosome scale *Hydractinia* genome ^14^. Interestingly, both genes annotated as *Shematrin* and *Prisilkin* were located in the same genomic region in chromosome 2 (Figure 5A). We observed that several other genes surrounding this genomic location also had similar repeats, indicating the presence of a genomic cluster likely originated by tandem duplications (Figure 5A). Most of these genes had substantial specific expression in the *Shematrin-like*+ cell cluster (Figure 5B).

**Figure 5:**
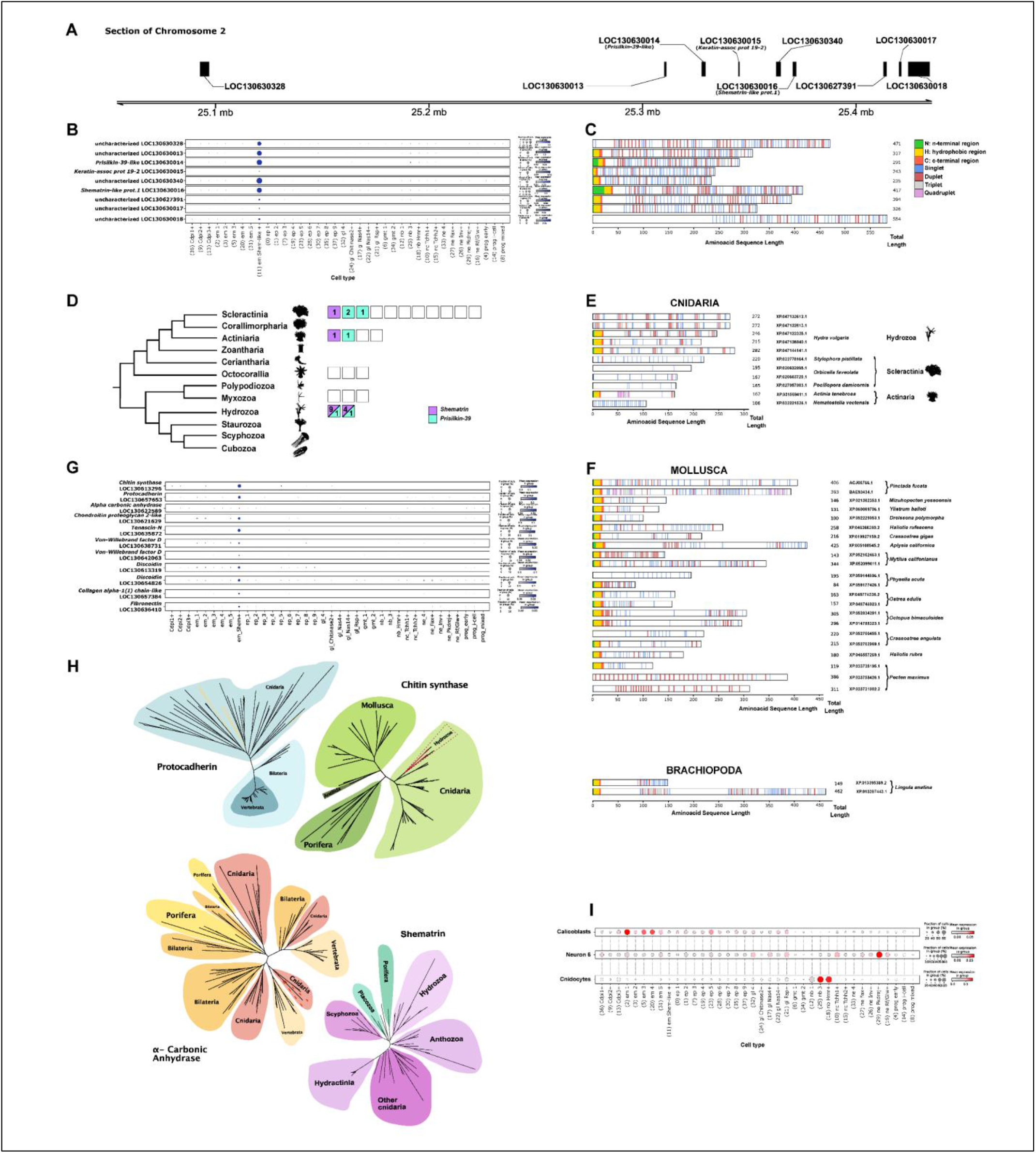
Architecture and phylogenetic relationships of genes primarily expressed in *Shematrin-like*+ cells. **A**: Region on the chromosome 2 of *Hydractinia* depicting the genomic location of *prisilkin-39* and one annotated *Shematrin-like* gene. **B**: Dot plot showing the individual expression levels of genes specific to the *Shematrin-like*+ cluster in the same genomic location ranges of A. **C**: Signal peptide regions (N: n-terminal region, H: hydrophobic region, C: c-terminal region) and glycine repetitions (singlet, duplet, triplet, quadruplet) across the complete amino acid sequences of all genes in B. (tomato red; C, lime green; N, gold; H, cornflower blue; singlets, indian red; duplets, light grey; triplets, plum; quadruplets). **D**: Phylogenetic relationships in cnidaria adapted from De Biasse *et al*. depicting the number of annotated proteomes in NCBI (empty blocks), the number of annotated *Shematrin-like* (Lilac blocks) and *prisilkin-39* proteins (Turquoise blocks). The clade silhouettes were obtained from www.phylopic.org and are either licensed under Creative Commons Attribution 3.0 Unported or are available under public domain. **E**: Signal peptide regions (N: n-terminal region, H: hydrophobic region, C: c-terminal region) and glycine repetitions (singlet, duplet, triplet, quadruplet) across the complete amino acid sequences of all *Shematrin-like* and *Prisilkin-39-like* annotated proteins in cnidarians. **F**: Signal peptide regions (N: n-terminal region, H: hydrophobic region, C: c-terminal region) and glycine repetitions (singlet, duplet, triplet, quadruplet) across the complete amino acid sequences of a selection of *Shematrin-like* and *Prisilkin-39-like* annotated proteins in molluscs. **G**: Dot plots showing the individual expression levels of genes found *Hydractinia* related to biomineralization pathways. **H**: Unrooted phylogenetic trees depicting relationships of genes that are involved in biomineralization pathways. **I**: Selection of dot plots showing the scored expression of one to one ortholog genes present in annotated clusters of *Stylophora pistillata* with a minimum fold change of 1 (Calicoblasts, Neurone 6 and Cnidocytes).

To understand their sequence similarities, we performed a series of preliminary BLAST analyses which failed to provide robust results due to the highly repetitive nature of these sequences. We also aimed to obtain gene alignments of the *Hydractinia Shematrin-like+* cluster genes and other genes annotated as *Shematrins*, which resulted in poor alignment and support values. Finally, to structurally characterise these genes we devised a repeat finder algorithm and used SignalP 6.0 ^53^ to predict signal peptides. Our repeat finding algorithm searches for repeats with a XG_n_X (where X represents amino acids I, L, V, Y) structure where n can have a value between 1 and 4 (singlets, duplets, triplets, quadruplets), typical of molluscan *Shematrins* ^39^. *Hydractinia Shematrin-like+* cluster genes all had a majority of singlet and duplet repeats, and 7 out of 9 had a predicted signal peptide (Figure 5C). We found another gene cluster in a nearby genomic location containing 8 further repeat-containing genes with a very similar structure in chromosome 2 and two more genes very close together in chromosome 3 (Supplementary Figure 4).

We then wondered if other *Shematrin*-like genes are present in other cnidarian species. Since these are difficult to identify by BLAST algorithms, but are sometimes annotated as “*Prisilkin*” and “*Shematrin*” in NCBI annotated genomes, we searched currently available cnidarian genomes for these annotations. This analysis uncovered annotated *Shematrins* or *Prisilkins* in 7 out of 23 species with annotated genomes ^54^ (Figure 5D). *Hydra vulgaris, Stylophora pistillata*, *Orbicella faveolata*, *Pocillopora damicornis*, *Actinia tenebrosa* and *Nematostella vectensis* possess *Shematrin* genes with similar repeats (Figure 5E)

To compare these results to molluscan *Shematrin* genes, we detected repeats using the same algorithm in 22 molluscan *Shematrins* (Figure 5F). We included 2 *Shematrins* from Brachiopoda for reference (Figure 5F). This analysis revealed that their structure is very similar, with a signal peptide followed by repeats with a singlet and duplet majority. Altogether, these analyses show that *Hydractinia* stolon enriched *Shematrin-like+* cells express a genomic cluster of *Shematrin-like* genes similar to molluscan *Shematrins*. These genes are present in other related cnidarians but their wider presence in the cnidarian phylum cannot be identified with current annotation methods.

As *Shematrins* have been characterised as molluscan shell proteins and are involved in biomineralization in other organisms too, we searched for the expression of other bona-fide biomineralization genes. We identified 11 genes involved in biomineralization with high expression or enriched expression in *Shematrin-like*+ cells (Figure 5G). We confirmed that this gene and other genes are bona fide *Hydractinia* homologues by a phylogenomic approach (Figure 5H, Supplementary Figure 5). These analyses confirmed that *Hydractinia Shematrin*-*like*+ cells express a biomineralization gene programme. Taken together, we discovered a stolon-enriched cell type that expresses biomineralization genes, including *Shematrin-like* genes akin to genes involved in molluscan shell formation as well as chitin synthesis genes. Similar cell types and genes have not been found in other cnidarians, raising the possibility that they are a colonial cnidarian cell type innovation.

Finally, we aimed to elucidate the evolutionary relationship between *Shematrin-like*+ cells and other cnidarian types involved in biomineralization. Recently, the cell type atlas of the stony coral *Stylophora pistillata* has revealed that these corals have calicoblasts, a type of epidermal cells that express carbonic anhydrases among other enzymes ^25,55^. To elucidate if *Stylophora* calicoblasts are similar at the transcriptomic level to *Hydractinia Shematrin-like*+ cells we obtained the top markers of calicoblasts, identified one-to-one orthologs in *Hydractinia*, and scored their expression in our single cell dataset. These scores were high in several epitheliomuscular and epithelial cell types, but not in *Shematrin-like*+ cells (Figure 5I). As a comparison, we performed the same analyses using markers of *Stylophora* neuronal types and cnidocytes. These scores were indeed high in *Hydractinia* neuronal types and nematoblasts, evidencing the transcriptomic similarity of these types, likely originated by an evolutionary homology relationship. In contrast, calicoblasts and *Shematrin-like*+ cells are not transcriptionally similar and therefore have likely originated as two independent cell type innovation events in the evolution of the *Stylophora* and *Hydractinia* lineages.

## Discussion

Our *Hydractinia* cell atlas reveals the cellular composition of distinct colony parts of a colonial cnidarian. Previous cnidarian cell atlases have profiled solitary species such as *Hydra* and *Nematostella* or considered colonial cnidarians as a whole ^19–29,56^. We show that most cell types are present in all parts of the colony, albeit in different proportions, with fewer cell types being part-specific. Therefore, it appears that a cell ‘mix-and-match’ approach has been the major mechanism in the evolution of new colony parts in *Hydractinia*. In a broader sense, using different combinations of a similar cellular repertoire may have facilitated the multiple gains and losses of characters, such as medusa stage and coloniality, in hydrozoan evolution ^2,57^ and could be a common mechanism of colony evolution across Metazoa. We demonstrate that single-cell methods are ideal for studying cellular composition in colonial animals. Future studies will apply these methods to other colonial animals, such as colonial ascidians and bryozoans.

We annotated 53 cell clusters and validated them through several *in silico* methodologies. The atlas shows known cell types that are found in several single-cell atlases, such as nematocytes, epithelia, and neurones as well as multiple *Hydractinia*-specific novel cell types. We analysed gene sets with expression signatures that can be correlated mostly one to one to the obtained cell types and used them to define regulatory profiles from 38 gene coexpression modules and transcription factors. This fine-grained characterization of signalling pathways controlling colony homeostatic processes, allows us to understand better molecular mechanisms and gene regulatory networks underlying evolutionary patterns of colony plasticity.

*Hydractinia* has pluripotent stem cells which can give rise to all somatic cells and to gametes that have been confirmed experimentally at single-cell resolution ^9^. Our cell atlas and the PAGA analysis computationally confirm what has been known experimentally. The congruence between the two approaches provides confidence to our results and places this model organism and its cell atlas in a unique position to study pluripotency and lineage commitment *in vivo* in a fully genetically tractable system. The PAGA analysis recapitulated the *Hydractinia* cell lineages’ derivation from i-cells; however, Cluster 30, a sub-cluster of epithelial cells, appears to derive directly from i-cells rather than from the larger epithelial cluster as could be expected. This could be explained by the fact that most polyps from which the atlas was generated were fully grown, sampled from the non-growing middle of the colony rather than from the growing edges. Since epithelial turnover in *Hydractinia* is slow ^41^, committed epithelial progenitors would be rare in our sample, making their precise position in the PAGA cladogram difficult to determine.

Our analysis reveals new cnidarian cell types, such as the *Conodipine*+ cells that share many transcriptomic signatures with nematocytes, neurones, and epithelial cells. These cells express genes encoding for venom such as membrane protein proteolysis, and at least one homolog of the Membrane Attack Complex (MAC) and perforin family, a group of proteins studied before for being recruited into venom-injecting cells in cnidarians ^58^. However, they lack the nematocyst apparatus. Venom-expressing cells that are not nematocytes are also present in *Nematostella* ^59^. Further studies are required to fully confirm the identity and function of this broad cell type in *Hydractinia* colonies.

Cnidarian nematocytes are a unique cell type across the metazoan tree but appear to have co-opted gene programs and transcription factors used for specification of mechanosensory cells in other organisms. Previous work in *Nematostella* highlighted a role for POU4 in the terminal differentiation of nematocytes ^51^. Here we demonstrate nematocyte specific expression of a bona fide *Pou4* homologue, along with co-expression of *Trichohyalin1* and *2*, known markers of hair follicle development in other organisms ^60^ and venom genes. As such, it appears that nematocytes, although a unique cell type, do not rely on a unique gene program for their function; rather that they employ a unique combination of common gene programs, shared with other organisms.

*Hydractinia* and other closely related cnidarians have adapted to form colonies on molluscan shells inhabited by hermit crabs. *Hydractinia* colonies possess chitinous stolons ^41–43^ and form a crystalline mat with calcium carbonate ^61^, though the cellular and molecular basis for this process was previously unknown. Our data show that *Hydractinia* has a biomineralizing cell type that expresses enzymes for both chitin synthesis and biomineralization. This cell type is not transcriptionally similar to the recently described calicoblasts of stony corals ^25,62^, suggesting that they may have arisen independently in the evolution of cnidarians.

Cnidarian skeletons have been morphologically classified as calcareous, corneous, or coriaceous, with an additional type, bilayered, described later ^63^. Our data reveal that *Hydractinia Shematrin-*like+ cells are responsible for chitin and biomineralization, likely forming the basis for coriaceous biomineralization. Stony corals, which have calcareous skeletons, possess calicoblasts that are likely not homologous to *Shematrin-like*+ cells. Along with the recent finding that biomineralization in stony corals and octocorals arose independently ^64,65^, our data support a scenario where biomineralization evolved independently in several cnidarian lineages. Therefore, we propose that cell type innovation happens at a low frequency when compared to the innovation of colony parts using preexisting cell types. Nonetheless, cell type innovation might have played important roles in the evolution of coloniality in cnidarians. In the future, single-cell techniques will enable the identification of additional biomineralizing cell types and other cell type innovations, allowing further investigation into the cellular and molecular basis of adaptation.

## Data availability

The datasets supporting the conclusions of this article are available in: GEO (scRNA-seq reads): GSE269914

## Code availability

Code: https://github.com/scbe-lab/hydractinia_sc_atlas

## Acknowledgments

Research at the Solana lab at Oxford Brookes University and at the Living Systems Institute is supported by MRC grants (MR/S007849/1 and MR/W017539/1), a BBSRC Grant (BB/V014447/1) and a Leverhulme Trust grant (RPG-2019-332 and RPG-2023-330) to JS. HG-C and EE were supported by Nigel Groome studentships from Oxford Brookes University. Work in the Frank lab is funded by a Wellcome Trust Investigator Award in Science (grant no. 210722/Z/18/Z). HRH was a doctoral student at the Science Foundation Ireland Centre for Research Training in Genomic Data Science (grant no. 18/CRT/6214). MS-S was a Human Frontier Science Program Long-Term Postdoctoral Fellow (grant no. LT000756/2020-L). MER is funded by NERC DTP GW4+, MAP is supported by a fellowship from the Fundación General CSIĆs ComFuturo programme funded from the European Union’s Horizon 2020 research and innovation programme under the Marie Skłodowska-Curie grant agreement No. 101034263; and MAP and JP are supported by the Wellcome Trust (210101/Z/18/Z) and the School of Biological Sciences (University of Bristol). Flow cytometry was performed at the Sir William Dunn School of Pathology Flow Cytometry Facility, University of Oxford with the assistance of Dr Robert Hedley and University of Galway Flow Cytometry Core Facility with the assistance of Dr Shirley Hanley. The authors would like to extend thanks to all members of the Solana, Frank and Paps labs, especially Maria Roselló, Camille Curantz, Paris Kovi Weavers and Yasmine Lund-Ricard, for useful discussion, input and assistance. Finally, to Amy Duclaux and Laura Ryan for animal care and culturing.

## Authors’ Contributions

D.A.S.-D, H.R.H., H.G.-C., E.E., N.J.K., M.S.-S., U.F. and J.S. conceived the study and designed the experiments. M.S.-S., H.G.-C., H.R.H., F. and E.E. generated cell dissociations and performed single-cell transcriptomic experiments using *Hydractinia* cell suspensions. D.A.S.-D. and H.R.H. performed bioinformatic analyses, N.J.K. performed preliminary bioinformatic single-cell analyses. A.P.-P. assisted in performing bioinformatic analyses on the transcriptional landscape of *Hydractinia*. R.M.G performed phylogenetic analysis of POU domain proteins. M.E.R. and M.A.-P. supervised by J.P. performed phylogenetic analyses. D.A.S.-D., H.R.H., M.S.-S., F, U.F. and J.S. contributed to the interpretation of the single-cell analysis data, with contributions from all other authors. D.A.S.-D., H.R.H., U.F. and J.S. wrote the manuscript and generated the figures, with contributions from all other authors. All authors read and approved the final version of the manuscript.

## Supplementary Note 1: Explanation of cluster naming and grouping

**i-cells, early and mixed progenitors (4, 8, 14):** We identified i-cells and their early progeny based on high *Piwi1* (LOC130623353) and Piwi2 (LOC130628528) expression and an absence of cell type specific marker expression, indicating an undifferentiated cell state. We determined that these clusters represent i-cells early and mixed progenitor cells as the cluster contains 8.5% of the total number of cells captured in the atlas, while in the animal true pluripotent i-cells represent 1% or less of the total cell population, as verified by flow cytometry of transgenic reporter animals ^66^.*Piwi1* and *Piwi2* were also expressed in cluster 8 but this cluster exhibits markers of specific cell types, as well as epigenetic regulators.

**sperm and spermatogenesis (6, 34):** Clusters 6 and 34 were determined to be sperm and spermatogenesis due to expression of *Piwi1* and *Piwi2* (germline marker genes and expression of spermatogenesis specific markers including; known marker *Histone H2B.3/4* (LOC130657957), verified by RNA in-situ hybridization ^67^, and sperm specific antigen 16 (HSymV2.0_g12.22020 LOC130622507), *Histone H2B.6* (LOC130636493), *Tubulin alpha 1 chain* (LOC130641471), and *Myosin 4-like* (LOC130621975).

**neurones (16, 26, 27, 29, 33):** This group of clusters was identified as neuronal due to expression ELAV orthologs (*Elav1*; LOC130630563, *Elav2*; LOC130612043 and *Elav3*; LOC130630562) ^26,68^. Cluster 16 was further identified as *RFamide* and *GLWamide* neurones (LOC130657354, LOC130624206 respectively) ^20^, clusters 26 and 33 were identified as *PQRFVamide* neurones (LOC130628716) and cluster 26 was further named Involucrin+ neurones (LOC130654945). Cluster 27 was identified as *Fax*+ neurones (LOC130645007) and cluster 29 was identified as *Pkdrej*+ neurones (LOC130623103). Additionally, all of these clusters co-occur in Supplementary Figure 1D.

**nematoblasts and nematocytes (10, 12, 15, 18, 25):** We identified cluster 12 as nematoblasts 1 ‘nb 1’ due to expression of *Piwi1*, *Piwi2* and *SoxB2* and shared expression of marker genes with cluster 25. Clusters 12, 18 and 25 express *Ncol1* (LOC130622551) a nematogenesis marker ^46^ and as such are nematoblasts. We named cluster 25 ‘nb 3+ nematoblast’ because even though the expression of *HsymJFT1c-I* (LOC130642394) predominates in this cluster, it shares many markers with other nematoblast clusters. Cluster 18 ‘*Hrnr*+ nematoblast’ expresses *HsymJFT2a* and *Hornerin-like* orthologs (LOC130641429 and LOC130653621 respectively). Additionally, clusters 12, 18 and 25 co-occur in Supplementary Figure 1D. Clusters 10 and 15 were showing expression of trichohyalin-like orthologs and *ARSTNd2* (LOC130657211, LOC130662747 and LOC130644310 respectively), and do not co-occur with any other cluster. We classify them as terminally differentiated nematocytes and name them ‘*Tchh1* & *2* + nematocytes’.

***Conodipine* + cells (9, 13, 36):** Cluster 9 and 36 are labelled ‘*Conodipine* + cells’ as they share markers with epithelial cells but have specific expression of *Conodipine* orthologs (LOC130644743, LOC130636858, LOC130642091). Also, they express at least one homolog of the Membrane Attack Complex and perforin family (MAC), a group of proteins found in cnidarians (LOC130635919) expressed in venom cells, so we classify these cells as venomous epithelial cells ^58^. While neither of these clusters co-occur with any particular other in Supplementary Figure 1D, cluster 13 groups with number 9 and is likely to be a *Conodipine* cell type.

**digestive gland cells (17, 22, 24):** Cluster 17 was identified as a gland cell type due to strong expression of known gland cell markers *Astacin1* (LOC130623614), *Astacin2* ( LOC130614501) ^69^, along with lower expression of *Chitinases* 1, 2 and 3 (LOC130649211, LOC130624325, LOC130655910 respectively) and a strong expression of zinc metalloproteinase *Nas 4-like* (LOC130614209). Cluster 22 reveals a strong expression of zinc metalloproteinase *Nas14-like* orthologs (LOC130636648, LOC130614442), together with expression of *Astacins* 1 and 2, at a lower level than in cluster 17. Cluster 24, *Antistasin*+ cells, had weak expression of *Astacins* 1 and 2 but strong expression of *Chitinases* 1, 2 and 3 along with specific expression of *Antistasin* orthologs (LOC130646107,LOC130657200, LOC130646090). Clusters 17, 22 and 24 co-occur and are likely to be digestive gland cell types (Supplementary Figure 1D).

**mucosal gland cells (21, 32):** *Rsp*+ cells, cluster 21, were named after the specific ortholog markers of rhamnospondin (LOC130621200, LOC130655212, LOC130622532), a known gland cell marker likely for production of the outer mucosal layer (glycocalyx) ^70^. Cluster 32 was identified as gland 4 cells due to the shared low expression of one rhamnospondin ortholog (LOC130622532). Due to co-occurrence of clusters 21, 32 (Supplementary Figure 1D) we classify these cells as mucosal gland cells based on their secretory nature.

**epithelial (0, 1, 7, 19, 23, 28, 30, 35, 37):** Clusters 0, 1, 7, 19, 23, 28, 30, 35, 37 were labelled as epithelial due to expression of general epithelial cell markers like *Proton-coupled zinc antiporter SLC30A1*, *Protocadherin Fat 4* orthologs and alpha *Catulin-like* (LOC130626173, LOC130648302, LOC130647734, LOC130655582 and LOC130641787 respectively) ^71^ and an absence of known muscle markers. Many markers of these clusters also closely co-occur with the epitheliomuscular group (Supplementary Figure 1D).

**epitheliomuscular (2, 3, 5, 20, 31):** Clusters 2, 3, 5, 20, and 31 were identified as epitheliomuscular cells due to expression of known muscle marker genes *Myosin heavy chain* (striated muscle-like) (LOC130649617), *Tropomyosin* (LOC130645893) ^72^, *Myophilin* orthologs (LOC130614185, LOC130613823, LOC130644608), titin-like (LOC120641779) ^73–75^, and close co-occurrence (Supplementary Figure 1D). This group also broadly co-occurs with the epithelial group and as such is a subclassification of epithelial cells. Cluster 2, 3, 31 is enriched in feeding polyps only and has specific expression of *Rhamnose-binding lectin-like* (LOC130635948) which may be specific to the mouth of feeding polyps ^76^.

***Shematrin-like*+ cells (11):** Cluster 11 was highly enriched in the stolon specific sample, but also contains cells from the mixed polyp type sample (likely due to the unclear interface between polyp and stolon). Cluster 11 co-occurs with epithelial cells but has specific expression of *Prisilkin* (LOC130630016) which is involved in mantle production in oyster and sea snail ^38^ and binds tightly to chitin.

**unannotated (37, 38, 39, 40, 41, 42, 43, 44, 45, 46, 47, 48, 49, 50, 51, 52):** These clusters were left unannotated either due to a lack of specific marker genes, a lack of annotated marker genes or lack of correlation with known clusters in the co-occurrence analysis. Additionally, these clusters appear to be small and distributed sporadically in the UMAP space, often with cells in multiple larger clusters. As such we believe these clusters are largely composed of artefacts such as doublets and cells with low read counts.

## Methods

### *Hydractinia* culture and experimental manipulation

*Hydractinia symbiolongicarpus* male clone 291-10 was cultured as described ^13^. Polyp samples were obtained by cutting them with fine surgical scissors close to the polyp-stolon boundary. Feeding and sexual polyps are easily distinguishable by morphology. Stolonal tissue was harvested by cutting away all polyps from a colony, followed by scraping the stolons with a single-sided razor blade.

### ACME dissociation

After sample collection, isolated colony parts were simultaneously fixed and dissociated using ACME solution ^30^ to generate distinct libraries per colony part. For libraries 09 (mixed feeding and sexual polyps) and 20 (mixed feeding and sexual polyps, and stolon only sub-libraries) ACME-a solution was prepared, on ice, using a 12:3:2:2:1 ratio of DEPC treated water, methanol, glacial acetic acid, glycerol and 7.5% N-Acetyl-Cysteine in DEPC water. Dissociation was performed by placing ±100 polyps, in 30 ppt filtered seawater (FSW), into nuclease free 1.5 mL tubes on ice. FSW was removed, followed by direct addition of 1 mL of ACME-a solution. Immediate vigorous pipetting was performed at the surface of the liquid to generate small bubbles (using a p1000 pipette, set to 700 uL), samples are completely dissociated after around 15 minutes of pipetting. After dissociation, samples in ACME-a were filtered through 40 µm Flowmi filters (Bel-Art™ SP Scienceware™ Flowmi™ Cell Strainers for 1000 μL Pipette Tips).

For libraries 27 (feeding polyps only) and 29 (sexual polyps only) ACME-b solution was prepared, on ice, using a 13:3:2:2 ratio of commercially sourced nuclease free water, methanol, glacial acetic acid and glycerol. Prior to dissociation, ±100 polyps were placed in 1.5 mL lobind tubes in FSW. FSW was removed and replaced with anaesthetic solution (filtered 4% MgCl_2_ made in 50:50 MilliQ H_2_O and FSW 30 ppt) and samples were left on ice for 15 minutes to anaesthetise. For removal of the glycocalyx and preservation of RNA, anaesthetic solution was removed and replaced by 200 uL of 7.5% N-Acetyl-Cysteine (in anaesthetic solution), which was immediately removed and replaced by 1 mL ACME-b solution. Samples were dissociated as above, followed by filtration through 100 and then 40 µm PluriSelect mini cell strainers (SKU 43-10040-40).

After filtration, samples were centrifuged for 5 minutes at 1200 g in a 4°C cooled swing out bucket centrifuge. Supernatant was aspirated and the pellet resuspended in 1X PBS in nuclease free water (volume dependant on the number of sub-samples to be concatenated e.g. 200 uL for concatenation of 5 samples, for a final volume of 1 mL). After resuspension, samples were concatenated for a final volume of 1 mL, followed again by 5 minutes centrifugation at 1200 g in a 4°C cooled swing out bucket centrifuge. Supernatant was aspirated and the pellet resuspended in 900 uL 1X PBS in nuclease free water, followed by addition of 100 uL DMSO. Tubes were gently inverted to homogenise the sample and DMSO, followed by storage at −80°C for up to 3 months.

### Flow Cytometry

Cells were prepared for SPLiT-seq by thawing on ice then centrifuging for 5 minutes at 1200 g in a 4°C cooled swing out bucket centrifuge. Supernatant was aspirated and the pellet resuspended in 500 uL of 1% BSA in PBS followed by another round of centrifugation. Supernatant was aspirated and the pellet resuspended in 450 uL of 1% BSA in PBS followed by filtration through a 50 μm CellTrics strainer (Sysmex). 50 uL of cell suspension was aliquoted and diluted by addition of 100 uL 1% BSA in PBS for a final 1 in 3 suspension. The 1 in 3 suspension was stained with 1.5 uL of DRAQ5 (1 in 10 dilution of a 5 mM stock, eBioscience) and 0.6 uL of Concanavalin-A (Con-A) conjugated with AlexaFluor 488 (1 mg/mL stock, Invitrogen) and incubated in the dark at room temperature for 25 minutes.

The stained 1 in 3 cell solution was assessed by flow cytometry (CytoFlex S Flow Cytometer, Beckman Coulter) and the number of singlet (based on FSC-H vs FSC-A), nucleated (based on DRAQ5 vs FSC-A, gated as a sub-population of singlets) and intact (Con-A positive vs FSC-A, gated as a sub-population of nucleated) was measured per 10 uL, in triplicate. An average of the three readings for each sample was used to calculate the number of cells per microlitre in the original cell suspension.

### SPLiT-seq Barcoding

SPLiT-seq barcoding was performed as in García-Castro *et al*. ^30^ with several modifications. Cells from libraries 09 and 20 were diluted in 0.5X PBS for a final concentration of 625 cells/uL and 8 uL of this suspension was added to each well of the RT/R1 barcoding plate (for a final concentration of 5000 cells per well). For libraries 27 and 29 cells were diluted in 0.5X PBS to a final concentration of 770 cells/uL and 6.5 uL of this suspension was added to each well of the RT/R1 barcoding plate (for a final concentration of 5000 cells per well). The addition of 6.5 uL of cell suspension, rather than 8 uL, in libraries 27 and 29 was to account for a modification to the RT master mix in these libraries. The RT master mix for libraries 27 and 29 contained 10% w/v PEG8000, to help with pellet aggregation in later centrifugation steps, thus increasing the volume of the master mix to 9.5 uL per well in the RT/R1 plate. All further steps of barcoding, sorting, lysis, cDNA purification, template switching, PCR and qPCR amplification and size selection were performed as per the original protocol.

Tagmentation and round 4 barcoding was performed in the same way for all libraries. The concentration and average fragment size of the size selected libraries was assessed by Qubit dsDNA HS assay (Invitrogen Q33230) and Bioanalyzer High Sensitivity DNA Analysis (Agilent Technologies AGLS5067-4626) respectively. The Nextera XT DNA Library Preparation Kit (Illumina 15032354) was used to perform tagmentation of 1 ng of cDNA, as calculated previously, per the manufacturer’s guidelines. The final Round 4 barcode was added by PCR by addition of 20-22 uL of tagmented cDNA. 15 uL of Nextera XT Kit PCR mix, 1 uL of 100 uM Tagmentation Master Primer (BC_0018) and 1 uL of 100 uM Round 4 Barcode (one of BC_0076-BC_0083 per sub-library) into a PCR tube. The reaction was incubated in a thermocycler at 72°C for 3 minutes, 95°C for 30 seconds, then 13 cycles of 95°C for 10 seconds, 55°C for 30 seconds and 72°C for 30 seconds, followed by 72°C for 5 minutes and 4°C hold.

The resultant libraries were again size selected at 0.7 and 0.6x as per the original protocol and assessed by Qubit and Bioanalyzer for pooling in an equimolar mix for sequencing. Pooled libraries were sequenced both shallow (10M reads per 10k cells) and deep (133M reads per 10k cells).

### DIAMOND

We implemented diamond v2.0.8.146 ^31^ to corroborate orthologs present in our reference genome on top of the annotation provided by NCBI. This software performed a blastp search against the whole downloaded database with default settings and organised the results into a table with the settings -- salltitles -b8 -c1 -p8 --outfmt 6 qseqid sseqid pident evalue stitle.

### eggNOG annotation

The annotated proteome of the assembled genome of *Hydractinia symbiolongicarpus* was downloaded from NCBI ^14^. We modified the headers of each gene so it only had their own protein id. The resulting translated genome (from now on referred to as proteome) was queried using EggNOG mapper ^77^ with the parameters: ‘-m diamond --sensmode sensitive --target_orthologs all --go_evidence non-electronic‘ against the EggNOG metazoa database. From the EggNOG output, GO term, functional category COG, and gene name association files, were generated using custom bash code. Full code is available at the project repository.

### SPLiT-seq read processing

Each library was sequenced using a NovaSeq 6000 platform (Illumina) by Novogene (China) and is available for download. The sequencing data comprises the following read counts: 112614850 (9_1), 137678940 (9_2), 139158858 (9_3), 131010556 (9_4), 154135270 (9_5), 578661976 (20_1), 802450388 (20_2), 960633216 (20_3), 669218030 (20_4), 690902500 (27_1), 676659158 (27_2), 807680330 (29_1), and 1026403228 (29_2). Initially, the quality of the output reads was assessed using FastQC (https://www.bioinformatics.babraham.ac.uk/projects/fastqc/). Subsequently, CutAdapt v2.8 ^78^ was employed to remove adapter sequences, short reads, and low-quality reads. We applied distinct strategies for each pair-end read file. Specifically, the command cutadapt -j 4 -m 60 -q 10 -b AGATCGGAAGAG was executed on each read 1 file to eliminate the Illumina universal adapter and any reads below 60 bp in length. For read 2 files, we ran the command cutadapt -j 4 -m 94 –trim -n -q 10 -b CTGTCTCTTATA to remove Nextera adapter sequences, reads shorter than 94 bp, and terminal Ns. Additionally, a “phase” step was performed using the command grep to identify sequences from read 2 files and ensure the corresponding barcodes were in the correct position. Finally, paired reads were generated using pairfq makepairs v0.17 (https://github.com/sestaton/Pairfq) to proceed with further analysis.

The previously assembled reference genome ^14^ served as the basis for creating a reference database for read mapping. Dropseq_tools-2.3.0 (https://github.com/broadinstitute/Drop-seq/releases/tag/v2.3.0) was then utilized to process the generated GTF file, resulting in the creation of a sequence dictionary, a refFlat file, a reduced GTF file, and corresponding interval files. To build the reference index, we employed STAR-2.7.3a (https://github.com/alexdobin/STAR/releases/tag/2.7.3a) with the parameters --sjdbOverhang 99, - -genomeSAindexNbases 13, and --genomeChrBinNbits 14. Each sub-library was individually processed and later combined in the analysis. We leveraged the SPLiT-seq toolbox (https://github.com/RebekkaWegmann/splitseq_toolbox), which incorporates algorithms from Drop-seq_tools-2.3.0, to retrieve, correct, and label the barcodes. The barcodes were labeled with a hamming distance ≤1. For mapping to the reference genome, we used STAR-2.7.3a (https://github.com/alexdobin/STAR/releases/tag/2.7.3a) with --quantMode GeneCounts and all other default settings, except for --outFilterMultimapNmax with 1 and 200 values, which created two modalities of the data and allowed us to retain and analyse reads that mapped up to one and to two hundred different loci in the reference. We employed Picard v2.21.1-SNAPSHOT (https://github.com/broadinstitute/picard) to re-order, merge, align, and tag reads for each sub-library, utilizing the SortSam and MergeBamAlignment features. To create expression matrices for each library, we implemented the Drop-seq_tools-2.3.0 features TagReadWithInterval and TagReadWithGeneFunction. We used the feature DigitalExpression from Drop-seq_tools-2.3.0 with the following settings: READ_MQ=0, EDIT_DISTANCE=1, MIN_NUM_GENES_PER_CELL=50, and LOCUS_FUNCTION_LIST=INTRONIC. For further details, the complete code and documentation can be found in the project repository. The resulting expression matrices, along with the gene models and raw reads, have been uploaded to GEO under the accession code GSE269914.

### Single cell analysis

We started processing this dataset with initially 241,340 cells in the “no multimappers” modality and 277,529 in the “with multimappers” modality. We retained both versions of the dataset in the same object following a multimodal integrative strategy with the python package MUON ^32^. All the following downstream analyses were performed to the “no multimappers” modality. The processing eliminated genes with high counts using sc.pp.filter_cells with min_counts=50 and min_genes=50. We calculated metrics using sc.pp.calculate_qc_metrics, sliced the matrix genes_by_counts < 700 and total_counts < 750. These steps eliminated 42,227 cells, giving us our final dataset of 199,113 cells. We normalised the matrix using sc.pp.normalize_total with a target_sum=1e4. We selected high variable genes using sc.pp.highly_variable_genes with n_top_genes = 18000, and sliced the matrix to contain only those genes, storing the raw in an adata.raw object. We then scaled the matrix with sc.pp.scale, performed pca with sc.tl.pca with n_comps=100, performed a batch correction with the python version of harmony ^33^ on the ‘X_pca’ column calculated in the previous step, we constructed a kNN graph with sc.pp.neighbors, with 40 neighbours, 75 principal components, made use of the representation ‘X_pca_harmony’, and calculated a UMAP visualisation with sc.tl.umap (min_dist=0.1, spread = 0.5, alpha = 1, gamma = 1.0). We run the Leiden clustering algorithm using sc.tl.leiden with resolutions 1, 1.5 and 2, which gave 47, 53 and 57 clusters respectively. We calculated marker genes for each cluster using sc.tl.rank_genes_groups, using the clusters of obtained with all 3 resolution parameters, and using both the Wilcoxon (method=’wilcoxon’) and the Logistic Regression (method=’logreg’) We selected resolution 1.5 for further downstream analyses.

### Compositional Analysis

To assess if cell clusters were significantly enriched across the different libraries representing the various colony parts of the animal. We implemented two approaches for testing this, one with the python package scCODA ^79^, which uses a Bayesian framework to model cell type counts while accounting for uncertainty in cell-type proportions and the negative correlative bias via modelling all the joint cell-type proportions. The other approach we also employed was a Fisher exact test. It’s important to note that this test is meant for small sample sizes, and it tends to greatly exaggerate p-values. To account for this statistical effect, we added small positive values, ensuring that variations in enrichment between clusters were not influenced or discarded due to statistical artefacts. For further details, the complete code and documentation can be found in the project repository.

### Count Per Million (CPM) calculation

A custom Python script (available in the project repository) was used to extract raw counts. This script sliced the raw unprocessed matrix to include only cells present in the processed matrix. Subsequently, cluster information was transferred from the processed matrix to the unprocessed one using a pandas script. To obtain the sum of all counts for each gene within each cluster, numpy was utilized on the matrix. The resulting raw summed counts dataset was then normalized by pseudobulk “library size” using the DESeqDataSetFromMatrix() function, where the parameter design = ∼condition, and the counts() function with parameter normalised = TRUE from the DESeq2 package ^80^. For further details, the complete code and documentation can be found in the project repository.

### Co-occurrence analysis

The cell type co-occurrence analysis was performed using the treeFromEnsembleClustering() function, sourced from the provided code ^25^. The function was executed with the following parameters: h = c(0.65, 0.95), clustering_algorithm = “hclust”, clustering_method = “average”, cor_method = “pearson”, p = 0.15, n = 1000, bootstrap=FALSE. The analysis performed 1000 iterations of cross-cell type Pearson correlation, utilising a downsampling of 85% on highly variable genes (FC > 1.5). Subsequently, hierarchical clustering of cell types was performed, and co-occurring pairs of cell types were quantified across iterations, generating a co-occurrence matrix. The final cell type tree was produced by hierarchically clustering the co-occurrence matrix. For further details, the complete code and documentation can be found in the project repository.

### Transcription factor annotation

The translated proteome of *Hydractinia* was queried for evidence of Transcription Factor (TF) homology using (i) InterProScan ^81^ against the Pfam ^82^, PANTHER ^83^, and (ii) SUPERFAMILY ^84^ domain databases with standard parameters, (iii) using BLAST reciprocal best hits ^85^ against swissprot transcription factors ^86^, and (iv) using OrthoFinder ^87^ with standard parameters against a set of model organisms (Human, Zebrafish, Mouse, Drosophila) with well annotated transcription factor databases (following AnimalTFDB v3.0) ^88^. For the latter, a given *Hydractinia* gene was counted as TF if at least another TF gene from any of the species belonged to the same orthogroup as the *Hydractinia* gene. The different sources of evidence were pooled together and we kept those *Hydractinia* genes with at least two independent sources of TF evidence. Every TF gene was assigned a class based on their sources of evidence. For complete transparency, the code is available at the project repository.

### Transcription factor analysis

The table of cpms was subsetted to retrieve TFs from *Hydractinia*, and gene expression across cell types was scaled and visualised using the ComplexHeatmap R package ^89^. To understand the TFs at a broader level, we calculated statistics for each TF class. For each class, we looked at how much gene expression varied across different cell types, with the median and average coefficient of variation (CV), the number of genes in that class and the cumulative gene counts. We visualised the relationship between CV and number of genes using the base and ggplot2 packages (https://ggplot2.tidyverse.org) in R v4.1.0 (https://www.R-project.org/).

To represent these findings visually, we quantified the prominence of each TF class in terms of gene counts. For each TF class, we calculated its prominence across different cell clusters by summing up the gene counts in that class within each cluster, and then dividing by the number of genes from that class expressed in that cluster. The resulting matrix was normalised and visualised using a custom ggplot2 wrapper function in R v4.1.0. Fully documented code is available at the project repository.

### WGCNA analysis

We conducted an analysis using Weighted Gene Co-Expression Network Analysis (WGCNA) ^40^ to elucidate the intricacies of gene interactions in our dataset. We subsetted genes which had a coefficient of variation (CV) greater than 1 from the calculated CPM table, and we set the softPower parameter to 14 following the assessment of Scale-Free Topology Model Fit. We then computed adjacency and Topological Overlap Matrices (TOM) using default settings. For the hierarchical clustering of genes, we chose a minimum module size of 75 genes and set the deepSplit parameter to 4. We named and colour-coded all resulting gene modules manually, in a similar fashion to how cell clusters are designated (Supplementary Dataset). These modules were used to reorganise the expression dataset, and the output was visualised using ComplexHeatmap.

To evaluate and represent the association between TF classes and gene modules, we calculated the mean connectivity of each TF gene to the module eigengenes. For each TF class, we calculated the number of genes in that class with a Spearman correlation coefficient equal to or greater than 0.01 with each module eigengene. We normalised the resulting matrix and visualised it using the ComplexHeatmap package. We also subsetted the WGCNA results matrix to only show TFs present in the gene modules.

We constructed WGCNA graphs using the TOM matrix, trimming sparse interactions with a low connectedness threshold (> 0.01). We employed the igraph package ^90^ and the Kamada-Kawai layout algorithm ^91^ for graph visualisation with parameters ‘maxiter = 100 * NUM_GENES_GRAPH, kkconst = NUM_GENES_GRAPH‘, where NUM_GENES_GRAPH is the number of genes present in the analysed graph. We assessed the membership of connected components with the function components() from the igraph package, and their agreement with WGCNA module membership using the adjusted Rand Index implementation adjustedRandIndex() from the package mclust ^92^.

The analysed graph was divided into subgraphs representing connected components using a custom wrapper function that employs the induced_subgraph() function from the igraph package. Following this, we calculated the centrality of TFs in each subgraph using the closeness() function from the igraph package. Output visualisation was done with ggplot2.

To explore cross-module connections, we created a ’gene x module’ matrix that counted how many genes from each module were direct neighbours of a given gene. This matrix was normalised by dividing the number of connections of a gene to each module by the size of the module to which the gene belonged. We aggregated these numbers at the module level to retrieve the number of normalised cross-connections between modules. The resulting matrix was converted into a graph using graph_from_adjacency_matrix() from igraph with parameters ‘mode = “upper”, weighted = TRUE, diag = FALSÈ, highlighting the strongest cross-connections based on edge size. Fully documented code is available at the project repository.

### Gene Ontology analysis

We conducted Gene Ontology (GO) analyses using the R package topGO v2.52.0 ^93^. We used the ’elim’ method with a custom wrapper function. To keep the analysis stringent, we excluded GO terms with fewer than three genes that were significantly linked to them. Unless stated otherwise, we used all the genes found in *Hydractinia* as the gene universe set to compare against.

### Flanking amino acid analysis of *Shematrin-like* sequences

To assess architecture of the gene set expressed in *Shematrin-like*+ cells, we developed a custom script searching for glycine repeats (singlets, duplets, triplets quadruplets in XG_n_X) between flanking aminoacids (I, L, V, Y) known for being characteristic of *Shematrins* (see code repository) ^38,39^. Furthermore, we identified signal peptide regions (N: n-terminal region, H: hydrophobic region, C: c-terminal region) of these genes running a locally installed version SignalP 6.0 ^53^ with default parameters except for ‘organism=eukarya, format=all, mode=slow-sequential’.

### *Stylophora* cell types gene scores

To understand and compare the evolutionary history of our annotated cell states, we observed clusters from another species which has publicly available single-cell data, *Stylophora pistillata* ^25^. We subset lists of genes corresponding to each cell state in *Stylophora* which had at least a value of 1 in their fold change expression. Later, we filtered these genes with the lists of one-to-one orthologs product of reciprocal blasts between *Hydractinia* and *Stylophora*. The filtered table of gene names in *Hydractinia* was later parsed through our dataset and scored across all cell states with the scanpy function sc.tl.score_genes. Where the distribution of a certain set of genes in each cell group is compared against the same distribution in all other cells not in the group.

### Phylogenetic inference

To assess conservation of POU family transcription factors we sampled POU domain containing protein sequences from 14 species, using known human POU protein sequences as Blast queries. Top hits were retained and confirmed by reciprocal Blast. Sequences were trimmed to include only functional domains as identified by NCBI Conserved Domains ^94^. Sequences were aligned using Clustal Omega ^95^ and vacancies and blur sites were removed manually. Phylogenies were inferred by Bayesian analysis using MrBayes ^96^ under aamodelpr fixed(poisson) with 1 heated chain and 8 cold chains, for 3 million generations.

Following the identification of sequences related to biomineralisation in the atlas of *Hydractinia*, we decided to infer their phylogenetic relationships. For each set of genes, we downloaded the corresponding sequences from Genbank. Identical sequences were removed with CD-HIT ^97^. The sequences of each gene were aligned using MAFFT L-INS-i v 7.429 ^97^. The maximum likelihood topologies were inferred with IQTree 2 v.2.2.0.3 ^98^. The most appropriate evolutionary model was selected with the Model Finder Plus (-MFP) ^99^ implementation in IQTree with 1000 replicates of ultrafast bootstrap sampling (-bb) ^100^. Visualisation of the final topologies was performed with FigTree v1.4.4 (https://github.com/rambaut/figtree) and IToL v5 ^101^.

## Supplementary Figure Legends

**Supplementary Figure 1.**

A: Violin plots of the number of genes and counts across all libraries. B: Violin plot showing the number of genes per cluster. C: Violin plot showing the number of counts per cluster. D-E: Violin plots showing number of genes and counts per colony part. Heatmap of the co-occurrence matrix showing similarities between cell types based on gene expression correlation.

**Supplementary Figure 2.**

A: Barplot showing number of genes quantified with more than 5 UMI counts per million (cpm) per cluster in a pseudobulk analysis. B: Boxplot of the log transformation of the gene counts per cluster showing the distribution of the data.

**Supplementary Figure 3.**

Gene tree without any modifications of the POU orthologs used to identify Pou4 in *Hydractinia*.

**Supplementary Figure 4**

Sections of Chromosome 2 and 3 from the genome assembly of *Hydractinia* showing the expression dotplots in the atlas, signal peptide regions, glycine repetitions and genomic locations of all the genes closely located or similar in sequence to *Shematrin* or *Prisilkin* orthologs.

**Supplementary Figure 5.**

Gene trees without any modifications of the biomineralization pathway components.

## Supplementary Files and Dataset Description

**Supplementary Dataset.**

Compiled table of the genes input of the WGCNA and Transcription Factor analysis pipeline. In one tab is the output of diamond blast. Each column corresponds to gene id, protein id, name given by NCBI, accession code of homolog output of diamond blast, sequence coverage of diamond blast, sequence e-value of diamond blast, ortholog match of diamond blast. In the next tab, is the annotation given to the genes input of the WGCNA and Transcription Factor analysis. Each column corresponds to Gene Ontology term product of the eggnog pipeline, Functional categorization product of the eggnog pipeline, WGCNA initial module name, WGCNA renamed module, WGCNA module renamed to be human friendly, Pfam domain name of the transcription factor if found, Pfam superfamily name of the transcription factor if found, reciprocal blast with swissprot transcription factor database, orthofinder symbol of match with transcription factor database, orthofinder family match with transcription factor database, preferred name of the transcription factor if is classified as such, transcription factor class and summary name of the gene id together with transcription factor class.

**Supplementary File 1.**

Description of treatments per library and sublibrary, and general statistics of mapping when multimapping to multiple loci is and is not allowed. For details on this, check the materials and method section and the code repository.

**Supplementary File 2.**

Top 100 marker genes per cluster in resolution 1.5 of the Leiden algorithm calculated through the Wilcoxon rank-sum method. Each column in the corresponding tabs refers to gene names, logarithmic fold changes, p-values, adjusted p-values, scores, ortholog name output of a diamond blast and name given by ncbi.

**Supplementary File 3.**

Top 30 marker genes per cluster in resolution 1.5 of the Leiden algorithm calculated through the logarithmic regression method. Each column in the corresponding tabs refers to gene names, scores, ortholog name output of a diamond blast and name given by ncbi.

**Supplementary File 4.**

UMAPs corresponding to the genes found in both methods used to calculate defining markers per cluster.

**Supplementary File 5.**

Aggregated cell counts for each broad cell type category and individual cell type, together with the corresponding percentages of these cell counts and hex colour code used per cluster and per broad category.

**Supplementary File 6**

GO biological process terms enrichment (log10(p-value)) per WGCNA module.

