## Supplementary material for "The *Hydractinia* cell atlas reveals cellular and molecular principles of cnidarian coloniality": Hydractinia Cell Atlas Supplementary File 4 vs1 20240213.pdf

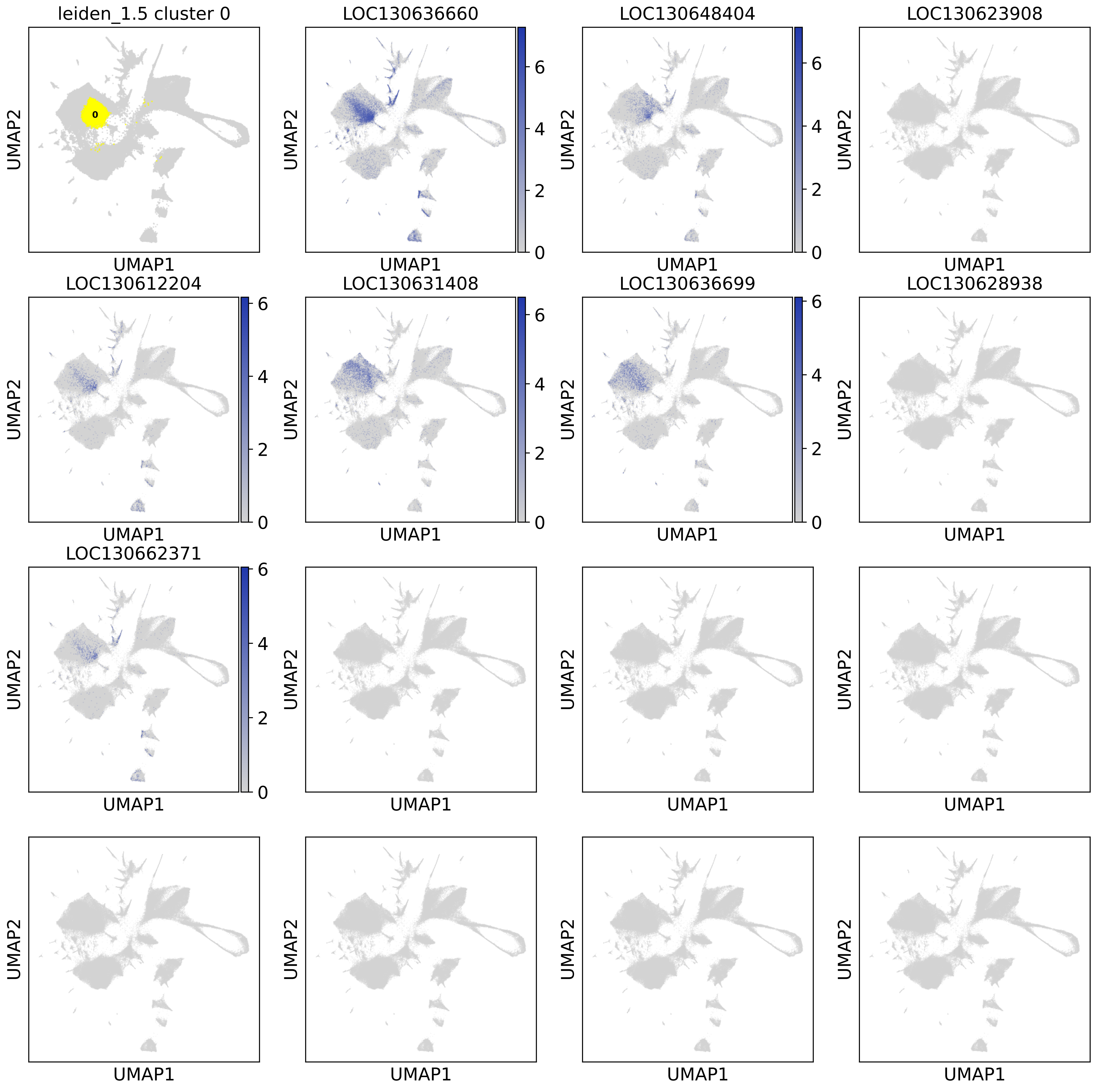

leiden_1.5 cluster 0
0
LOC130636660
LOC130648404
LOC130623908
LOC130612204
LOC130631408
LOC130636699
LOC130628938
LOC130662371
0
2
4
6
UMAP1
UMAP2

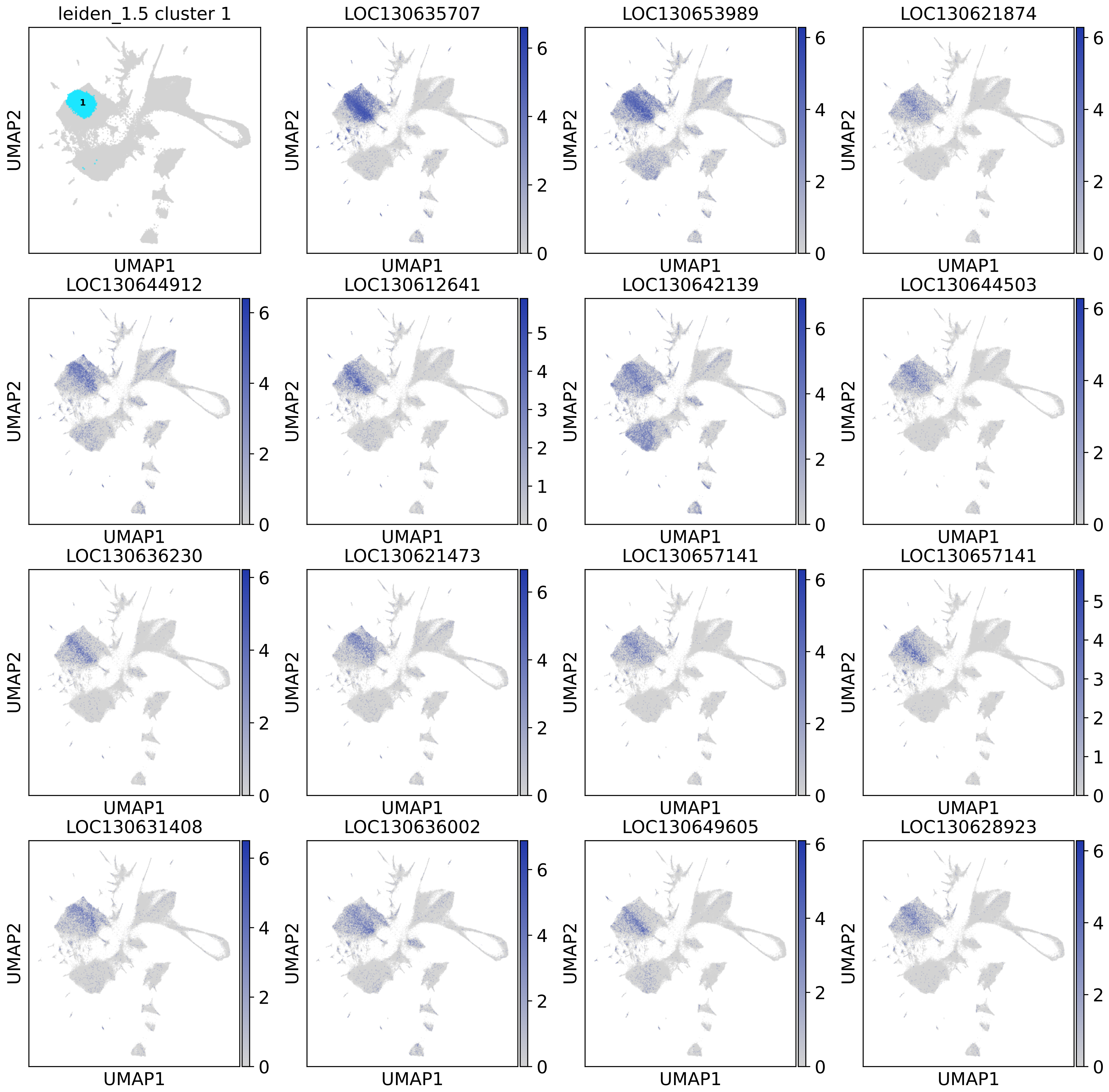
leiden_1.5 cluster 1
1
UMAP1
UMAP2
LOC130635707
LOC130653989
LOC130621874
LOC130644912
LOC130612641
LOC130642139
LOC130644503
LOC130636230
LOC130621473
LOC130657141
LOC130657141
LOC130631408
LOC130636002
LOC130649605
LOC130628923
0
1
2
3
4
5
6

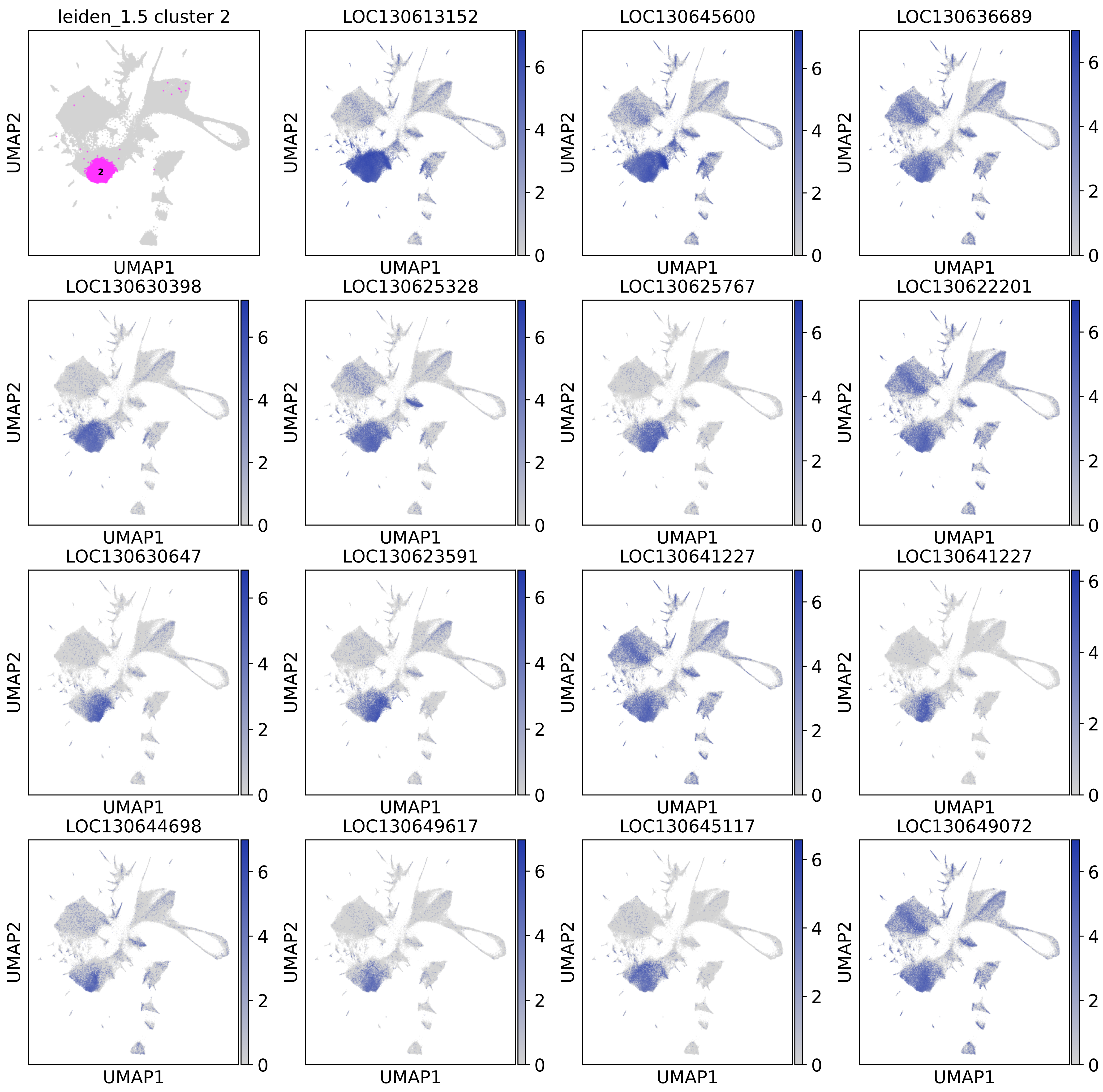
leiden_1.5 cluster 2
LOC130613152
LOC130645600
LOC130636689
LOC130630398
LOC130625328
LOC130625767
LOC130622201
LOC130630647
LOC130623591
LOC130641227
LOC130641227
LOC130644698
LOC130649617
LOC130645117
LOC130649072
UMAP1
UMAP2

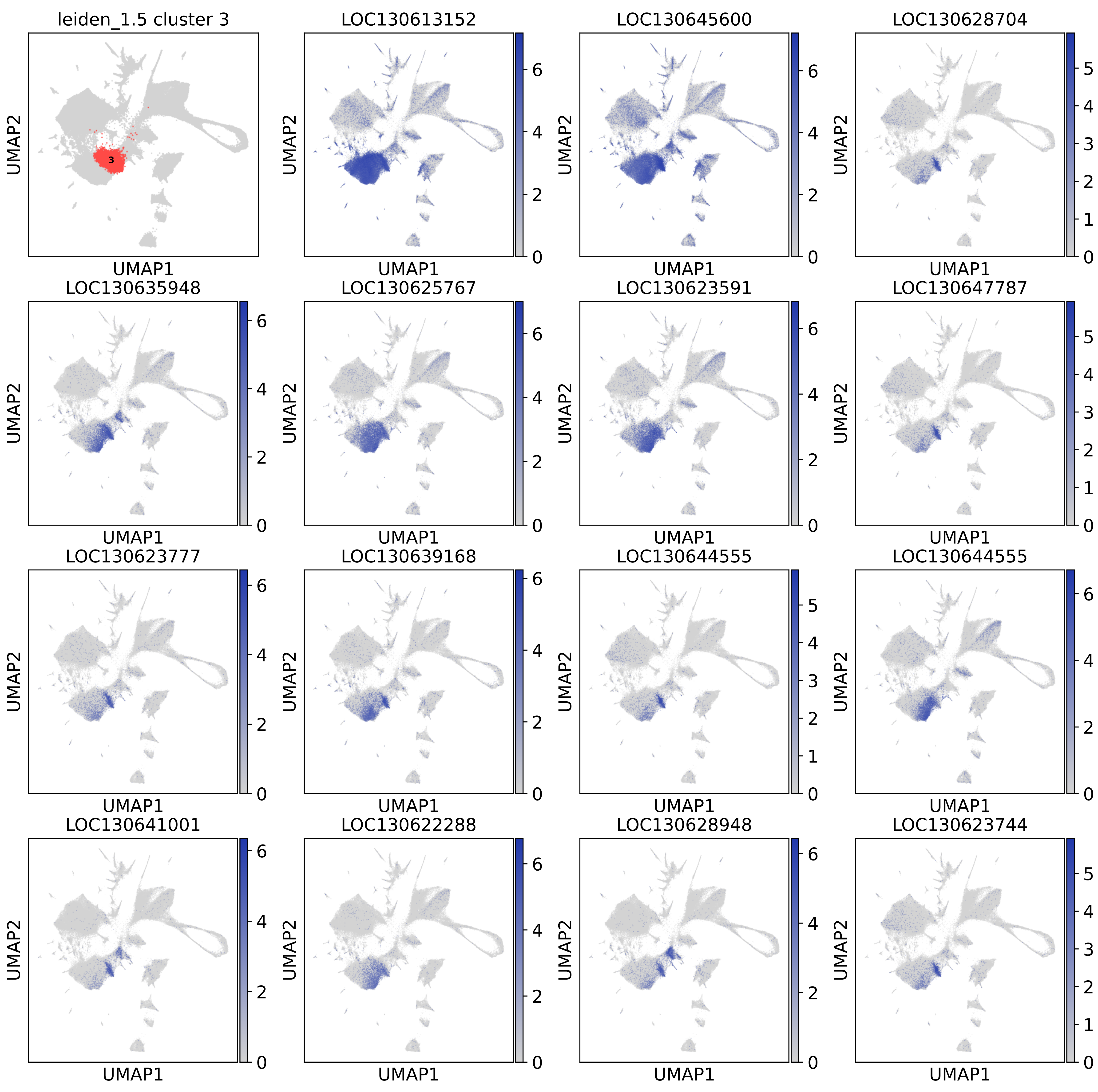
leiden_1.5 cluster 3
LOC130613152
LOC130645600
LOC130628704
LOC130635948
LOC130625767
LOC130623591
LOC130647787
LOC130623777
LOC130639168
LOC130644555
LOC130644555
LOC130641001
LOC130622288
LOC130628948
LOC130623744
UMAP1
UMAP2

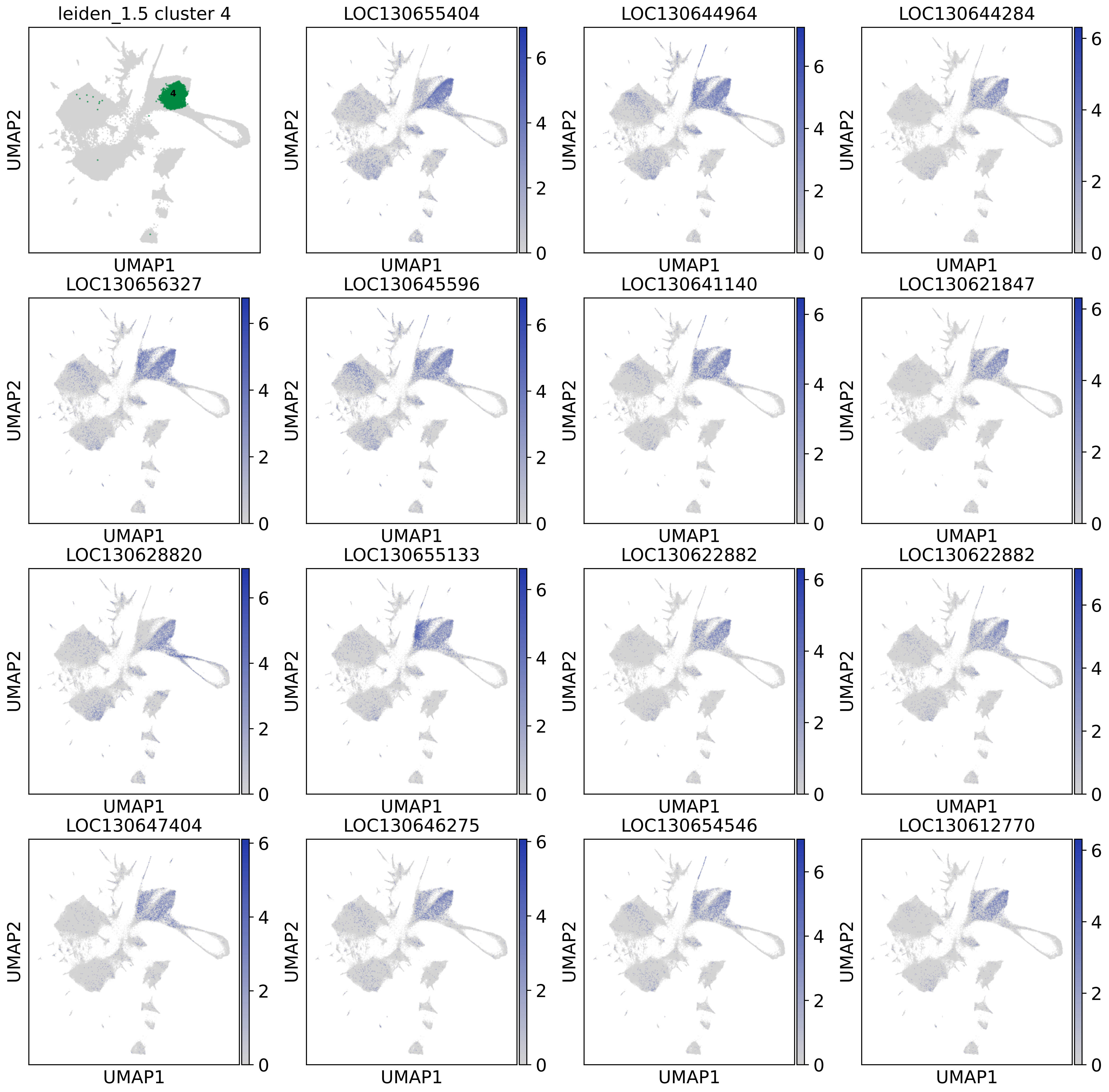
leiden_1.5 cluster 4
4
LOC130655404
LOC130644964
LOC130644284
LOC130656327
LOC130645596
LOC130641140
LOC130621847
LOC130628820
LOC130655133
LOC130622882
LOC130622882
LOC130647404
LOC130646275
LOC130654546
LOC130612770
UMAP1
UMAP2
0
2
4
6

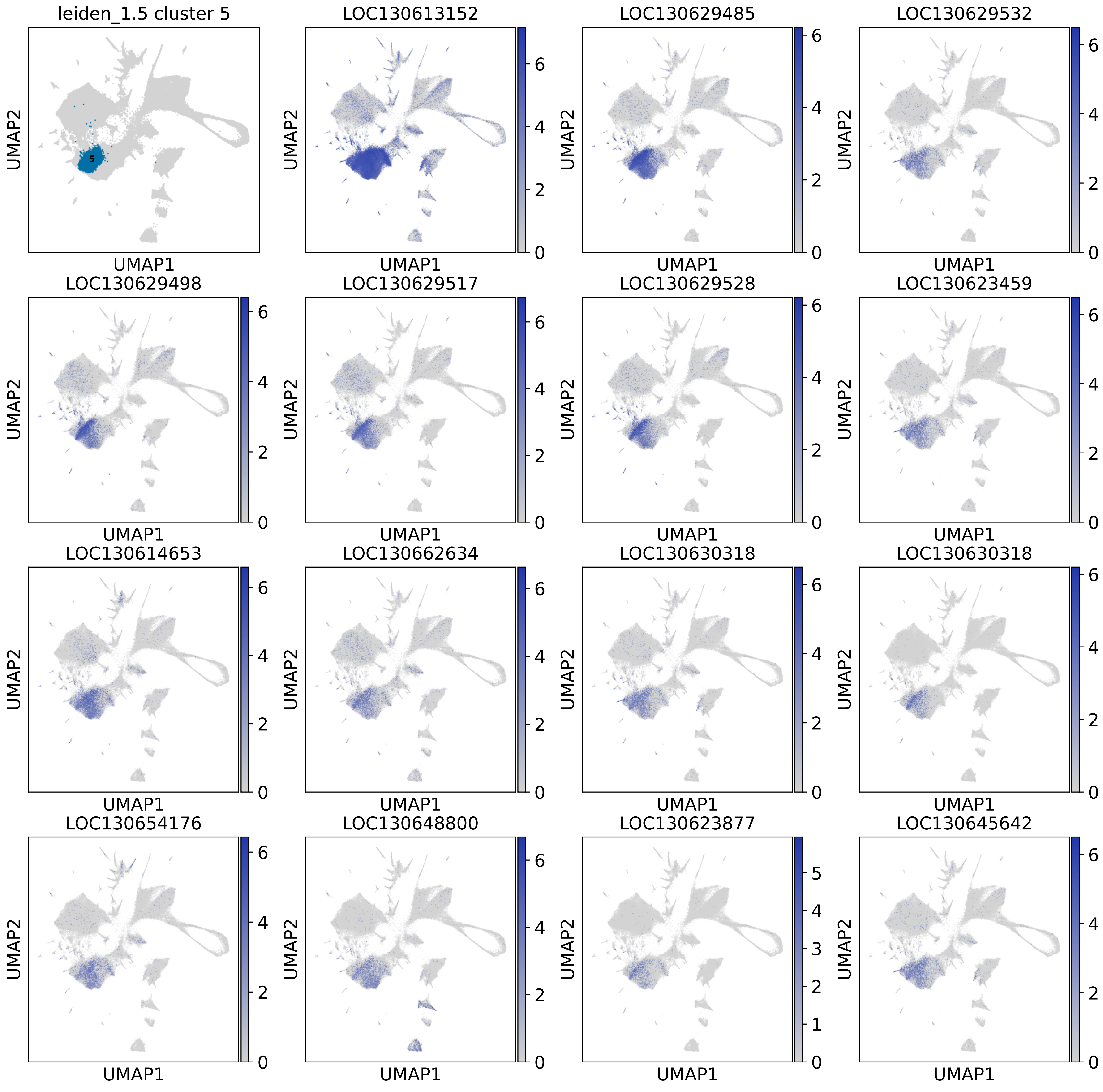
leiden_1.5 cluster 5
5
LOC130613152
LOC130629485
LOC130629532
LOC130629498
LOC130629517
LOC130629528
LOC130623459
LOC130614653
LOC130662634
LOC130630318
LOC130630318
LOC130654176
LOC130648800
LOC130623877
LOC130645642
UMAP1
UMAP2

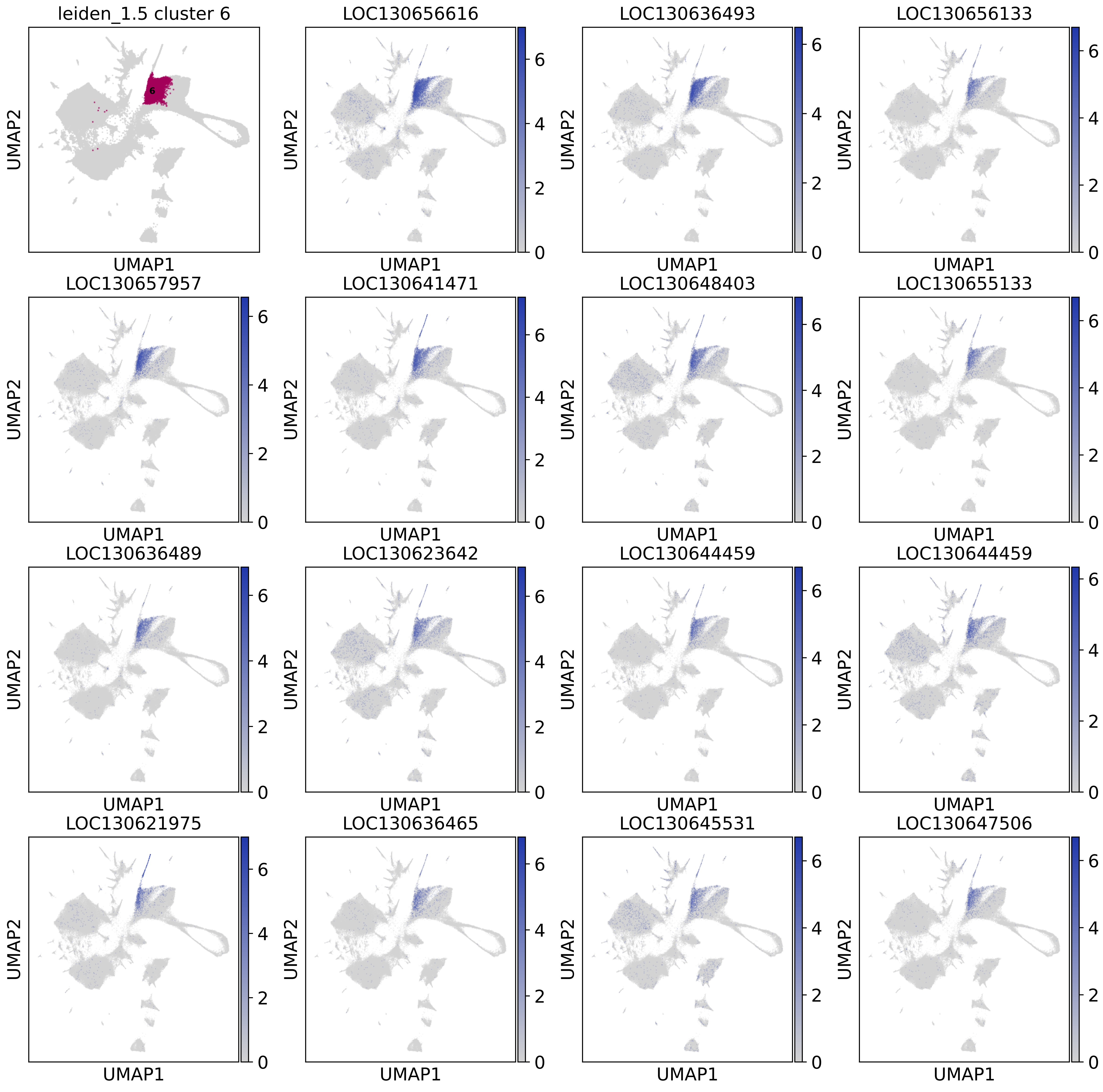
leiden_1.5 cluster 6
6
LOC130656616
LOC130636493
LOC130656133
LOC130657957
LOC130641471
LOC130648403
LOC130655133
LOC130636489
LOC130623642
LOC130644459
LOC130644459
LOC130621975
LOC130636465
LOC130645531
LOC130647506
UMAP1
UMAP2
0
2
4
6

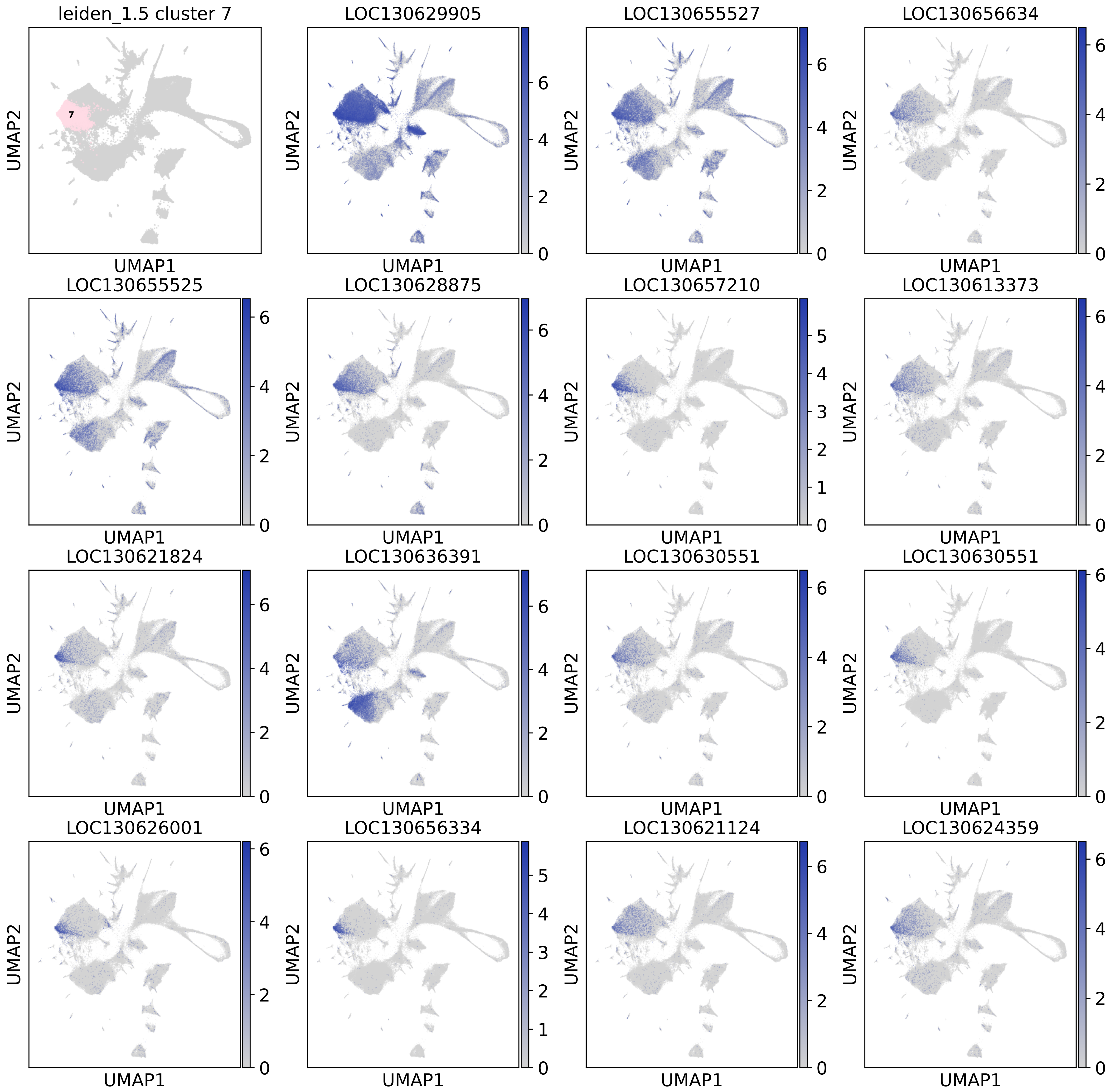
leiden_1.5 cluster 7
LOC130629905
LOC130655527
LOC130656634
LOC130655525
LOC130628875
LOC130657210
LOC130613373
LOC130621824
LOC130636391
LOC130630551
LOC130630551
LOC130626001
LOC130656334
LOC130621124
LOC130624359
UMAP1
UMAP2

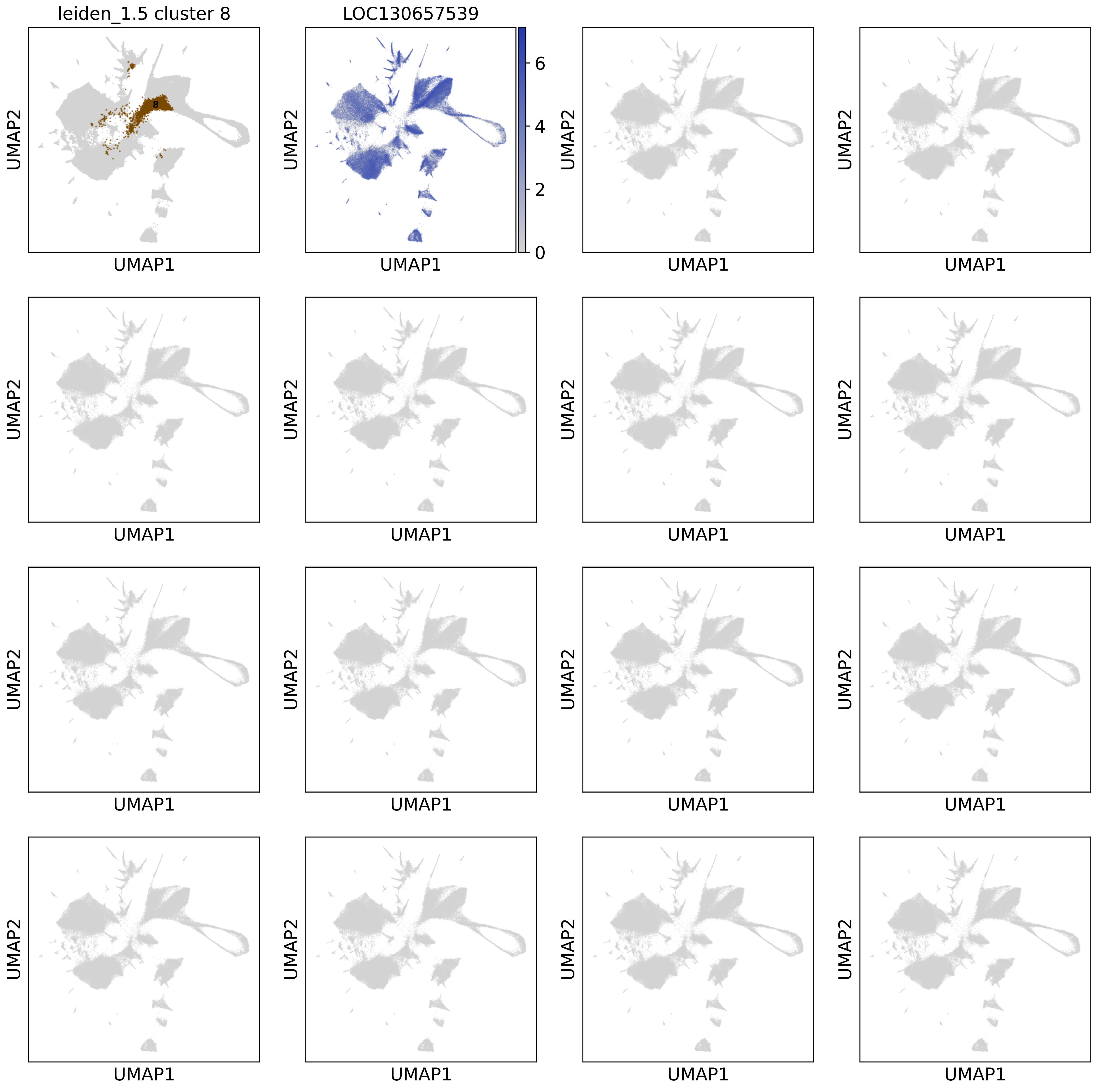
leiden_1.5 cluster 8
LOC130657539
8
0
2
4
6
UMAP1
UMAP2

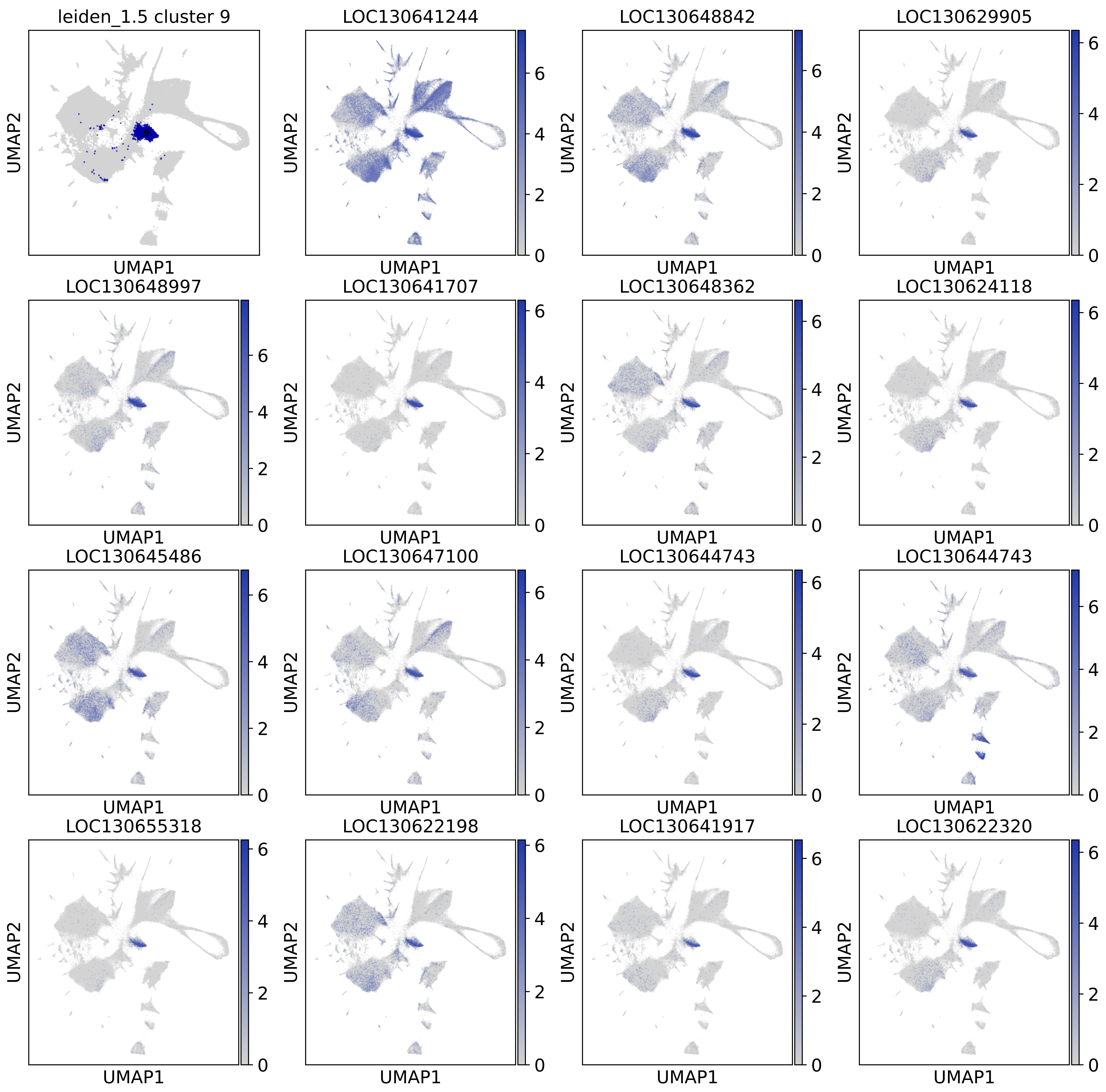
leiden_1.5 cluster 9
9
LOC130641244
LOC130648842
LOC130629905
LOC130648997
LOC130641707
LOC130648362
LOC130624118
LOC130645486
LOC130647100
LOC130644743
LOC130644743
LOC130655318
LOC130622198
LOC130641917
LOC130622320
UMAP1
UMAP2
0
2
4
6

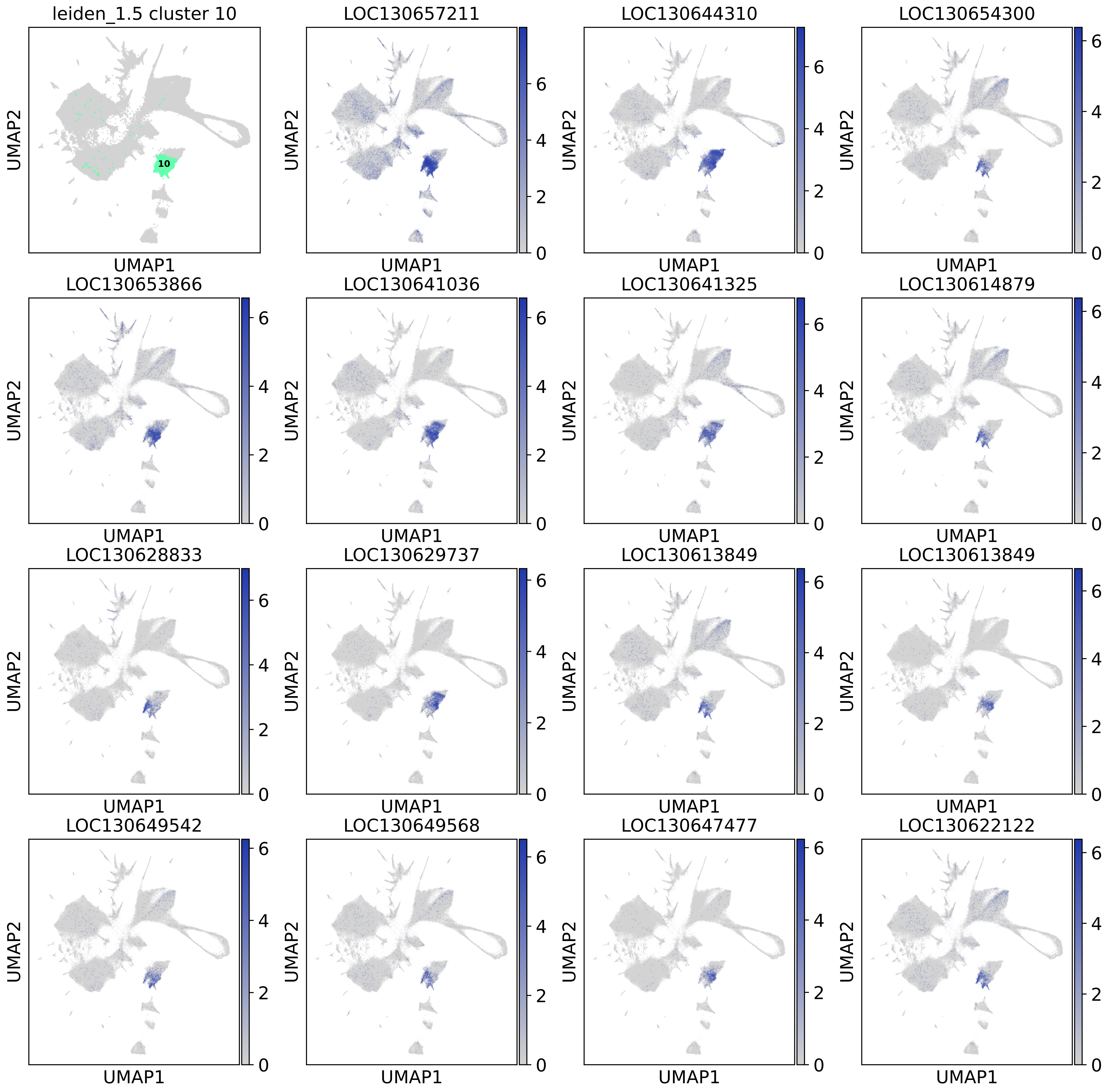

leiden_1.5 cluster 10
10
LOC130657211
LOC130644310
LOC130654300
LOC130653866
LOC130641036
LOC130641325
LOC130614879
LOC130628833
LOC130629737
LOC130613849
LOC130613849
LOC130649542
LOC130649568
LOC130647477
LOC130622122
UMAP1
UMAP2
0
2
4
6

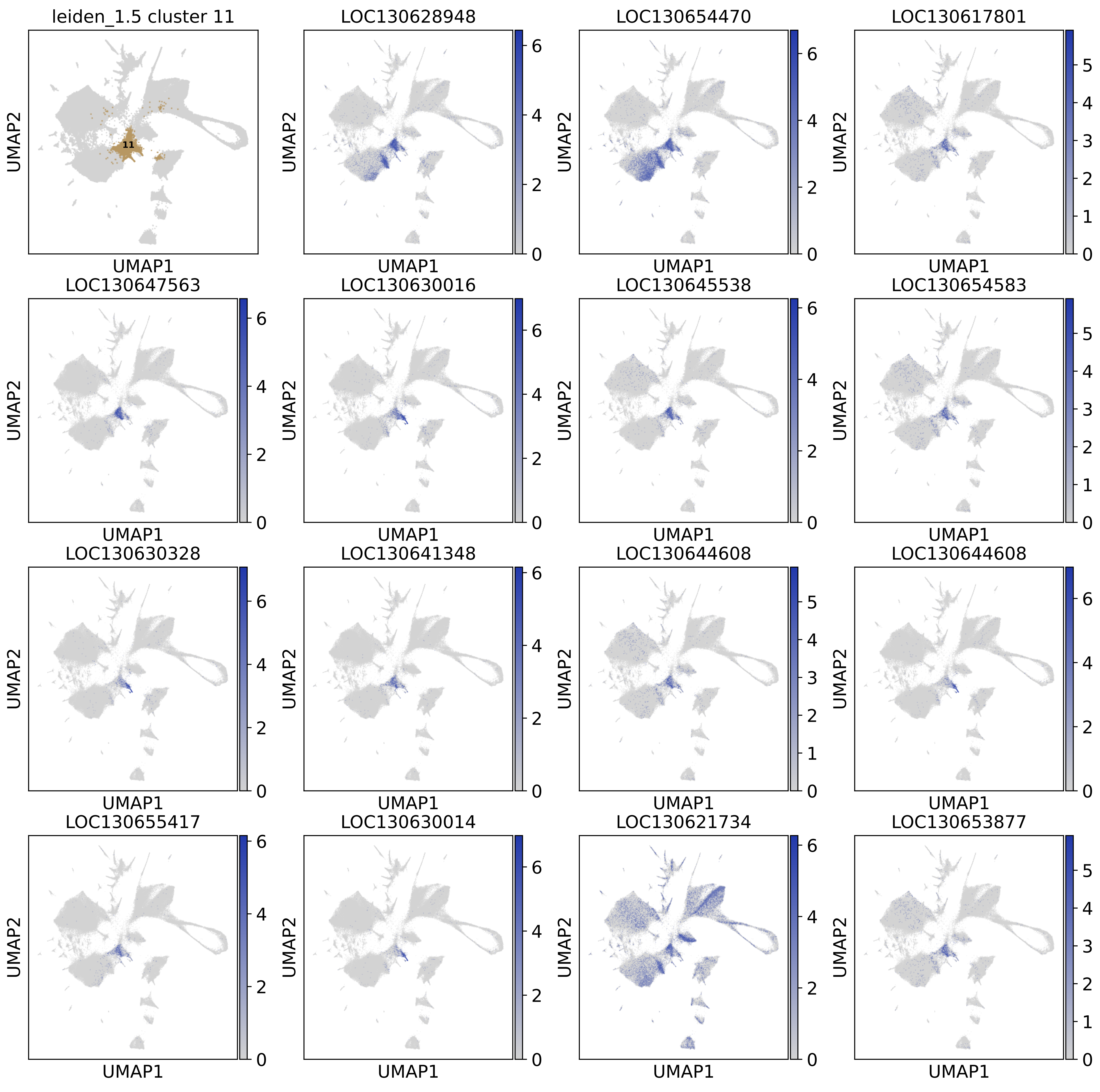
leiden_1.5 cluster 11
11
LOC130628948
LOC130654470
LOC130617801
LOC130647563
LOC130630016
LOC130645538
LOC130654583
LOC130630328
LOC130641348
LOC130644608
LOC130644608
LOC130655417
LOC130630014
LOC130621734
LOC130653877
UMAP1
UMAP2

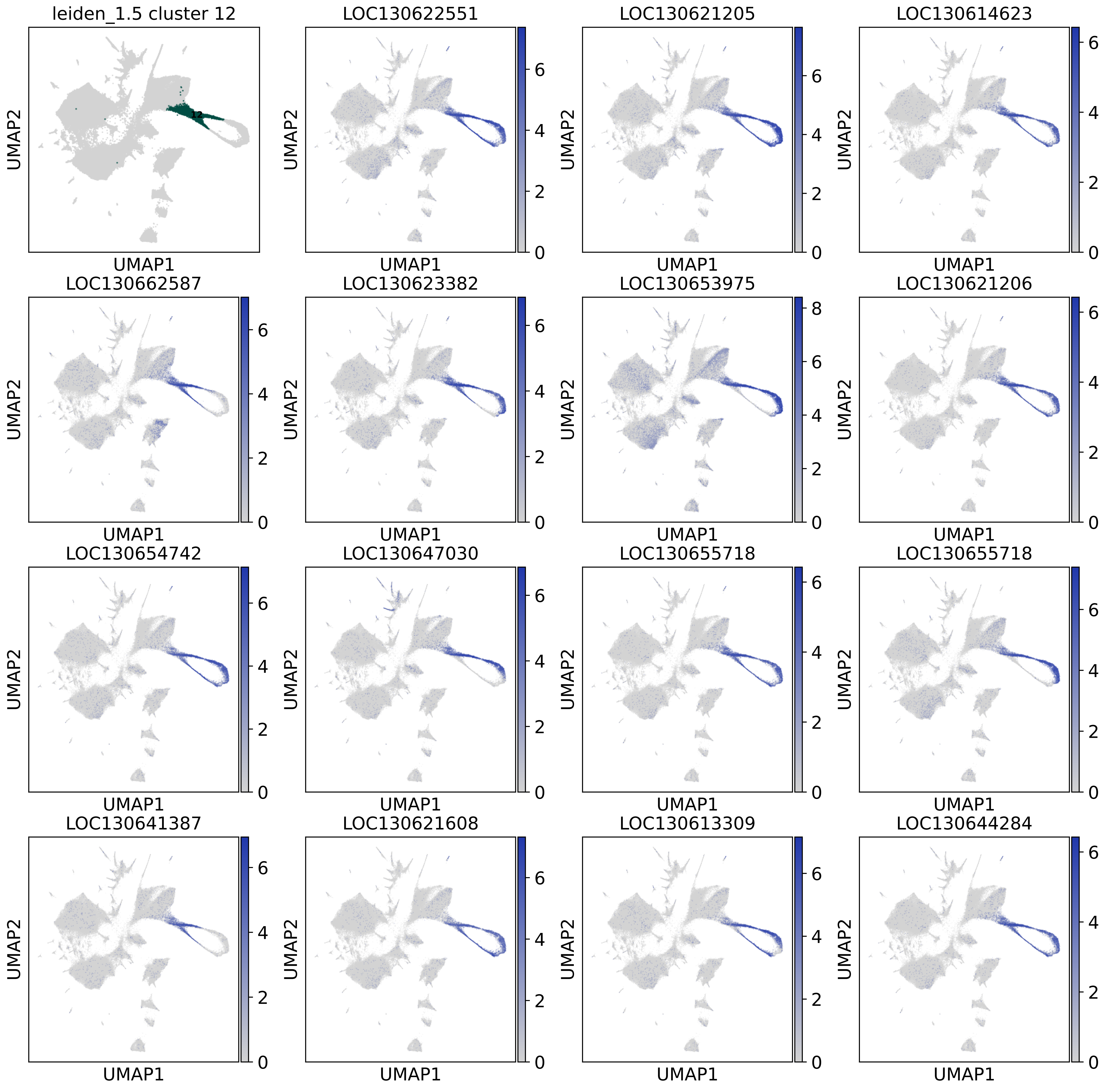
leiden_1.5 cluster 12
12
LOC130622551
LOC130621205
LOC130614623
LOC130662587
LOC130623382
LOC130653975
LOC130621206
LOC130654742
LOC130647030
LOC130655718
LOC130655718
LOC130641387
LOC130621608
LOC130613309
LOC130644284
UMAP1
UMAP2

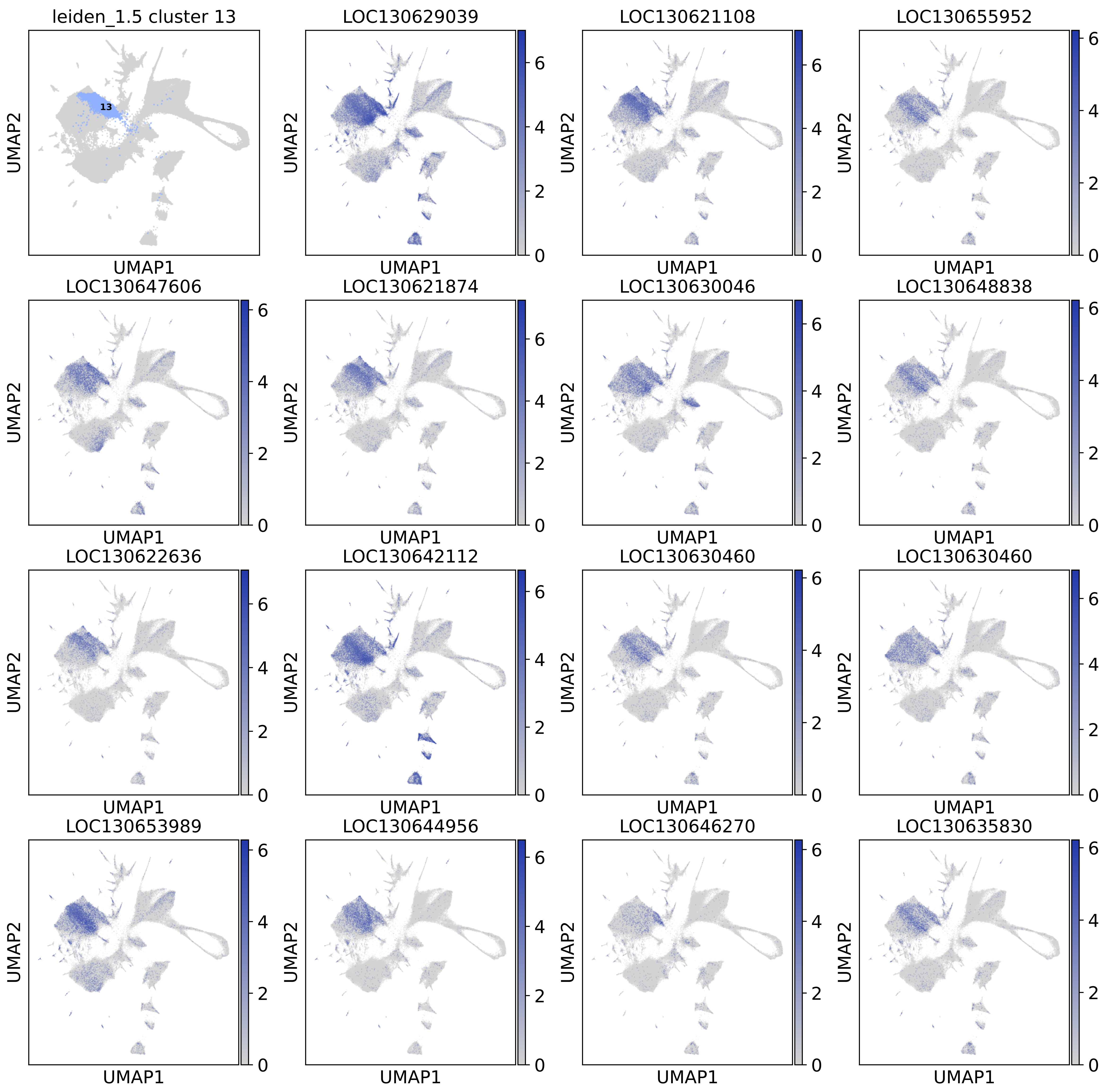
leiden_1.5 cluster 13
13
LOC130629039
LOC130621108
LOC130655952
LOC130647606
LOC130621874
LOC130630046
LOC130648838
LOC130622636
LOC130642112
LOC130630460
LOC130630460
LOC130653989
LOC130644956
LOC130646270
LOC130635830
UMAP1
UMAP2

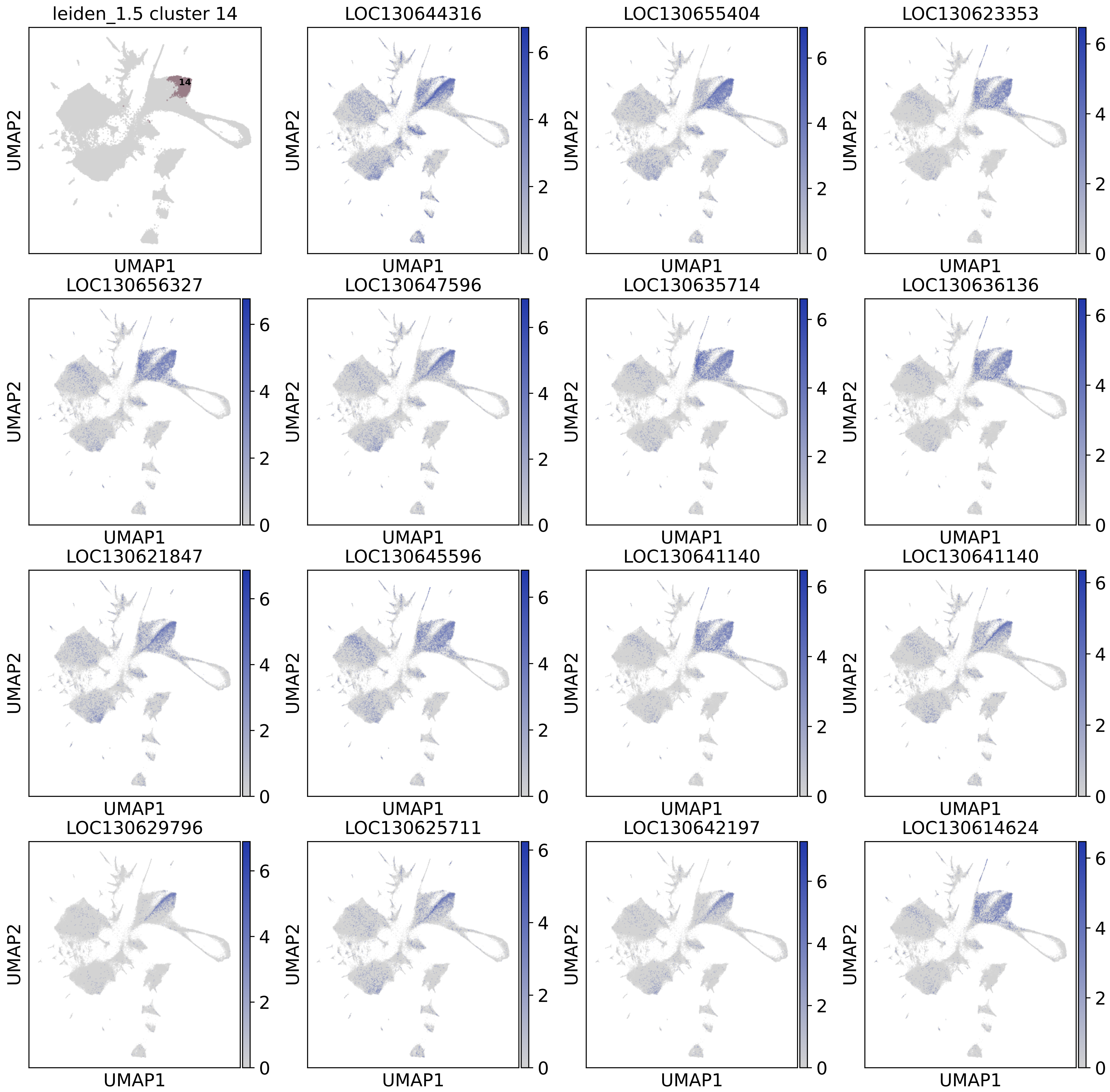

leiden_1.5 cluster 14
14
LOC130644316
LOC130655404
LOC130623353
LOC130656327
LOC130647596
LOC130635714
LOC130636136
LOC130621847
LOC130645596
LOC130641140
LOC130641140
LOC130629796
LOC130625711
LOC130642197
LOC130614624
UMAP1
UMAP2
0
2
4
6

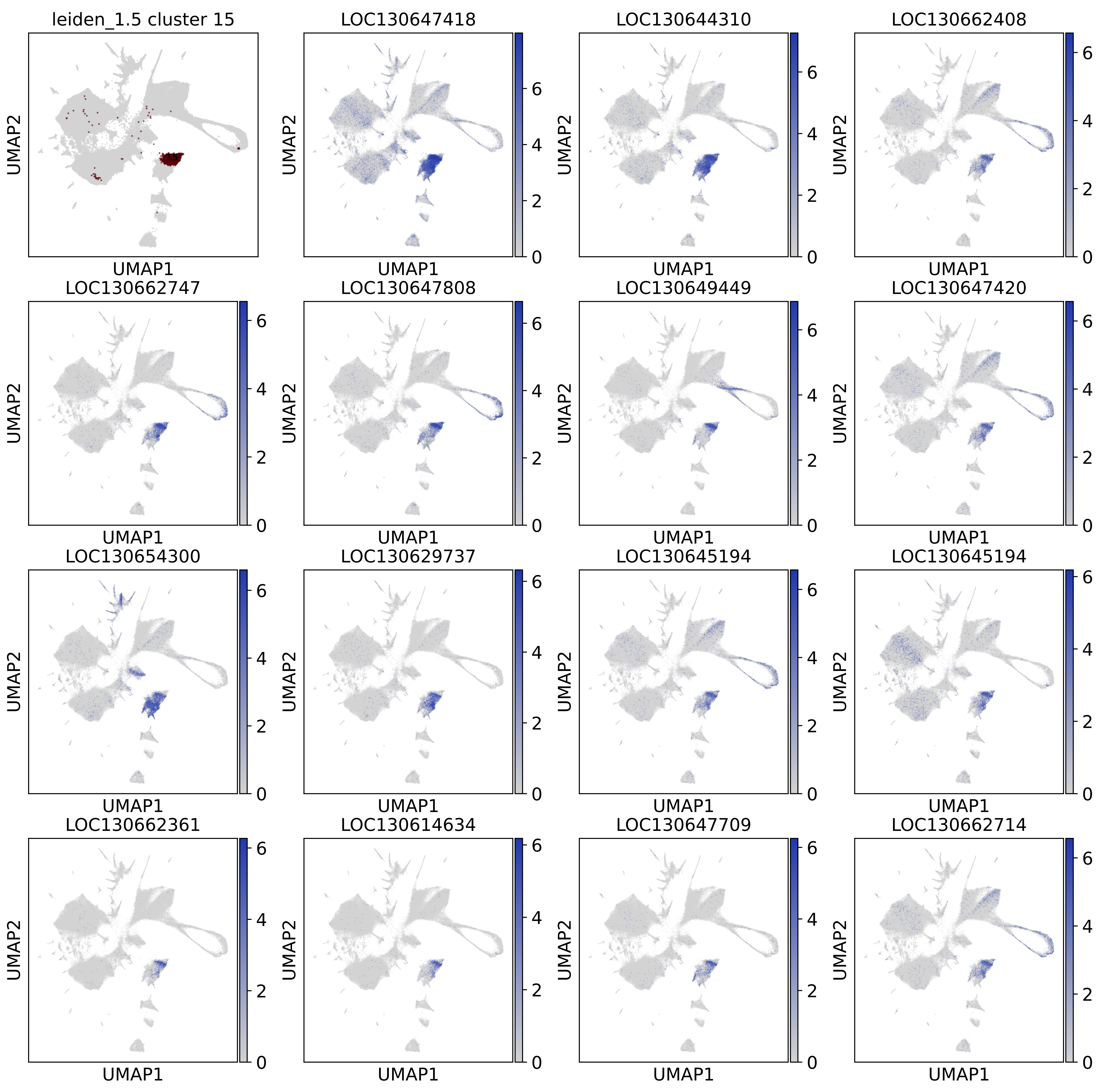
leiden_1.5 cluster 15
LOC130647418
LOC130644310
LOC130662408
LOC130662747
LOC130647808
LOC130649449
LOC130647420
LOC130654300
LOC130629737
LOC130645194
LOC130645194
LOC130662361
LOC130614634
LOC130647709
LOC130662714
UMAP1
UMAP2
0
2
4
6

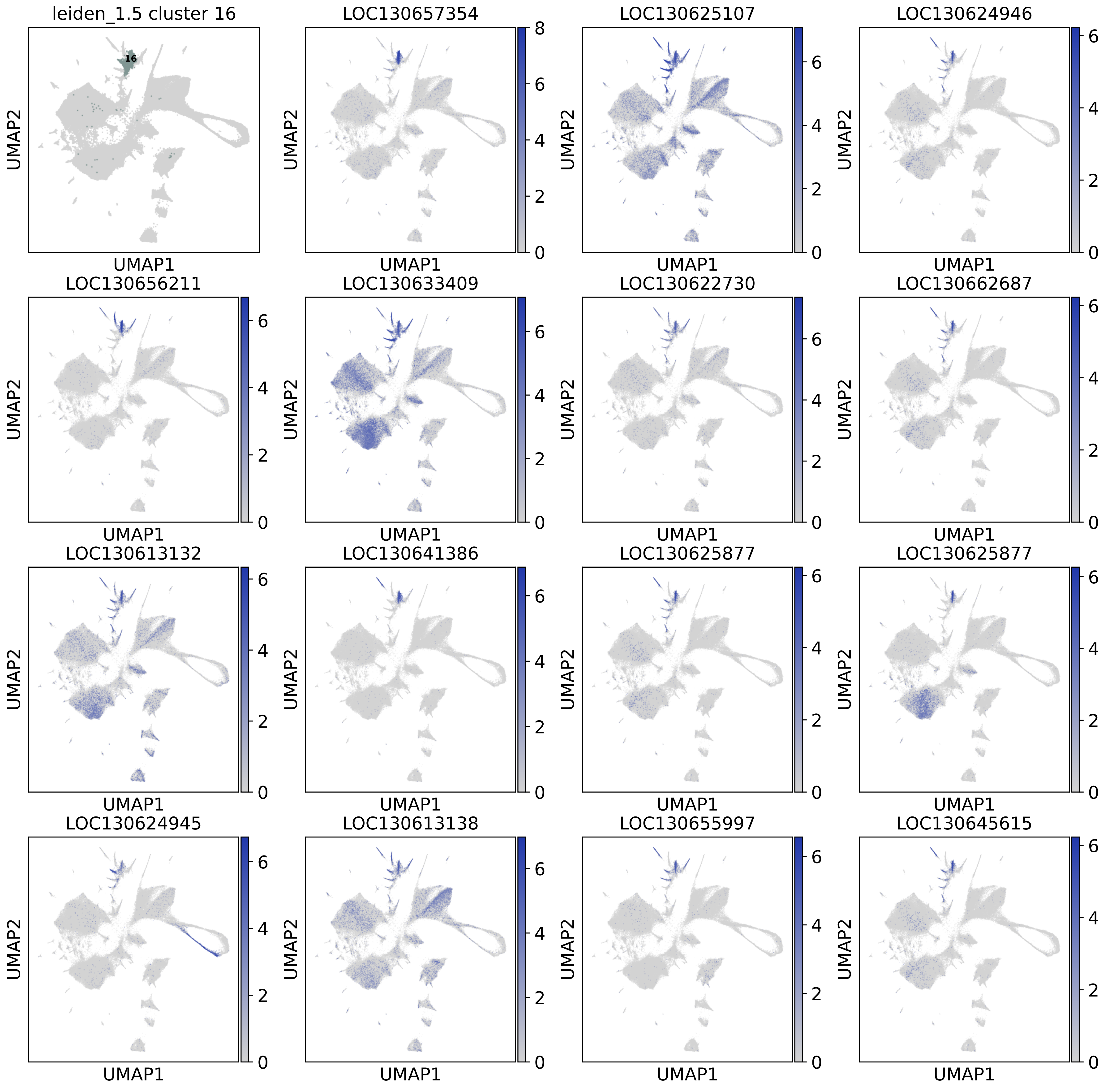

leiden_1.5 cluster 16
16
LOC130657354
LOC130625107
LOC130624946
LOC130656211
LOC130633409
LOC130622730
LOC130662687
LOC130613132
LOC130641386
LOC130625877
LOC130625877
LOC130624945
LOC130613138
LOC130655997
LOC130645615
UMAP1
UMAP2

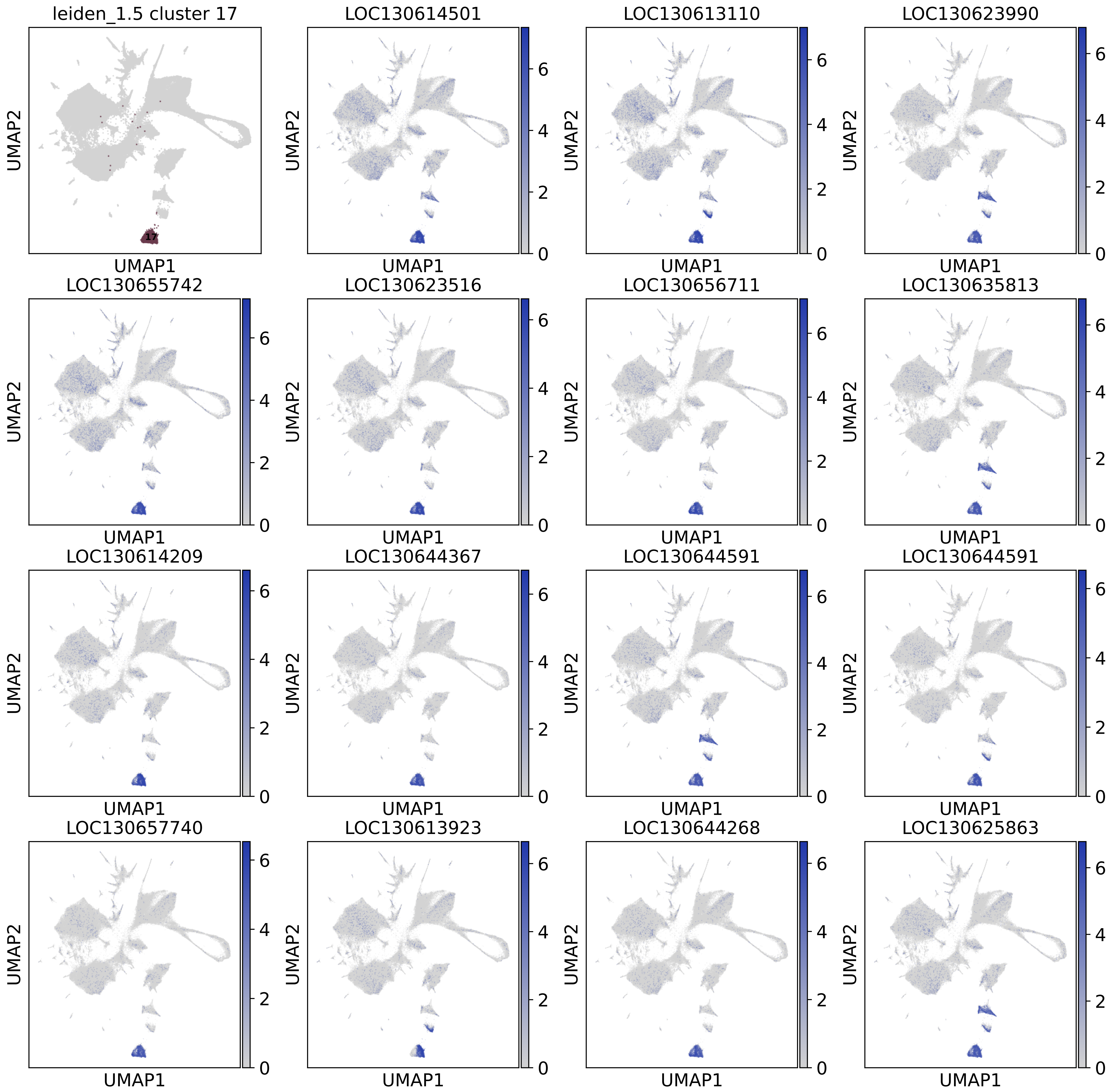

leiden_1.5 cluster 17
17
LOC130614501
LOC130613110
LOC130623990
LOC130655742
LOC130623516
LOC130656711
LOC130635813
LOC130614209
LOC130644367
LOC130644591
LOC130644591
LOC130657740
LOC130613923
LOC130644268
LOC130625863
UMAP1
UMAP2
0
2
4
6

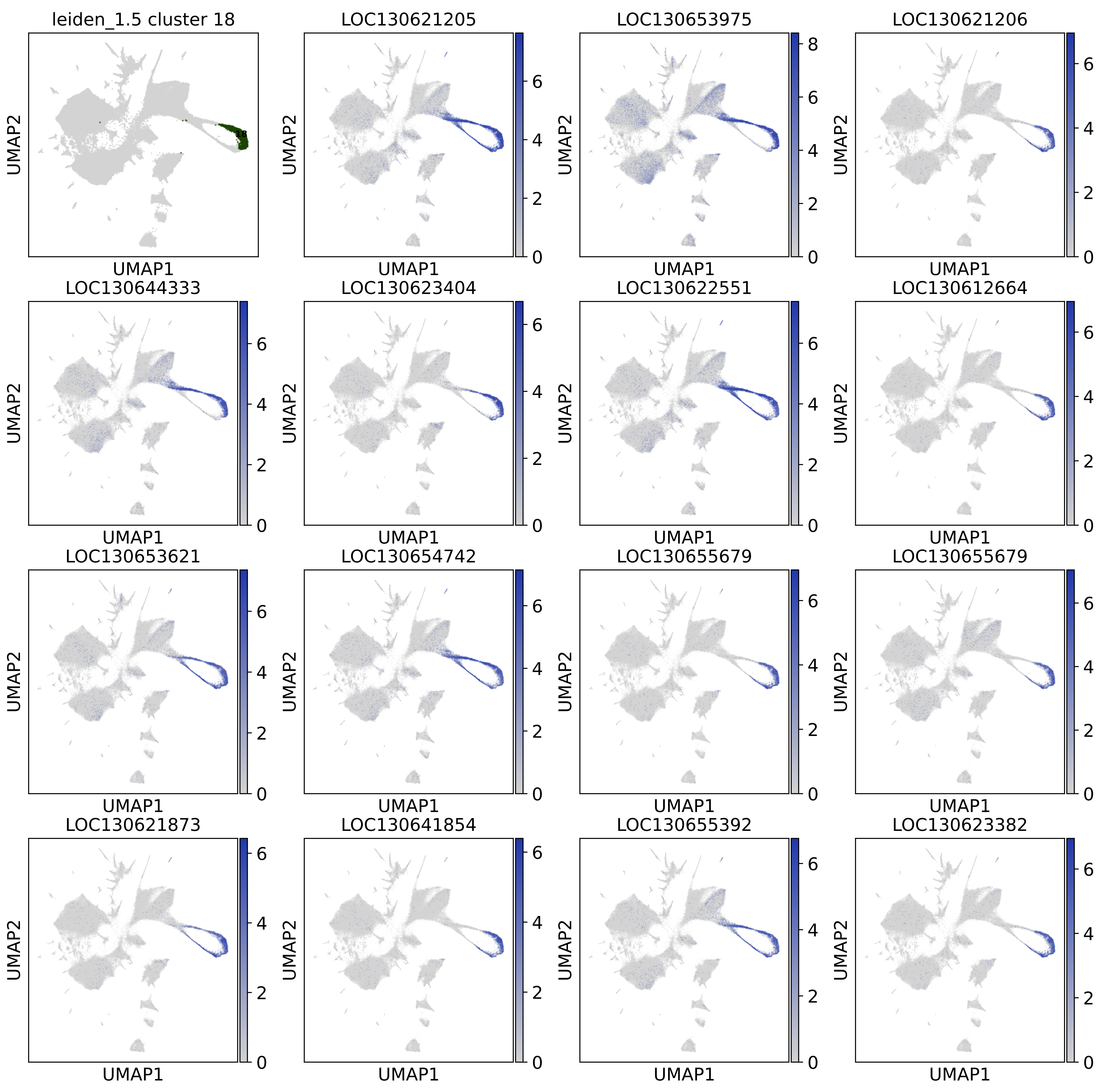
leiden_1.5 cluster 18
18
LOC130621205
LOC130653975
LOC130621206
LOC130644333
LOC130623404
LOC130622551
LOC130612664
LOC130653621
LOC130654742
LOC130655679
LOC130655679
LOC130621873
LOC130641854
LOC130655392
LOC130623382
UMAP1
UMAP2

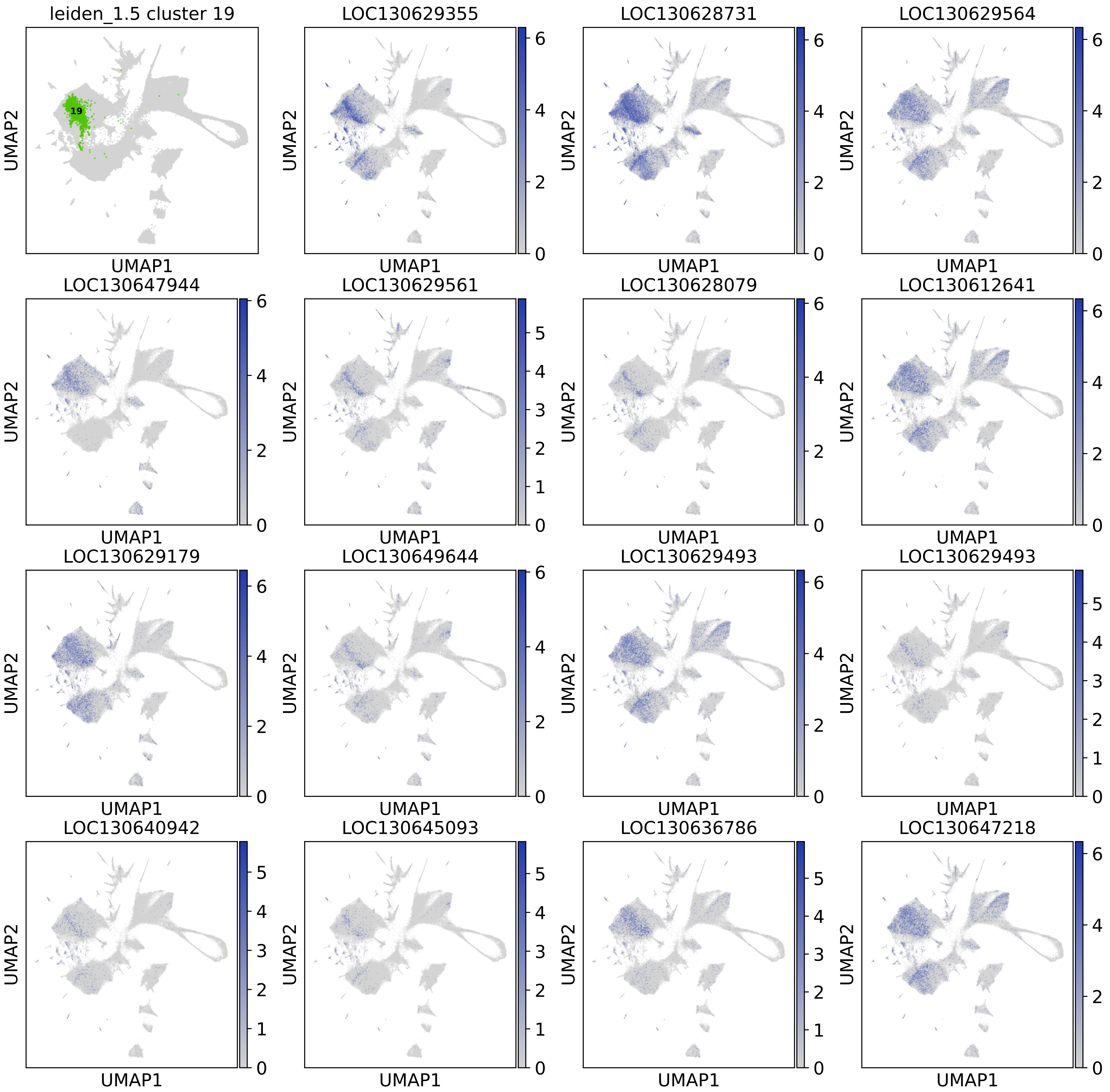
leiden_1.5 cluster 19
19
LOC130629355
LOC130628731
LOC130629564
LOC130647944
LOC130629561
LOC130628079
LOC130612641
LOC130629179
LOC130649644
LOC130629493
LOC130629493
LOC130640942
LOC130645093
LOC130636786
LOC130647218
UMAP1
UMAP2

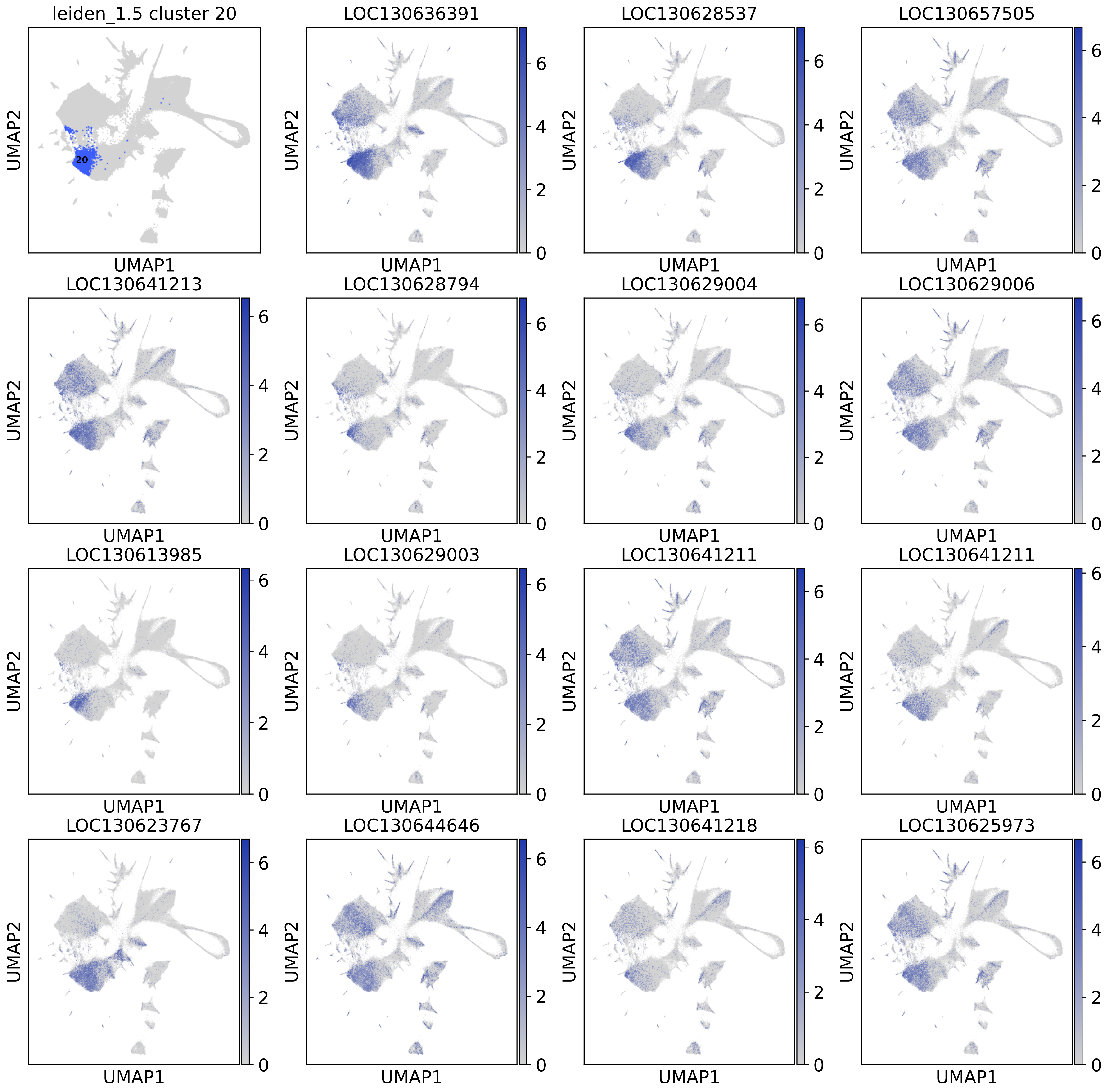
leiden_1.5 cluster 20
20
LOC130636391
LOC130628537
LOC130657505
LOC130641213
LOC130628794
LOC130629004
LOC130629006
LOC130613985
LOC130629003
LOC130641211
LOC130641211
LOC130623767
LOC130644646
LOC130641218
LOC130625973
UMAP1
UMAP2
0
2
4
6

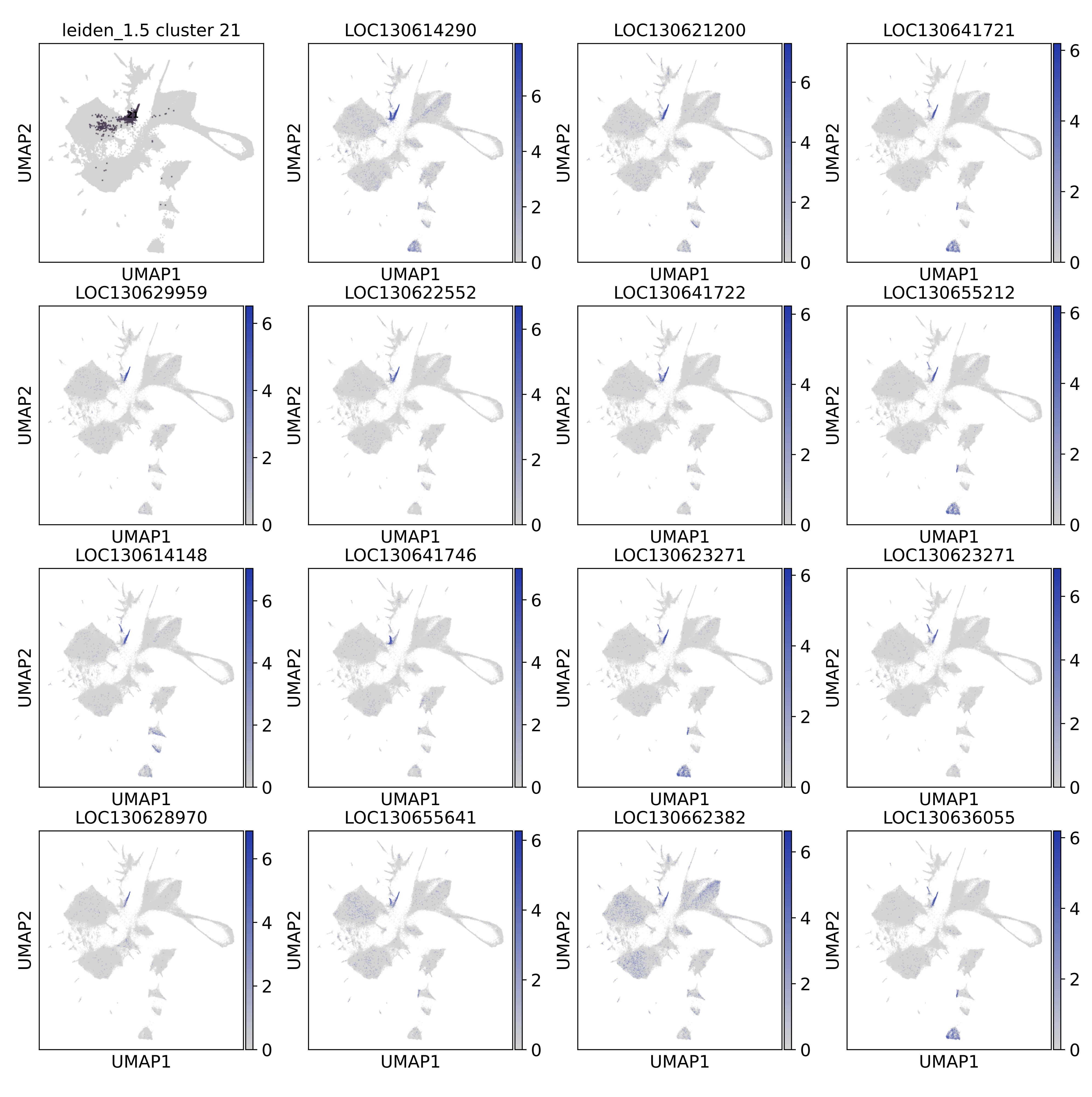
leiden_1.5 cluster 21
21
LOC130614290
LOC130621200
LOC130641721
LOC130629959
LOC130622552
LOC130641722
LOC130655212
LOC130614148
LOC130641746
LOC130623271
LOC130623271
LOC130628970
LOC130655641
LOC130662382
LOC130636055
UMAP1
UMAP2
0
2
4
6

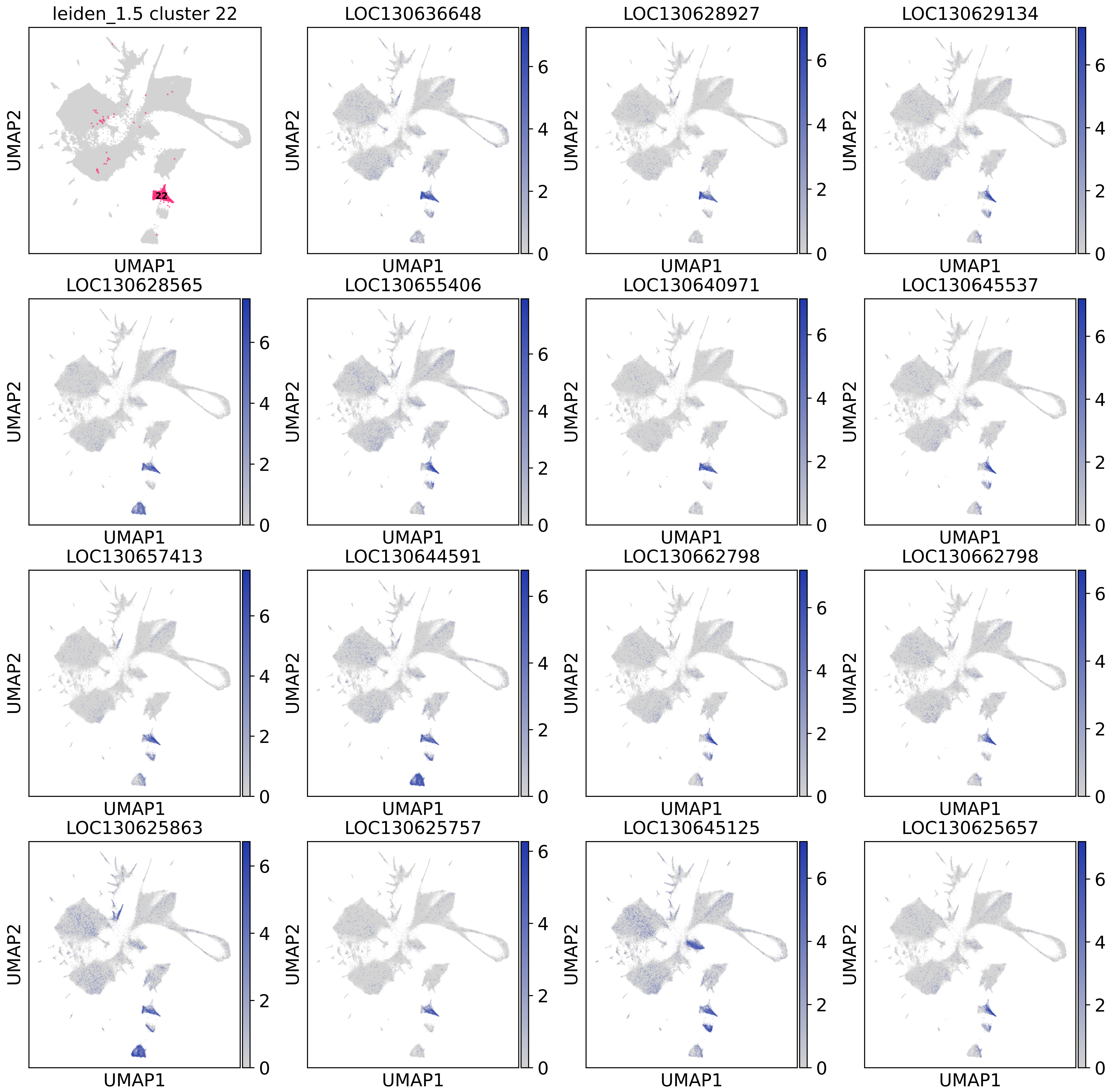
leiden_1.5 cluster 22
22
LOC130636648
LOC130628927
LOC130629134
LOC130628565
LOC130655406
LOC130640971
LOC130645537
LOC130657413
LOC130644591
LOC130662798
LOC130662798
LOC130625863
LOC130625757
LOC130645125
LOC130625657
UMAP1
UMAP2
0
2
4
6

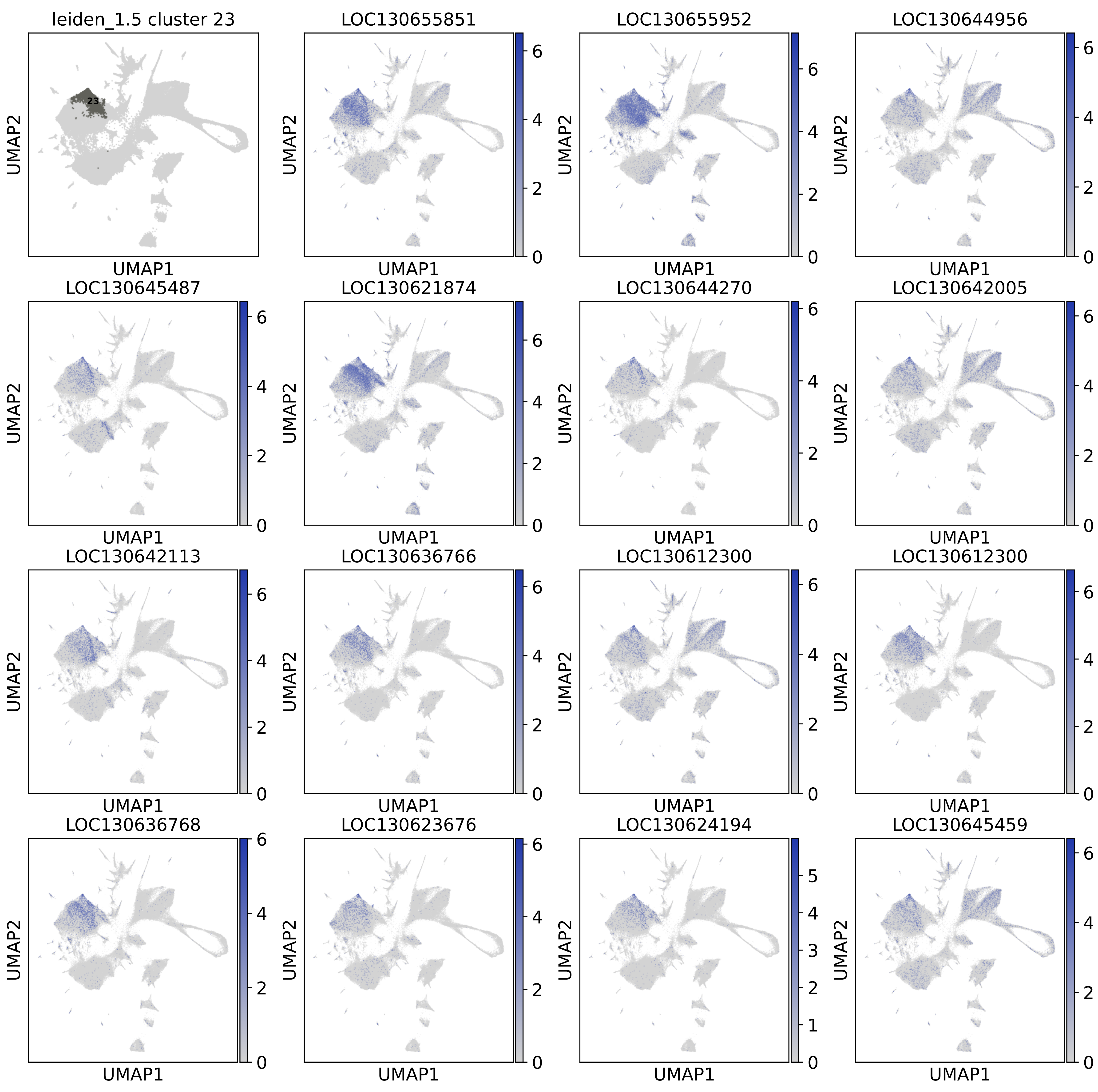

leiden_1.5 cluster 23
23
LOC130655851
LOC130655952
LOC130644956
LOC130645487
LOC130621874
LOC130644270
LOC130642005
LOC130642113
LOC130636766
LOC130612300
LOC130612300
LOC130636768
LOC130623676
LOC130624194
LOC130645459
UMAP1
UMAP2

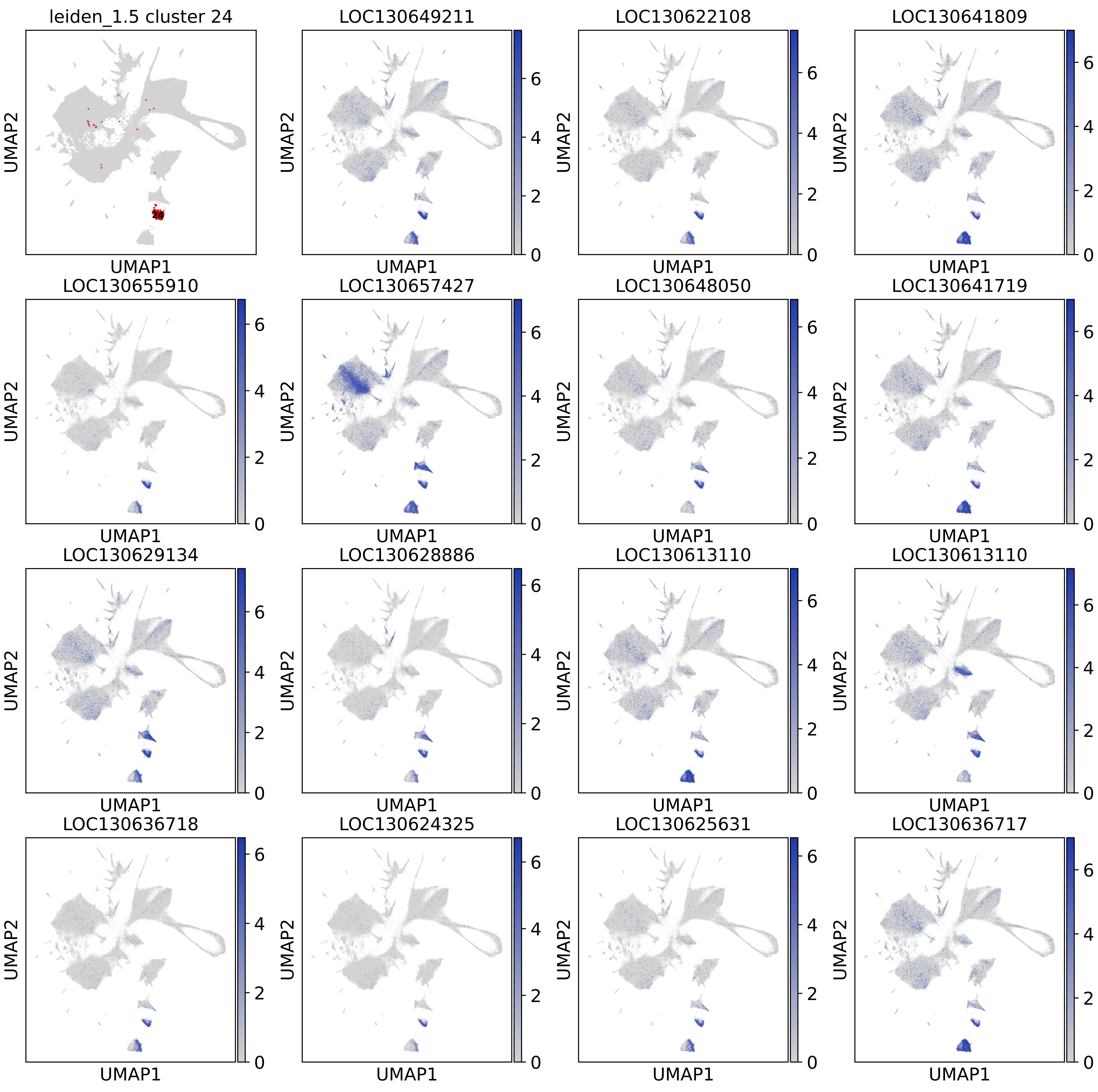

leiden_1.5 cluster 24
24
LOC130649211
LOC130622108
LOC130641809
LOC130655910
LOC130657427
LOC130648050
LOC130641719
LOC130629134
LOC130628886
LOC130613110
LOC130613110
LOC130636718
LOC130624325
LOC130625631
LOC130636717
UMAP1
UMAP2
0
2
4
6

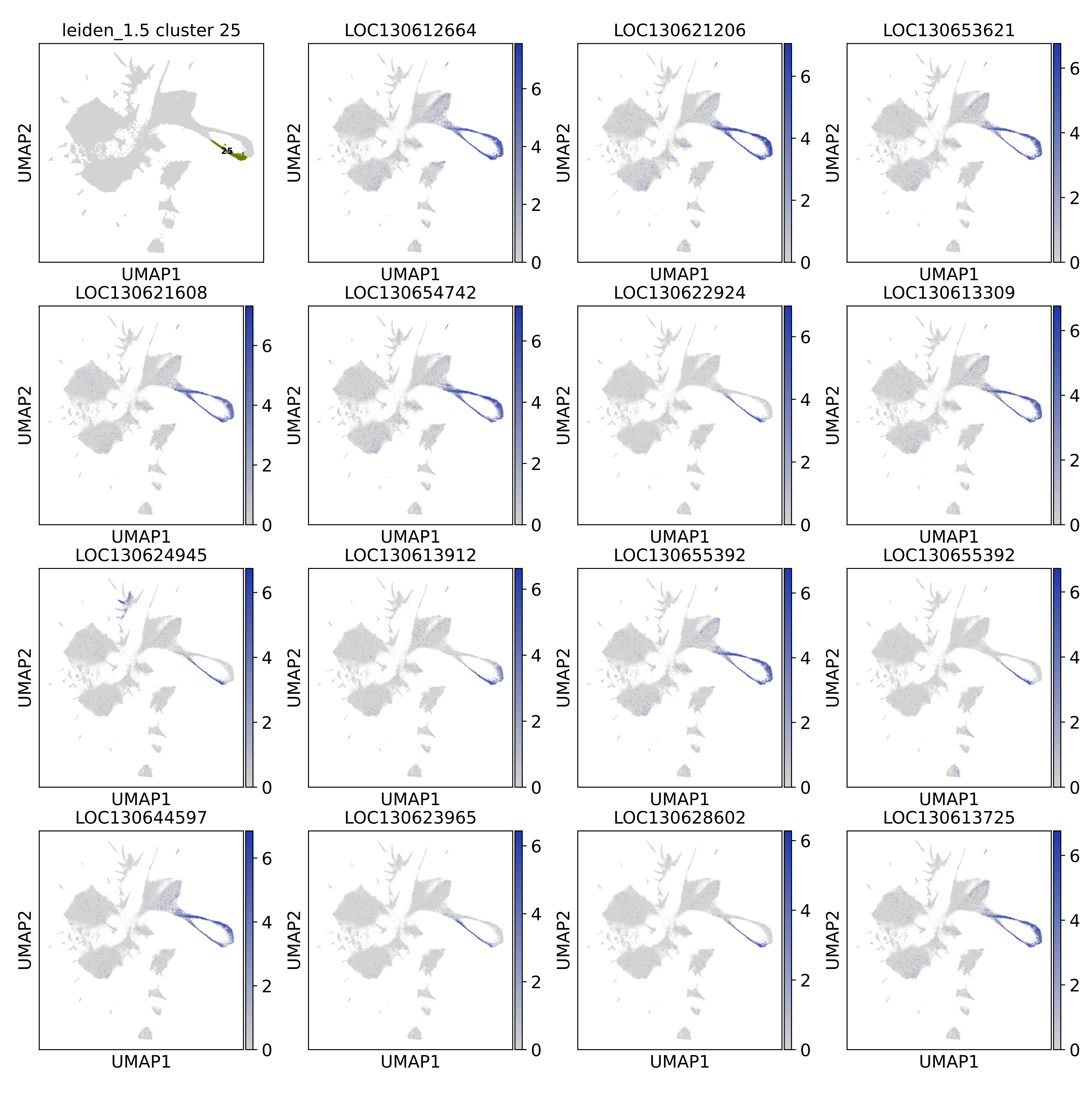
leiden_1.5 cluster 25
25
LOC130612664
LOC130621206
LOC130653621
LOC130621608
LOC130654742
LOC130622924
LOC130613309
LOC130624945
LOC130613912
LOC130655392
LOC130655392
LOC130644597
LOC130623965
LOC130628602
LOC130613725
UMAP1
UMAP2
0
2
4
6

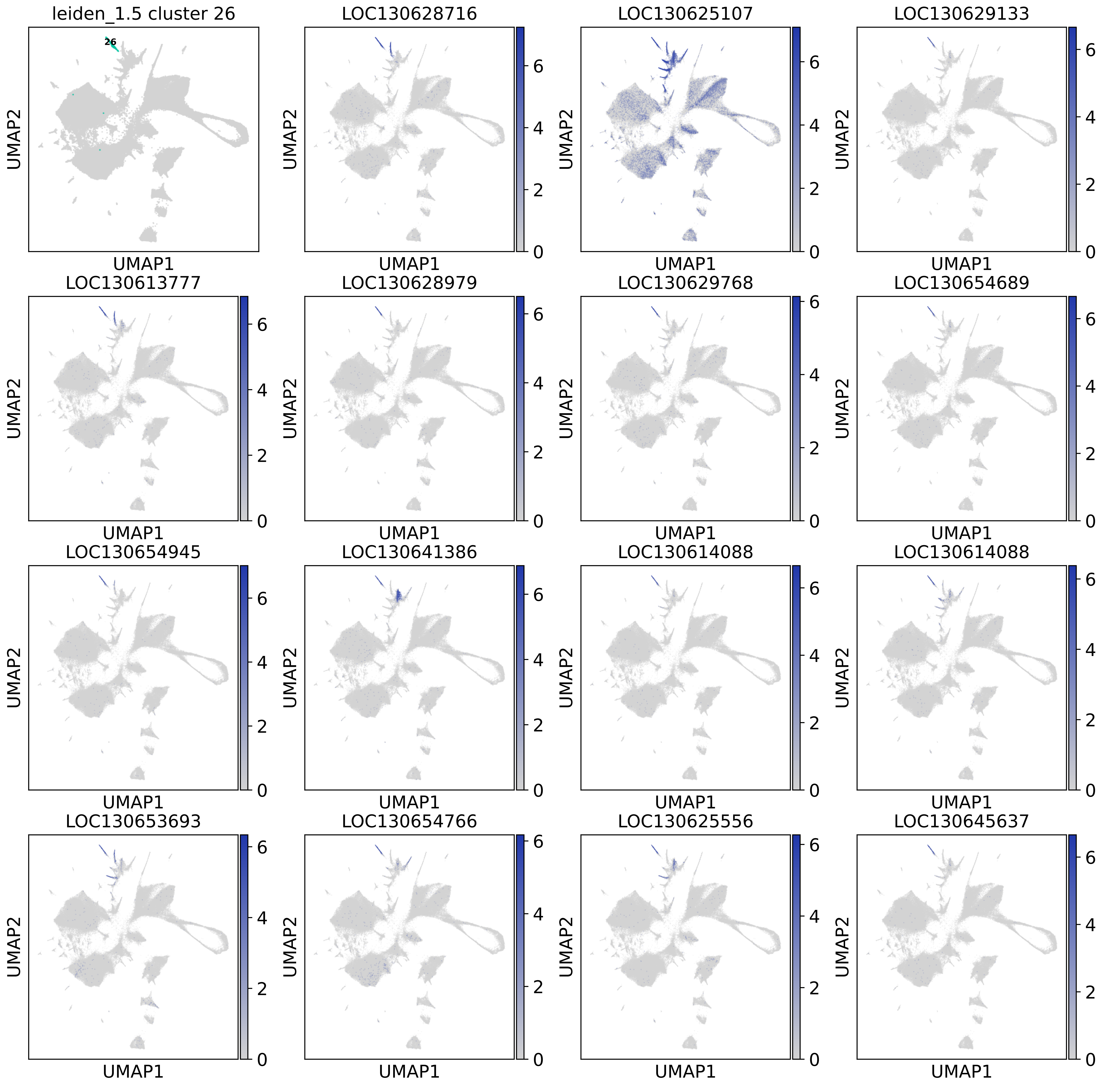
leiden_1.5 cluster 26
26
LOC130628716
LOC130625107
LOC130629133
LOC130613777
LOC130628979
LOC130629768
LOC130654689
LOC130654945
LOC130641386
LOC130614088
LOC130614088
LOC130653693
LOC130654766
LOC130625556
LOC130645637
UMAP1
UMAP2

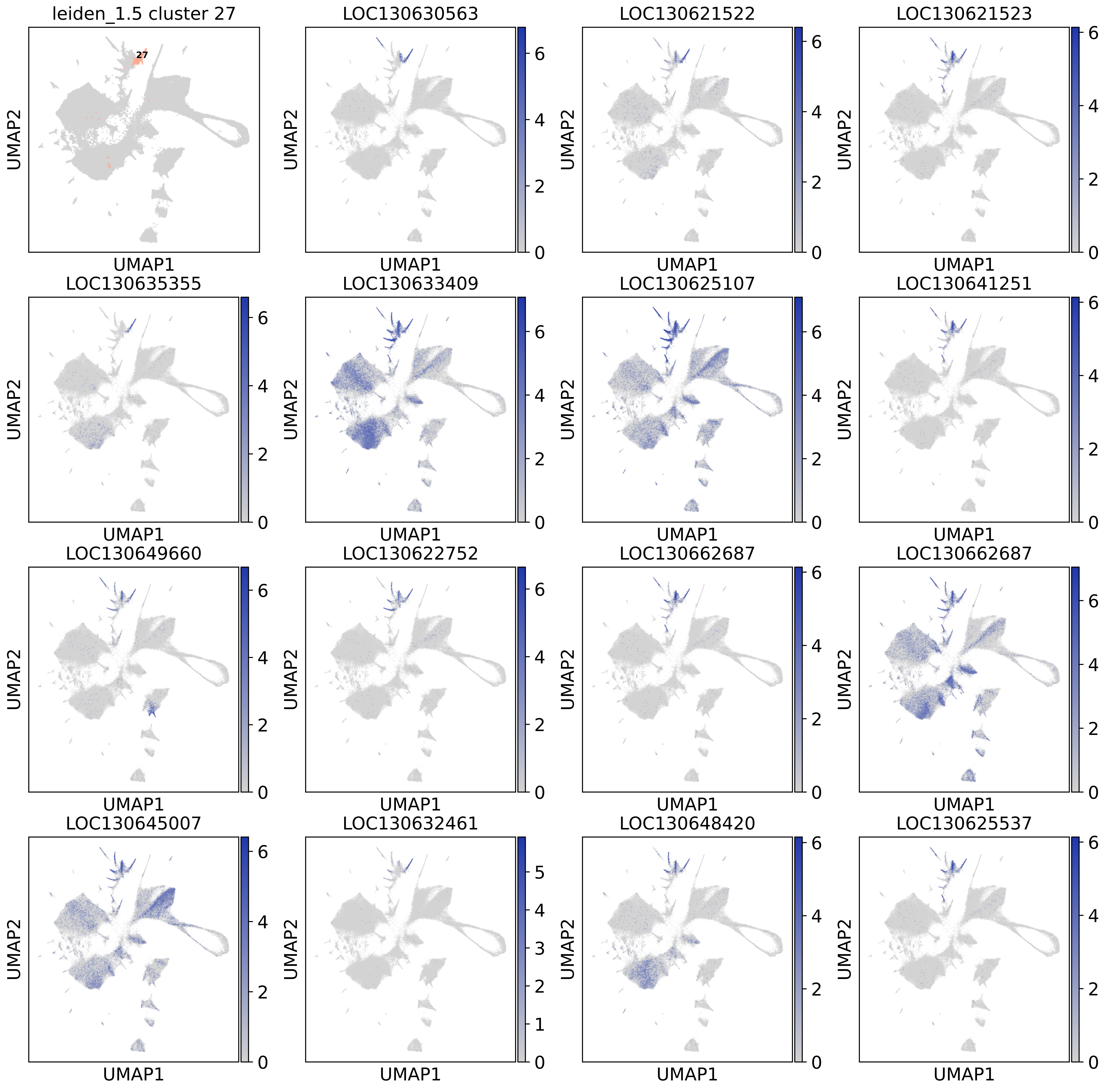
leiden_1.5 cluster 27
LOC130630563
LOC130621522
LOC130621523
LOC130635355
LOC130633409
LOC130625107
LOC130641251
LOC130649660
LOC130622752
LOC130662687
LOC130662687
LOC130645007
LOC130632461
LOC130648420
LOC130625537
UMAP1
UMAP2

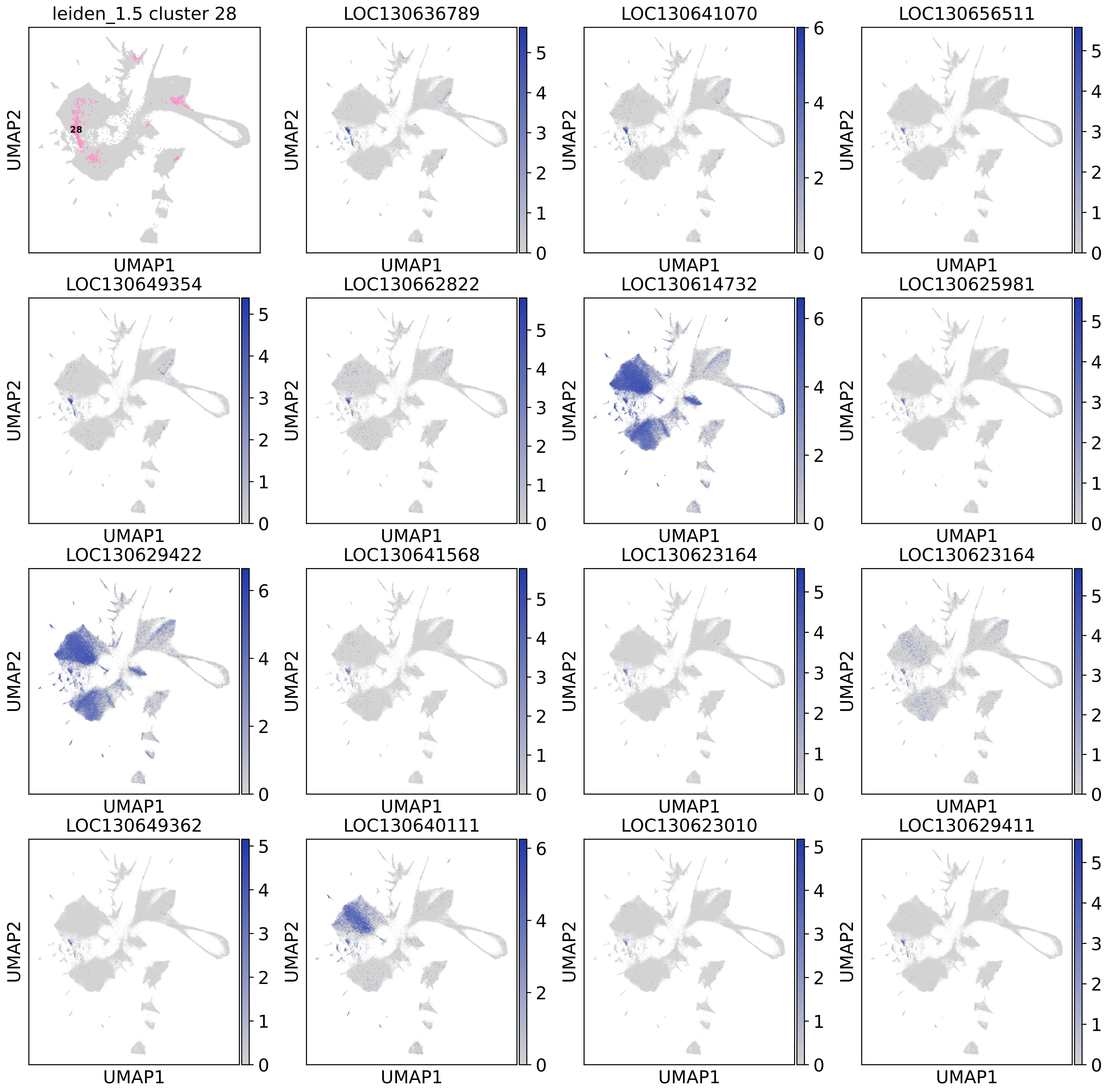
leiden_1.5 cluster 28
28
LOC130636789
LOC130641070
LOC130656511
LOC130649354
LOC130662822
LOC130614732
LOC130625981
LOC130629422
LOC130641568
LOC130623164
LOC130623164
LOC130649362
LOC130640111
LOC130623010
LOC130629411
UMAP1
UMAP2

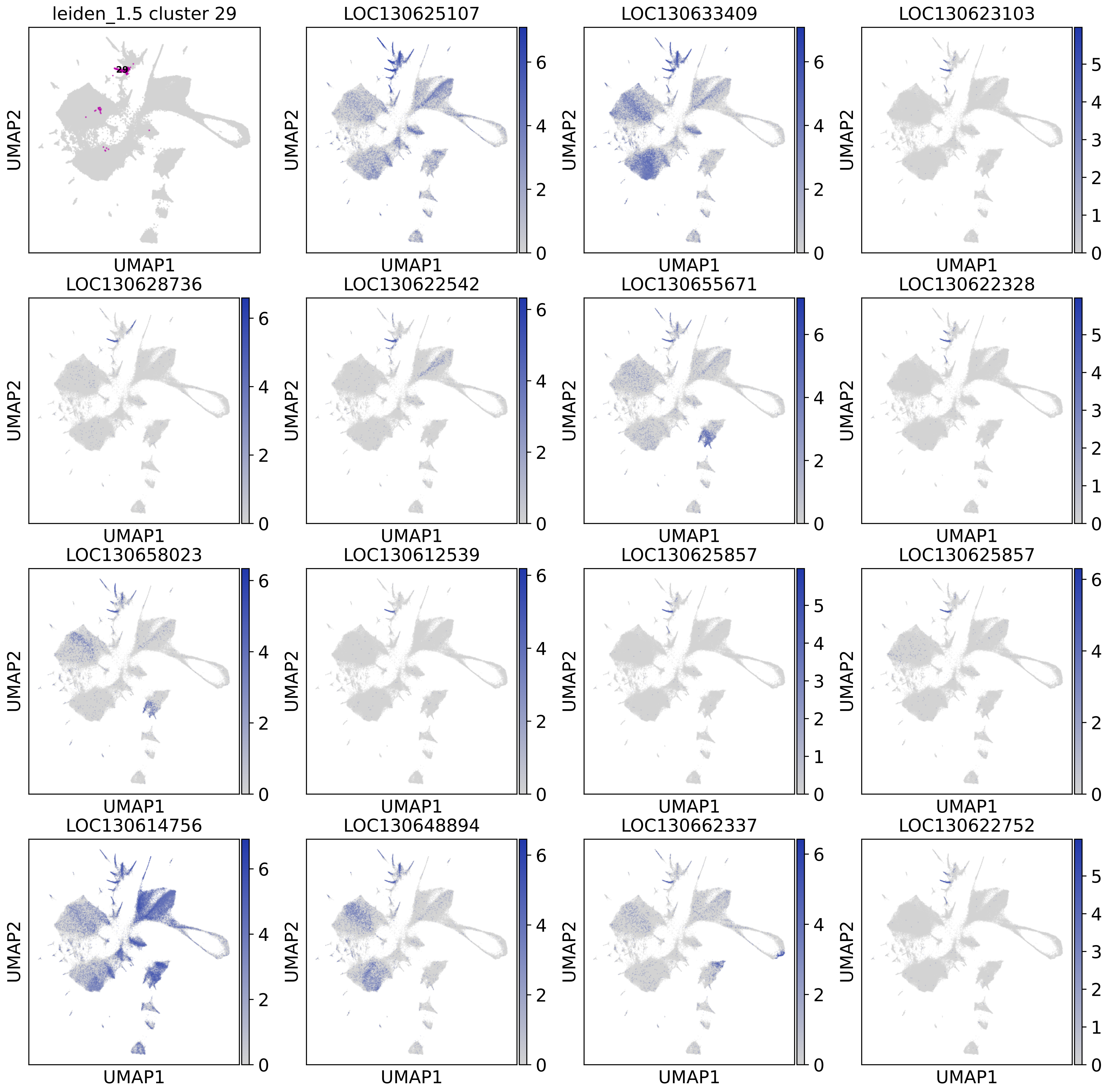

leiden_1.5 cluster 29
29
LOC130625107
LOC130633409
LOC130623103
LOC130628736
LOC130622542
LOC130655671
LOC130622328
LOC130658023
LOC130612539
LOC130625857
LOC130625857
LOC130614756
LOC130648894
LOC130662337
LOC130622752
UMAP1
UMAP2

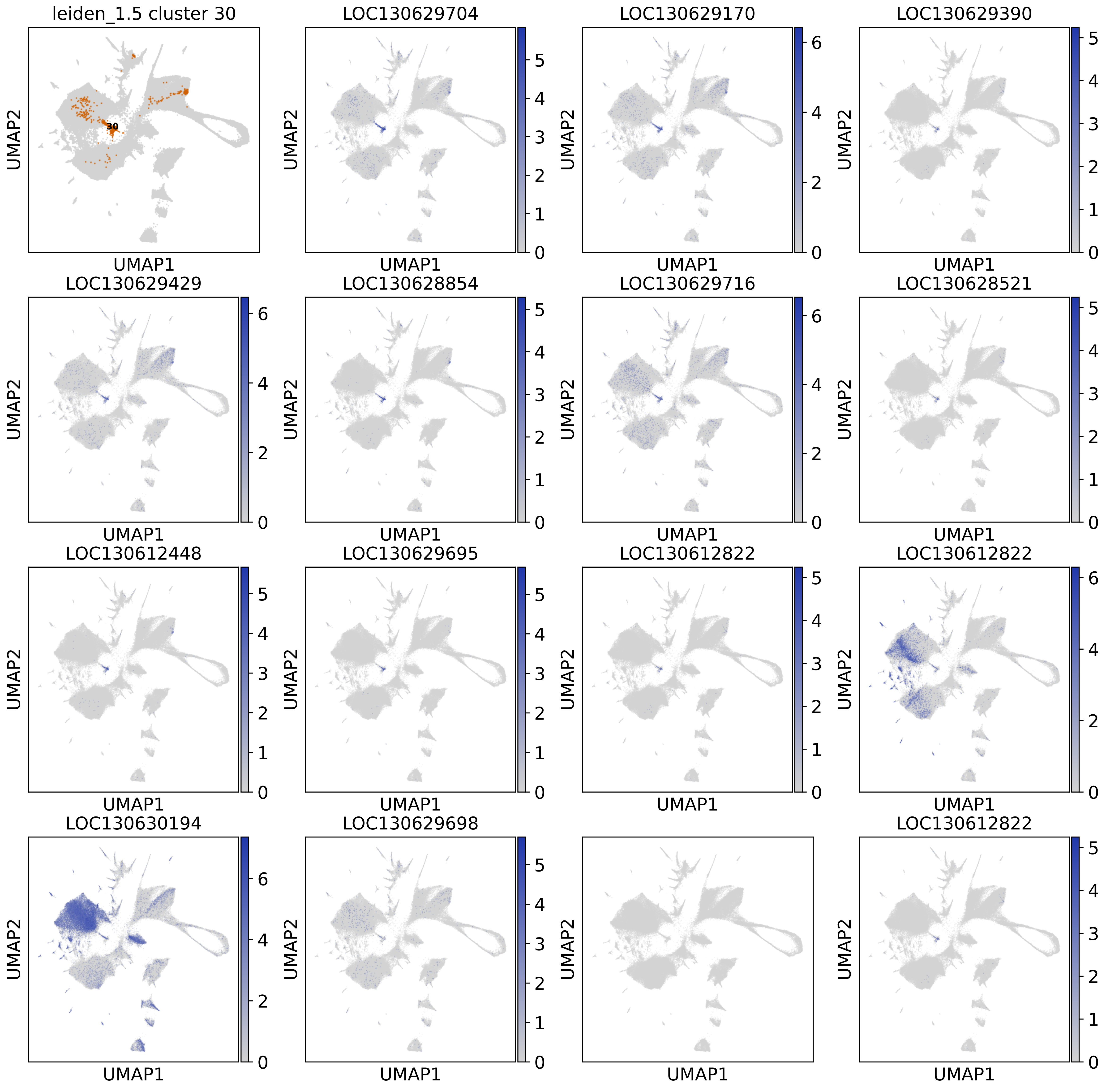
leiden_1.5 cluster 30
30
LOC130629704
LOC130629170
LOC130629390
LOC130629429
LOC130628854
LOC130629716
LOC130628521
LOC130612448
LOC130629695
LOC130612822
LOC130612822
LOC130630194
LOC130629698
LOC130612822
UMAP1
UMAP2

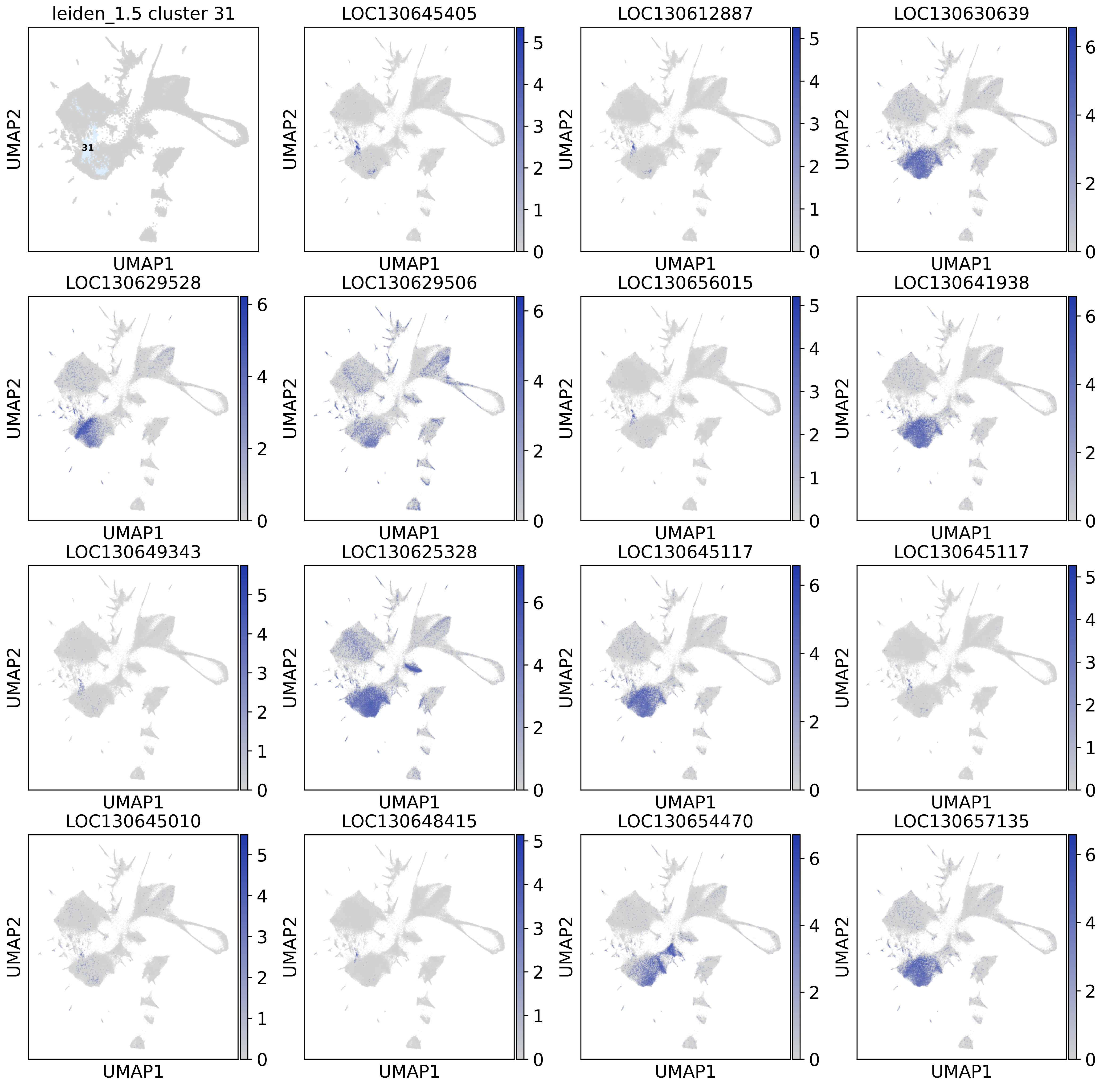

leiden_1.5 cluster 31
31
LOC130645405
LOC130612887
LOC130630639
LOC130629528
LOC130629506
LOC130656015
LOC130641938
LOC130649343
LOC130625328
LOC130645117
LOC130645117
LOC130645010
LOC130648415
LOC130654470
LOC130657135
UMAP1
UMAP2

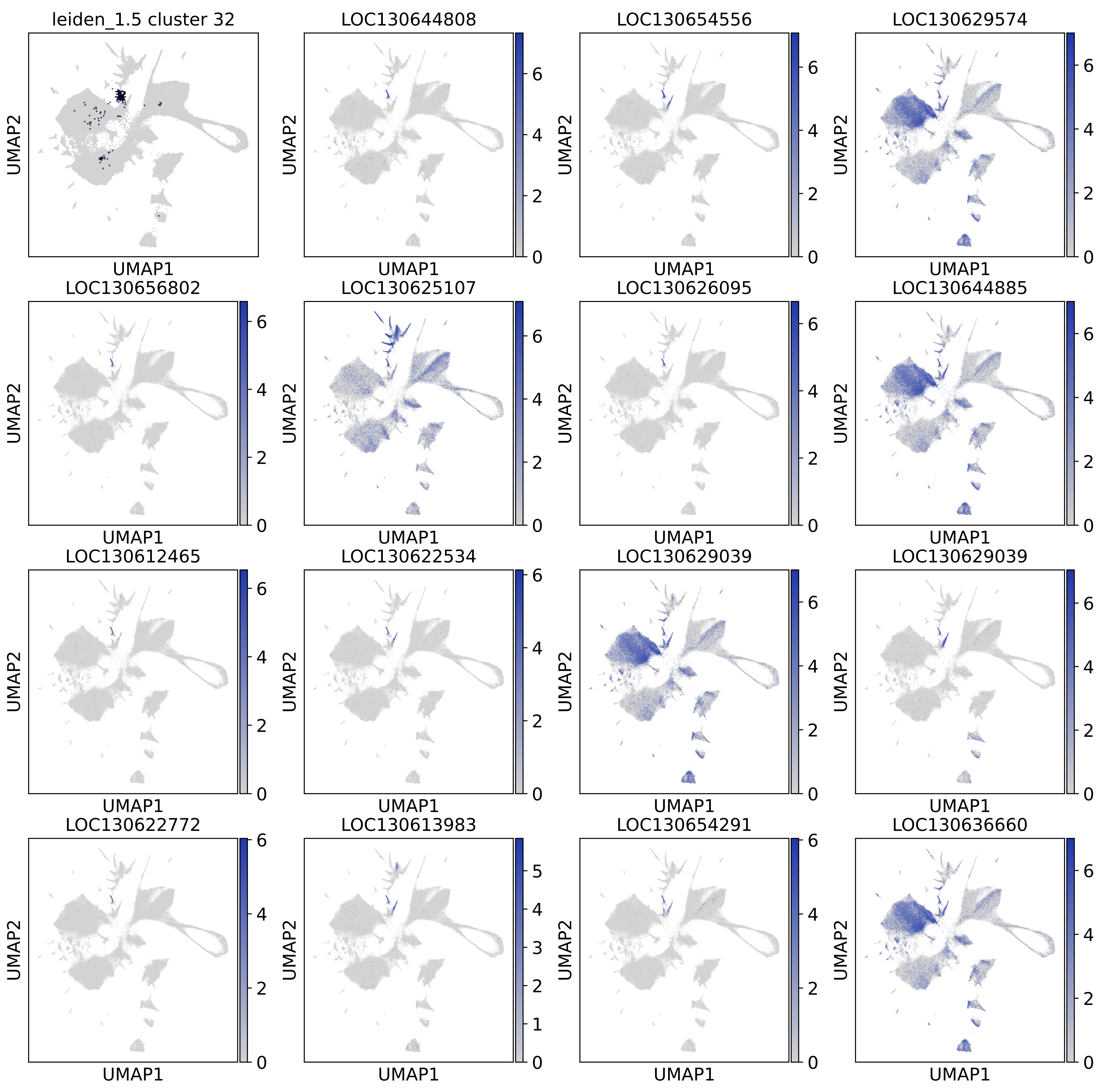
leiden_1.5 cluster 32
32
LOC130644808
LOC130654556
LOC130629574
LOC130656802
LOC130625107
LOC130626095
LOC130644885
LOC130612465
LOC130622534
LOC130629039
LOC130629039
LOC130622772
LOC130613983
LOC130654291
LOC130636660
UMAP1
UMAP2

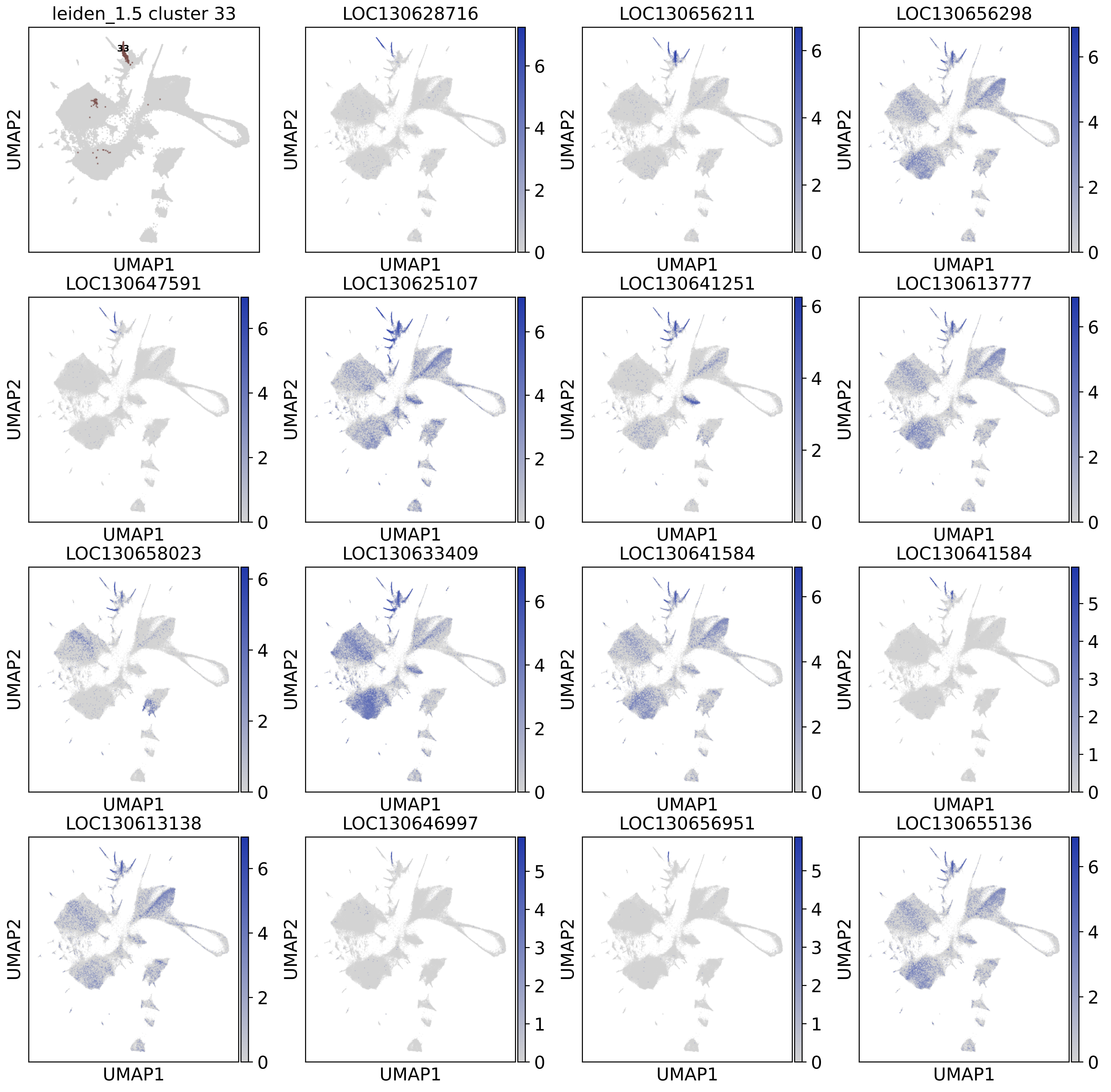
leiden_1.5 cluster 33
33
LOC130628716
LOC130656211
LOC130656298
LOC130647591
LOC130625107
LOC130641251
LOC130613777
LOC130658023
LOC130633409
LOC130641584
LOC130641584
LOC130613138
LOC130646997
LOC130656951
LOC130655136
UMAP1
UMAP2

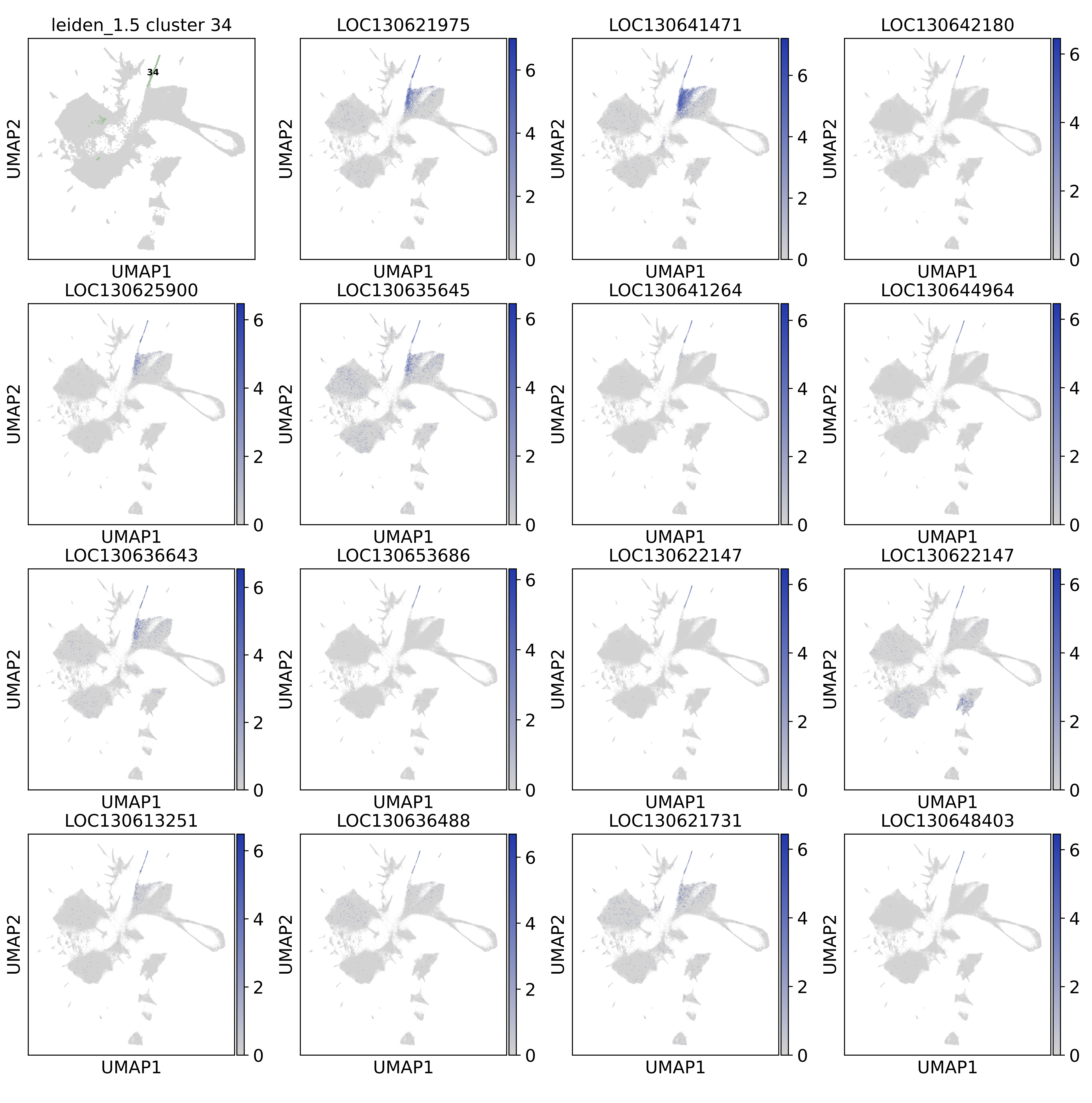

leiden_1.5 cluster 34
34
LOC130621975
LOC130641471
LOC130642180
LOC130625900
LOC130635645
LOC130641264
LOC130644964
LOC130636643
LOC130653686
LOC130622147
LOC130622147
LOC130613251
LOC130636488
LOC130621731
LOC130648403
UMAP1
UMAP2
0
2
4
6

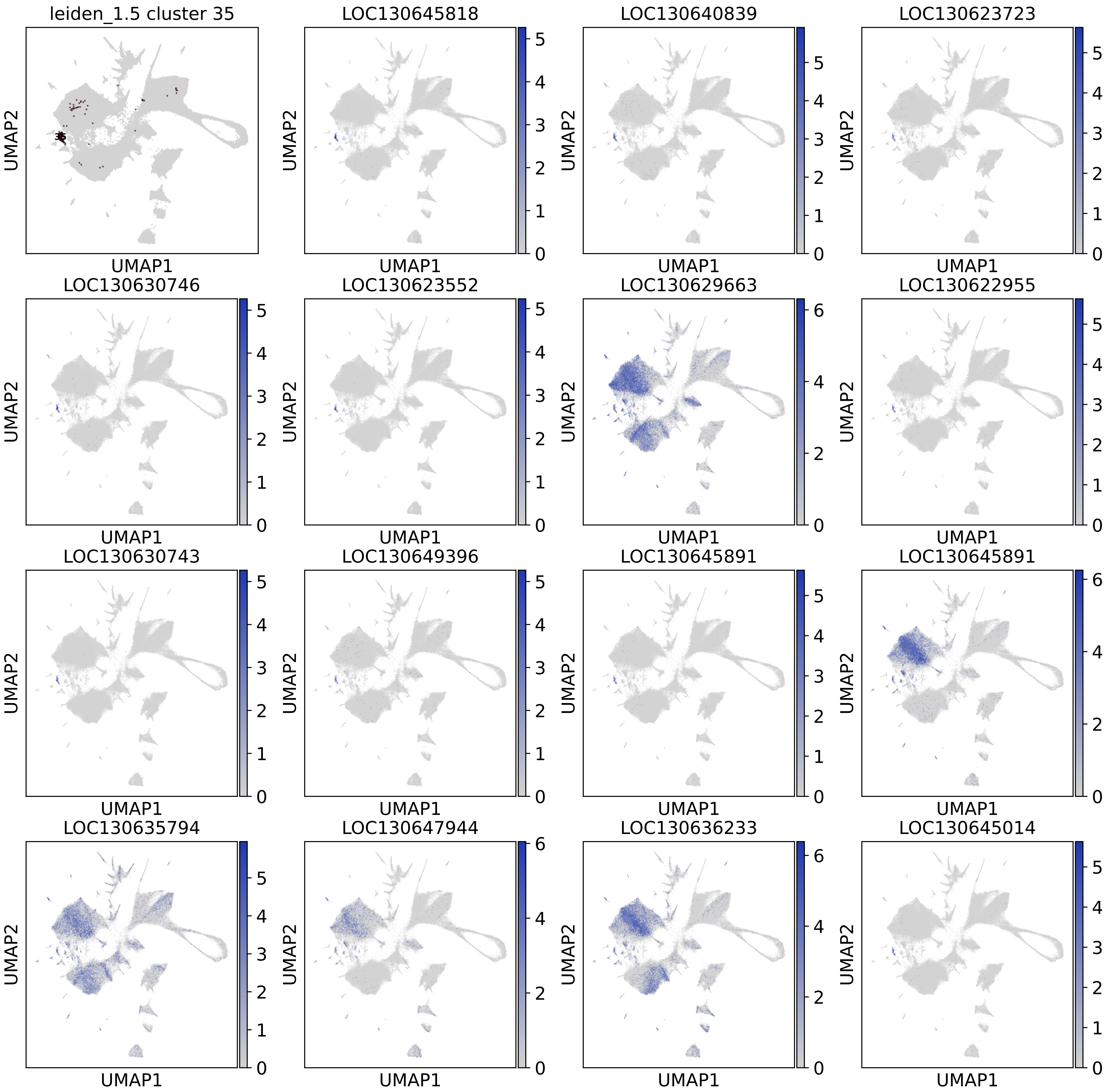

leiden_1.5 cluster 35
35
LOC130645818
LOC130640839
LOC130623723
LOC130630746
LOC130623552
LOC130629663
LOC130622955
LOC130630743
LOC130649396
LOC130645891
LOC130645891
LOC130635794
LOC130647944
LOC130636233
LOC130645014
UMAP1
UMAP2

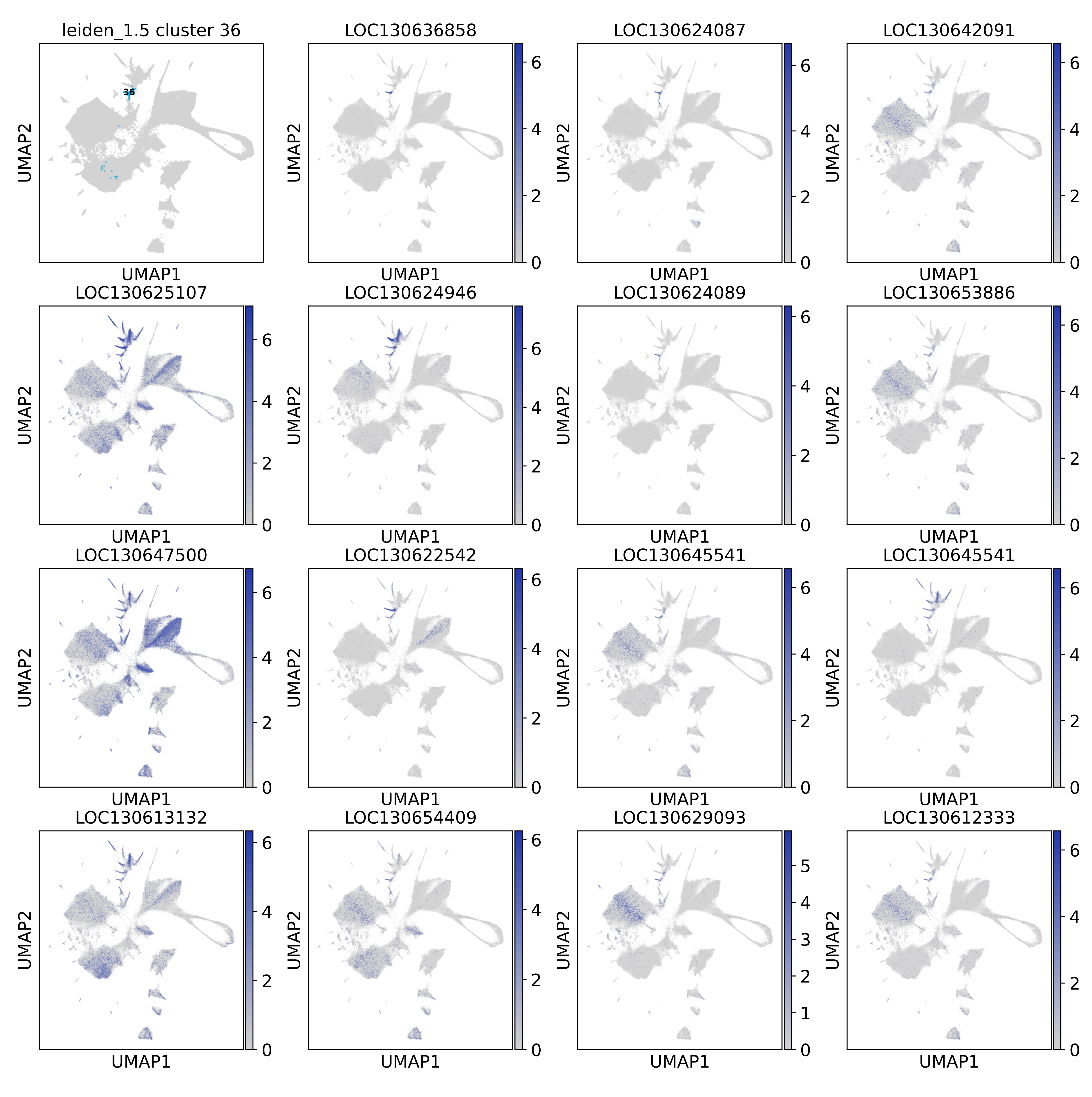
leiden_1.5 cluster 36
36
LOC130636858
LOC130624087
LOC130642091
LOC130625107
LOC130624946
LOC130624089
LOC130653886
LOC130647500
LOC130622542
LOC130645541
LOC130645541
LOC130613132
LOC130654409
LOC130629093
LOC130612333
UMAP1
UMAP2
0
2
4
6

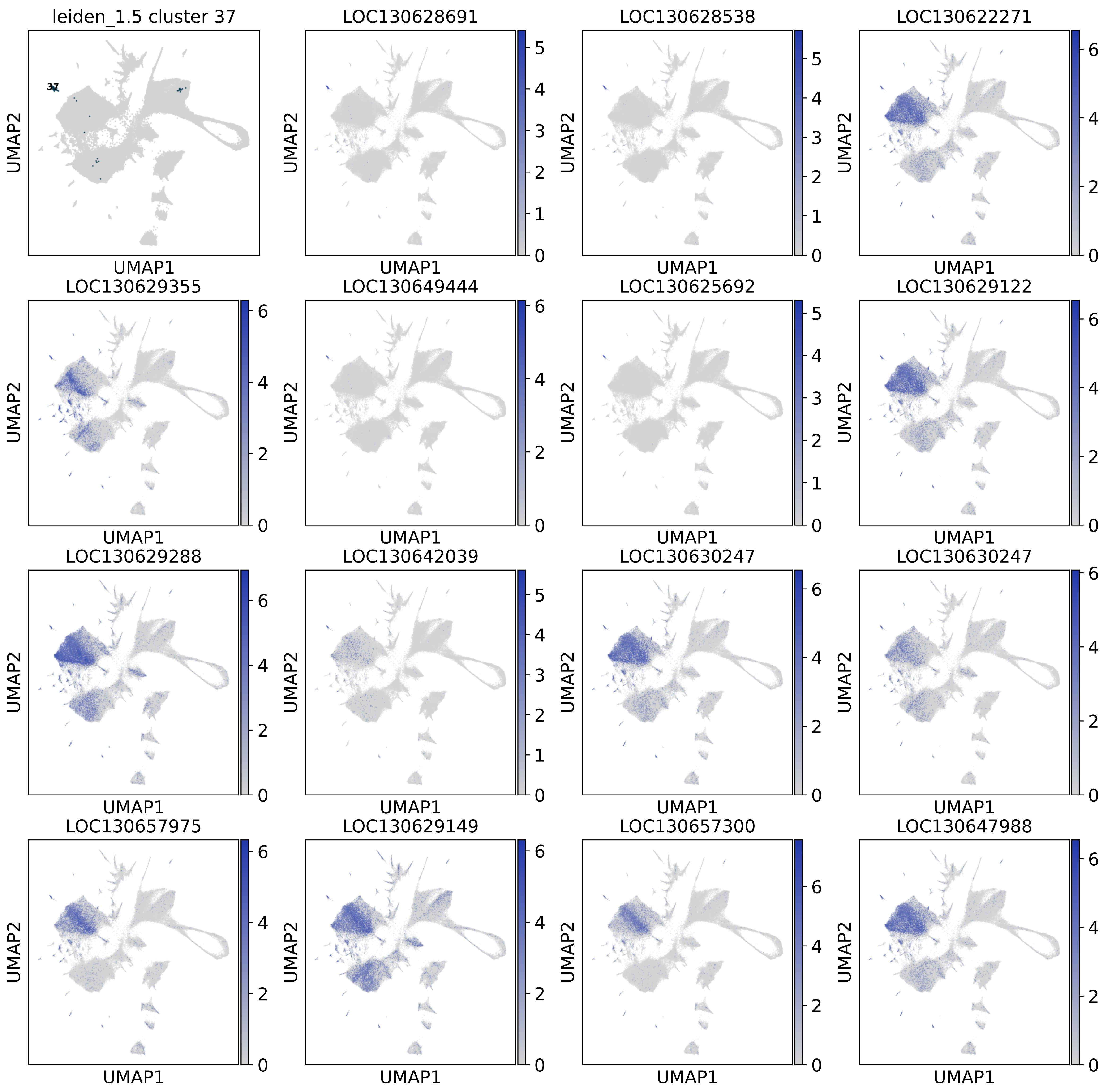
leiden_1.5 cluster 37
37
LOC130628691
LOC130628538
LOC130622271
LOC130629355
LOC130649444
LOC130625692
LOC130629122
LOC130629288
LOC130642039
LOC130630247
LOC130630247
LOC130657975
LOC130629149
LOC130657300
LOC130647988
UMAP1
UMAP2

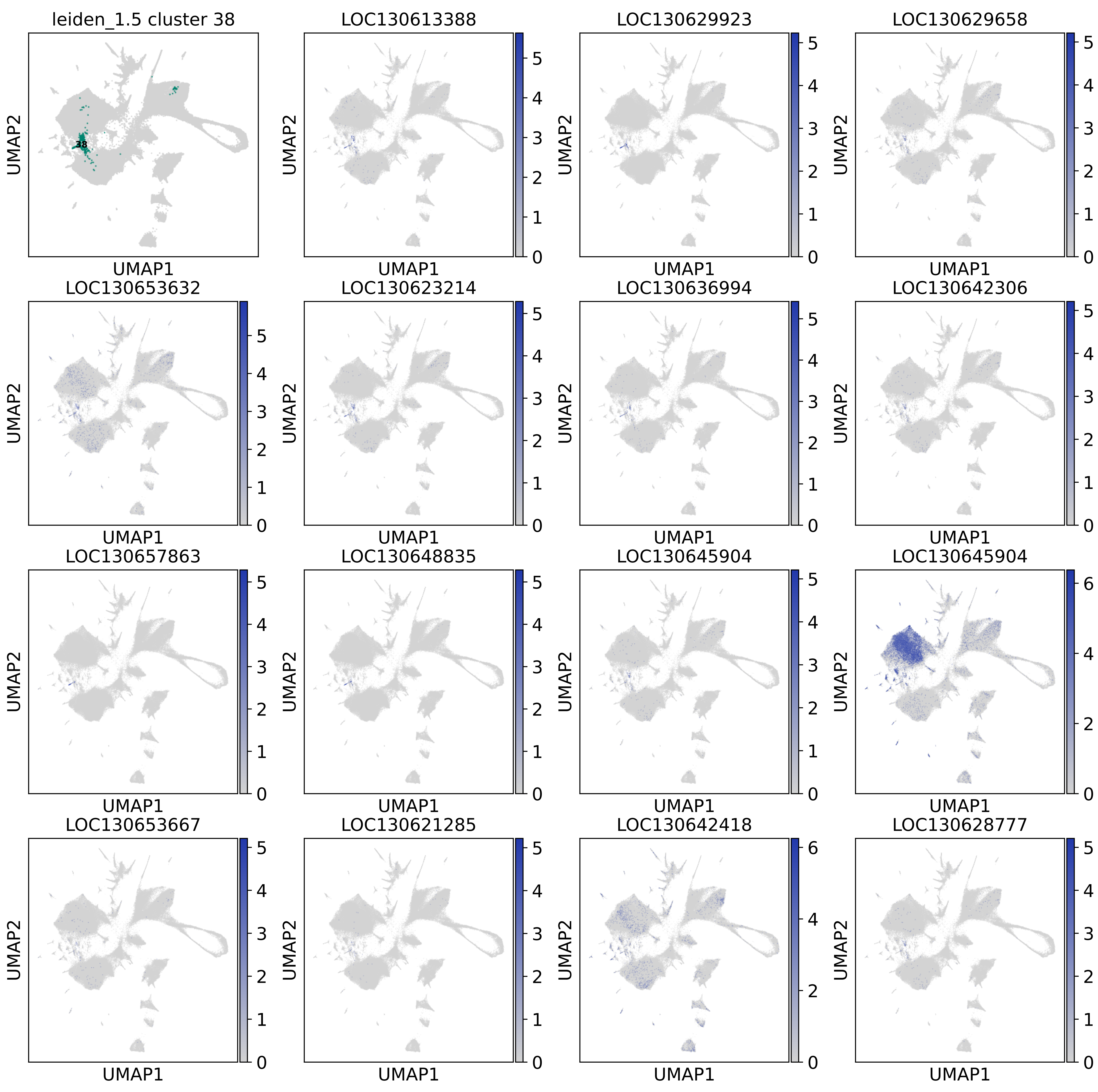

leiden_1.5 cluster 38
38
LOC130613388
LOC130629923
LOC130629658
LOC130653632
LOC130623214
LOC130636994
LOC130642306
LOC130657863
LOC130648835
LOC130645904
LOC130645904
LOC130653667
LOC130621285
LOC130642418
LOC130628777
UMAP1
UMAP2
0
1
2
3
4
5
6

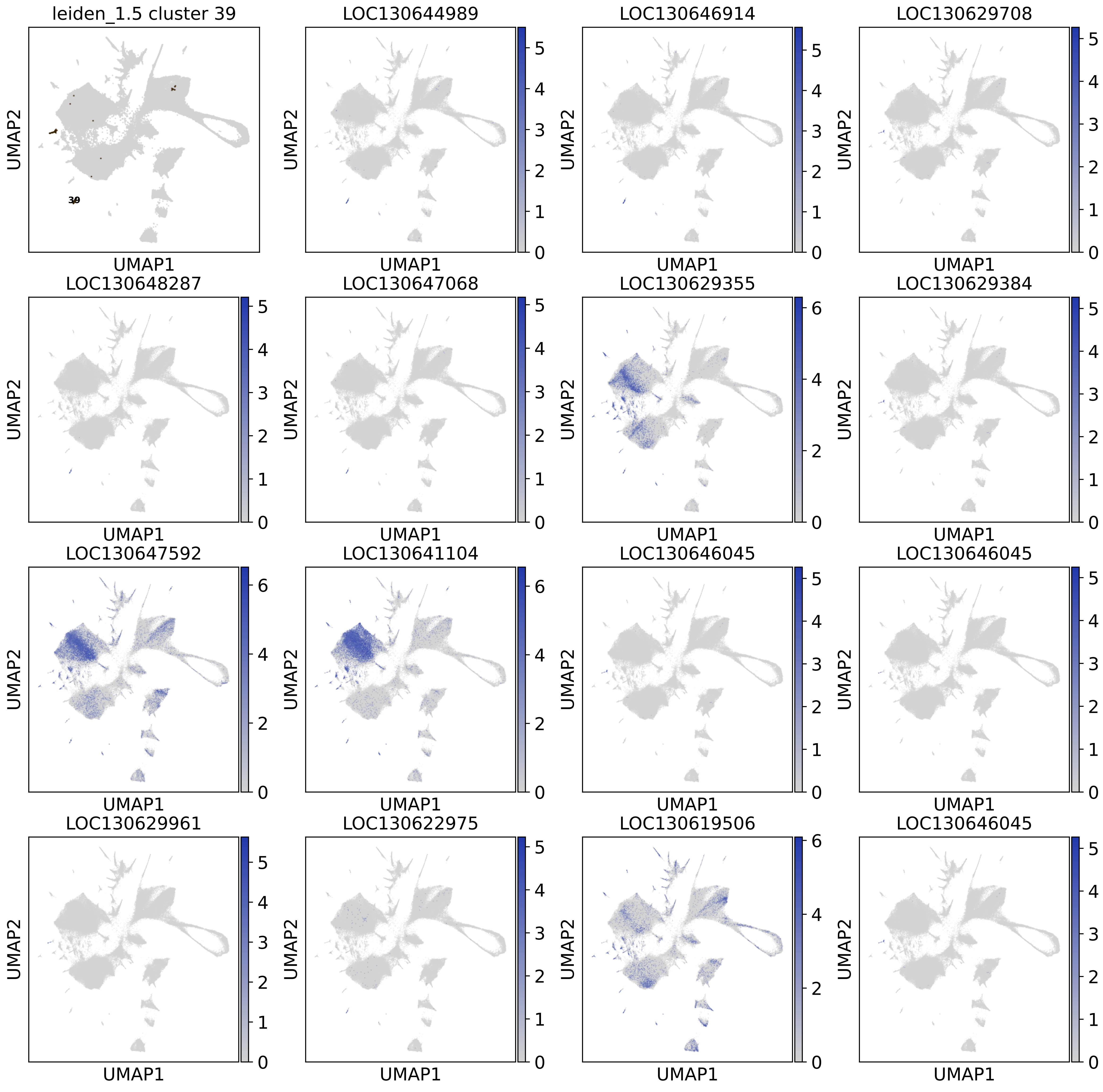
leiden_1.5 cluster 39
39
LOC130644989
LOC130646914
LOC130629708
LOC130648287
LOC130647068
LOC130629355
LOC130629384
LOC130647592
LOC130641104
LOC130646045
LOC130646045
LOC130629961
LOC130622975
LOC130619506
LOC130646045
UMAP1
UMAP2

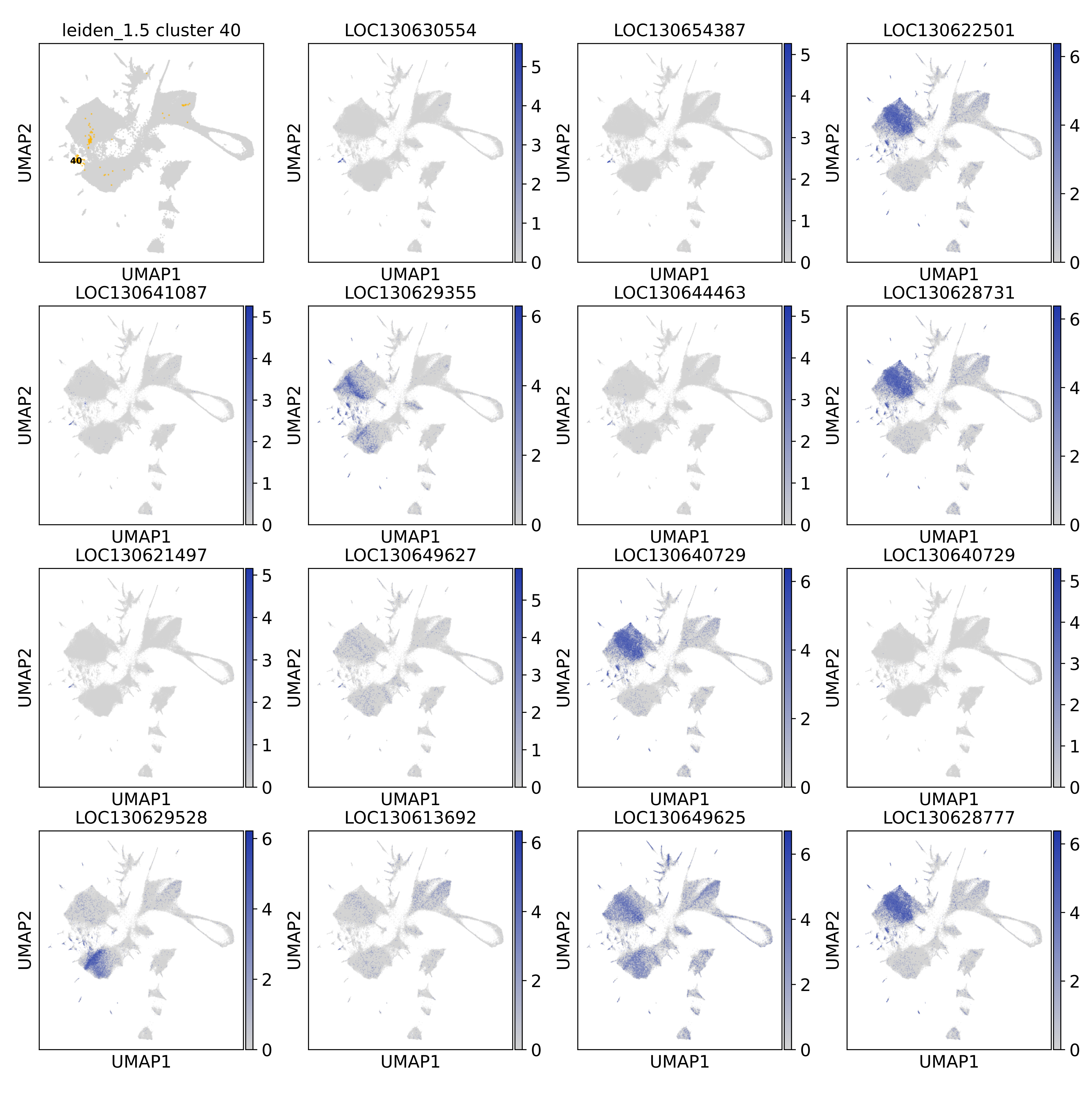
leiden_1.5 cluster 40
40
LOC130630554
LOC130654387
LOC130622501
LOC130641087
LOC130629355
LOC130644463
LOC130628731
LOC130621497
LOC130649627
LOC130640729
LOC130640729
LOC130629528
LOC130613692
LOC130649625
LOC130628777
UMAP1
UMAP2

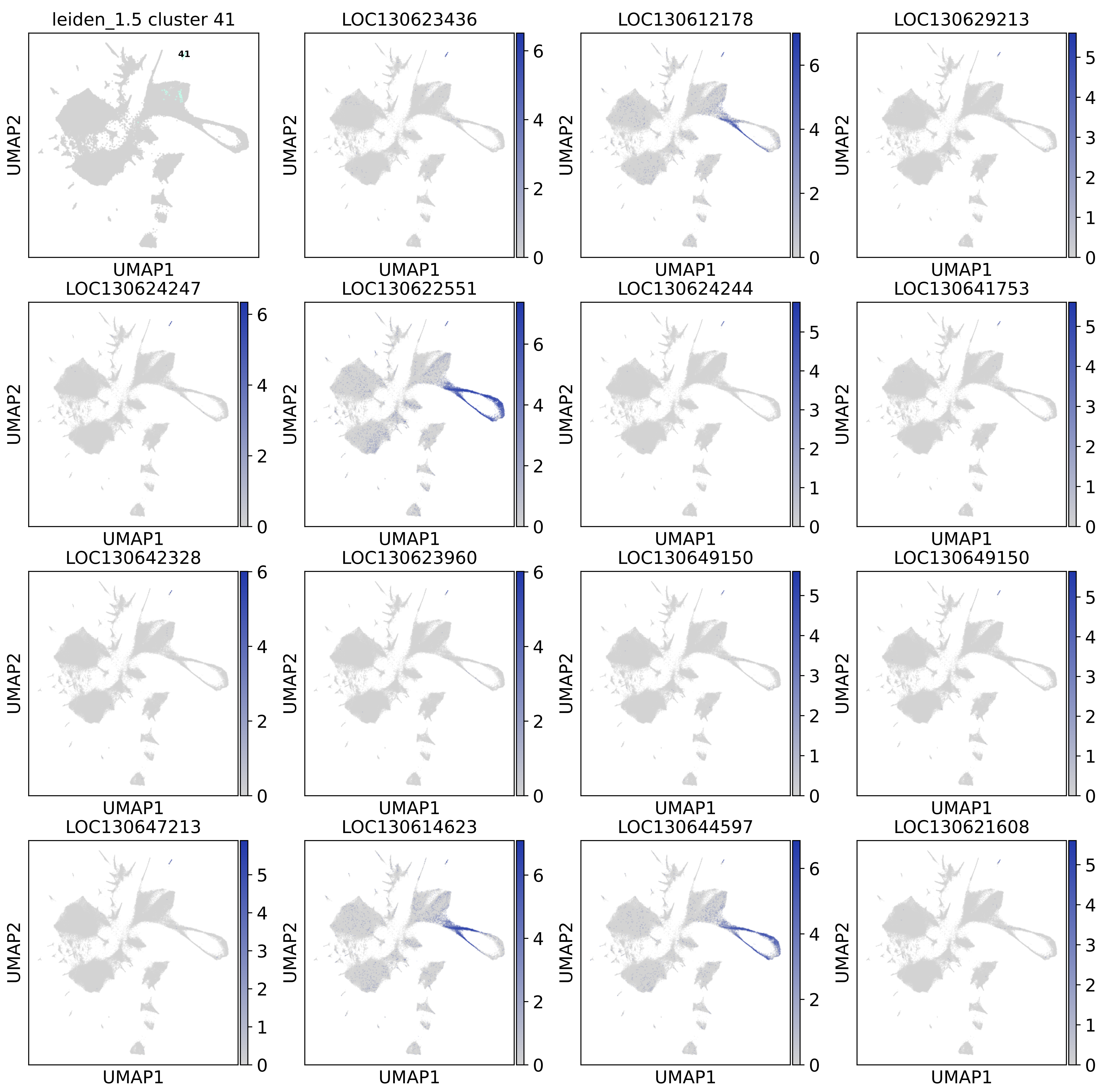
leiden_1.5 cluster 41
41
LOC130623436
LOC130612178
LOC130629213
LOC130624247
LOC130622551
LOC130624244
LOC130641753
LOC130642328
LOC130623960
LOC130649150
LOC130649150
LOC130647213
LOC130614623
LOC130644597
LOC130621608
UMAP1
UMAP2

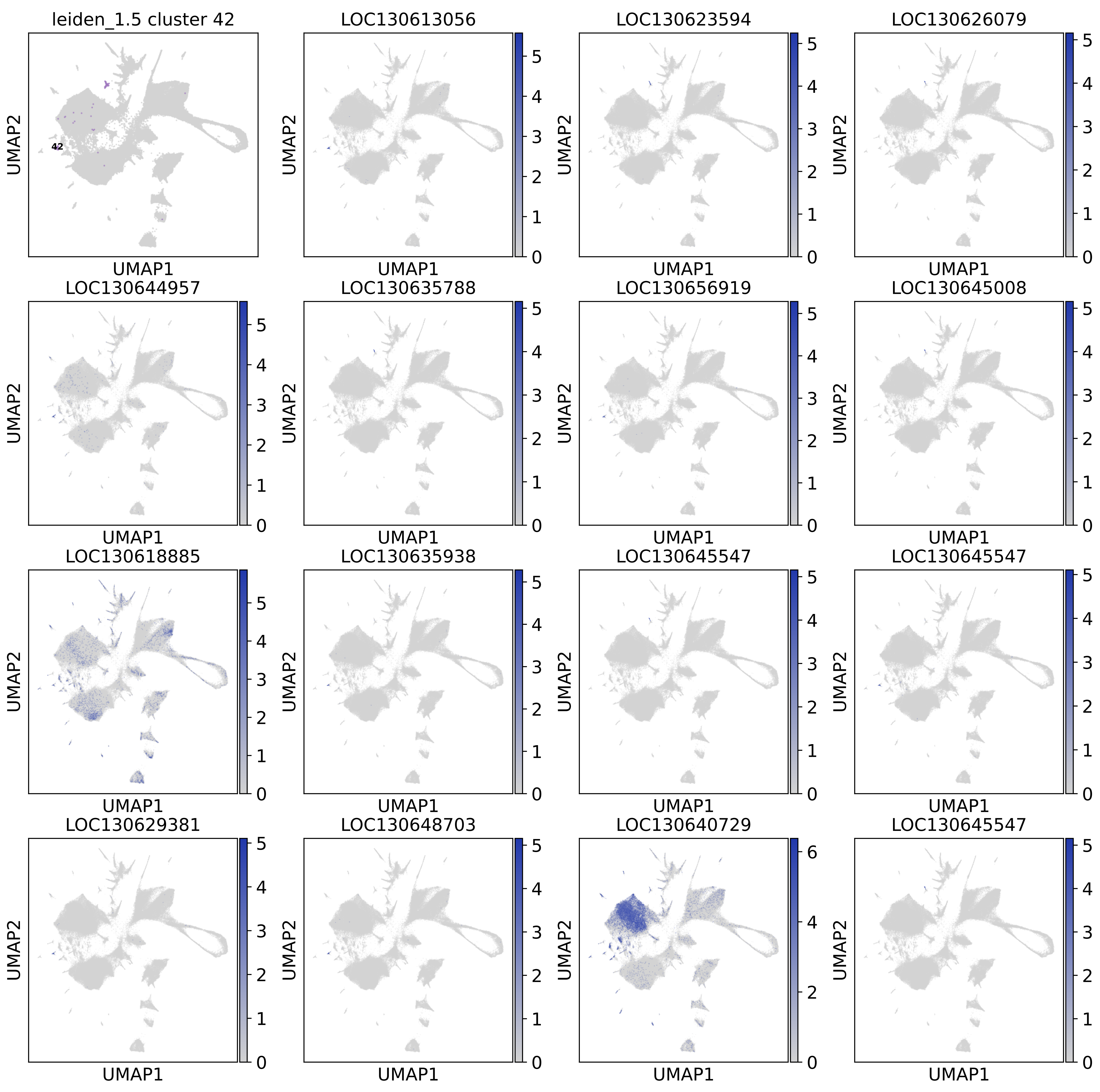

leiden_1.5 cluster 42
42
LOC130613056
LOC130623594
LOC130626079
LOC130644957
LOC130635788
LOC130656919
LOC130645008
LOC130618885
LOC130635938
LOC130645547
LOC130645547
LOC130629381
LOC130648703
LOC130640729
LOC130645547
UMAP1
UMAP2

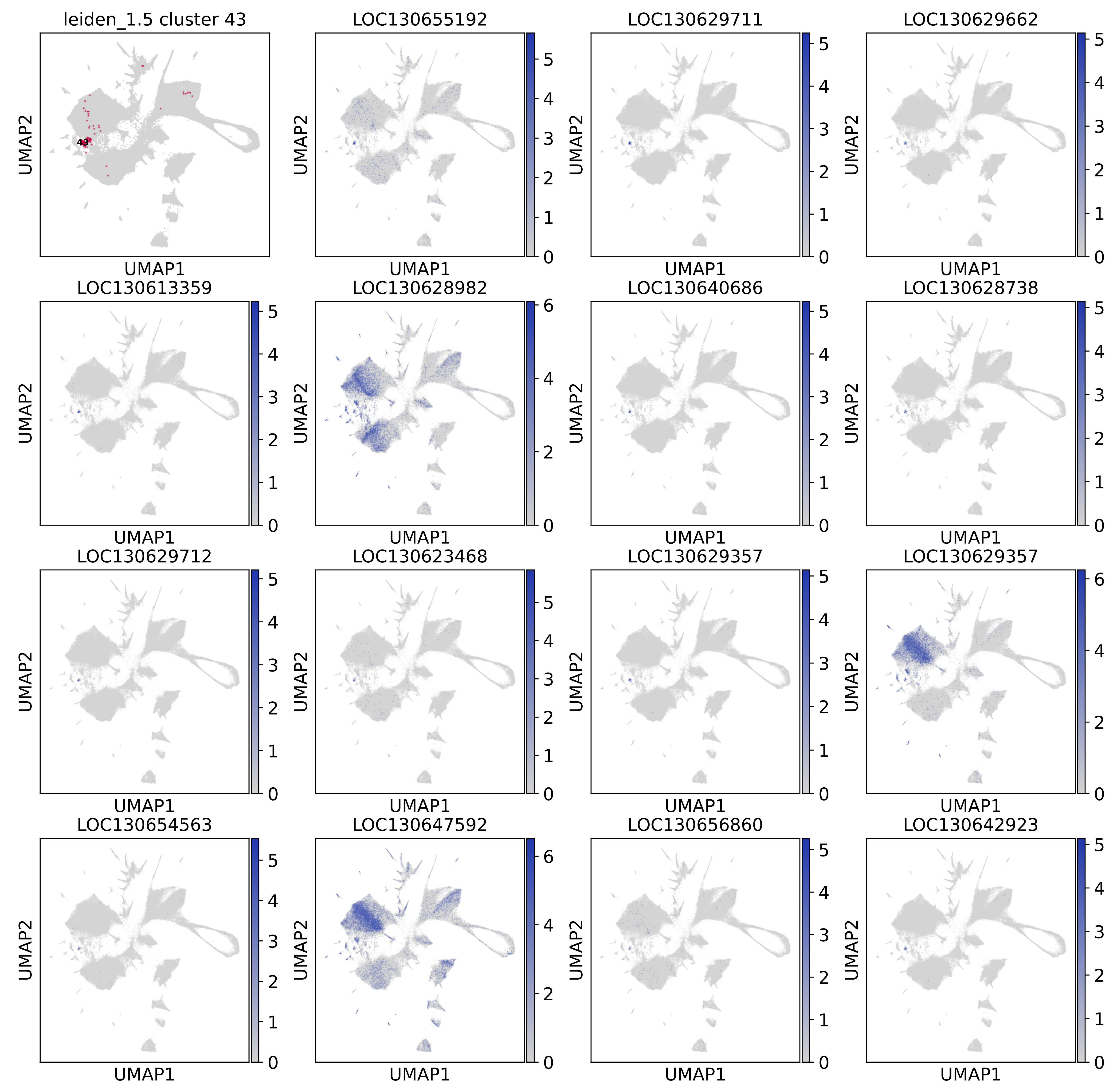
leiden_1.5 cluster 43
43
LOC130655192
LOC130629711
LOC130629662
LOC130613359
LOC130628982
LOC130640686
LOC130628738
LOC130629712
LOC130623468
LOC130629357
LOC130629357
LOC130654563
LOC130647592
LOC130656860
LOC130642923
UMAP1
UMAP2

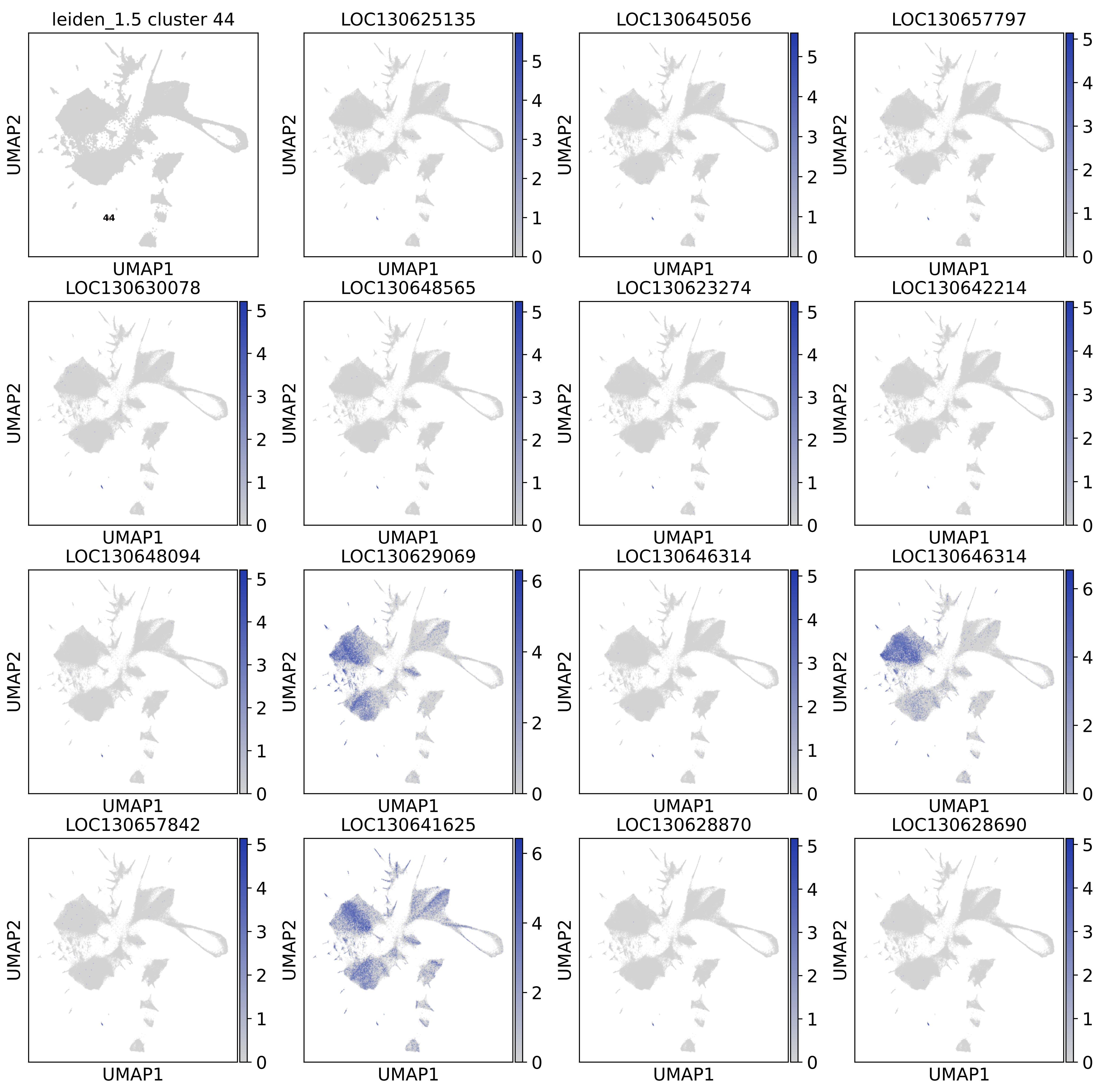
leiden_1.5 cluster 44
44
LOC130625135
LOC130645056
LOC130657797
LOC130630078
LOC130648565
LOC130623274
LOC130642214
LOC130648094
LOC130629069
LOC130646314
LOC130646314
LOC130657842
LOC130641625
LOC130628870
LOC130628690
UMAP1
UMAP2
0
1
2
3
4
5
6

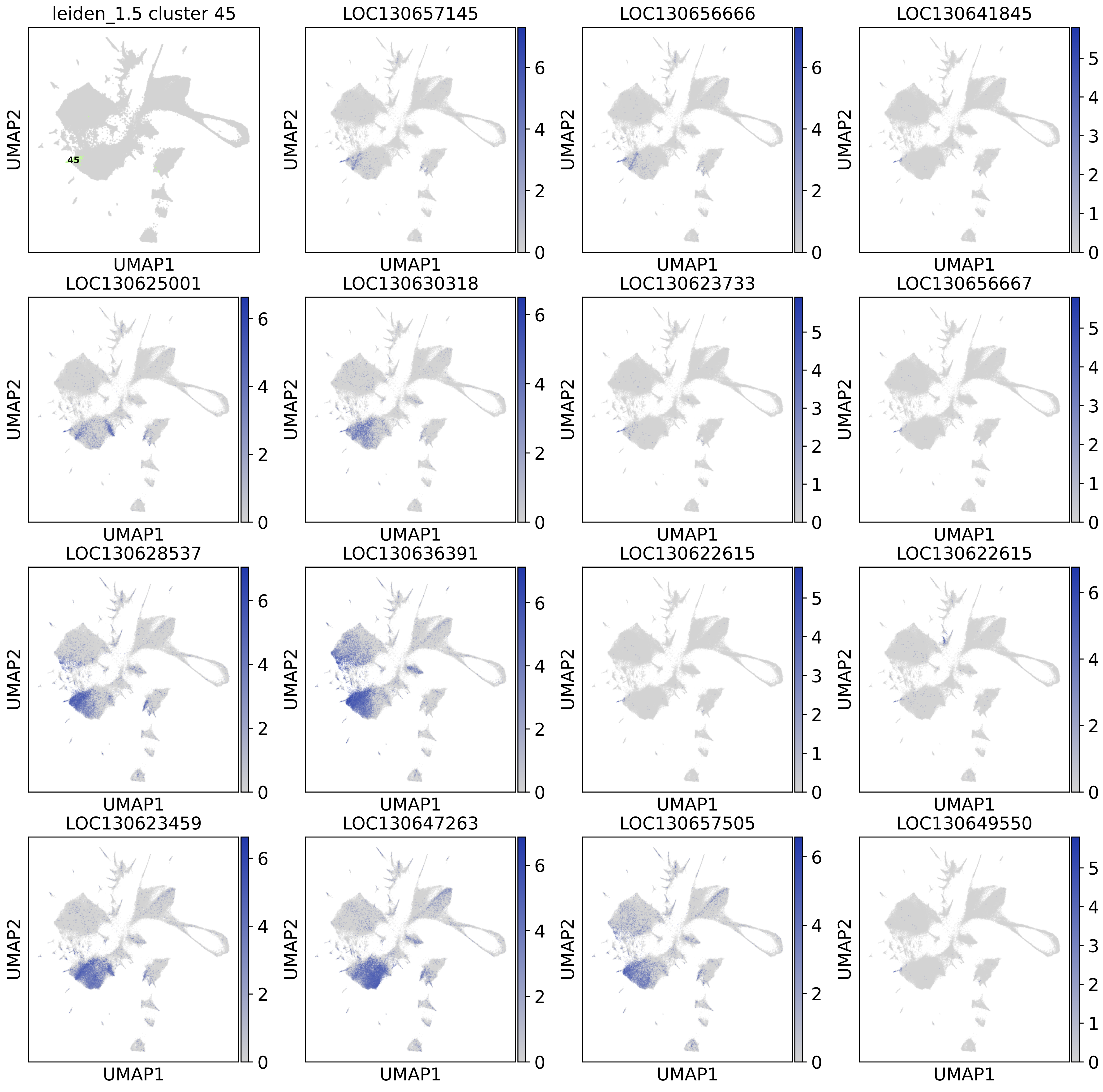
leiden_1.5 cluster 45
45
LOC130657145
LOC130656666
LOC130641845
LOC130625001
LOC130630318
LOC130623733
LOC130656667
LOC130628537
LOC130636391
LOC130622615
LOC130622615
LOC130623459
LOC130647263
LOC130657505
LOC130649550
UMAP1
UMAP2

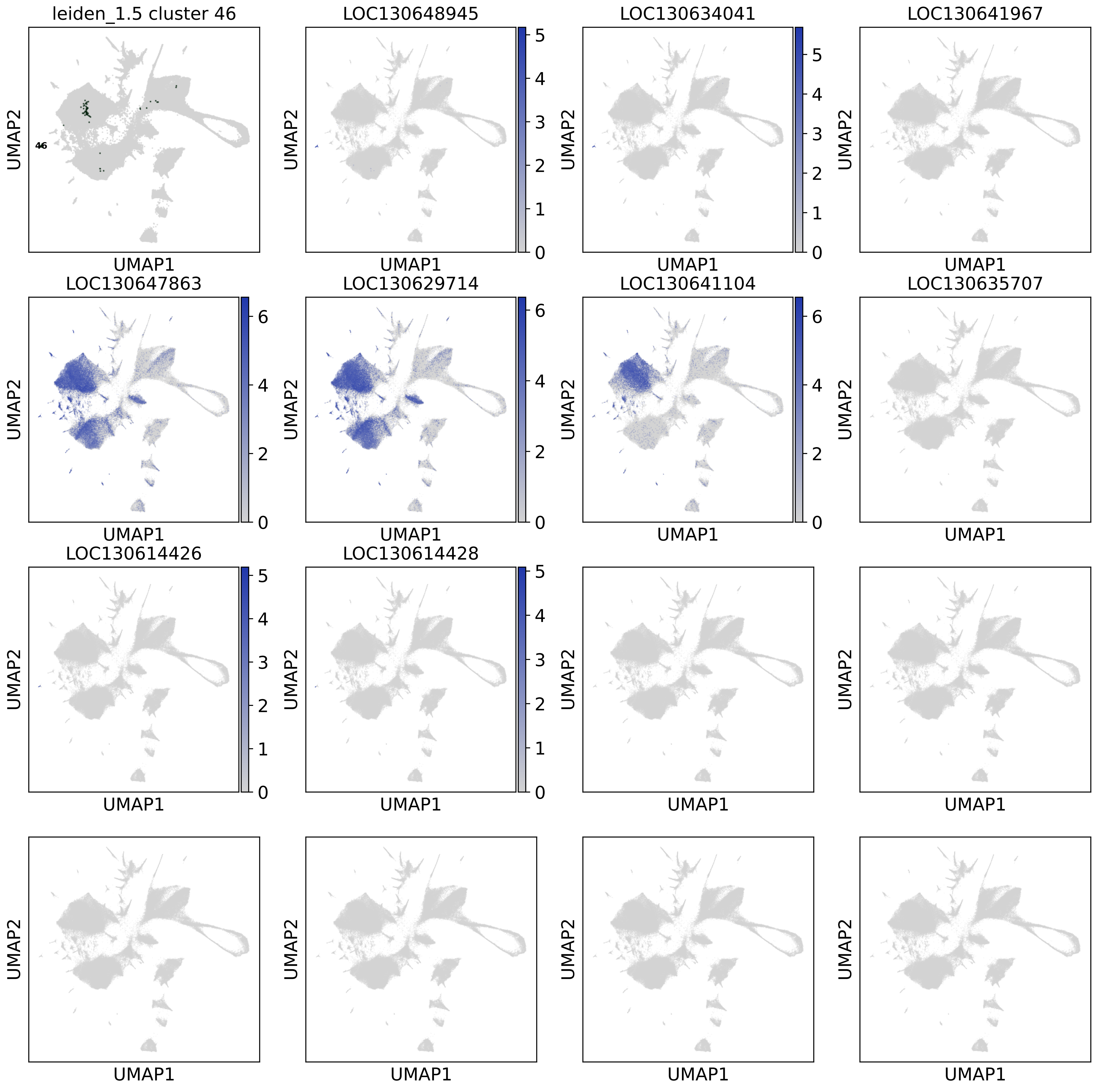

leiden_1.5 cluster 46
46
LOC130648945
LOC130634041
LOC130641967
LOC130647863
LOC130629714
LOC130641104
LOC130635707
LOC130614426
LOC130614428
UMAP1
UMAP2

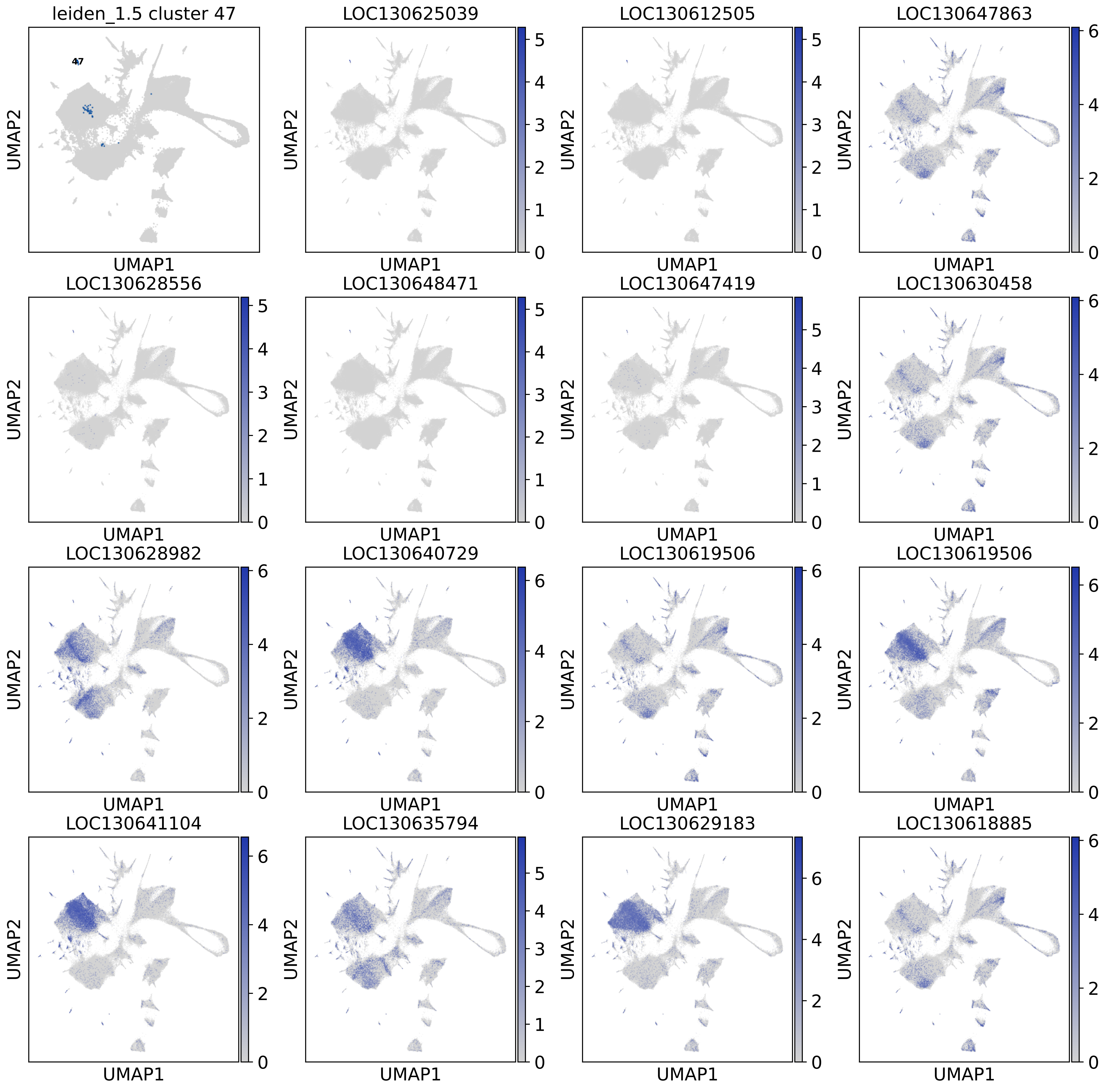

leiden_1.5 cluster 47
47
LOC130625039
LOC130612505
LOC130647863
LOC130628556
LOC130648471
LOC130647419
LOC130630458
LOC130628982
LOC130640729
LOC130619506
LOC130619506
LOC130641104
LOC130635794
LOC130629183
LOC130618885
UMAP1
UMAP2

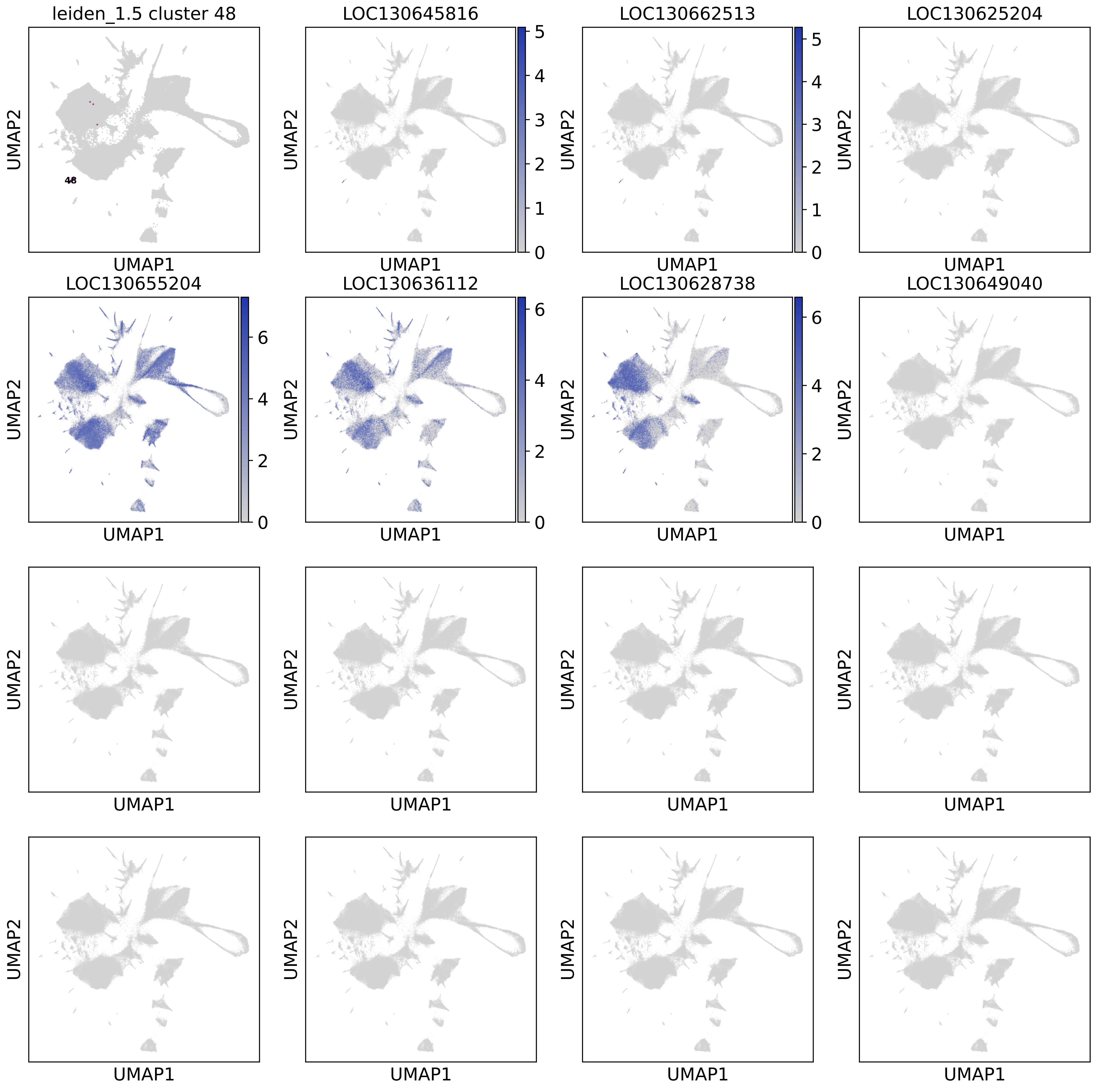

leiden_1.5 cluster 48
48
LOC130645816
LOC130662513
LOC130625204
LOC130655204
LOC130636112
LOC130628738
LOC130649040
UMAP1
UMAP2

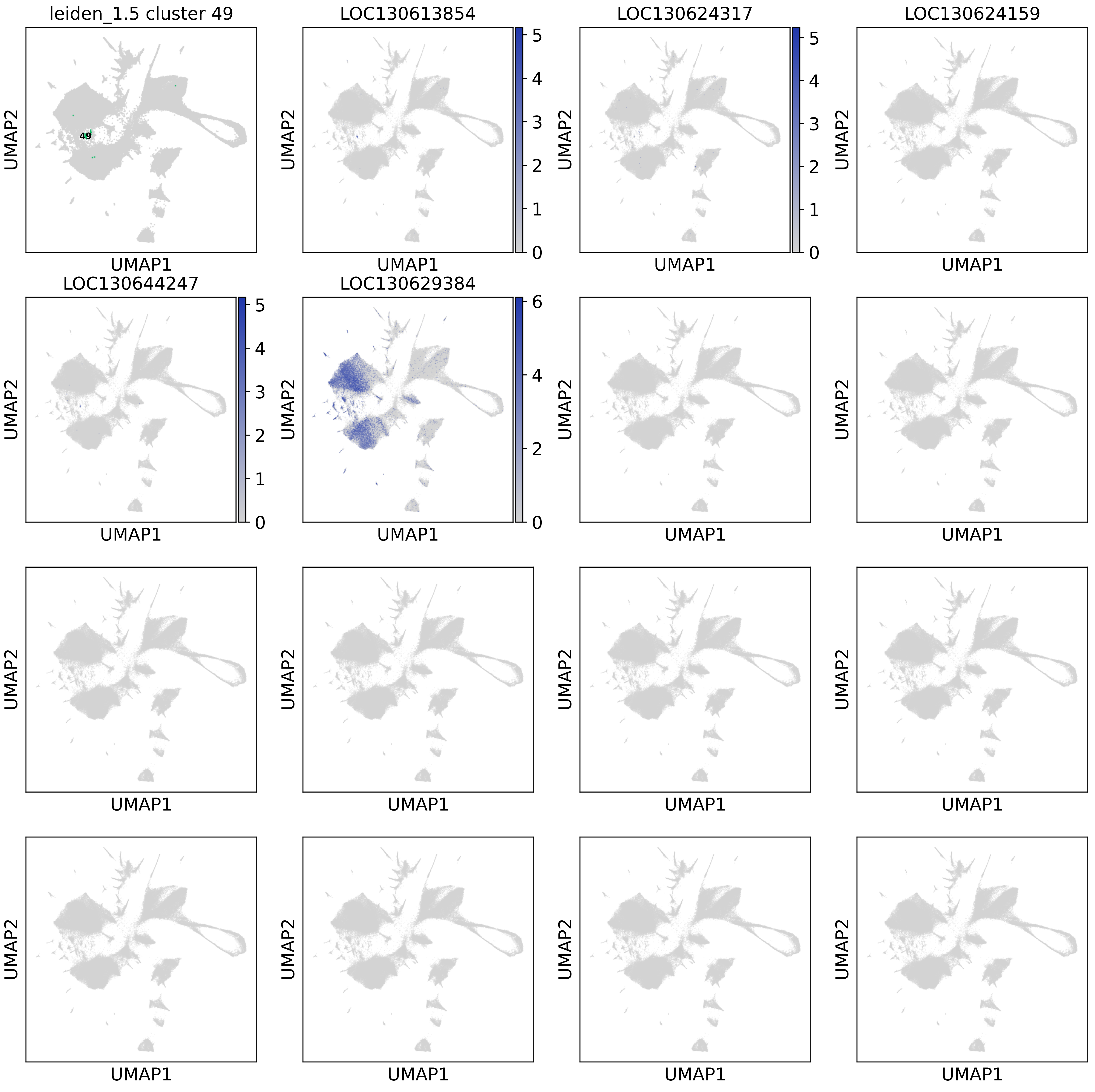

leiden_1.5 cluster 49
49
LOC130613854
LOC130624317
LOC130624159
LOC130644247
LOC130629384
UMAP1
UMAP2
0
1
2
3
4
5
6

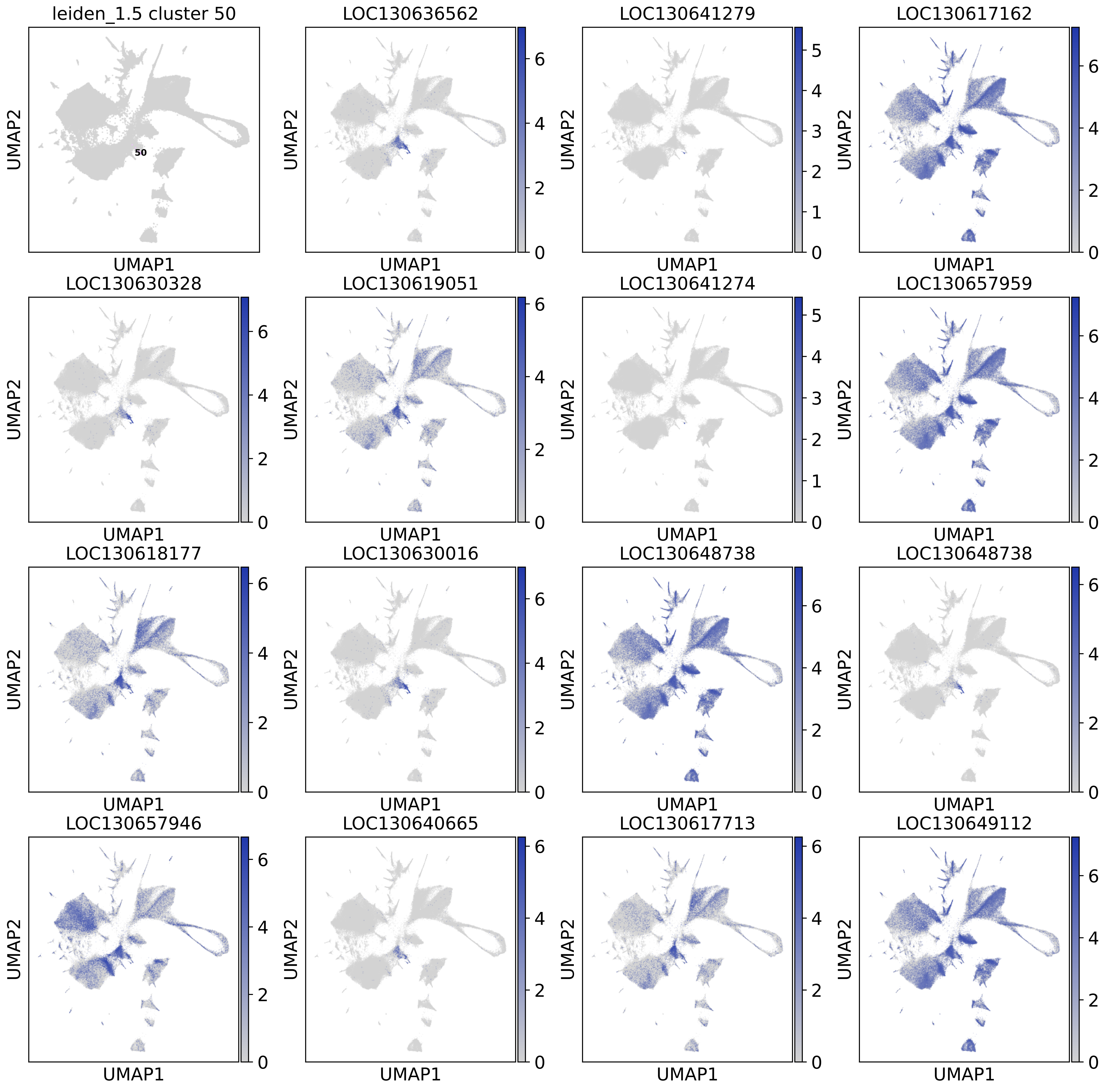
leiden_1.5 cluster 50
50
LOC130636562
LOC130641279
LOC130617162
LOC130630328
LOC130619051
LOC130641274
LOC130657959
LOC130618177
LOC130630016
LOC130648738
LOC130648738
LOC130657946
LOC130640665
LOC130617713
LOC130649112
UMAP1
UMAP2

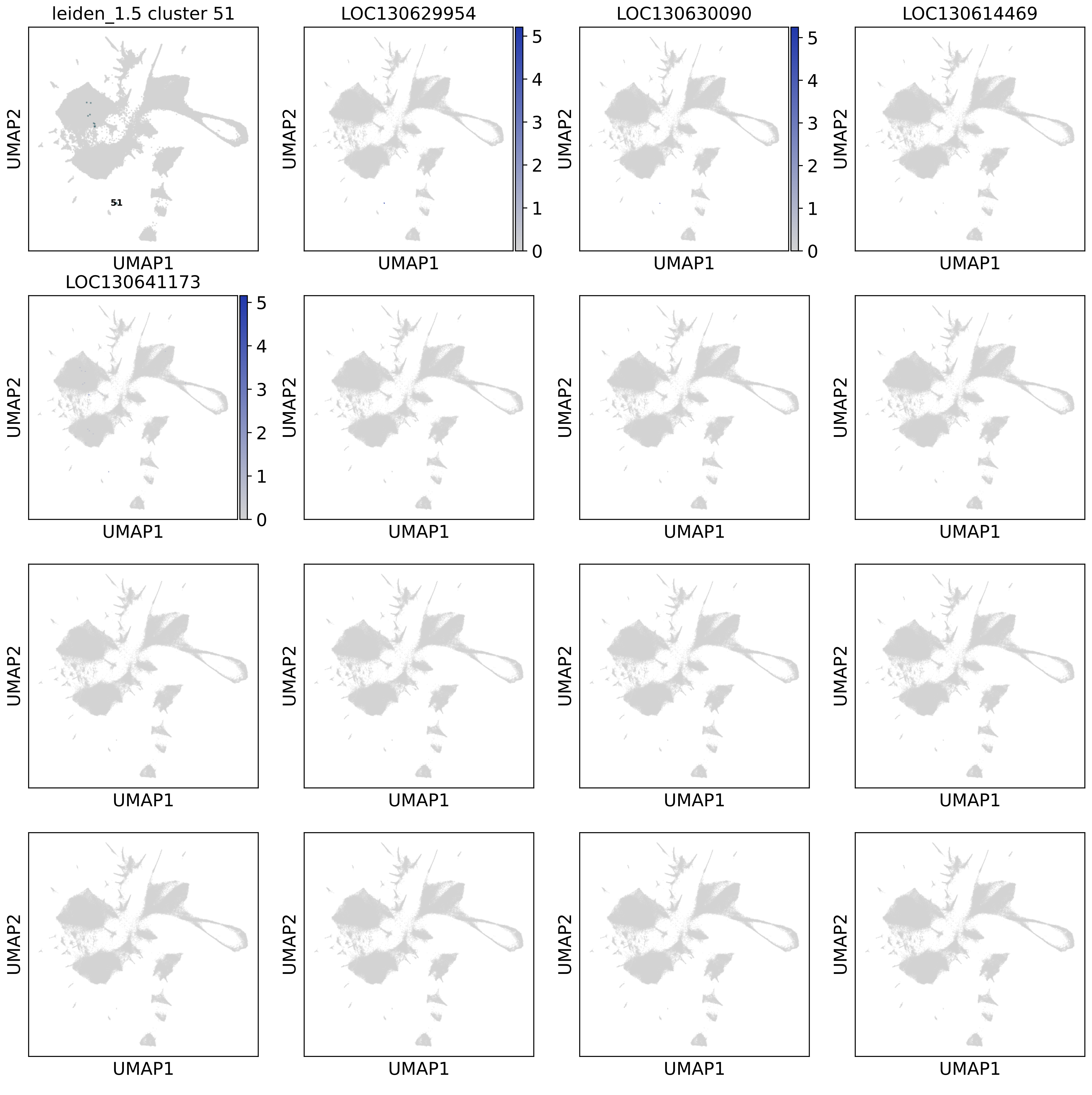

leiden_1.5 cluster 51
LOC130629954
LOC130630090
LOC130614469
LOC130641173
51
UMAP1
UMAP2
0
1
2
3
4
5

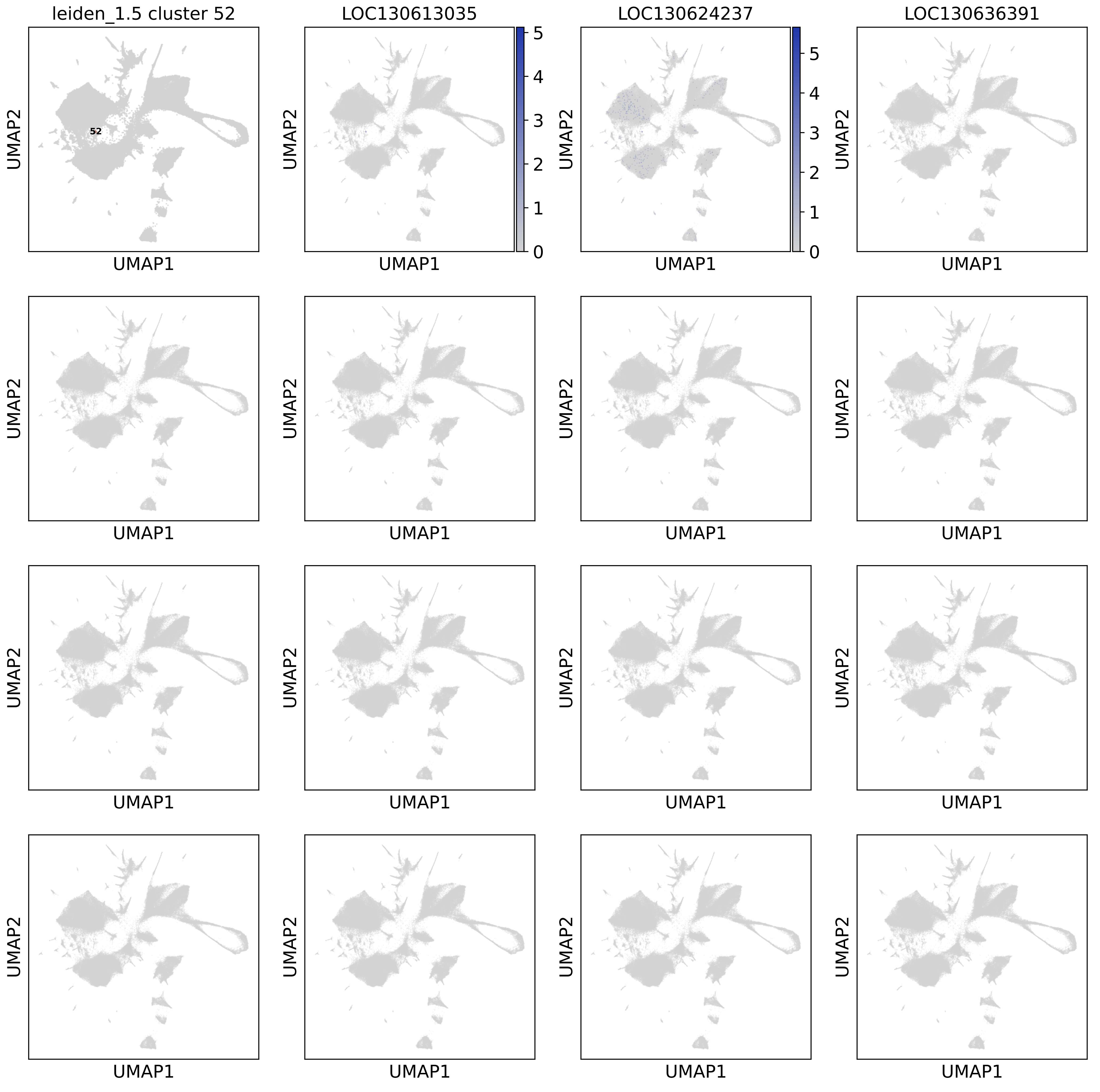
leiden_1.5 cluster 52
LOC130613035
LOC130624237
LOC130636391
52
UMAP1
UMAP2
0
1
2
3
4
5
