## Supplementary material for "The *Hydractinia* cell atlas reveals cellular and molecular principles of cnidarian coloniality": Supplementary File 6.pdf

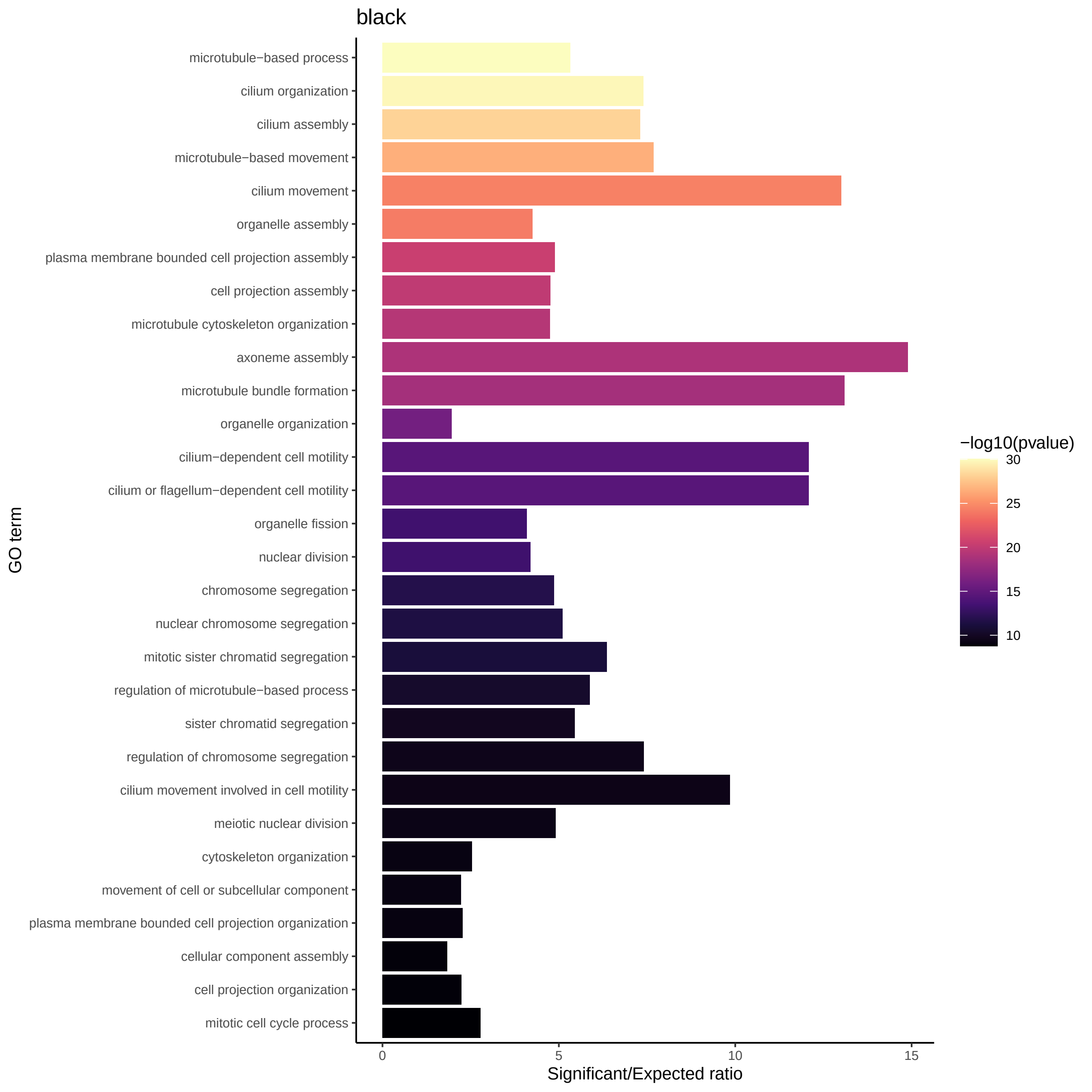

black
microtubule−based process
cilium organization
cilium assembly
microtubule−based movement
cilium movement
organelle assembly
plasma membrane bounded cell projection assembly
cell projection assembly
microtubule cytoskeleton organization
axoneme assembly
microtubule bundle formation
organelle organization
cilium−dependent cell motility
cilium or flagellum−dependent cell motility
organelle fission
nuclear division
chromosome segregation
nuclear chromosome segregation
mitotic sister chromatid segregation
regulation of microtubule−based process
sister chromatid segregation
regulation of chromosome segregation
cilium movement involved in cell motility
meiotic nuclear division
cytoskeleton organization
movement of cell or subcellular component
plasma membrane bounded cell projection organization
cellular component assembly
cell projection organization
mitotic cell cycle process
GO term
0
5
10
15
Significant/Expected ratio
−log10(pvalue)
30
25
20
15
10

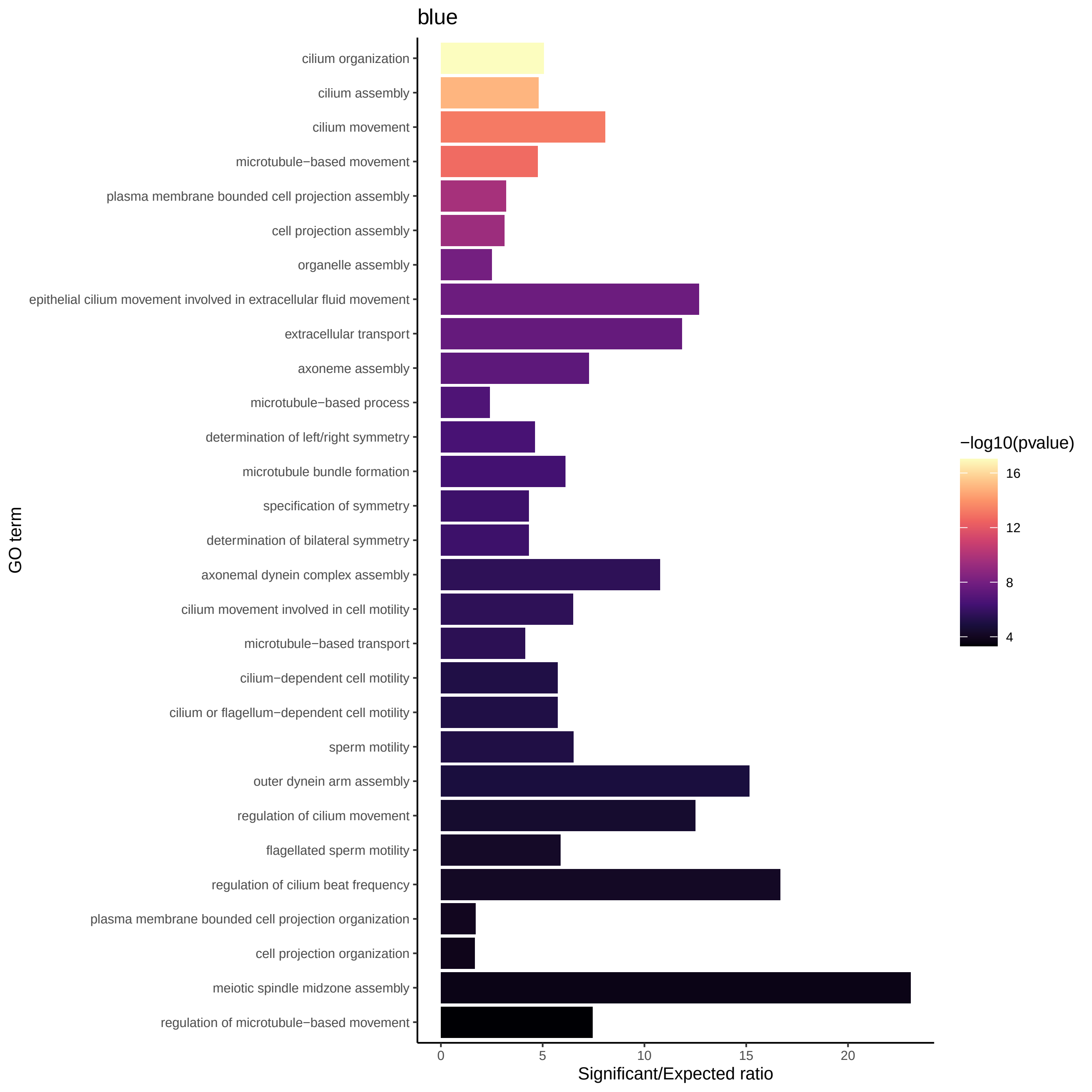
blue
cilium organization
cilium assembly
cilium movement
microtubule−based movement
plasma membrane bounded cell projection assembly
cell projection assembly
organelle assembly
epithelial cilium movement involved in extracellular fluid movement
extracellular transport
axoneme assembly
microtubule−based process
determination of left/right symmetry
microtubule bundle formation
specification of symmetry
determination of bilateral symmetry
axonemal dynein complex assembly
cilium movement involved in cell motility
microtubule−based transport
cilium−dependent cell motility
cilium or flagellum−dependent cell motility
sperm motility
outer dynein arm assembly
regulation of cilium movement
flagellated sperm motility
regulation of cilium beat frequency
plasma membrane bounded cell projection organization
cell projection organization
meiotic spindle midzone assembly
regulation of microtubule−based movement
GO term
0
5
10
15
20
Significant/Expected ratio
−log10(pvalue)
16
12
8
4

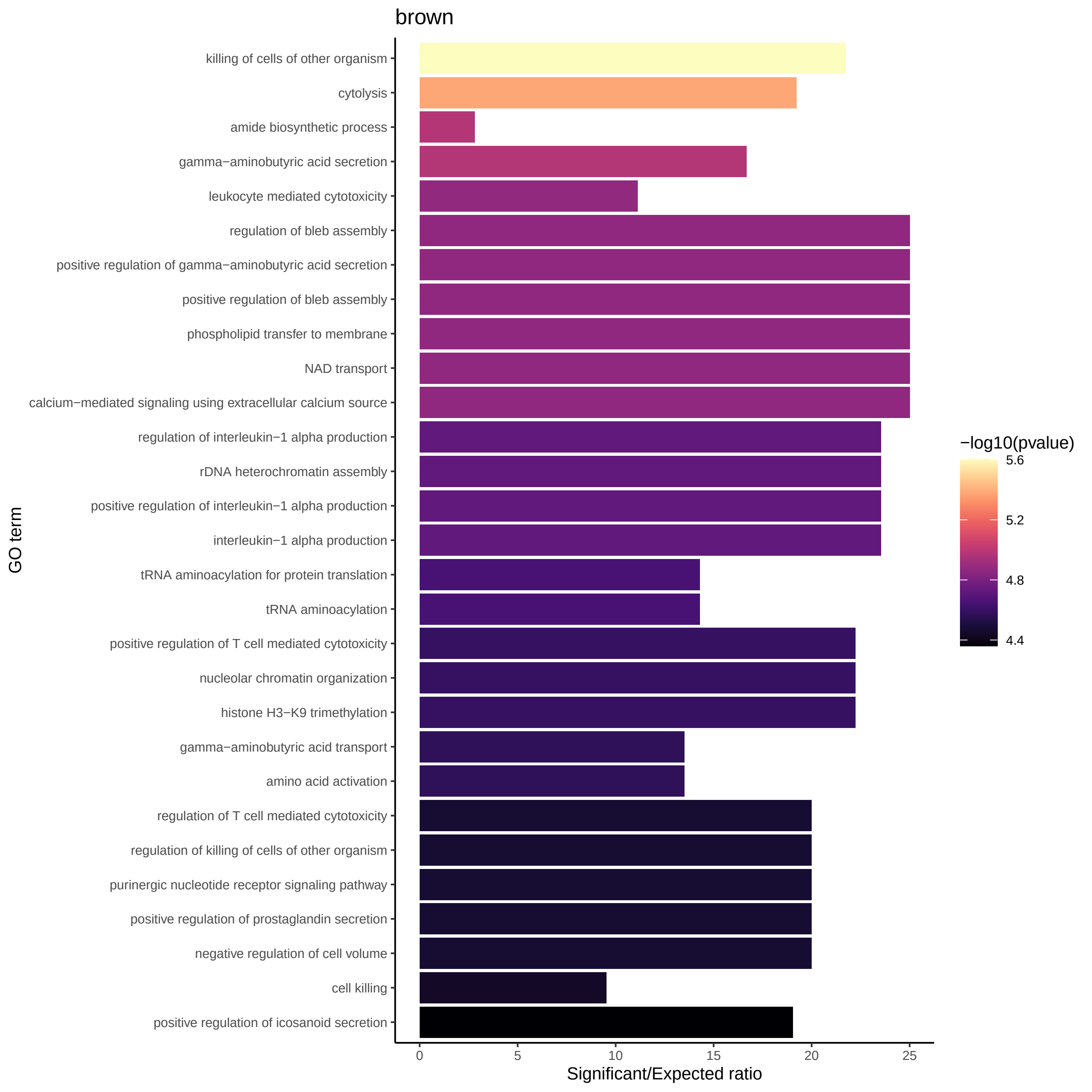

brown
killing of cells of other organism
cytolysis
amide biosynthetic process
gamma−aminobutyric acid secretion
leukocyte mediated cytotoxicity
regulation of bleb assembly
positive regulation of gamma−aminobutyric acid secretion
positive regulation of bleb assembly
phospholipid transfer to membrane
NAD transport
calcium−mediated signaling using extracellular calcium source
regulation of interleukin−1 alpha production
rDNA heterochromatin assembly
positive regulation of interleukin−1 alpha production
interleukin−1 alpha production
tRNA aminoacylation for protein translation
tRNA aminoacylation
positive regulation of T cell mediated cytotoxicity
nucleolar chromatin organization
histone H3−K9 trimethylation
gamma−aminobutyric acid transport
amino acid activation
regulation of T cell mediated cytotoxicity
regulation of killing of cells of other organism
purinergic nucleotide receptor signaling pathway
positive regulation of prostaglandin secretion
negative regulation of cell volume
cell killing
positive regulation of icosanoid secretion
GO term
0
5
10
15
20
25
Significant/Expected ratio
−log10(pvalue)
5.6
5.2
4.8
4.4

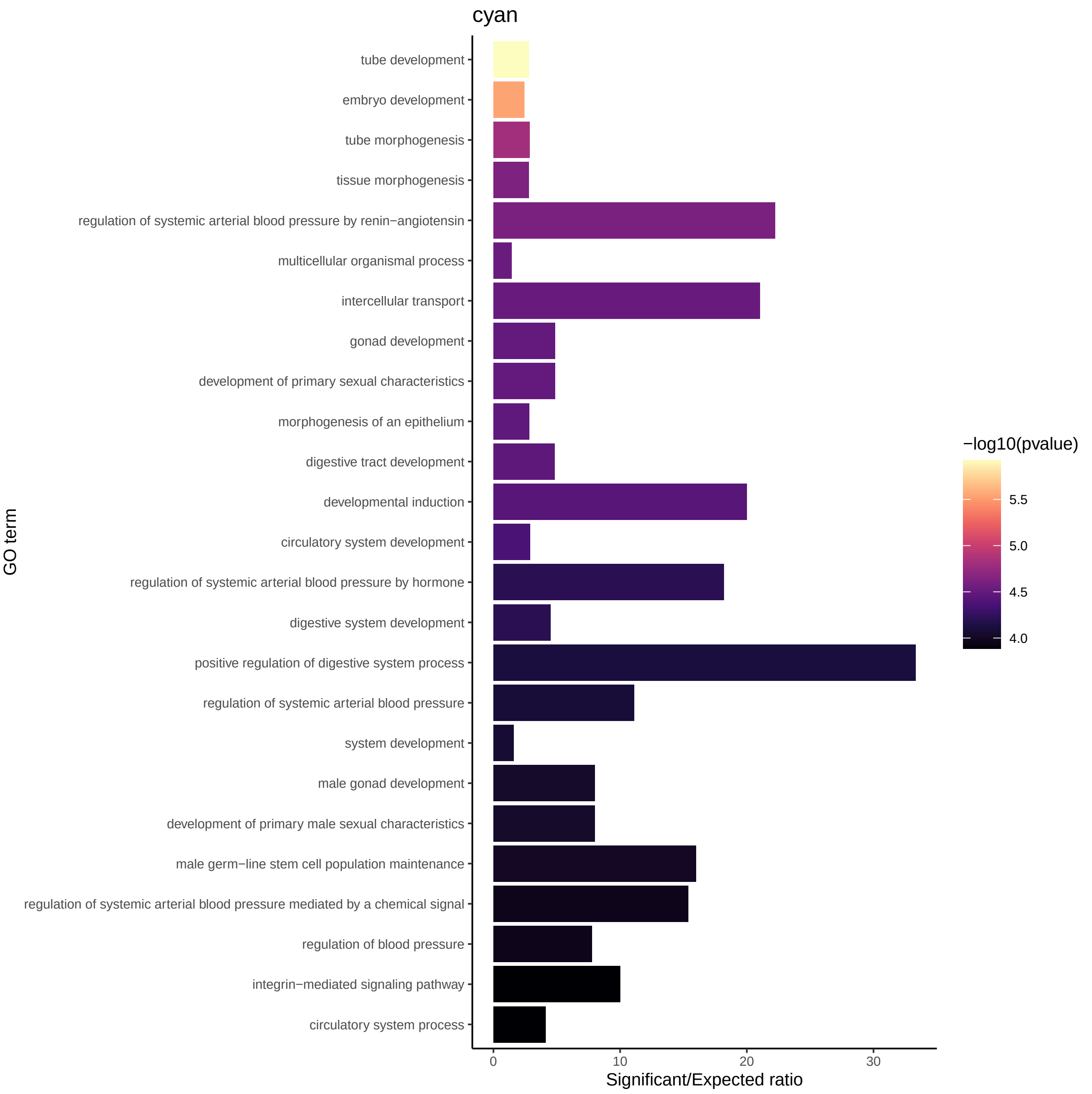
cyan
tube development
embryo development
tube morphogenesis
tissue morphogenesis
regulation of systemic arterial blood pressure by renin−angiotensin
multicellular organismal process
intercellular transport
gonad development
development of primary sexual characteristics
morphogenesis of an epithelium
digestive tract development
developmental induction
circulatory system development
regulation of systemic arterial blood pressure by hormone
digestive system development
positive regulation of digestive system process
regulation of systemic arterial blood pressure
system development
male gonad development
development of primary male sexual characteristics
male germ−line stem cell population maintenance
regulation of systemic arterial blood pressure mediated by a chemical signal
regulation of blood pressure
integrin−mediated signaling pathway
circulatory system process
GO term
0
10
20
30
Significant/Expected ratio
−log10(pvalue)
5.5
5.0
4.5
4.0

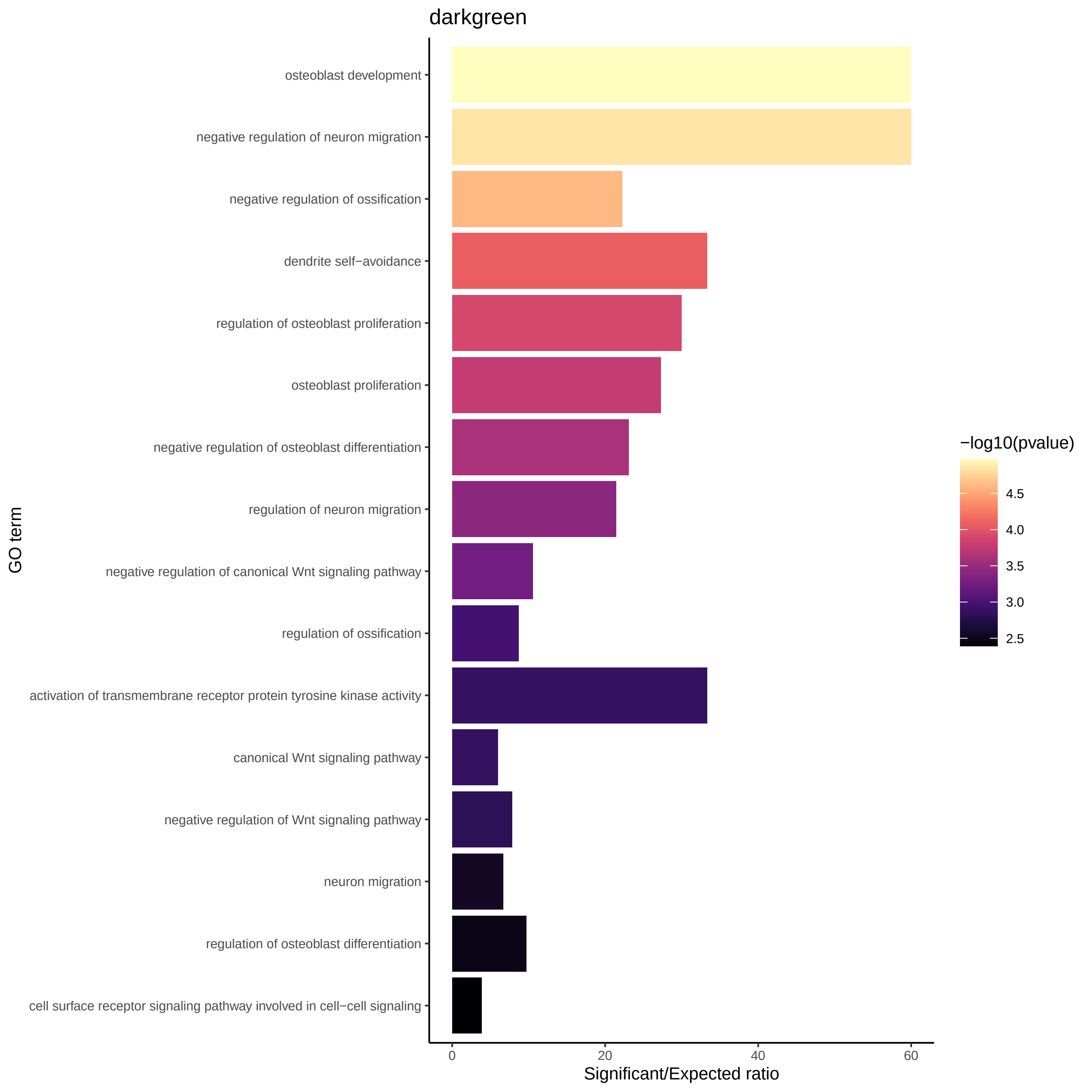

darkgreen
osteoblast development
negative regulation of neuron migration
negative regulation of ossification
dendrite self−avoidance
regulation of osteoblast proliferation
osteoblast proliferation
negative regulation of osteoblast differentiation
regulation of neuron migration
negative regulation of canonical Wnt signaling pathway
regulation of ossification
activation of transmembrane receptor protein tyrosine kinase activity
canonical Wnt signaling pathway
negative regulation of Wnt signaling pathway
neuron migration
regulation of osteoblast differentiation
cell surface receptor signaling pathway involved in cell−cell signaling
GO term
0
20
40
60
Significant/Expected ratio
−log10(pvalue)
4.5
4.0
3.5
3.0
2.5

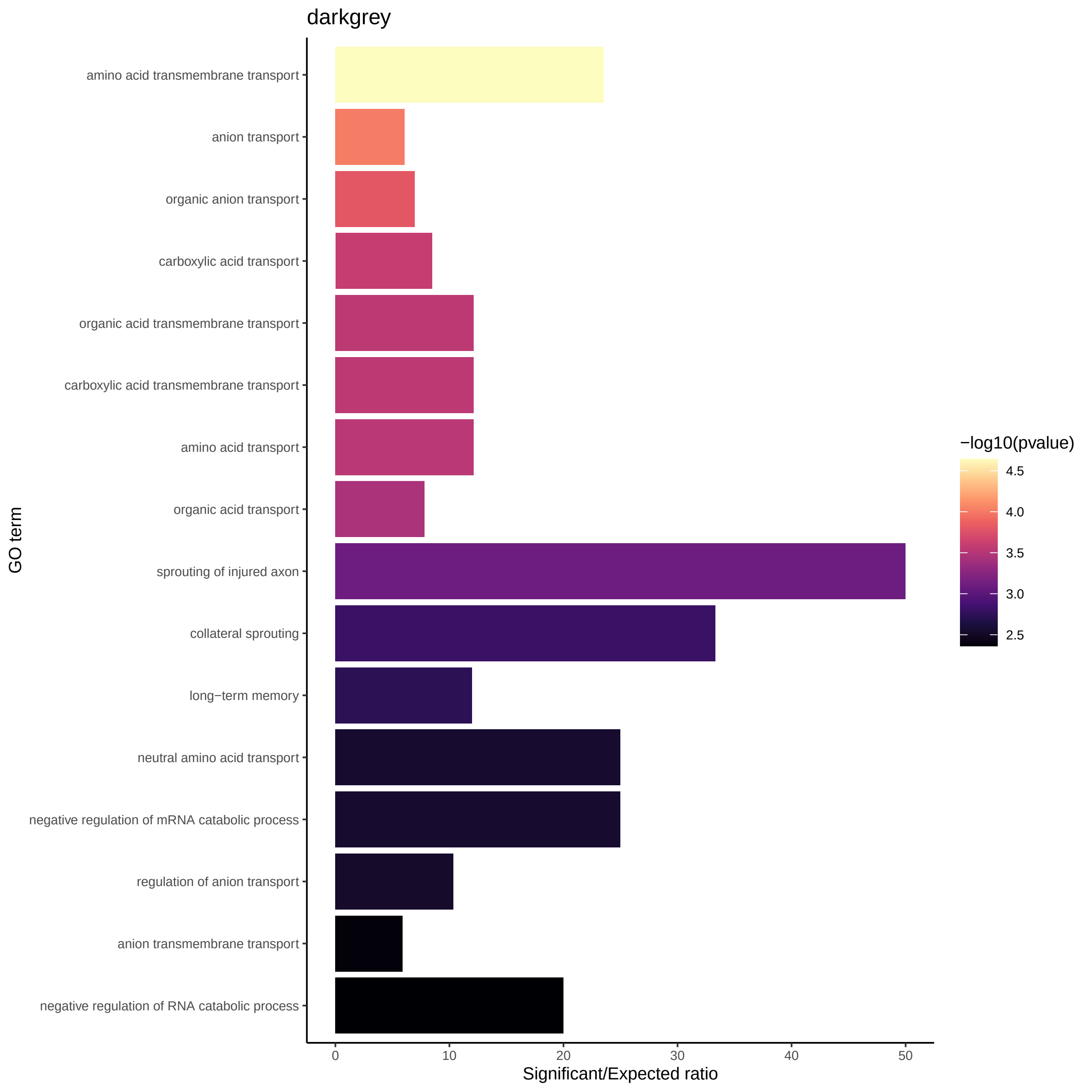
darkgrey
amino acid transmembrane transport
anion transport
organic anion transport
carboxylic acid transport
organic acid transmembrane transport
carboxylic acid transmembrane transport
amino acid transport
organic acid transport
sprouting of injured axon
collateral sprouting
long−term memory
neutral amino acid transport
negative regulation of mRNA catabolic process
regulation of anion transport
anion transmembrane transport
negative regulation of RNA catabolic process
GO term
0
10
20
30
40
50
Significant/Expected ratio
−log10(pvalue)
4.5
4.0
3.5
3.0
2.5

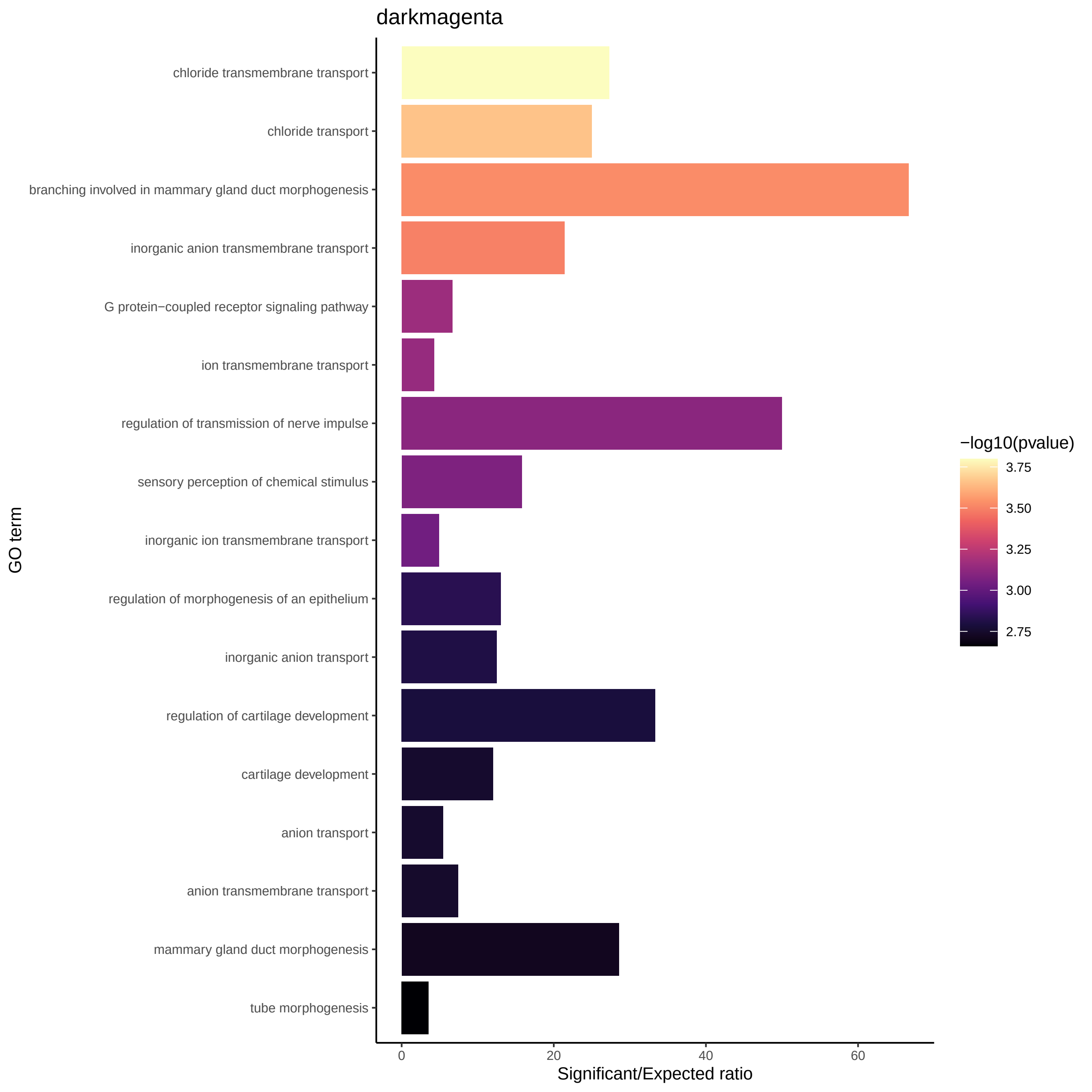

darkmagenta
chloride transmembrane transport
chloride transport
branching involved in mammary gland duct morphogenesis
inorganic anion transmembrane transport
G protein−coupled receptor signaling pathway
ion transmembrane transport
regulation of transmission of nerve impulse
sensory perception of chemical stimulus
inorganic ion transmembrane transport
regulation of morphogenesis of an epithelium
inorganic anion transport
regulation of cartilage development
cartilage development
anion transport
anion transmembrane transport
mammary gland duct morphogenesis
tube morphogenesis
GO term
0
20
40
60
Significant/Expected ratio
−log10(pvalue)
3.75
3.50
3.25
3.00
2.75

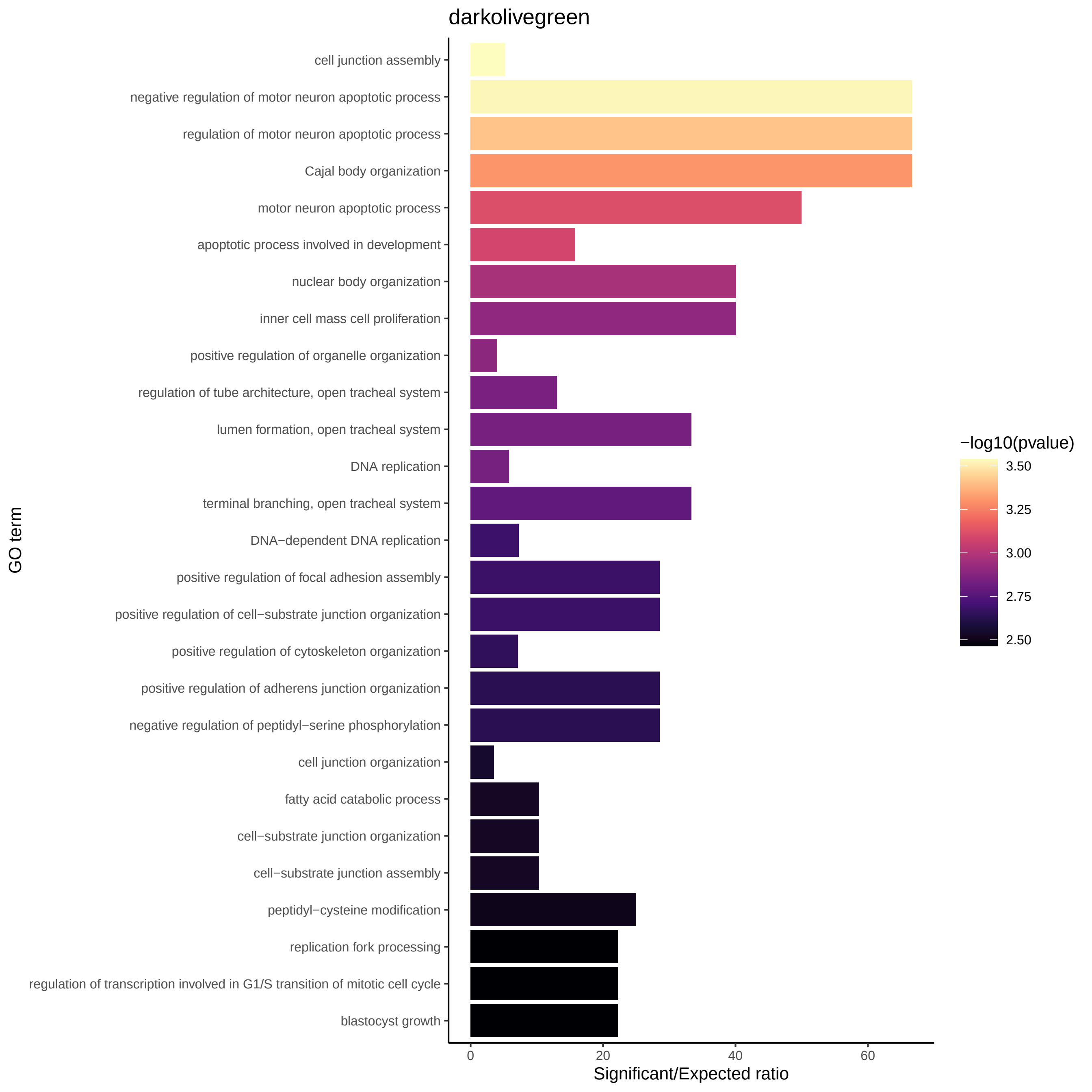
darkolivegreen
cell junction assembly
negative regulation of motor neuron apoptotic process
regulation of motor neuron apoptotic process
Cajal body organization
motor neuron apoptotic process
apoptotic process involved in development
nuclear body organization
inner cell mass cell proliferation
positive regulation of organelle organization
regulation of tube architecture, open tracheal system
lumen formation, open tracheal system
DNA replication
terminal branching, open tracheal system
DNA−dependent DNA replication
positive regulation of focal adhesion assembly
positive regulation of cell−substrate junction organization
positive regulation of cytoskeleton organization
positive regulation of adherens junction organization
negative regulation of peptidyl−serine phosphorylation
cell junction organization
fatty acid catabolic process
cell−substrate junction organization
cell−substrate junction assembly
peptidyl−cysteine modification
replication fork processing
regulation of transcription involved in G1/S transition of mitotic cell cycle
blastocyst growth
GO term
0
20
40
60
Significant/Expected ratio
−log10(pvalue)
3.50
3.25
3.00
2.75
2.50

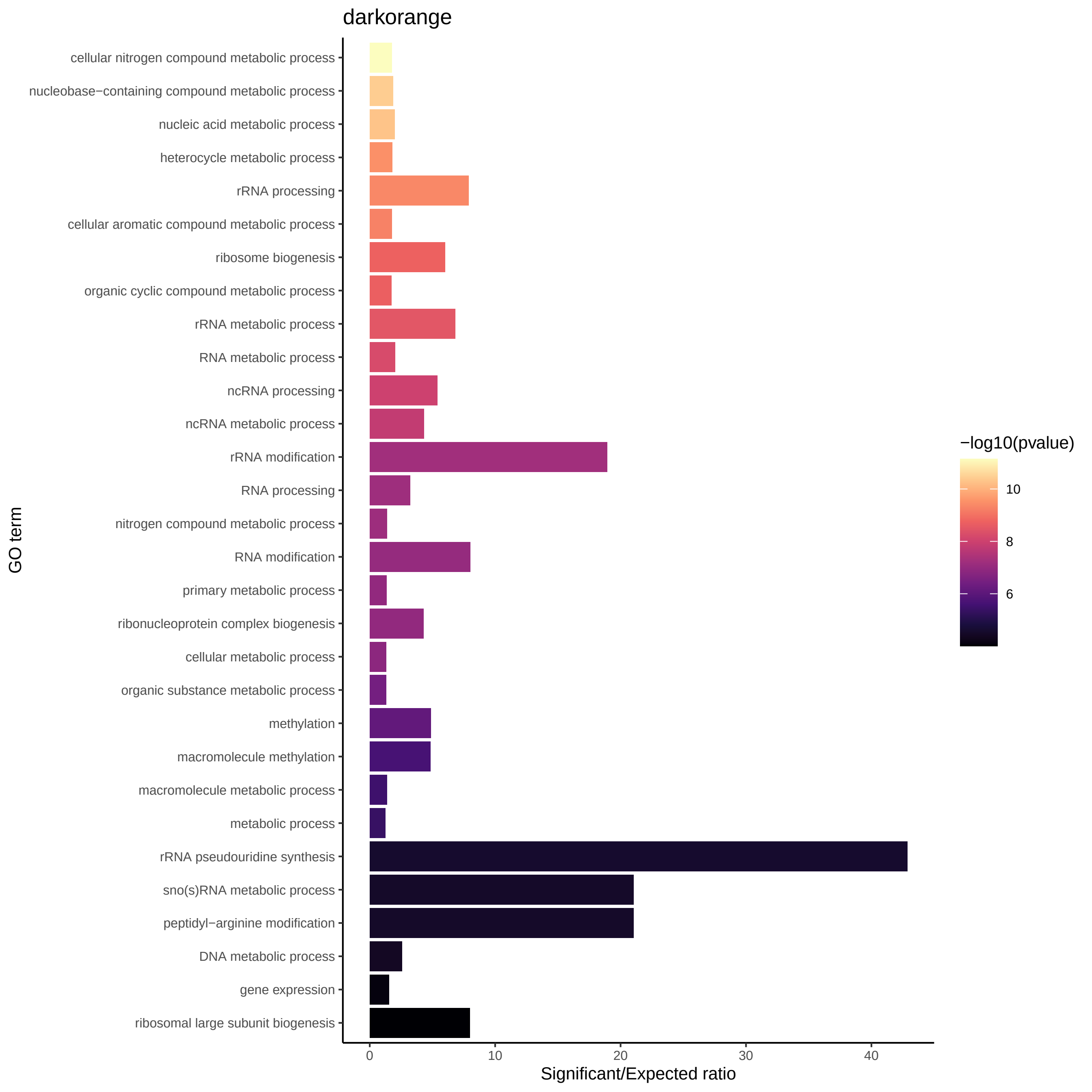
darkorange
cellular nitrogen compound metabolic process
nucleobase−containing compound metabolic process
nucleic acid metabolic process
heterocycle metabolic process
rRNA processing
cellular aromatic compound metabolic process
ribosome biogenesis
organic cyclic compound metabolic process
rRNA metabolic process
RNA metabolic process
ncRNA processing
ncRNA metabolic process
rRNA modification
RNA processing
nitrogen compound metabolic process
RNA modification
primary metabolic process
ribonucleoprotein complex biogenesis
cellular metabolic process
organic substance metabolic process
methylation
macromolecule methylation
macromolecule metabolic process
metabolic process
rRNA pseudouridine synthesis
sno(s)RNA metabolic process
peptidyl−arginine modification
DNA metabolic process
gene expression
ribosomal large subunit biogenesis
GO term
0
10
20
30
40
Significant/Expected ratio
−log10(pvalue)
10
8
6

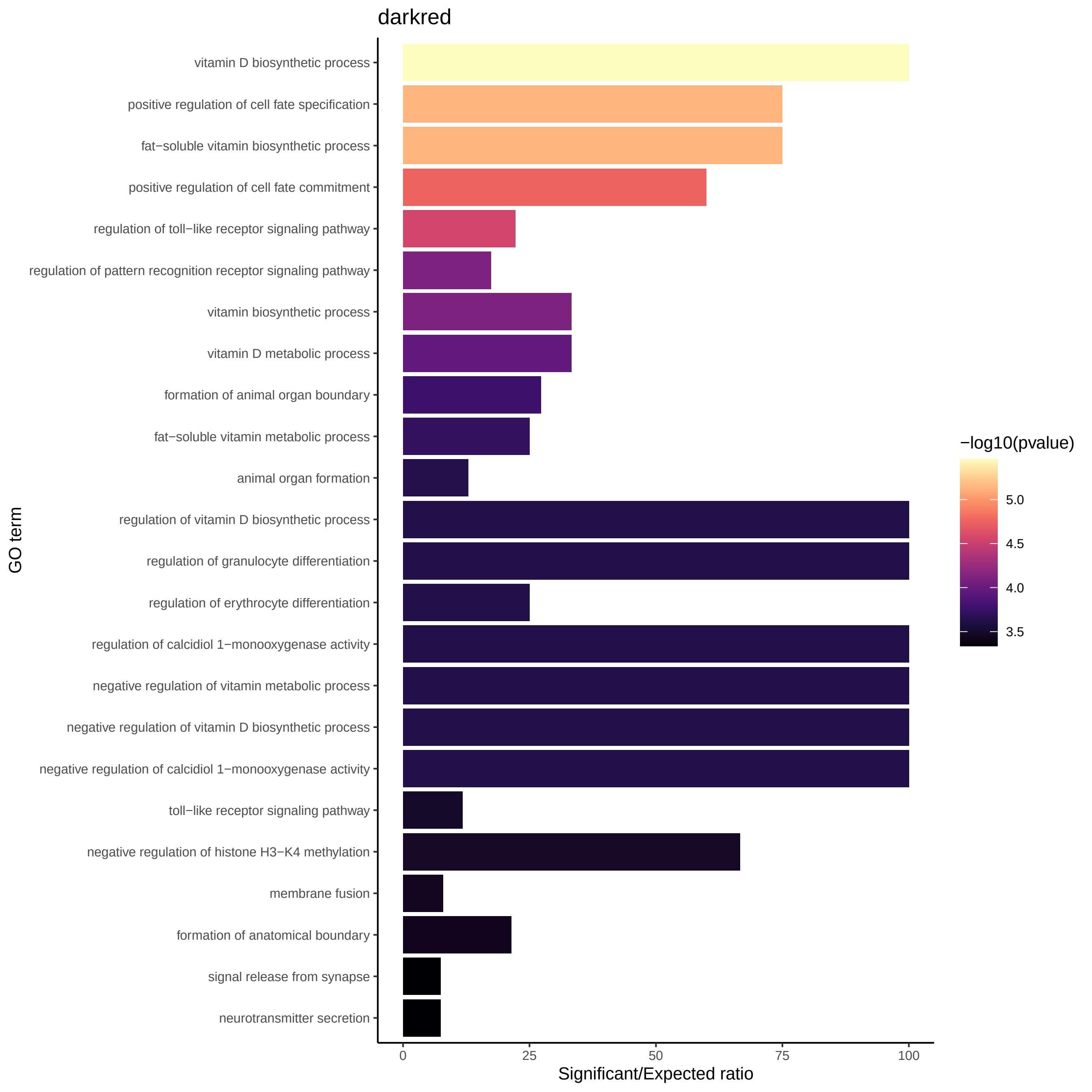
darkred
vitamin D biosynthetic process
positive regulation of cell fate specification
fat−soluble vitamin biosynthetic process
positive regulation of cell fate commitment
regulation of toll−like receptor signaling pathway
regulation of pattern recognition receptor signaling pathway
vitamin biosynthetic process
vitamin D metabolic process
formation of animal organ boundary
fat−soluble vitamin metabolic process
animal organ formation
regulation of vitamin D biosynthetic process
regulation of granulocyte differentiation
regulation of erythrocyte differentiation
regulation of calcidiol 1−monooxygenase activity
negative regulation of vitamin metabolic process
negative regulation of vitamin D biosynthetic process
negative regulation of calcidiol 1−monooxygenase activity
toll−like receptor signaling pathway
negative regulation of histone H3−K4 methylation
membrane fusion
formation of anatomical boundary
signal release from synapse
neurotransmitter secretion
GO term
0
25
50
75
100
Significant/Expected ratio
−log10(pvalue)
5.0
4.5
4.0
3.5

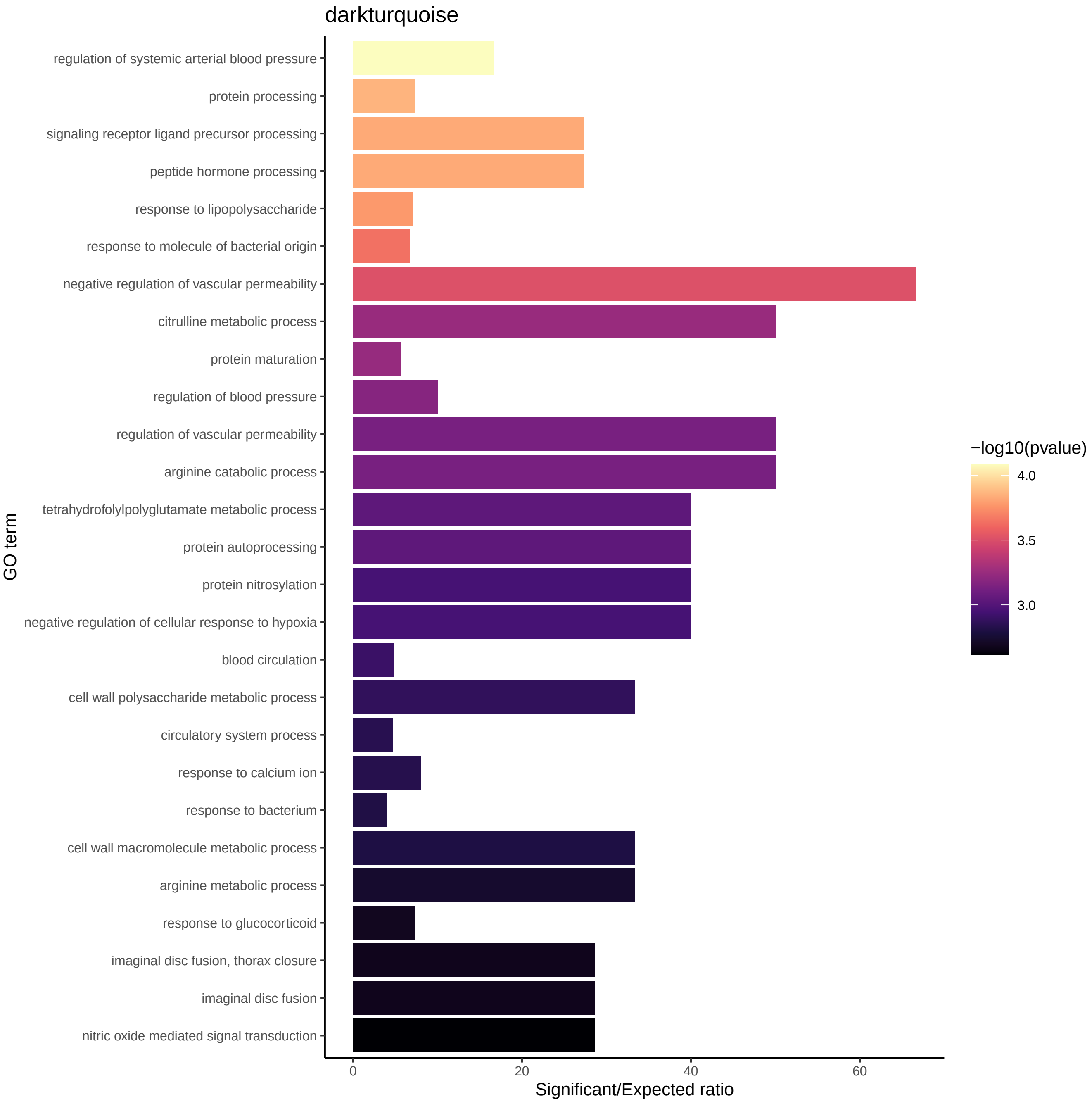
darkturquoise
regulation of systemic arterial blood pressure
protein processing
signaling receptor ligand precursor processing
peptide hormone processing
response to lipopolysaccharide
response to molecule of bacterial origin
negative regulation of vascular permeability
citrulline metabolic process
protein maturation
regulation of blood pressure
regulation of vascular permeability
arginine catabolic process
tetrahydrofolylpolyglutamate metabolic process
protein autoprocessing
protein nitrosylation
negative regulation of cellular response to hypoxia
blood circulation
cell wall polysaccharide metabolic process
circulatory system process
response to calcium ion
response to bacterium
cell wall macromolecule metabolic process
arginine metabolic process
response to glucocorticoid
imaginal disc fusion, thorax closure
imaginal disc fusion
nitric oxide mediated signal transduction
GO term
0
20
40
60
Significant/Expected ratio
−log10(pvalue)
4.0
3.5
3.0

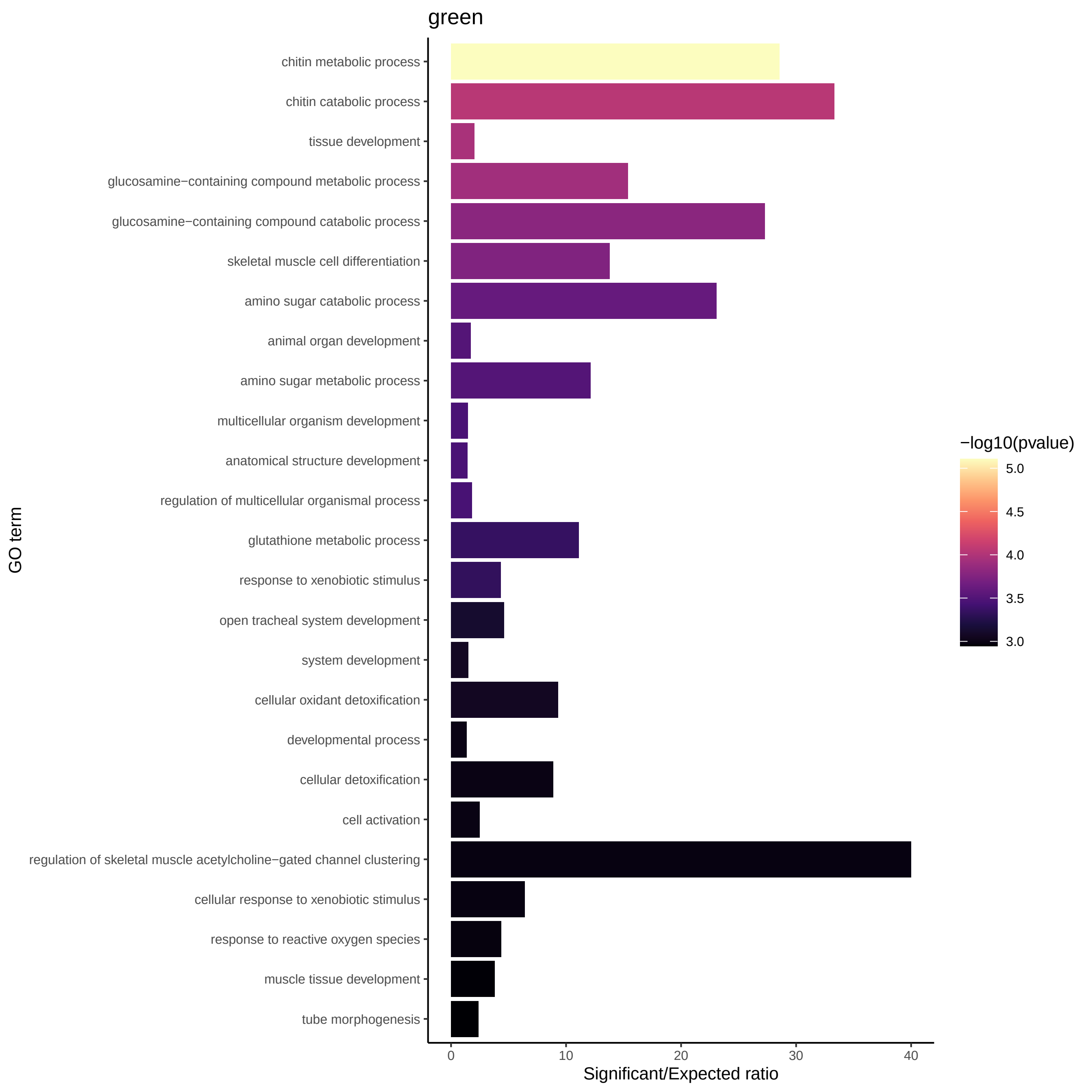
green
chitin metabolic process
chitin catabolic process
tissue development
glucosamine−containing compound metabolic process
glucosamine−containing compound catabolic process
skeletal muscle cell differentiation
amino sugar catabolic process
animal organ development
amino sugar metabolic process
multicellular organism development
anatomical structure development
regulation of multicellular organismal process
glutathione metabolic process
response to xenobiotic stimulus
open tracheal system development
system development
cellular oxidant detoxification
developmental process
cellular detoxification
cell activation
regulation of skeletal muscle acetylcholine−gated channel clustering
cellular response to xenobiotic stimulus
response to reactive oxygen species
muscle tissue development
tube morphogenesis
GO term
0
10
20
30
40
Significant/Expected ratio
−log10(pvalue)
5.0
4.5
4.0
3.5
3.0

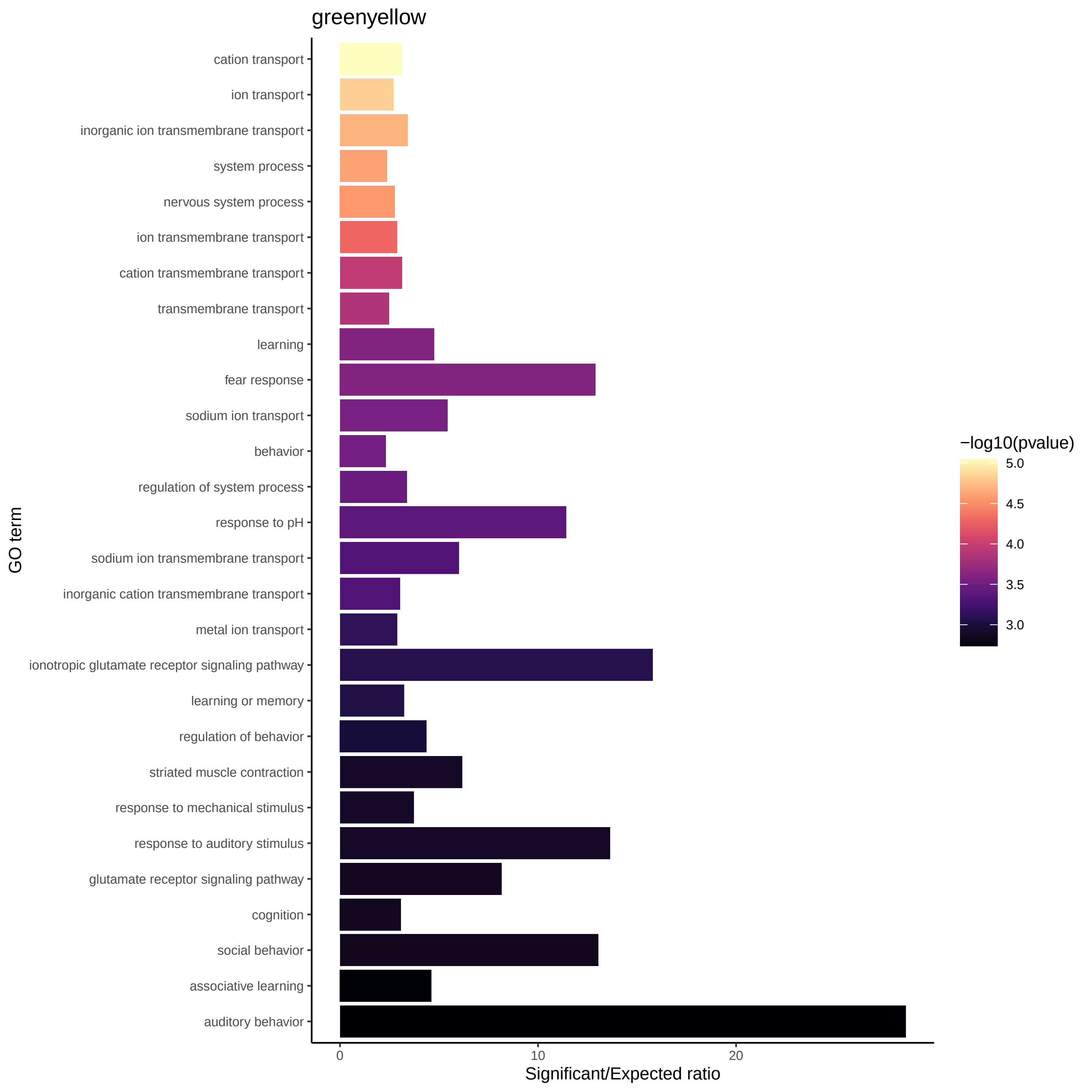
greenyellow
cation transport
ion transport
inorganic ion transmembrane transport
system process
nervous system process
ion transmembrane transport
cation transmembrane transport
transmembrane transport
learning
fear response
sodium ion transport
behavior
regulation of system process
response to pH
sodium ion transmembrane transport
inorganic cation transmembrane transport
metal ion transport
ionotropic glutamate receptor signaling pathway
learning or memory
regulation of behavior
striated muscle contraction
response to mechanical stimulus
response to auditory stimulus
glutamate receptor signaling pathway
cognition
social behavior
associative learning
auditory behavior
GO term
0
10
20
Significant/Expected ratio
−log10(pvalue)
5.0
4.5
4.0
3.5
3.0

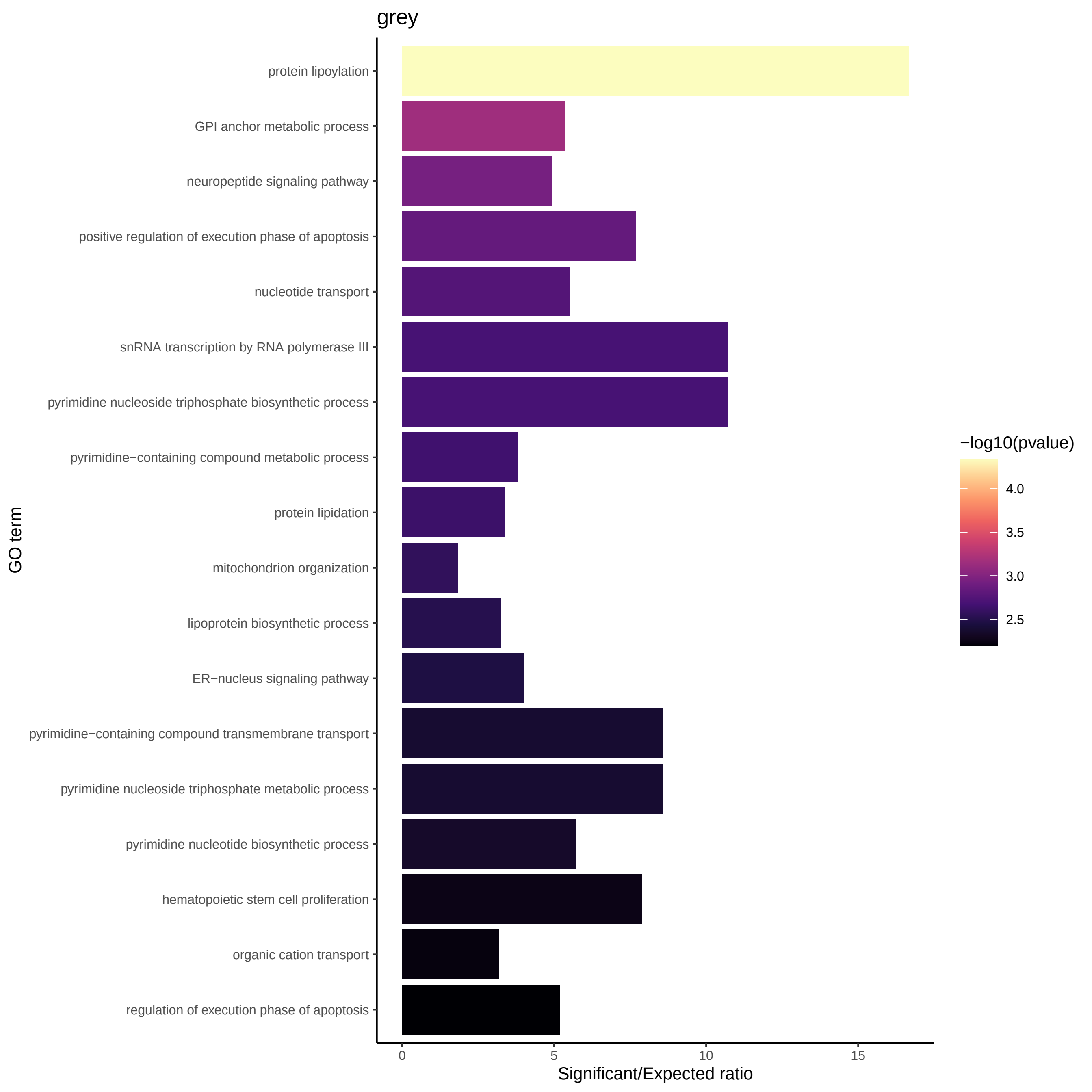
grey
protein lipoylation
GPI anchor metabolic process
neuropeptide signaling pathway
positive regulation of execution phase of apoptosis
nucleotide transport
snRNA transcription by RNA polymerase III
pyrimidine nucleoside triphosphate biosynthetic process
pyrimidine−containing compound metabolic process
protein lipidation
mitochondrion organization
lipoprotein biosynthetic process
ER−nucleus signaling pathway
pyrimidine−containing compound transmembrane transport
pyrimidine nucleoside triphosphate metabolic process
pyrimidine nucleotide biosynthetic process
hematopoietic stem cell proliferation
organic cation transport
regulation of execution phase of apoptosis
GO term
0
5
10
15
Significant/Expected ratio
−log10(pvalue)
4.0
3.5
3.0
2.5

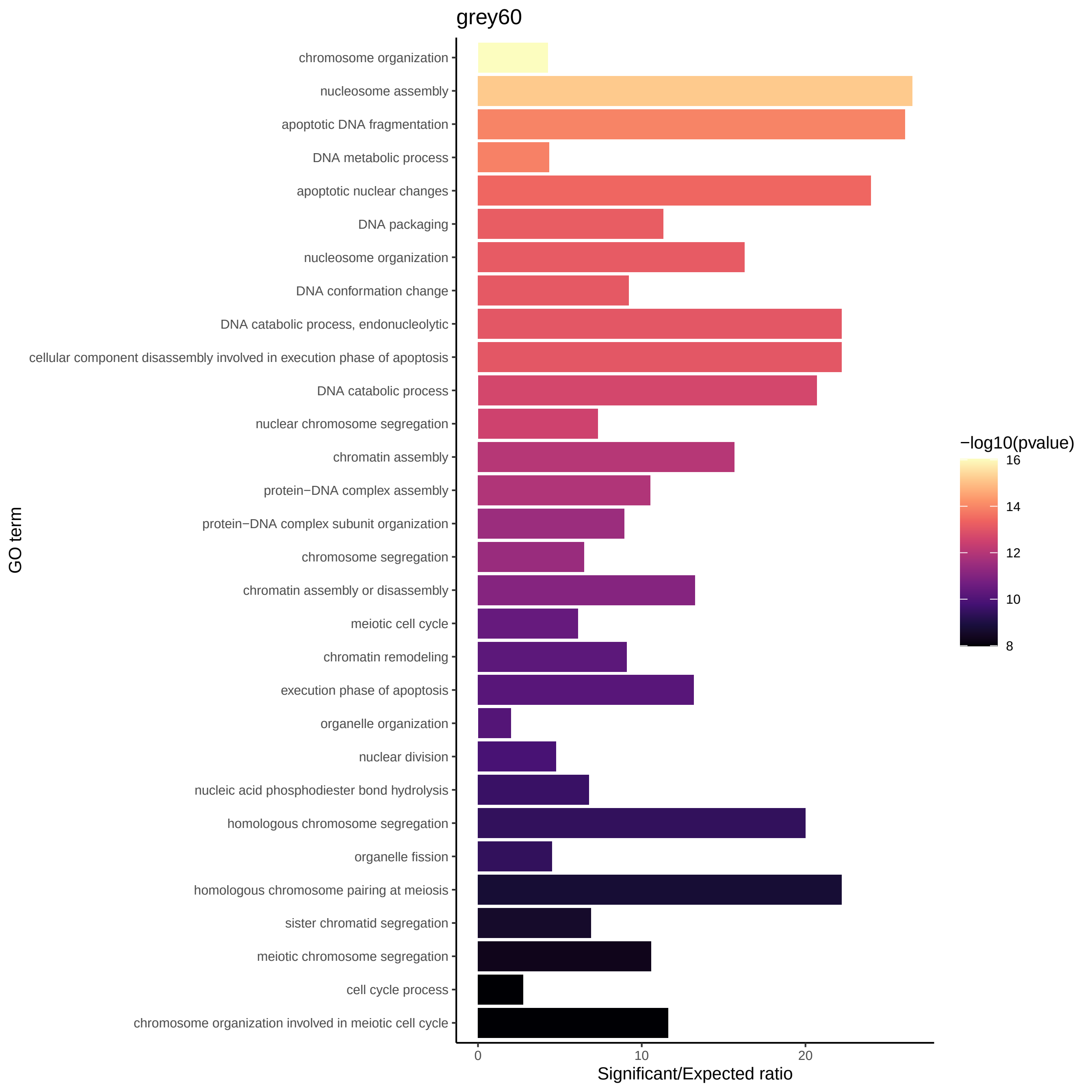

grey60
chromosome organization
nucleosome assembly
apoptotic DNA fragmentation
DNA metabolic process
apoptotic nuclear changes
DNA packaging
nucleosome organization
DNA conformation change
DNA catabolic process, endonucleolytic
cellular component disassembly involved in execution phase of apoptosis
DNA catabolic process
nuclear chromosome segregation
chromatin assembly
protein−DNA complex assembly
protein−DNA complex subunit organization
chromosome segregation
chromatin assembly or disassembly
meiotic cell cycle
chromatin remodeling
execution phase of apoptosis
organelle organization
nuclear division
nucleic acid phosphodiester bond hydrolysis
homologous chromosome segregation
organelle fission
homologous chromosome pairing at meiosis
sister chromatid segregation
meiotic chromosome segregation
cell cycle process
chromosome organization involved in meiotic cell cycle
GO term
0
10
20
Significant/Expected ratio
−log10(pvalue)
16
14
12
10
8

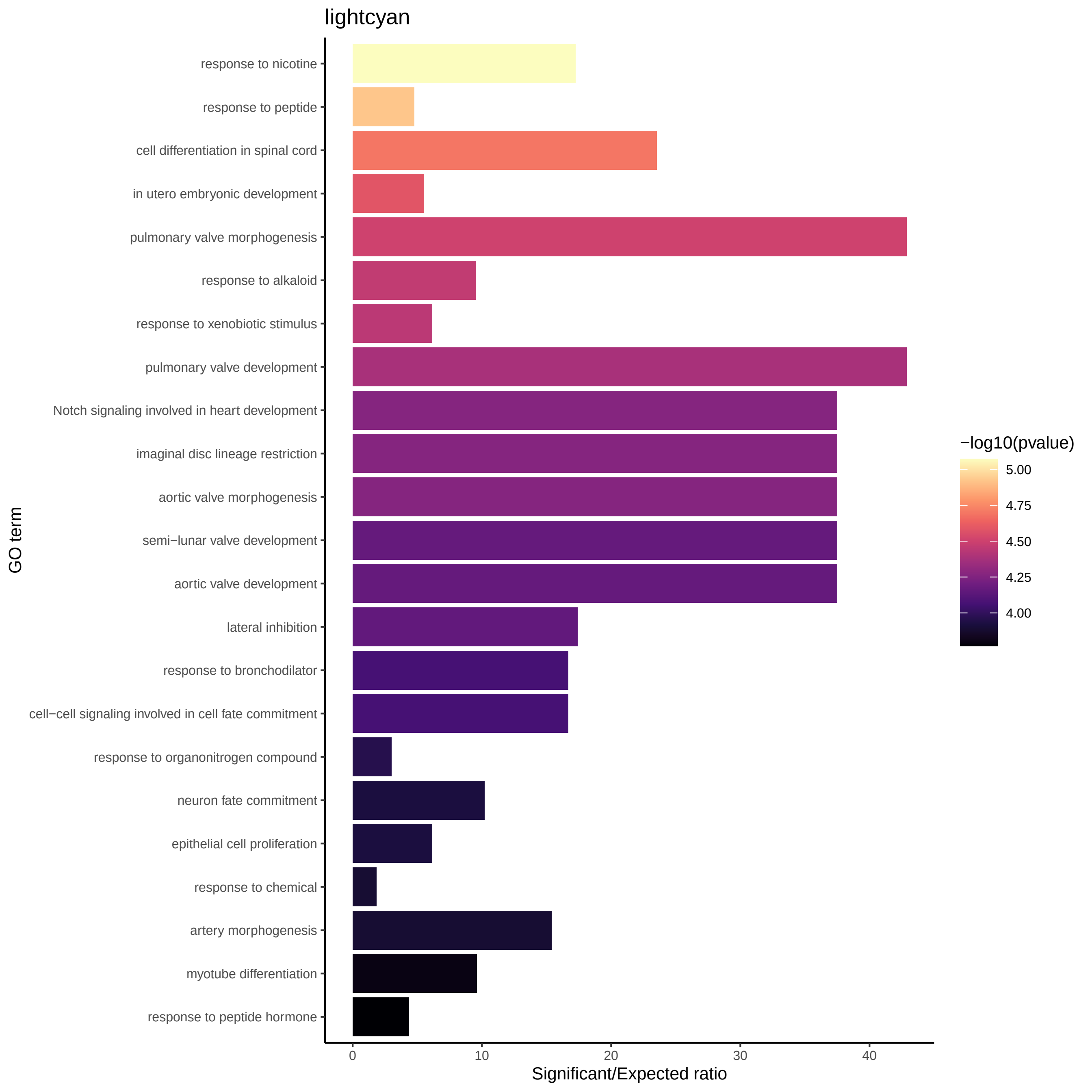
lightcyan
GO term
response to nicotine
response to peptide
cell differentiation in spinal cord
in utero embryonic development
pulmonary valve morphogenesis
response to alkaloid
response to xenobiotic stimulus
pulmonary valve development
Notch signaling involved in heart development
imaginal disc lineage restriction
aortic valve morphogenesis
semi−lunar valve development
aortic valve development
lateral inhibition
response to bronchodilator
cell−cell signaling involved in cell fate commitment
response to organonitrogen compound
neuron fate commitment
epithelial cell proliferation
response to chemical
artery morphogenesis
myotube differentiation
response to peptide hormone
0
10
20
30
40
Significant/Expected ratio
−log10(pvalue)
5.00
4.75
4.50
4.25
4.00

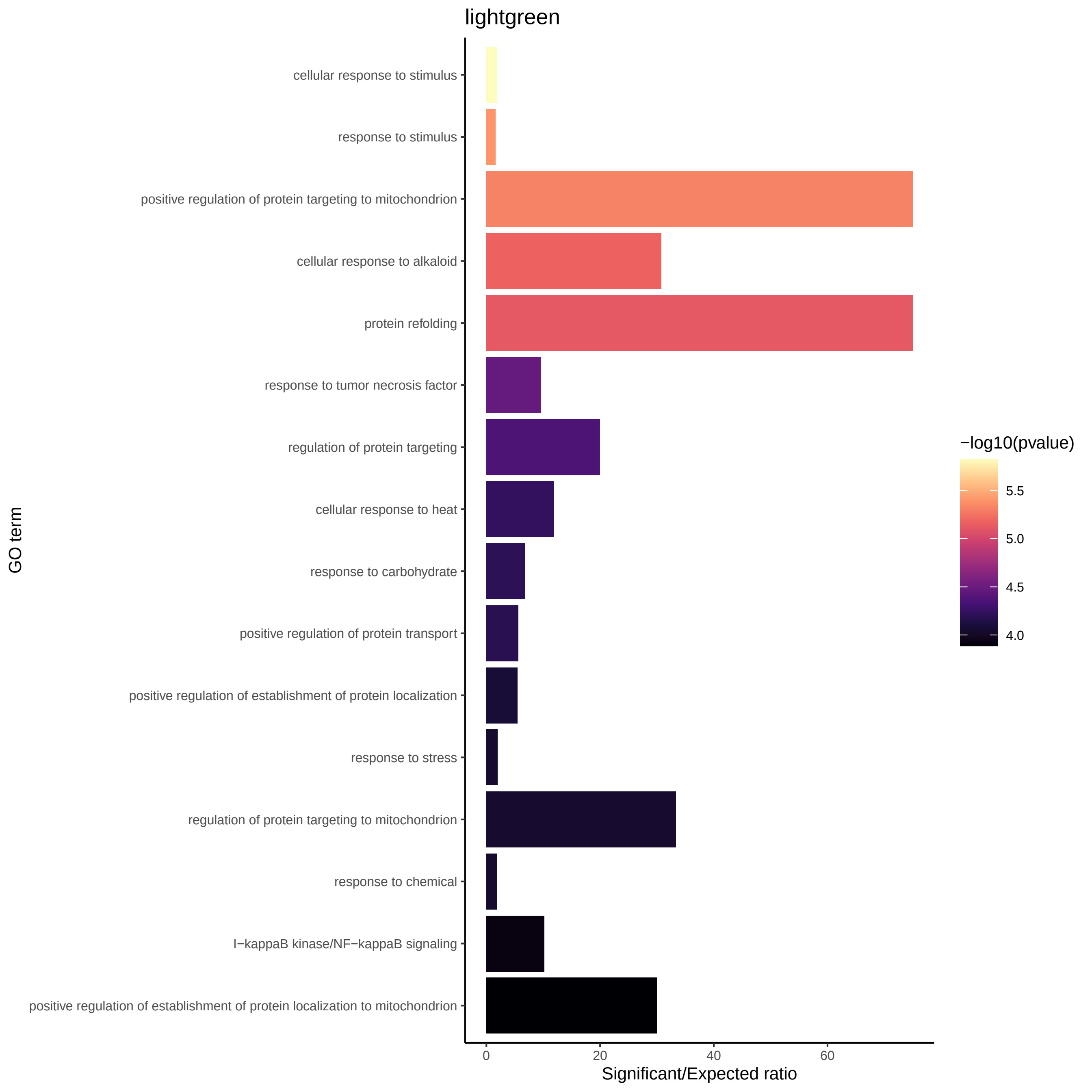
lightgreen
cellular response to stimulus
response to stimulus
positive regulation of protein targeting to mitochondrion
cellular response to alkaloid
protein refolding
response to tumor necrosis factor
regulation of protein targeting
cellular response to heat
response to carbohydrate
positive regulation of protein transport
positive regulation of establishment of protein localization
response to stress
regulation of protein targeting to mitochondrion
response to chemical
I−kappaB kinase/NF−kappaB signaling
positive regulation of establishment of protein localization to mitochondrion
GO term
0
20
40
60
Significant/Expected ratio
−log10(pvalue)
5.5
5.0
4.5
4.0

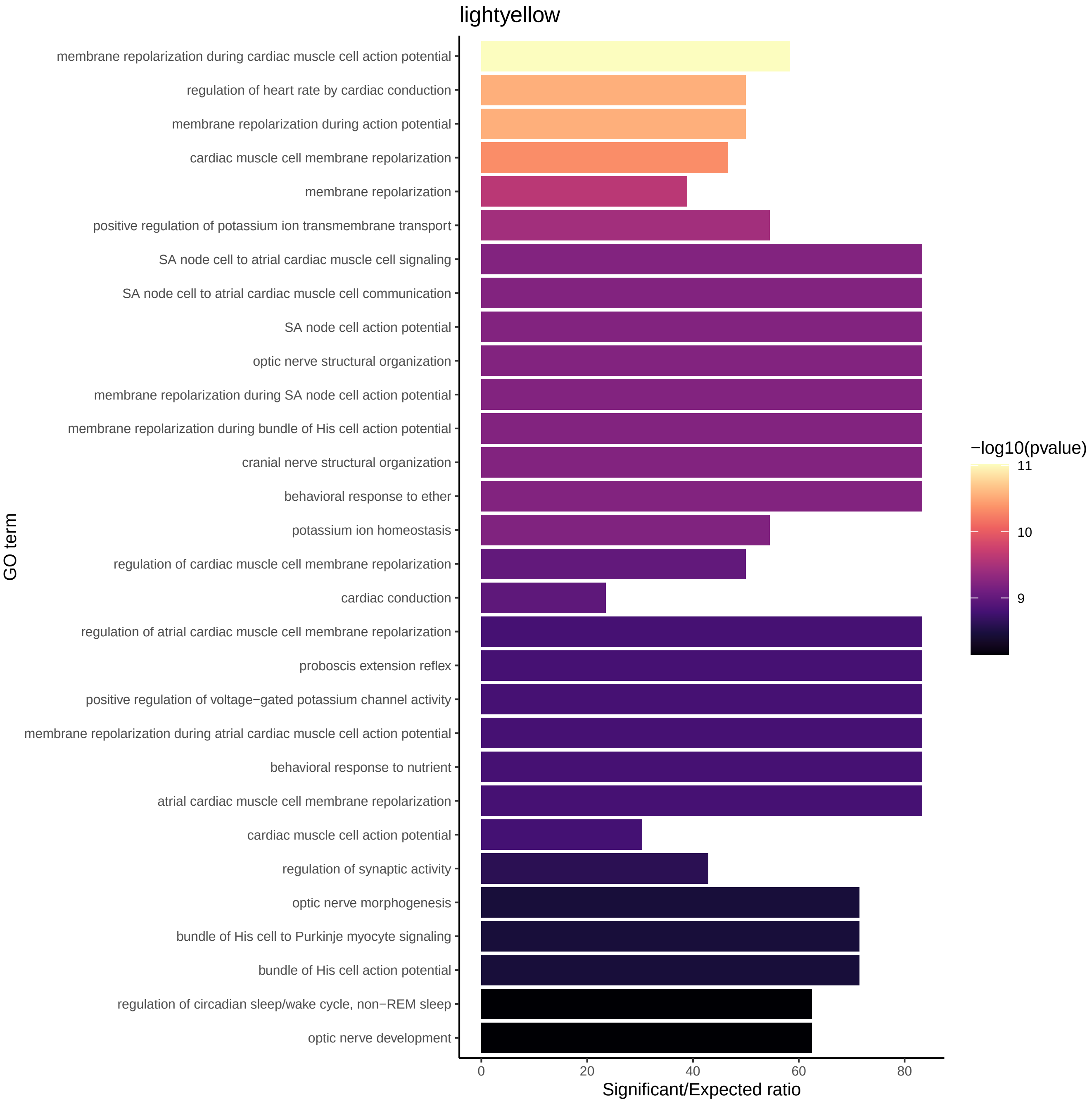

lightyellow
membrane repolarization during cardiac muscle cell action potential
regulation of heart rate by cardiac conduction
membrane repolarization during action potential
cardiac muscle cell membrane repolarization
membrane repolarization
positive regulation of potassium ion transmembrane transport
SA node cell to atrial cardiac muscle cell signaling
SA node cell to atrial cardiac muscle cell communication
SA node cell action potential
optic nerve structural organization
membrane repolarization during SA node cell action potential
membrane repolarization during bundle of His cell action potential
cranial nerve structural organization
behavioral response to ether
potassium ion homeostasis
regulation of cardiac muscle cell membrane repolarization
cardiac conduction
regulation of atrial cardiac muscle cell membrane repolarization
proboscis extension reflex
positive regulation of voltage−gated potassium channel activity
membrane repolarization during atrial cardiac muscle cell action potential
behavioral response to nutrient
atrial cardiac muscle cell membrane repolarization
cardiac muscle cell action potential
regulation of synaptic activity
optic nerve morphogenesis
bundle of His cell to Purkinje myocyte signaling
bundle of His cell action potential
regulation of circadian sleep/wake cycle, non−REM sleep
optic nerve development
GO term
0
20
40
60
80
Significant/Expected ratio
−log10(pvalue)
11
10
9

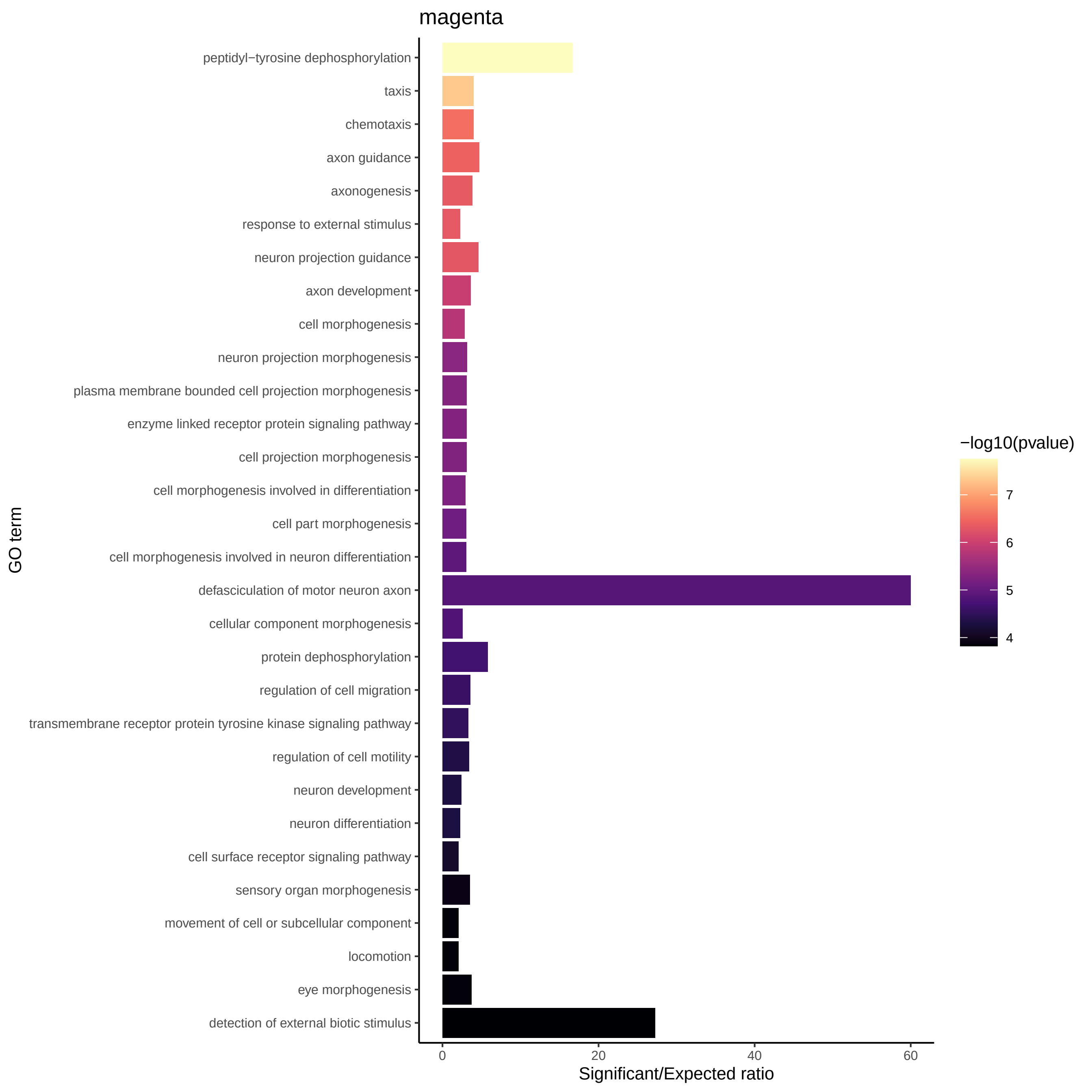
magenta
peptidyl−tyrosine dephosphorylation
taxis
chemotaxis
axon guidance
axonogenesis
response to external stimulus
neuron projection guidance
axon development
cell morphogenesis
neuron projection morphogenesis
plasma membrane bounded cell projection morphogenesis
enzyme linked receptor protein signaling pathway
cell projection morphogenesis
cell morphogenesis involved in differentiation
cell part morphogenesis
cell morphogenesis involved in neuron differentiation
defasciculation of motor neuron axon
cellular component morphogenesis
protein dephosphorylation
regulation of cell migration
transmembrane receptor protein tyrosine kinase signaling pathway
regulation of cell motility
neuron development
neuron differentiation
cell surface receptor signaling pathway
sensory organ morphogenesis
movement of cell or subcellular component
locomotion
eye morphogenesis
detection of external biotic stimulus
GO term
0
20
40
60
Significant/Expected ratio
−log10(pvalue)
7
6
5
4

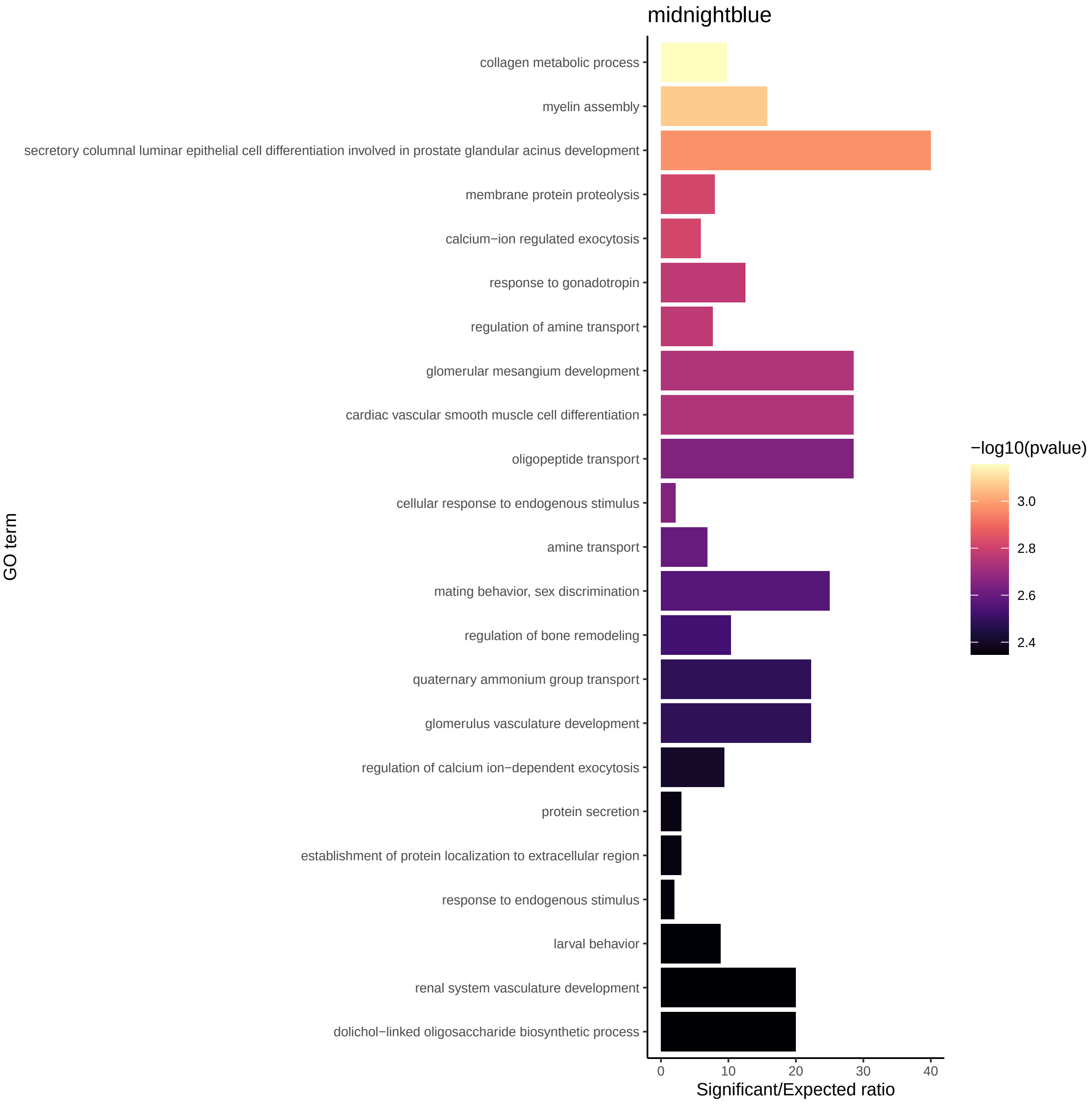
midnightblue
collagen metabolic process
myelin assembly
secretory columnal luminar epithelial cell differentiation involved in prostate glandular acinus development
membrane protein proteolysis
calcium−ion regulated exocytosis
response to gonadotropin
regulation of amine transport
glomerular mesangium development
cardiac vascular smooth muscle cell differentiation
oligopeptide transport
cellular response to endogenous stimulus
amine transport
mating behavior, sex discrimination
regulation of bone remodeling
quaternary ammonium group transport
glomerulus vasculature development
regulation of calcium ion−dependent exocytosis
protein secretion
establishment of protein localization to extracellular region
response to endogenous stimulus
larval behavior
renal system vasculature development
dolichol−linked oligosaccharide biosynthetic process
GO term
0
10
20
30
40
Significant/Expected ratio
−log10(pvalue)
3.0
2.8
2.6
2.4

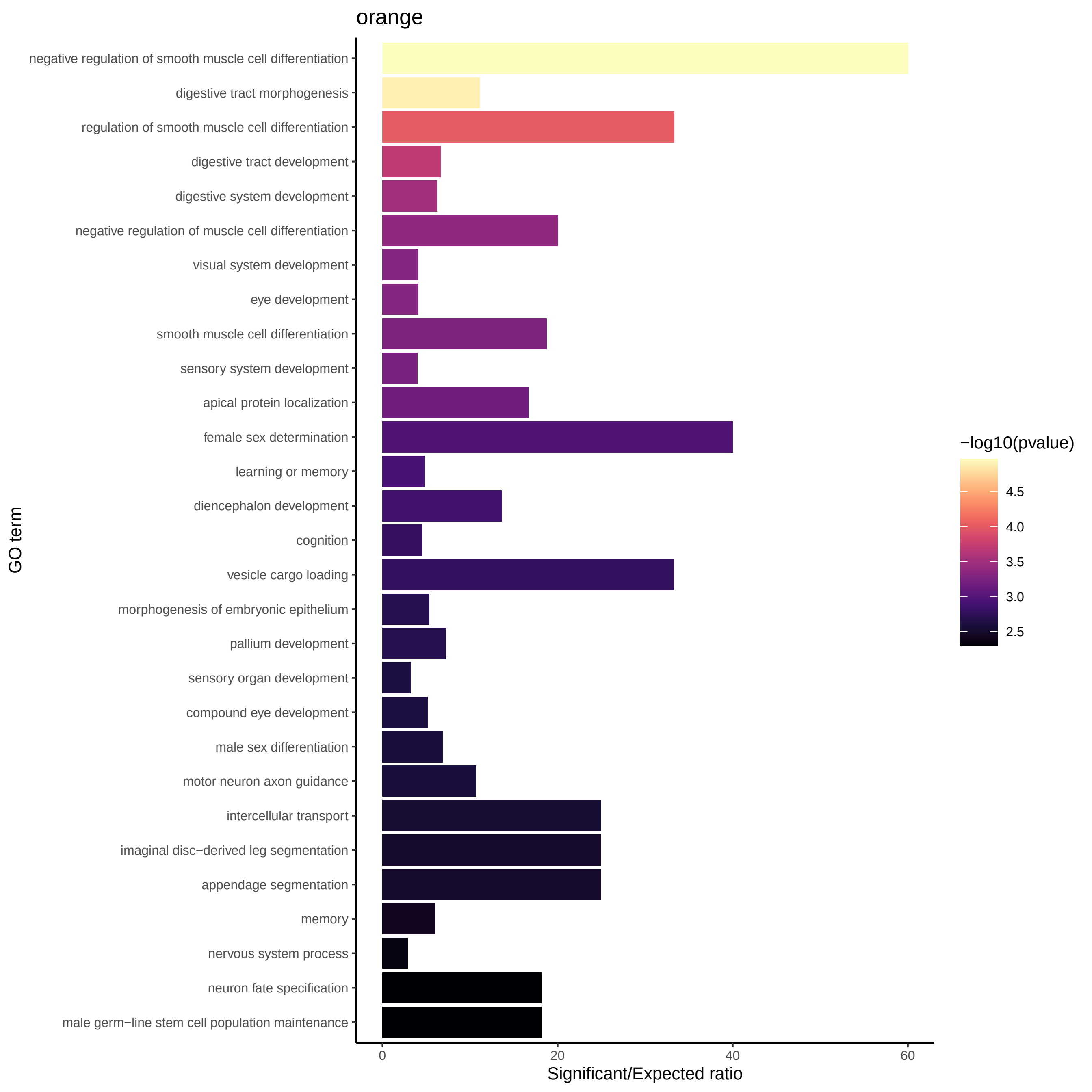
orange
negative regulation of smooth muscle cell differentiation
digestive tract morphogenesis
regulation of smooth muscle cell differentiation
digestive tract development
digestive system development
negative regulation of muscle cell differentiation
visual system development
eye development
smooth muscle cell differentiation
sensory system development
apical protein localization
female sex determination
learning or memory
diencephalon development
cognition
vesicle cargo loading
morphogenesis of embryonic epithelium
pallium development
sensory organ development
compound eye development
male sex differentiation
motor neuron axon guidance
intercellular transport
imaginal disc−derived leg segmentation
appendage segmentation
memory
nervous system process
neuron fate specification
male germ−line stem cell population maintenance
GO term
0
20
40
60
Significant/Expected ratio
−log10(pvalue)
4.5
4.0
3.5
3.0
2.5

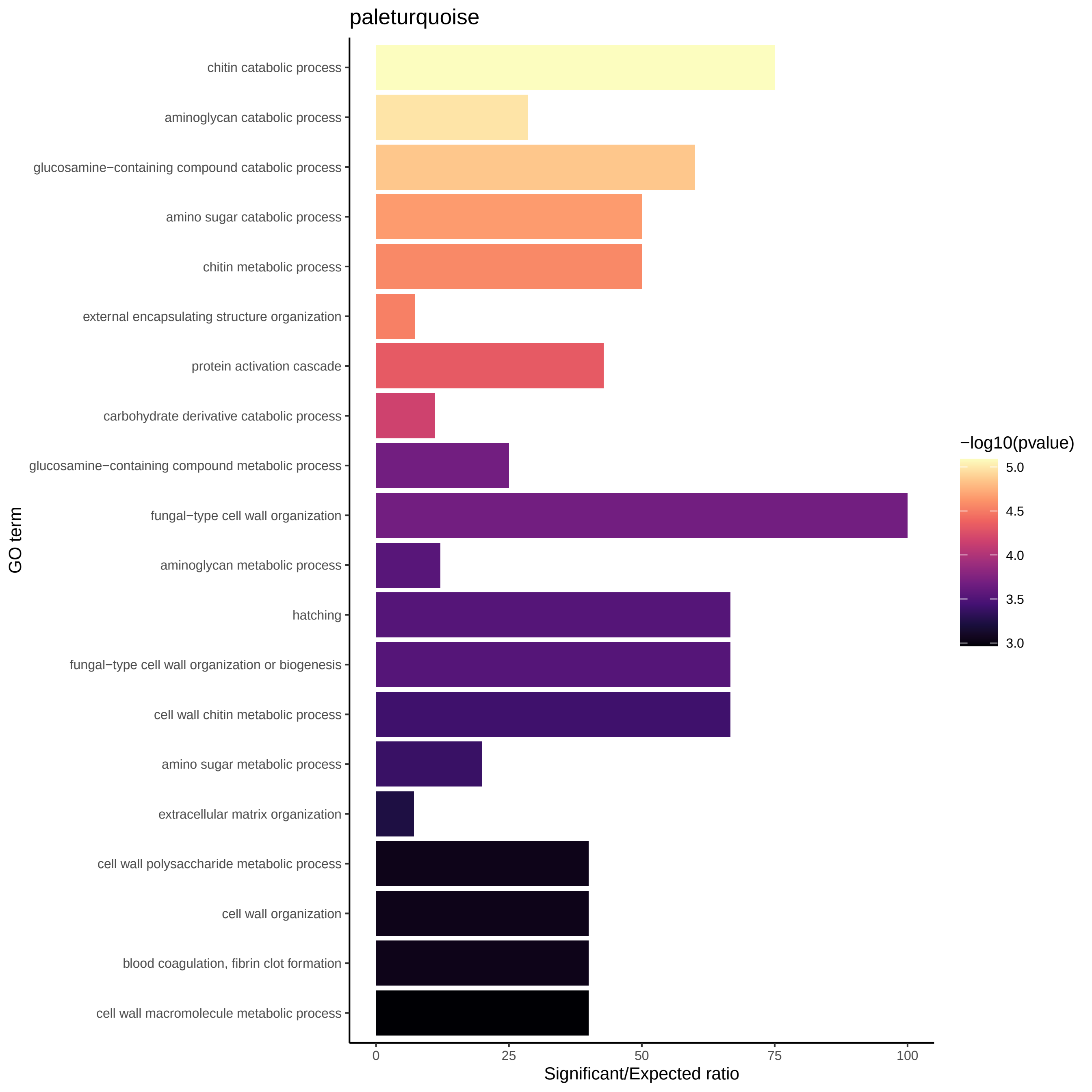

paleturquoise
chitin catabolic process
aminoglycan catabolic process
glucosamine−containing compound catabolic process
amino sugar catabolic process
chitin metabolic process
external encapsulating structure organization
protein activation cascade
carbohydrate derivative catabolic process
glucosamine−containing compound metabolic process
fungal−type cell wall organization
aminoglycan metabolic process
hatching
fungal−type cell wall organization or biogenesis
cell wall chitin metabolic process
amino sugar metabolic process
extracellular matrix organization
cell wall polysaccharide metabolic process
cell wall organization
blood coagulation, fibrin clot formation
cell wall macromolecule metabolic process
GO term
0
25
50
75
100
Significant/Expected ratio
−log10(pvalue)
5.0
4.5
4.0
3.5
3.0

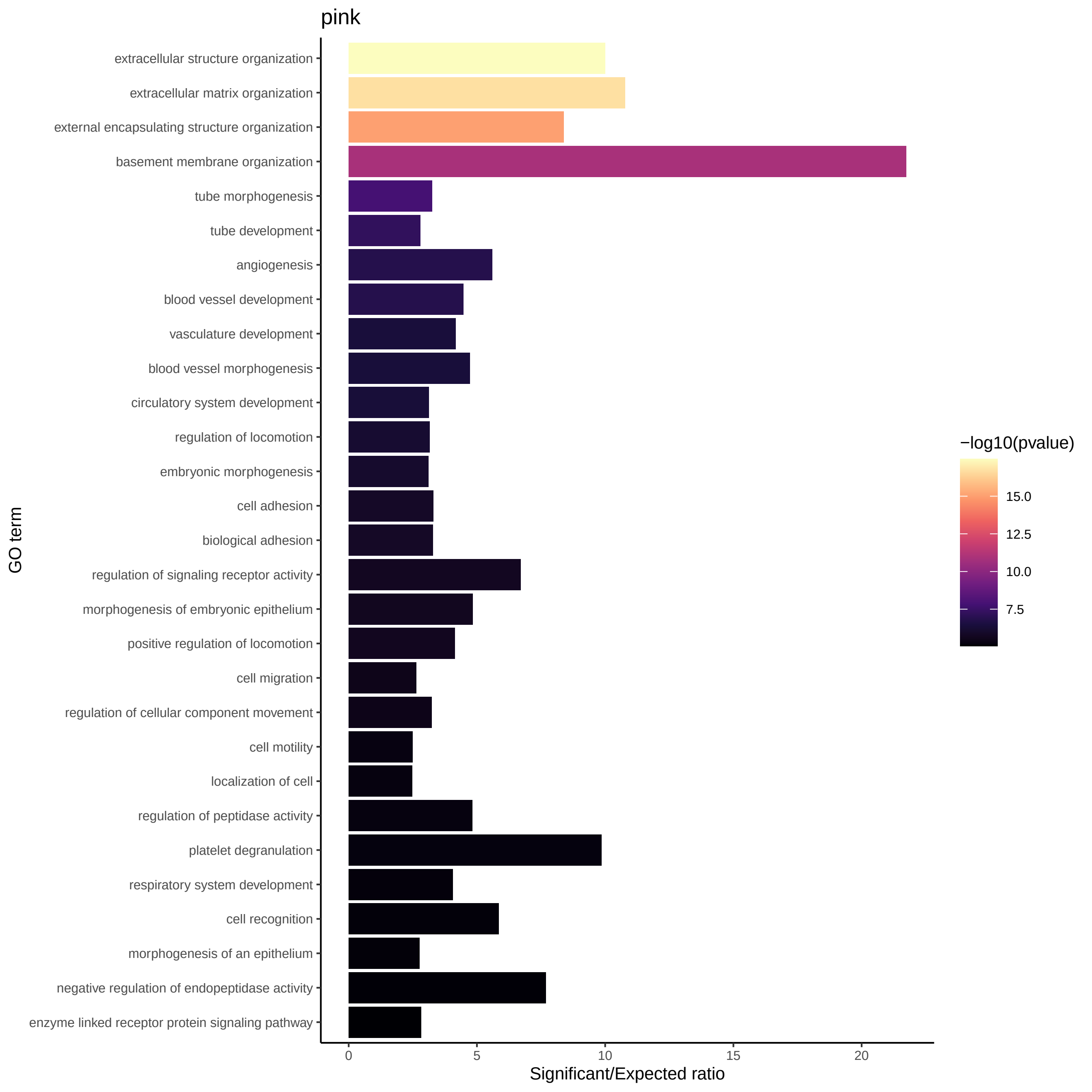

pink
extracellular structure organization
extracellular matrix organization
external encapsulating structure organization
basement membrane organization
tube morphogenesis
tube development
angiogenesis
blood vessel development
vasculature development
blood vessel morphogenesis
circulatory system development
regulation of locomotion
embryonic morphogenesis
cell adhesion
biological adhesion
regulation of signaling receptor activity
morphogenesis of embryonic epithelium
positive regulation of locomotion
cell migration
regulation of cellular component movement
cell motility
localization of cell
regulation of peptidase activity
platelet degranulation
respiratory system development
cell recognition
morphogenesis of an epithelium
negative regulation of endopeptidase activity
enzyme linked receptor protein signaling pathway
GO term
0
5
10
15
20
Significant/Expected ratio
−log10(pvalue)
15.0
12.5
10.0
7.5

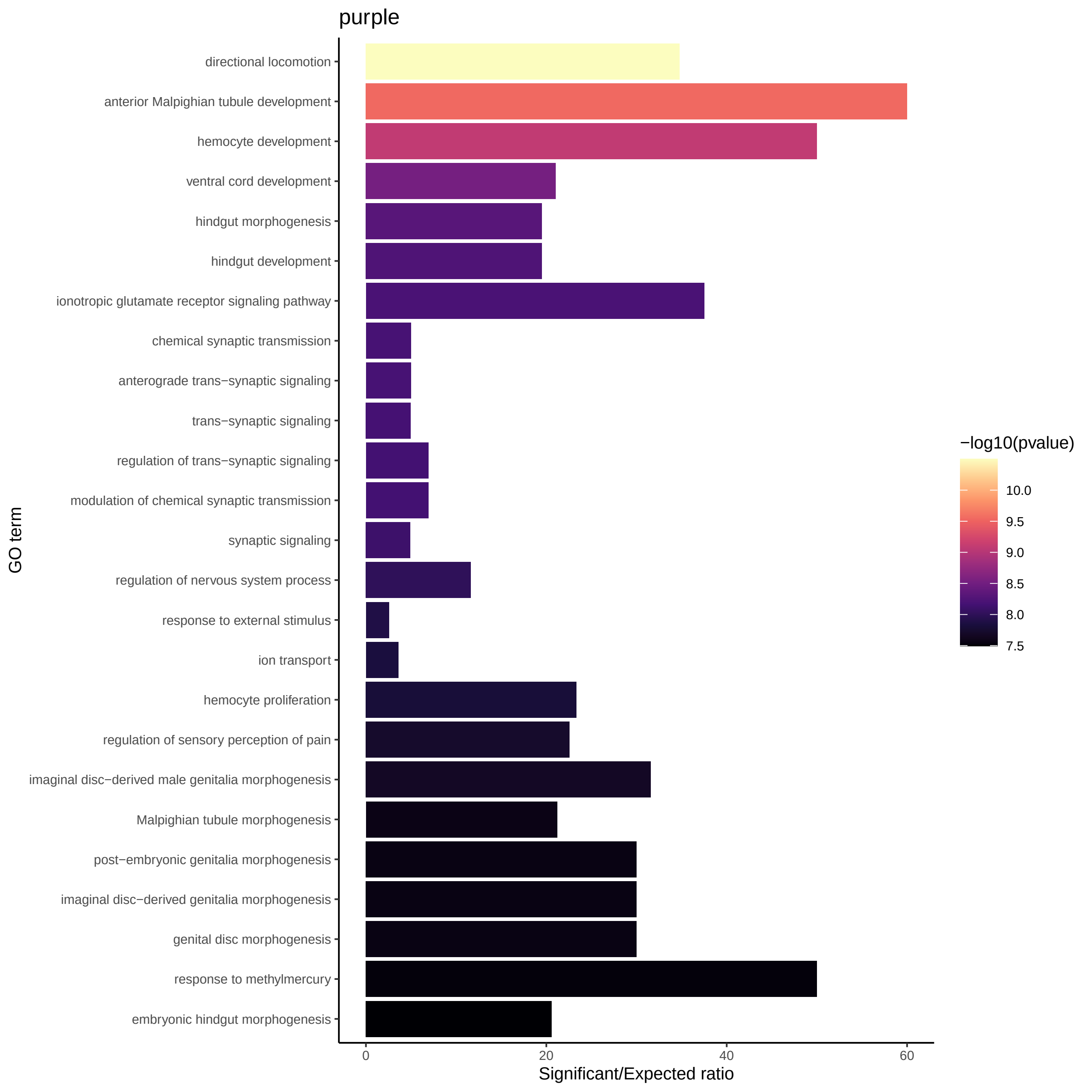
purple
directional locomotion
anterior Malpighian tubule development
hemocyte development
ventral cord development
hindgut morphogenesis
hindgut development
ionotropic glutamate receptor signaling pathway
chemical synaptic transmission
anterograde trans−synaptic signaling
trans−synaptic signaling
regulation of trans−synaptic signaling
modulation of chemical synaptic transmission
synaptic signaling
regulation of nervous system process
response to external stimulus
ion transport
hemocyte proliferation
regulation of sensory perception of pain
imaginal disc−derived male genitalia morphogenesis
Malpighian tubule morphogenesis
post−embryonic genitalia morphogenesis
imaginal disc−derived genitalia morphogenesis
genital disc morphogenesis
response to methylmercury
embryonic hindgut morphogenesis
GO term
0
20
40
60
Significant/Expected ratio
−log10(pvalue)
10.0
9.5
9.0
8.5
8.0
7.5

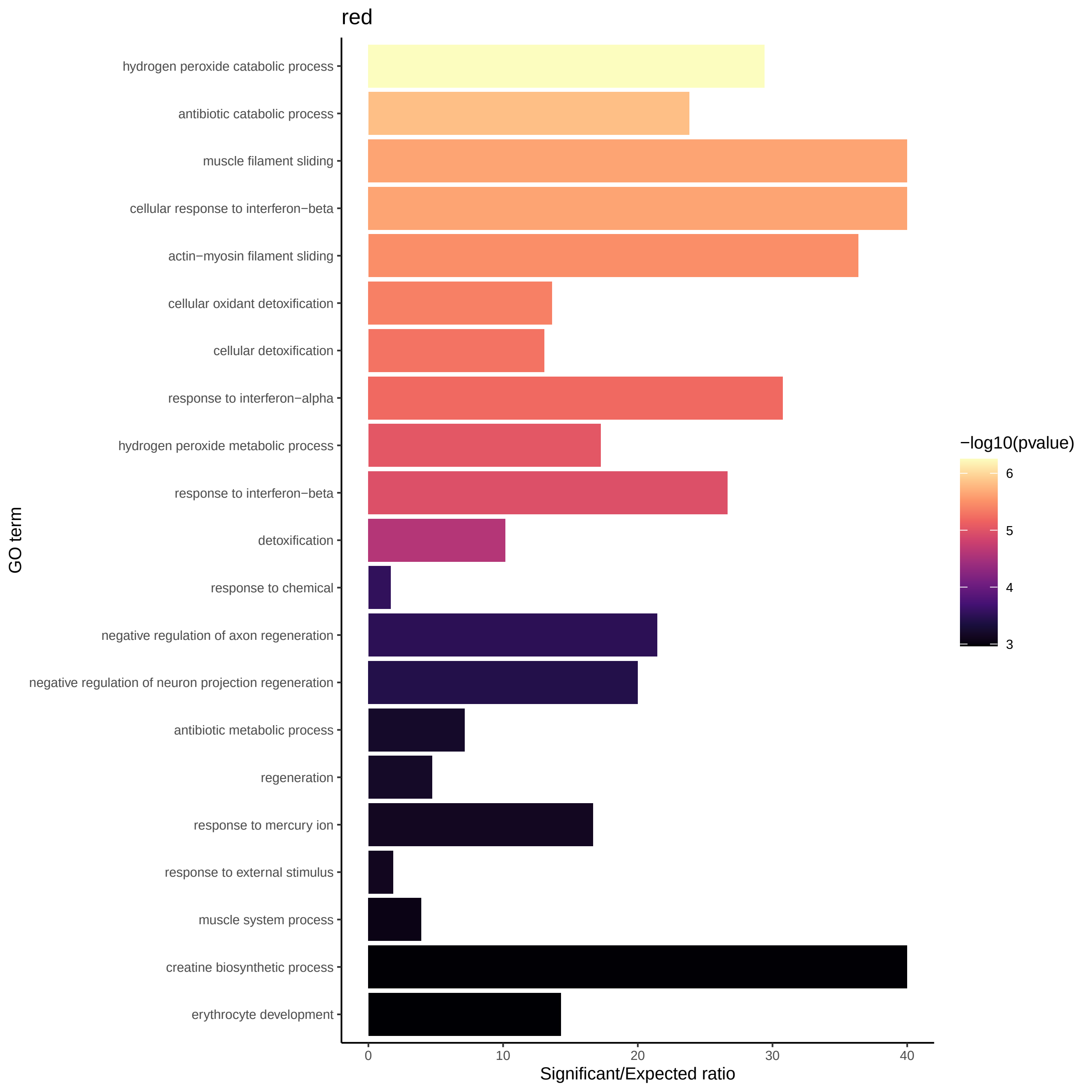

red
hydrogen peroxide catabolic process
antibiotic catabolic process
muscle filament sliding
cellular response to interferon−beta
actin−myosin filament sliding
cellular oxidant detoxification
cellular detoxification
response to interferon−alpha
hydrogen peroxide metabolic process
response to interferon−beta
detoxification
response to chemical
negative regulation of axon regeneration
negative regulation of neuron projection regeneration
antibiotic metabolic process
regeneration
response to mercury ion
response to external stimulus
muscle system process
creatine biosynthetic process
erythrocyte development
GO term
0
10
20
30
40
Significant/Expected ratio
−log10(pvalue)
6
5
4
3

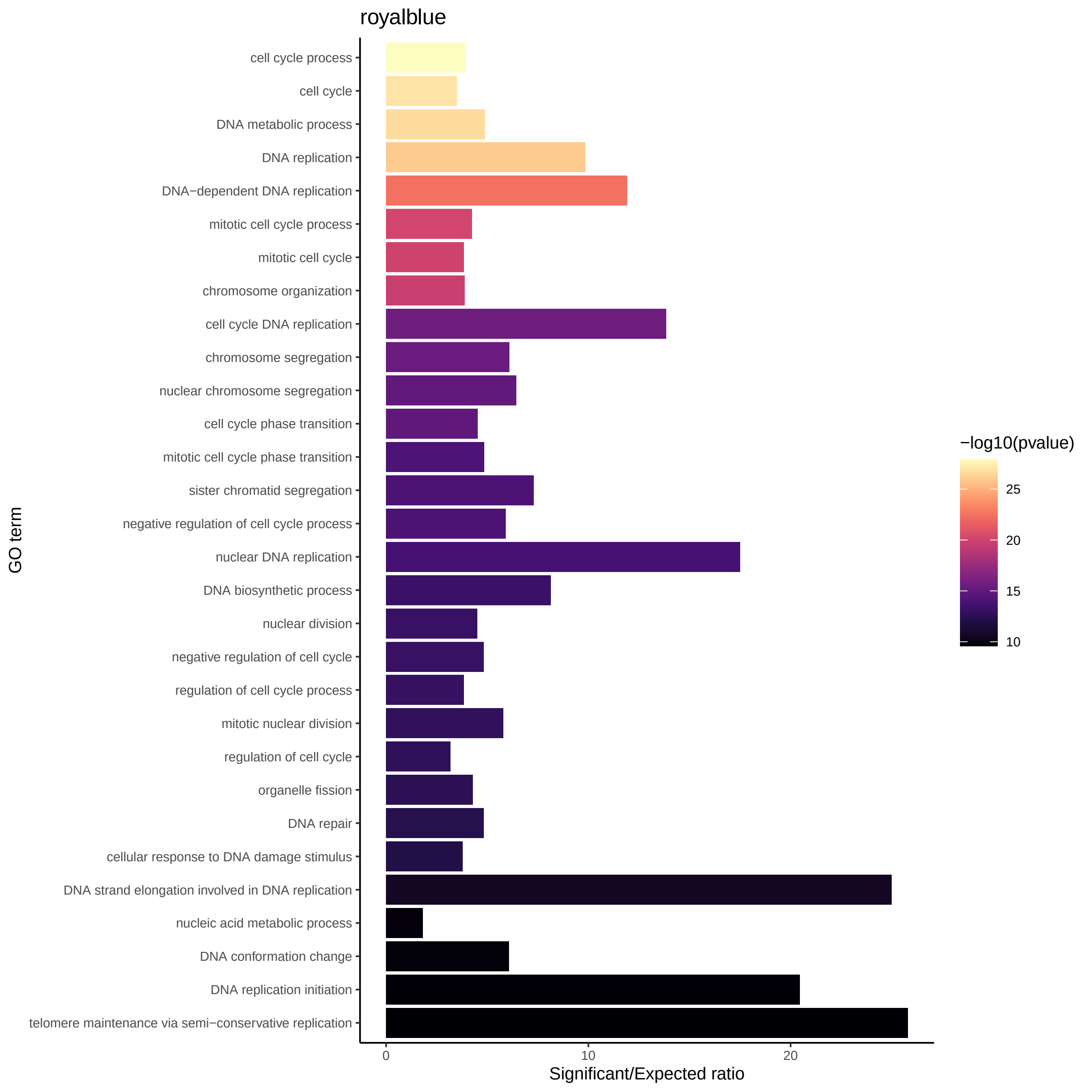
royalblue
cell cycle process
cell cycle
DNA metabolic process
DNA replication
DNA−dependent DNA replication
mitotic cell cycle process
mitotic cell cycle
chromosome organization
cell cycle DNA replication
chromosome segregation
nuclear chromosome segregation
cell cycle phase transition
mitotic cell cycle phase transition
sister chromatid segregation
negative regulation of cell cycle process
nuclear DNA replication
DNA biosynthetic process
nuclear division
negative regulation of cell cycle
regulation of cell cycle process
mitotic nuclear division
regulation of cell cycle
organelle fission
DNA repair
cellular response to DNA damage stimulus
DNA strand elongation involved in DNA replication
nucleic acid metabolic process
DNA conformation change
DNA replication initiation
telomere maintenance via semi−conservative replication
GO term
0
10
20
Significant/Expected ratio
−log10(pvalue)
25
20
15
10

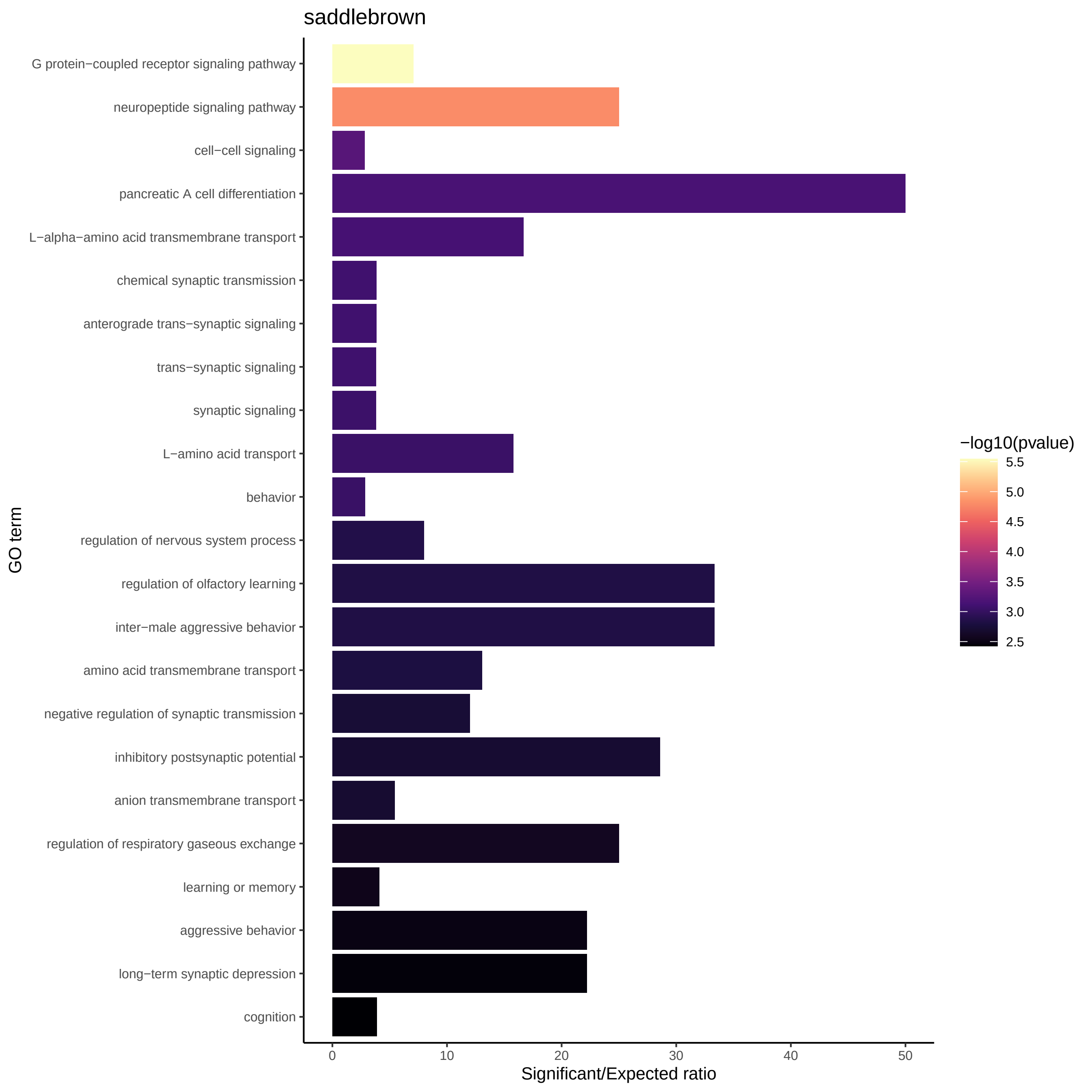
saddlebrown
G protein−coupled receptor signaling pathway
neuropeptide signaling pathway
cell−cell signaling
pancreatic A cell differentiation
L−alpha−amino acid transmembrane transport
chemical synaptic transmission
anterograde trans−synaptic signaling
trans−synaptic signaling
synaptic signaling
L−amino acid transport
behavior
regulation of nervous system process
regulation of olfactory learning
inter−male aggressive behavior
amino acid transmembrane transport
negative regulation of synaptic transmission
inhibitory postsynaptic potential
anion transmembrane transport
regulation of respiratory gaseous exchange
learning or memory
aggressive behavior
long−term synaptic depression
cognition
GO term
0
10
20
30
40
50
Significant/Expected ratio
−log10(pvalue)
5.5
5.0
4.5
4.0
3.5
3.0
2.5

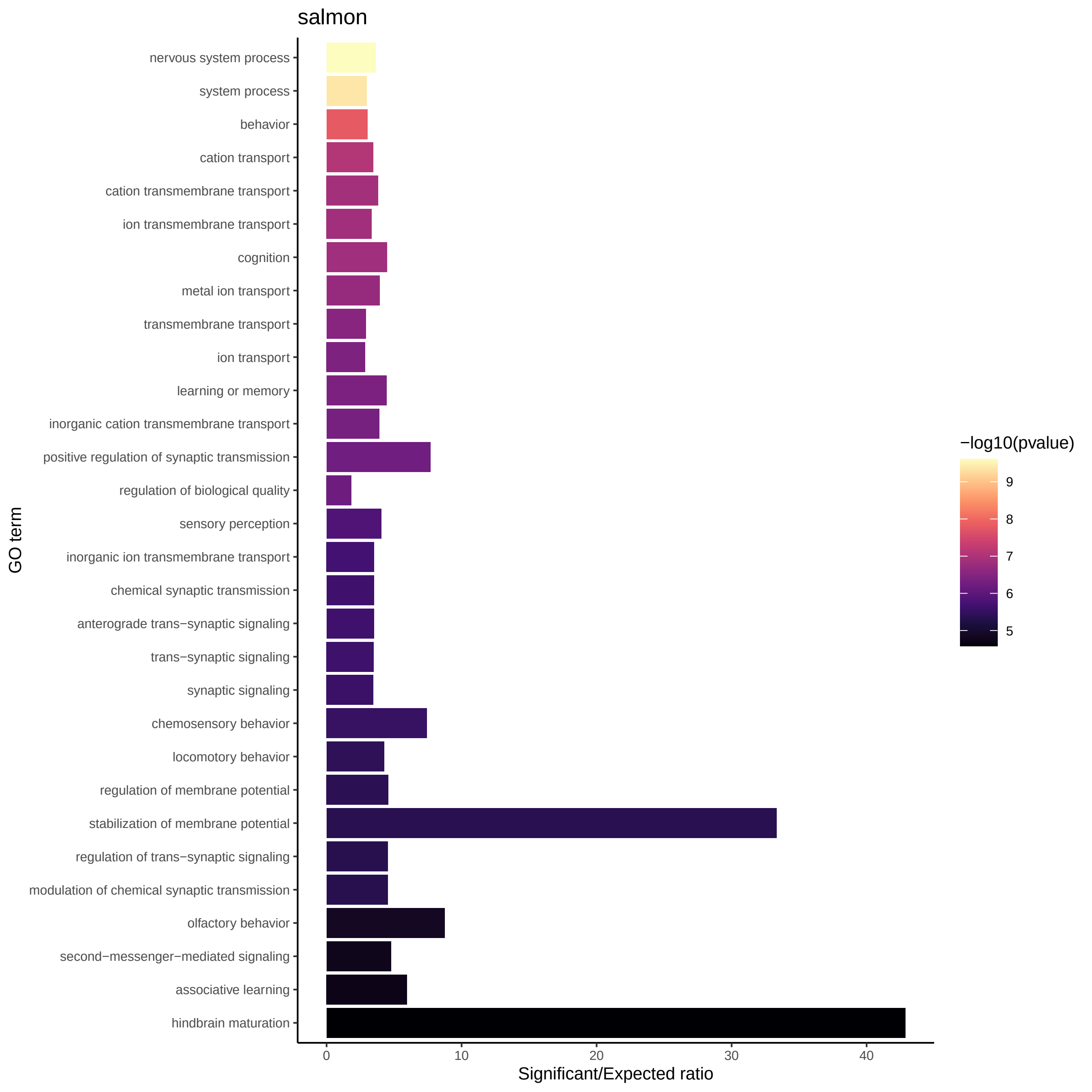

salmon
nervous system process
system process
behavior
cation transport
cation transmembrane transport
ion transmembrane transport
cognition
metal ion transport
transmembrane transport
ion transport
learning or memory
inorganic cation transmembrane transport
positive regulation of synaptic transmission
regulation of biological quality
sensory perception
inorganic ion transmembrane transport
chemical synaptic transmission
anterograde trans−synaptic signaling
trans−synaptic signaling
synaptic signaling
chemosensory behavior
locomotory behavior
regulation of membrane potential
stabilization of membrane potential
regulation of trans−synaptic signaling
modulation of chemical synaptic transmission
olfactory behavior
second−messenger−mediated signaling
associative learning
hindbrain maturation
GO term
0
10
20
30
40
Significant/Expected ratio
−log10(pvalue)
9
8
7
6
5

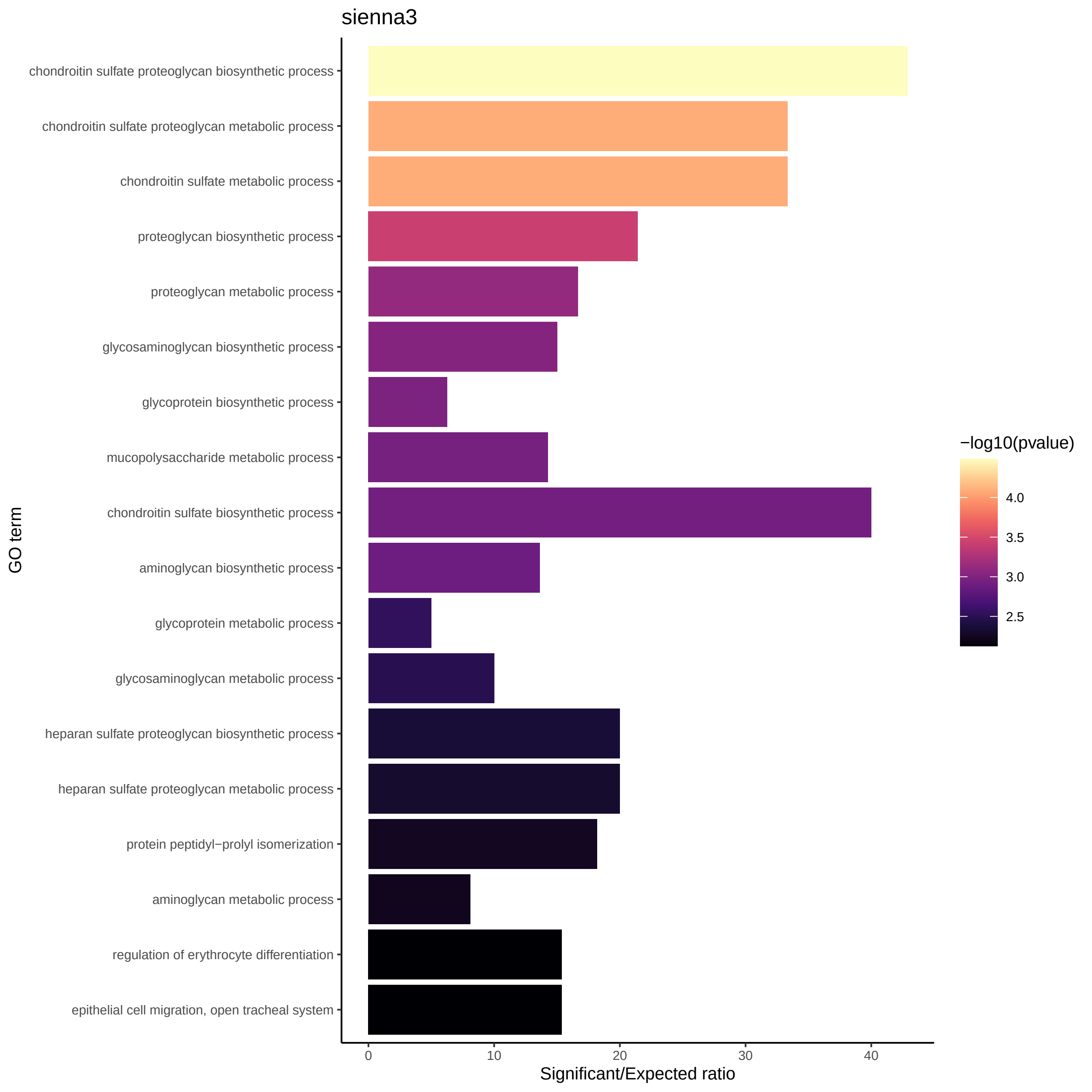
sienna3
chondroitin sulfate proteoglycan biosynthetic process
chondroitin sulfate proteoglycan metabolic process
chondroitin sulfate metabolic process
proteoglycan biosynthetic process
proteoglycan metabolic process
glycosaminoglycan biosynthetic process
glycoprotein biosynthetic process
mucopolysaccharide metabolic process
chondroitin sulfate biosynthetic process
aminoglycan biosynthetic process
glycoprotein metabolic process
glycosaminoglycan metabolic process
heparan sulfate proteoglycan biosynthetic process
heparan sulfate proteoglycan metabolic process
protein peptidyl−prolyl isomerization
aminoglycan metabolic process
regulation of erythrocyte differentiation
epithelial cell migration, open tracheal system
GO term
0
10
20
30
40
Significant/Expected ratio
−log10(pvalue)
4.0
3.5
3.0
2.5

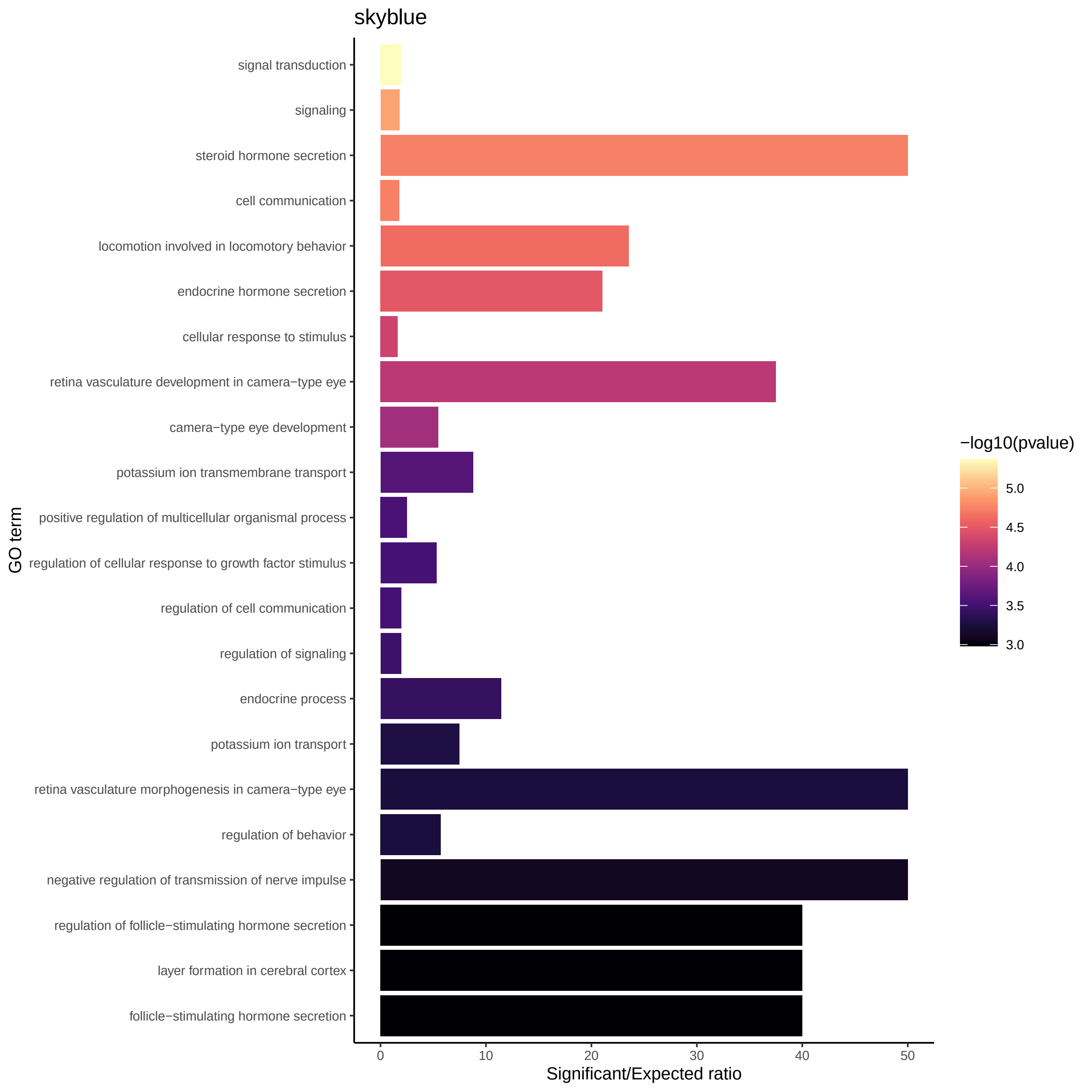
skyblue
signal transduction
signaling
steroid hormone secretion
cell communication
locomotion involved in locomotory behavior
endocrine hormone secretion
cellular response to stimulus
retina vasculature development in camera−type eye
camera−type eye development
potassium ion transmembrane transport
positive regulation of multicellular organismal process
regulation of cellular response to growth factor stimulus
regulation of cell communication
regulation of signaling
endocrine process
potassium ion transport
retina vasculature morphogenesis in camera−type eye
regulation of behavior
negative regulation of transmission of nerve impulse
regulation of follicle−stimulating hormone secretion
layer formation in cerebral cortex
follicle−stimulating hormone secretion
GO term
0
10
20
30
40
50
Significant/Expected ratio
−log10(pvalue)
5.0
4.5
4.0
3.5
3.0

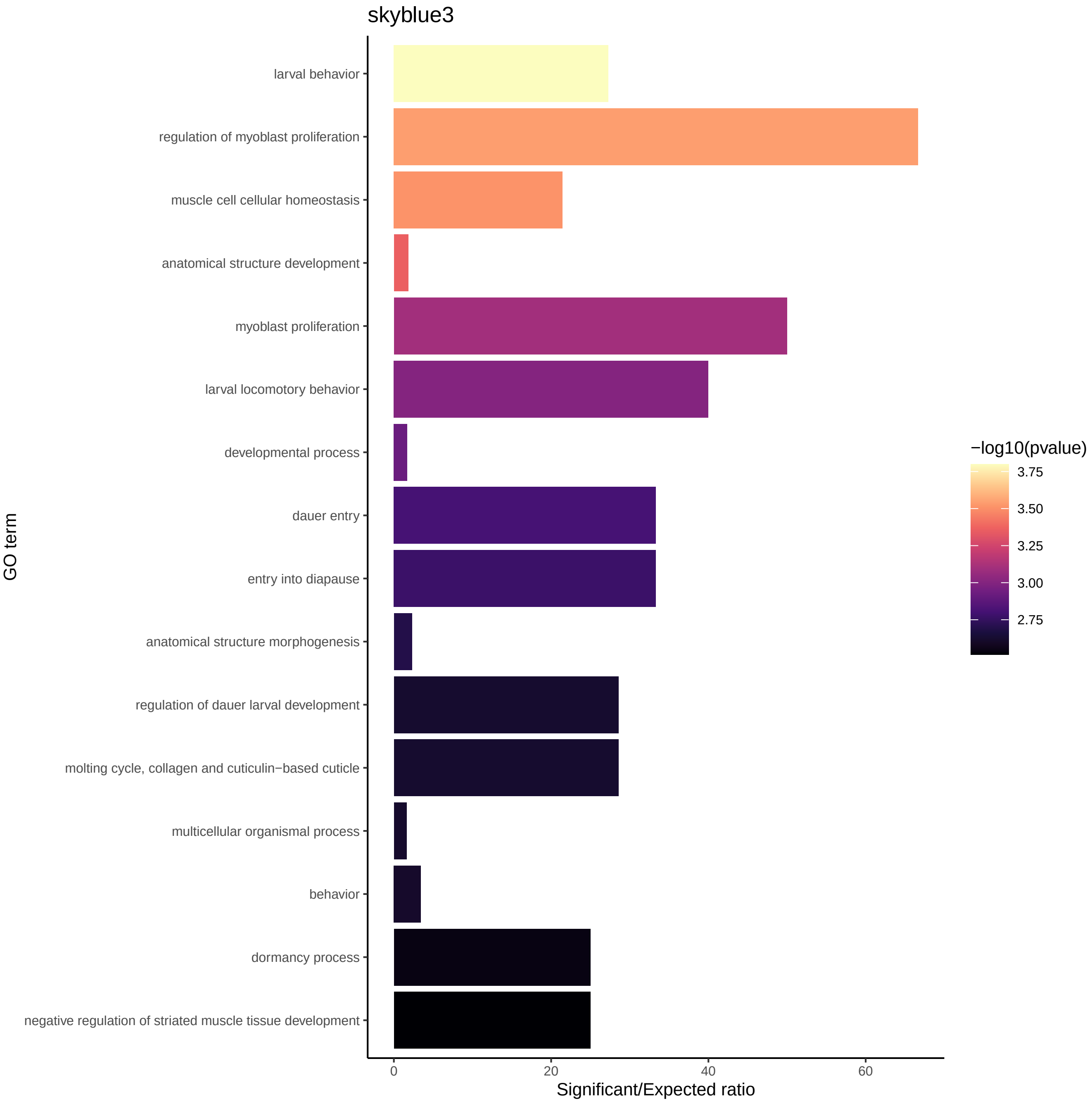
skyblue3
GO term
larval behavior
regulation of myoblast proliferation
muscle cell cellular homeostasis
anatomical structure development
myoblast proliferation
larval locomotory behavior
developmental process
dauer entry
entry into diapause
anatomical structure morphogenesis
regulation of dauer larval development
molting cycle, collagen and cuticulin−based cuticle
multicellular organismal process
behavior
dormancy process
negative regulation of striated muscle tissue development
0
20
40
60
Significant/Expected ratio
−log10(pvalue)
3.75
3.50
3.25
3.00
2.75

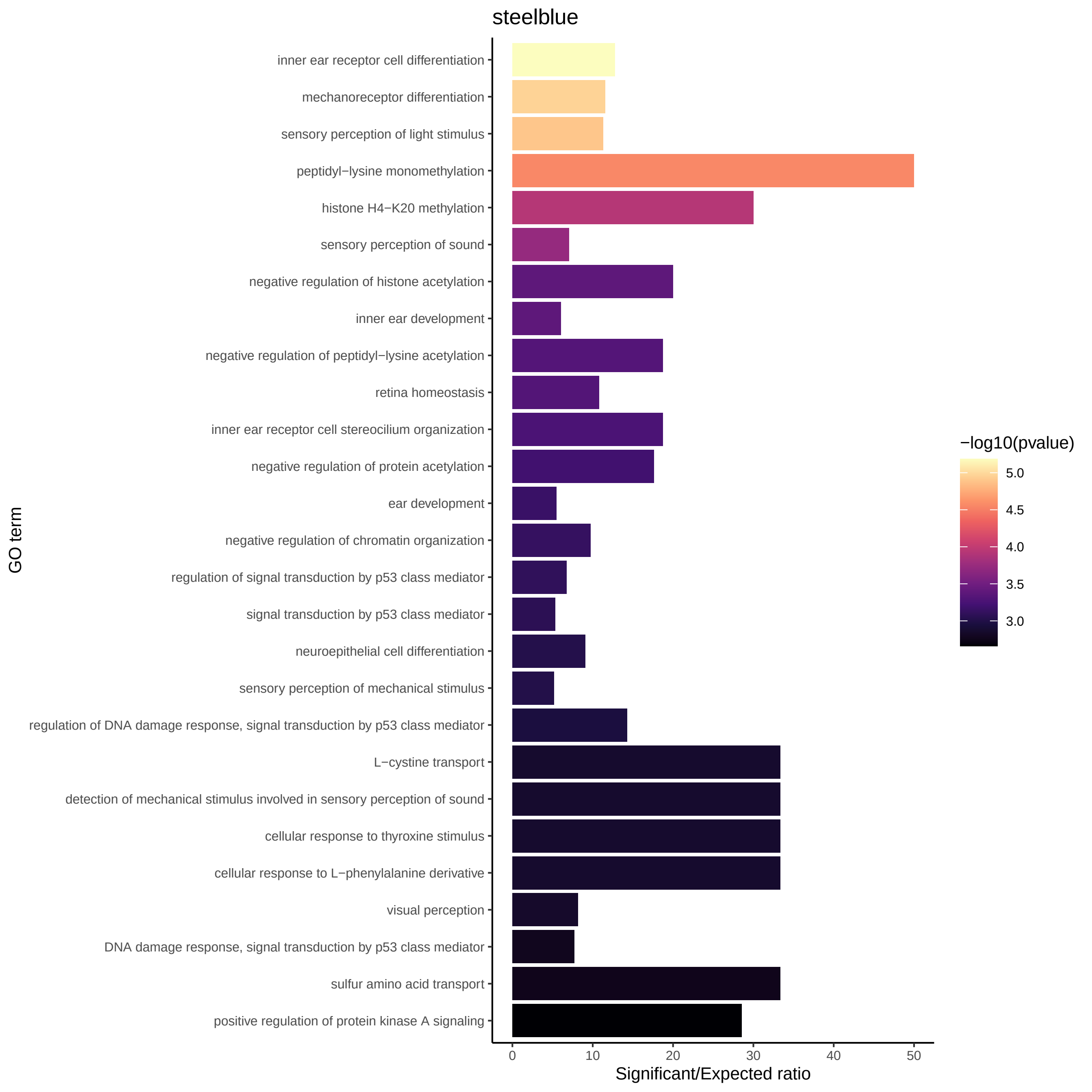

steelblue
inner ear receptor cell differentiation
mechanoreceptor differentiation
sensory perception of light stimulus
peptidyl−lysine monomethylation
histone H4−K20 methylation
sensory perception of sound
negative regulation of histone acetylation
inner ear development
negative regulation of peptidyl−lysine acetylation
retina homeostasis
inner ear receptor cell stereocilium organization
negative regulation of protein acetylation
ear development
negative regulation of chromatin organization
regulation of signal transduction by p53 class mediator
signal transduction by p53 class mediator
neuroepithelial cell differentiation
sensory perception of mechanical stimulus
regulation of DNA damage response, signal transduction by p53 class mediator
L−cystine transport
detection of mechanical stimulus involved in sensory perception of sound
cellular response to thyroxine stimulus
cellular response to L−phenylalanine derivative
visual perception
DNA damage response, signal transduction by p53 class mediator
sulfur amino acid transport
positive regulation of protein kinase A signaling
GO term
0
10
20
30
40
50
Significant/Expected ratio
−log10(pvalue)
5.0
4.5
4.0
3.5
3.0

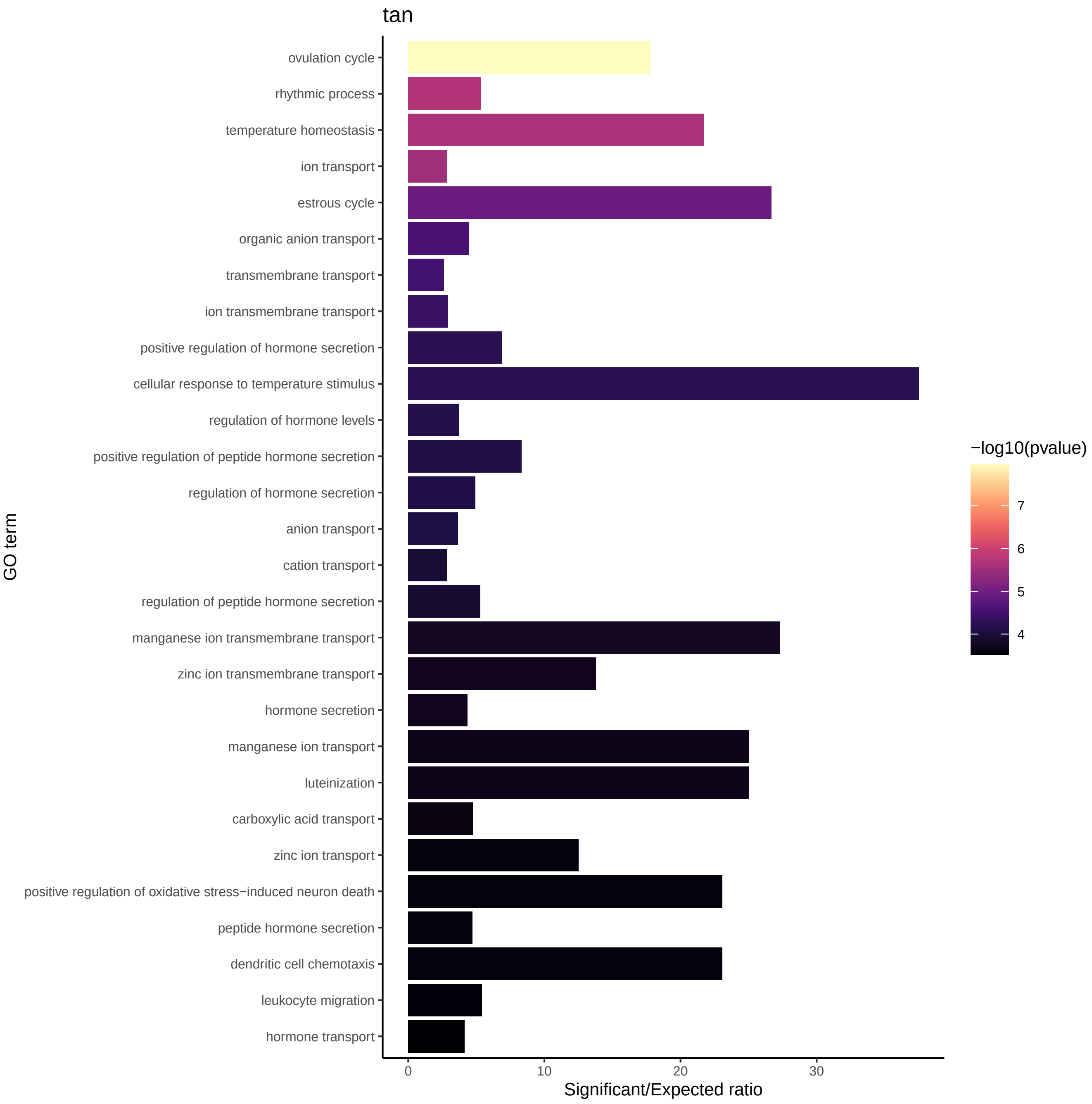
tan
ovulation cycle
rhythmic process
temperature homeostasis
ion transport
estrous cycle
organic anion transport
transmembrane transport
ion transmembrane transport
positive regulation of hormone secretion
cellular response to temperature stimulus
regulation of hormone levels
positive regulation of peptide hormone secretion
regulation of hormone secretion
anion transport
cation transport
regulation of peptide hormone secretion
manganese ion transmembrane transport
zinc ion transmembrane transport
hormone secretion
manganese ion transport
luteinization
carboxylic acid transport
zinc ion transport
positive regulation of oxidative stress−induced neuron death
peptide hormone secretion
dendritic cell chemotaxis
leukocyte migration
hormone transport
GO term
0
10
20
30
Significant/Expected ratio
−log10(pvalue)
7
6
5
4

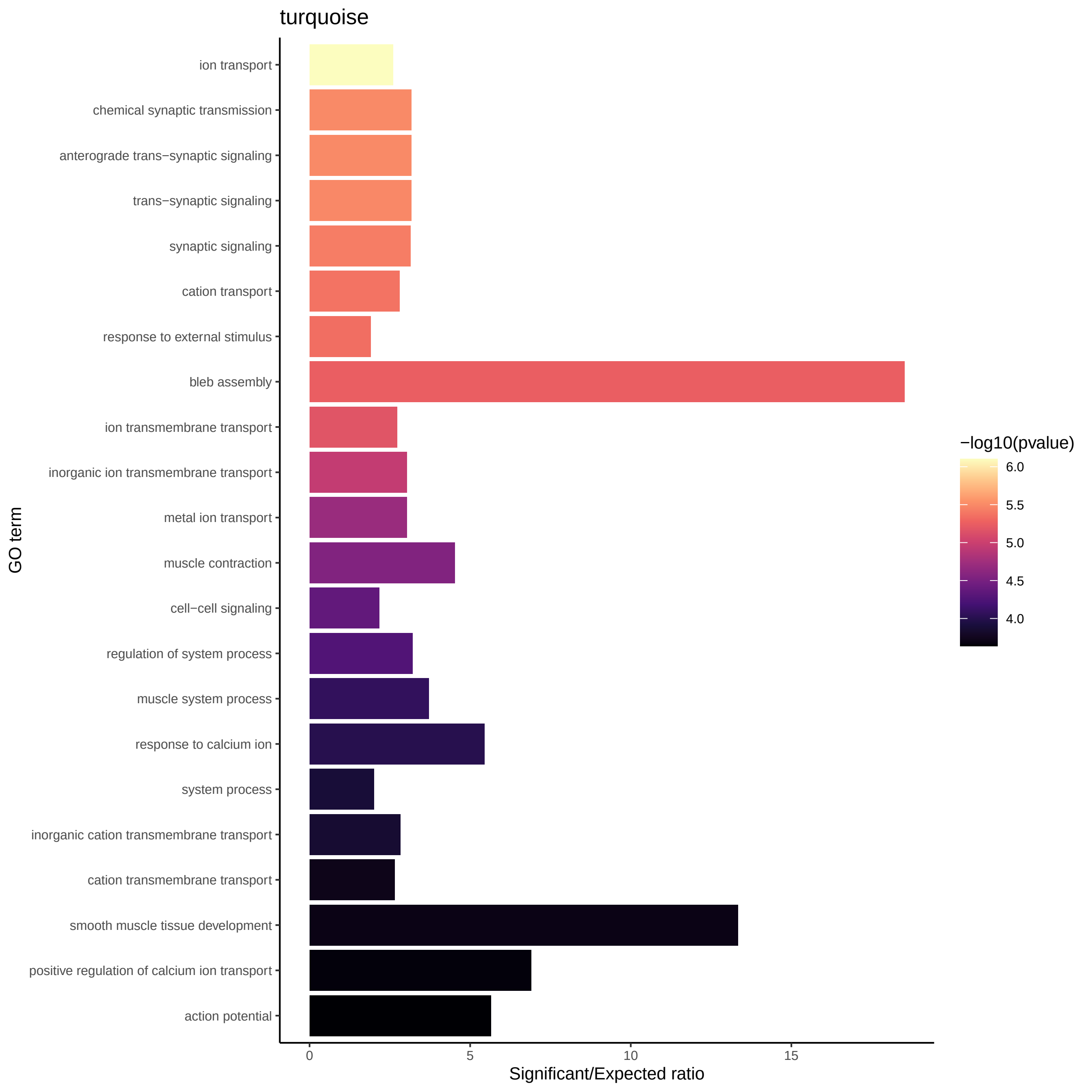
turquoise
ion transport
chemical synaptic transmission
anterograde trans−synaptic signaling
trans−synaptic signaling
synaptic signaling
cation transport
response to external stimulus
bleb assembly
ion transmembrane transport
inorganic ion transmembrane transport
metal ion transport
muscle contraction
cell−cell signaling
regulation of system process
muscle system process
response to calcium ion
system process
inorganic cation transmembrane transport
cation transmembrane transport
smooth muscle tissue development
positive regulation of calcium ion transport
action potential
GO term
0
5
10
15
Significant/Expected ratio
−log10(pvalue)
6.0
5.5
5.0
4.5
4.0

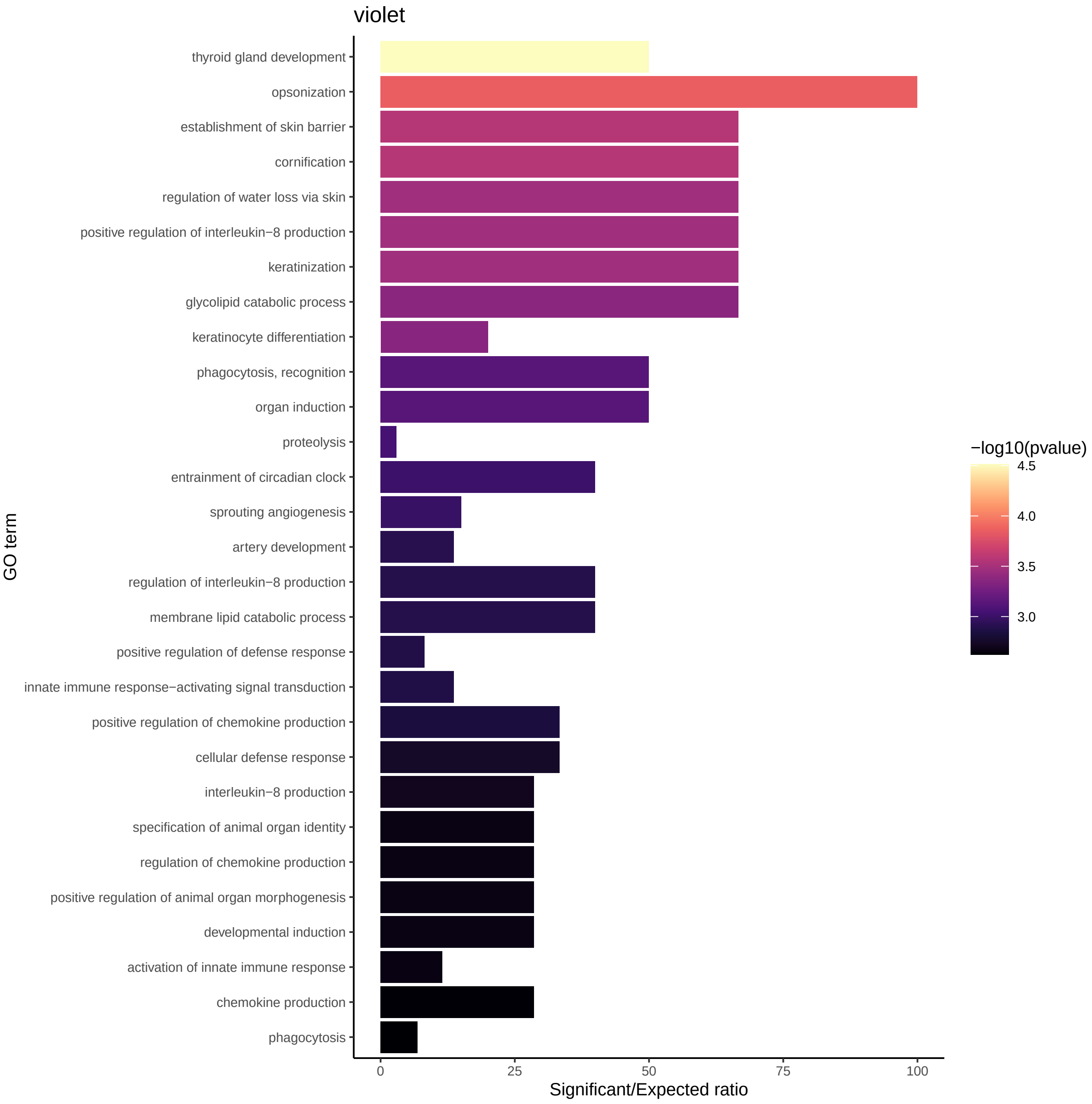

violet
thyroid gland development
opsonization
establishment of skin barrier
cornification
regulation of water loss via skin
positive regulation of interleukin−8 production
keratinization
glycolipid catabolic process
keratinocyte differentiation
phagocytosis, recognition
organ induction
proteolysis
entrainment of circadian clock
sprouting angiogenesis
artery development
regulation of interleukin−8 production
membrane lipid catabolic process
positive regulation of defense response
innate immune response−activating signal transduction
positive regulation of chemokine production
cellular defense response
interleukin−8 production
specification of animal organ identity
regulation of chemokine production
positive regulation of animal organ morphogenesis
developmental induction
activation of innate immune response
chemokine production
phagocytosis
GO term
0
25
50
75
100
Significant/Expected ratio
−log10(pvalue)
4.5
4.0
3.5
3.0

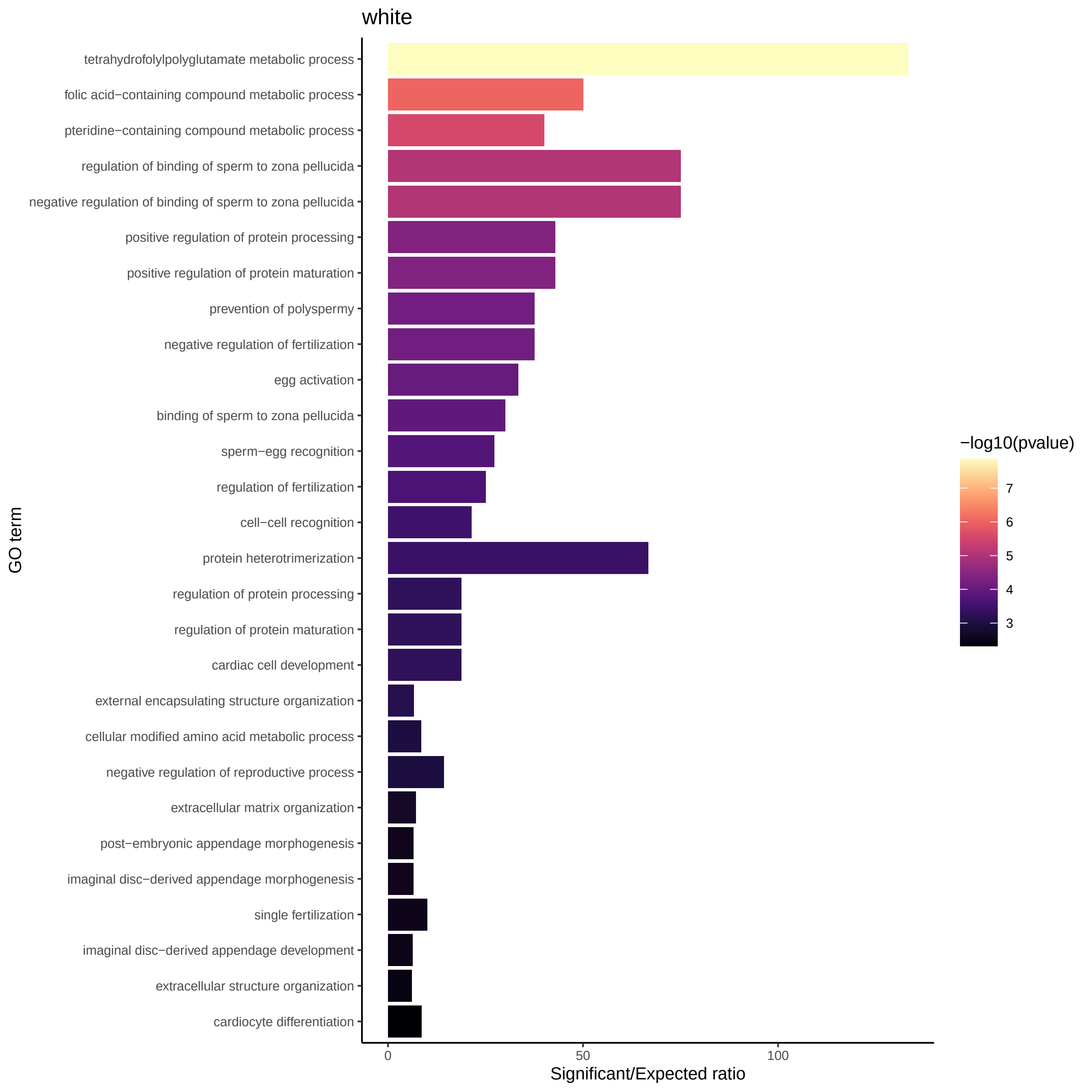
white
tetrahydrofolylpolyglutamate metabolic process
folic acid−containing compound metabolic process
pteridine−containing compound metabolic process
regulation of binding of sperm to zona pellucida
negative regulation of binding of sperm to zona pellucida
positive regulation of protein processing
positive regulation of protein maturation
prevention of polyspermy
negative regulation of fertilization
egg activation
binding of sperm to zona pellucida
sperm−egg recognition
regulation of fertilization
cell−cell recognition
protein heterotrimerization
regulation of protein processing
regulation of protein maturation
cardiac cell development
external encapsulating structure organization
cellular modified amino acid metabolic process
negative regulation of reproductive process
extracellular matrix organization
post−embryonic appendage morphogenesis
imaginal disc−derived appendage morphogenesis
single fertilization
imaginal disc−derived appendage development
extracellular structure organization
cardiocyte differentiation
GO term
0
50
100
Significant/Expected ratio
−log10(pvalue)
7
6
5
4
3

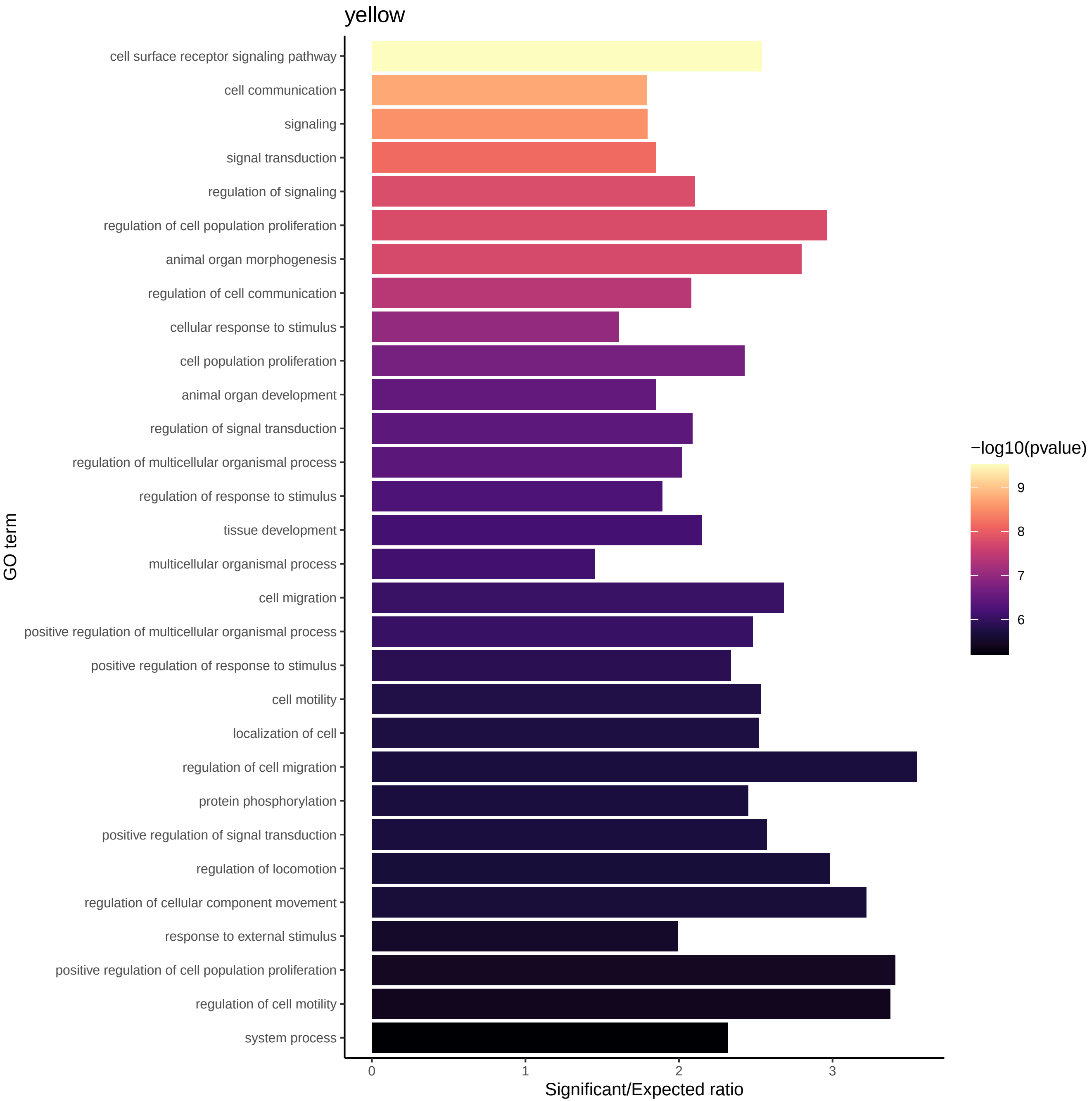
yellow
cell surface receptor signaling pathway
cell communication
signaling
signal transduction
regulation of signaling
regulation of cell population proliferation
animal organ morphogenesis
regulation of cell communication
cellular response to stimulus
cell population proliferation
animal organ development
regulation of signal transduction
regulation of multicellular organismal process
regulation of response to stimulus
tissue development
multicellular organismal process
cell migration
positive regulation of multicellular organismal process
positive regulation of response to stimulus
cell motility
localization of cell
regulation of cell migration
protein phosphorylation
positive regulation of signal transduction
regulation of locomotion
regulation of cellular component movement
response to external stimulus
positive regulation of cell population proliferation
regulation of cell motility
system process
GO term
0
1
2
3
Significant/Expected ratio
−log10(pvalue)
9
8
7
6

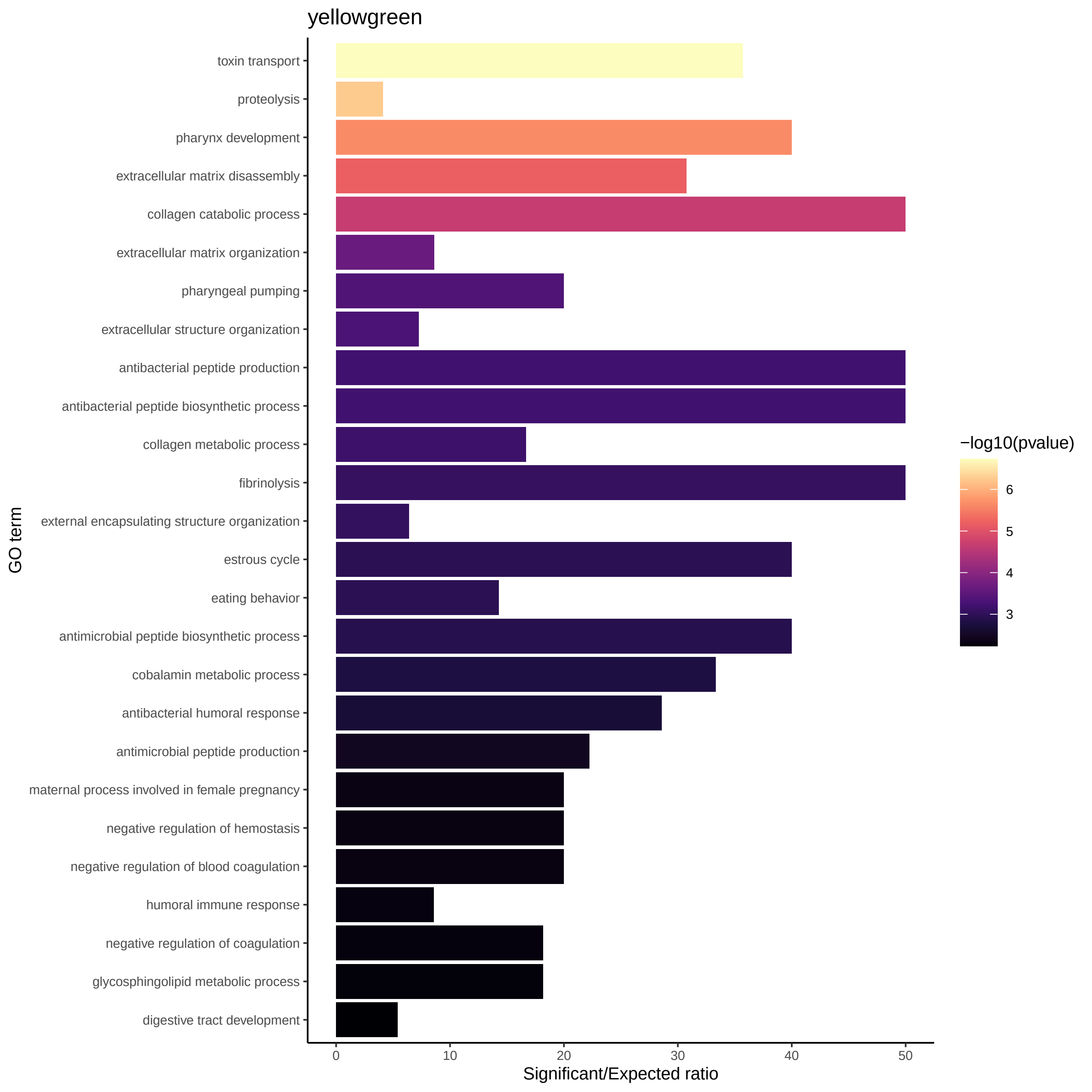
yellowgreen
toxin transport
proteolysis
pharynx development
extracellular matrix disassembly
collagen catabolic process
extracellular matrix organization
pharyngeal pumping
extracellular structure organization
antibacterial peptide production
antibacterial peptide biosynthetic process
collagen metabolic process
fibrinolysis
external encapsulating structure organization
estrous cycle
eating behavior
antimicrobial peptide biosynthetic process
cobalamin metabolic process
antibacterial humoral response
antimicrobial peptide production
maternal process involved in female pregnancy
negative regulation of hemostasis
negative regulation of blood coagulation
humoral immune response
negative regulation of coagulation
glycosphingolipid metabolic process
digestive tract development
GO term
0
10
20
30
40
50
Significant/Expected ratio
−log10(pvalue)
6
5
4
3
