## Supplementary figures and images for "The *Hydractinia* cell atlas reveals cellular and molecular principles of cnidarian coloniality"

### Supplementary Figures

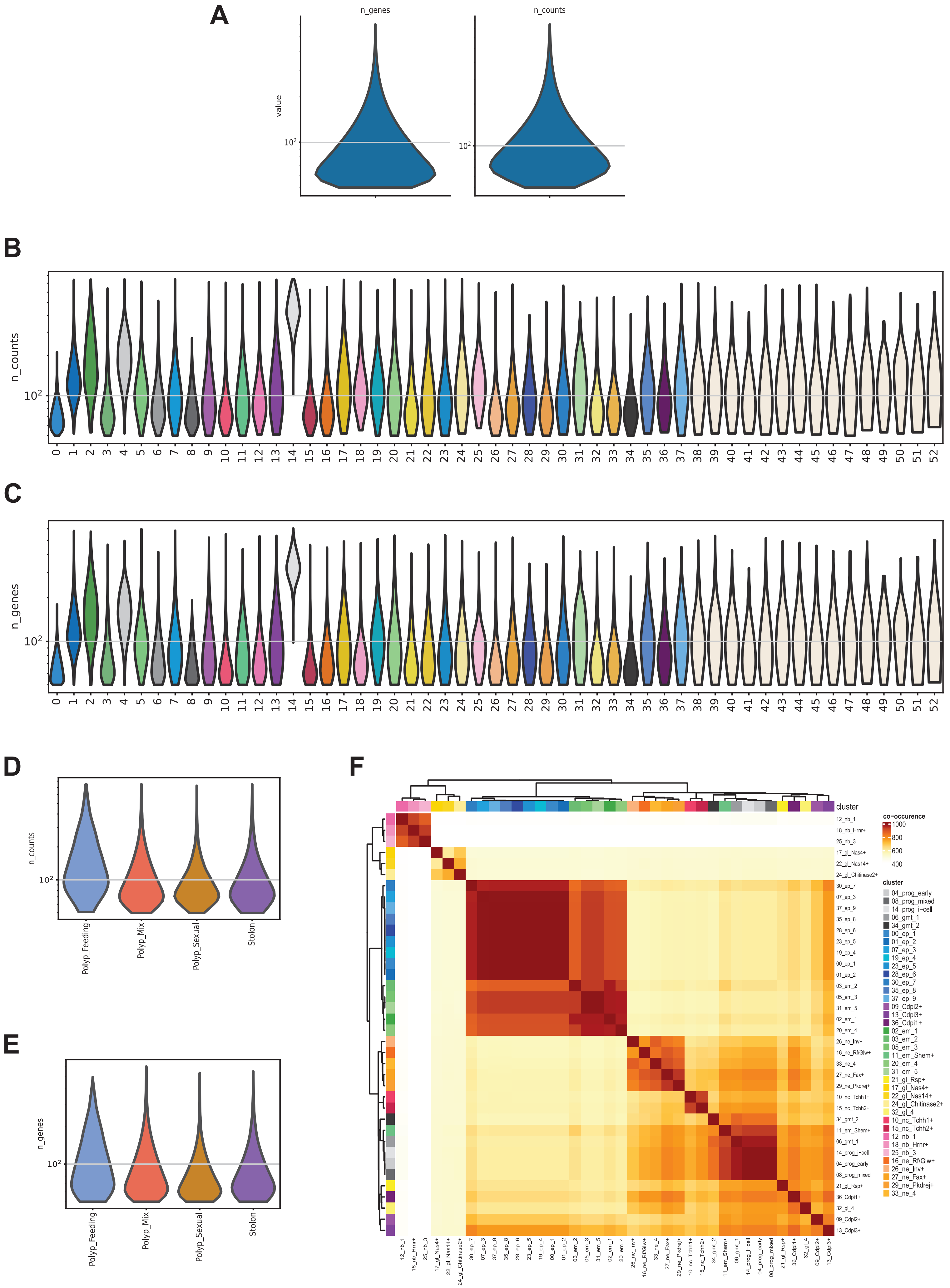

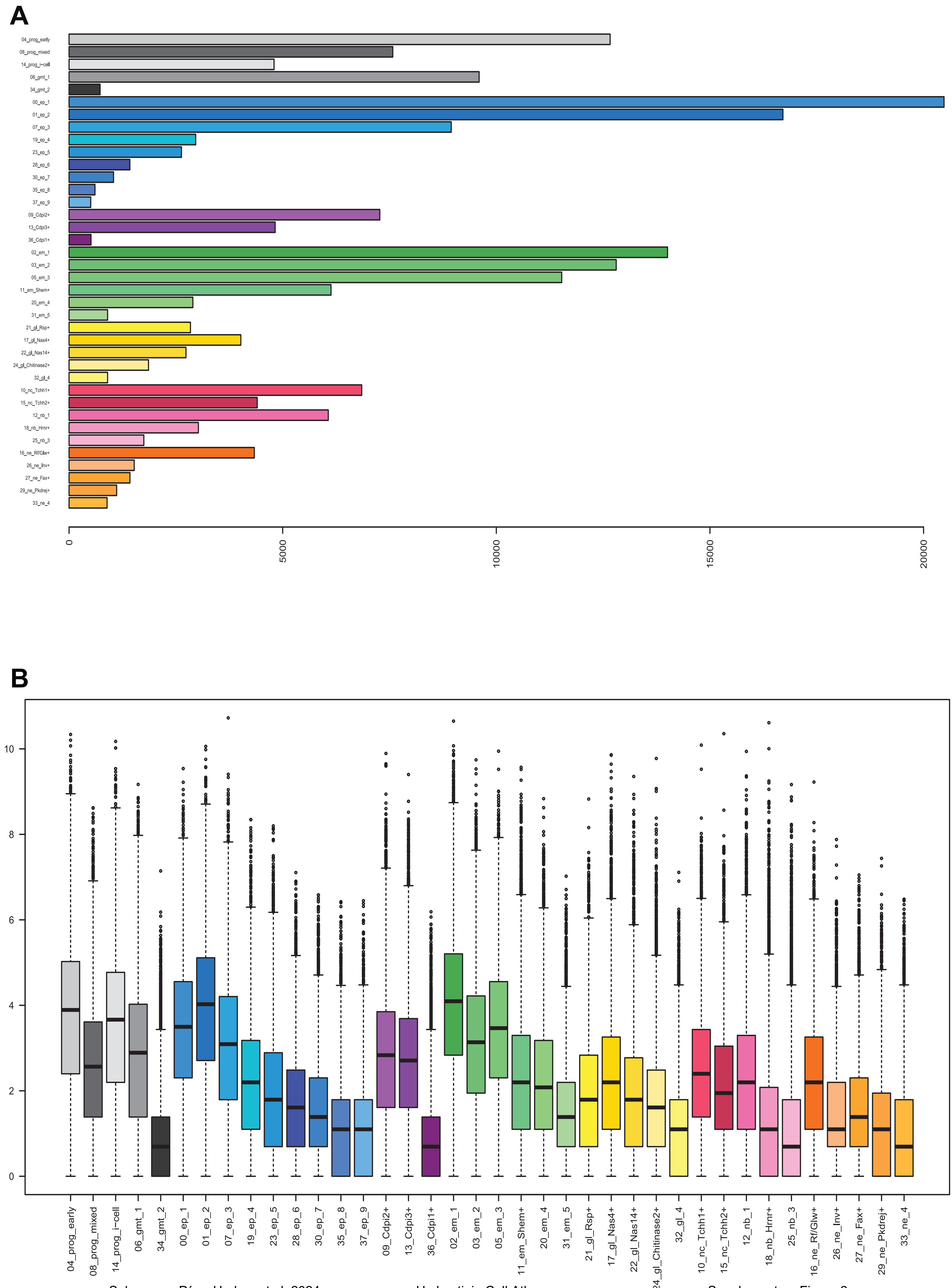

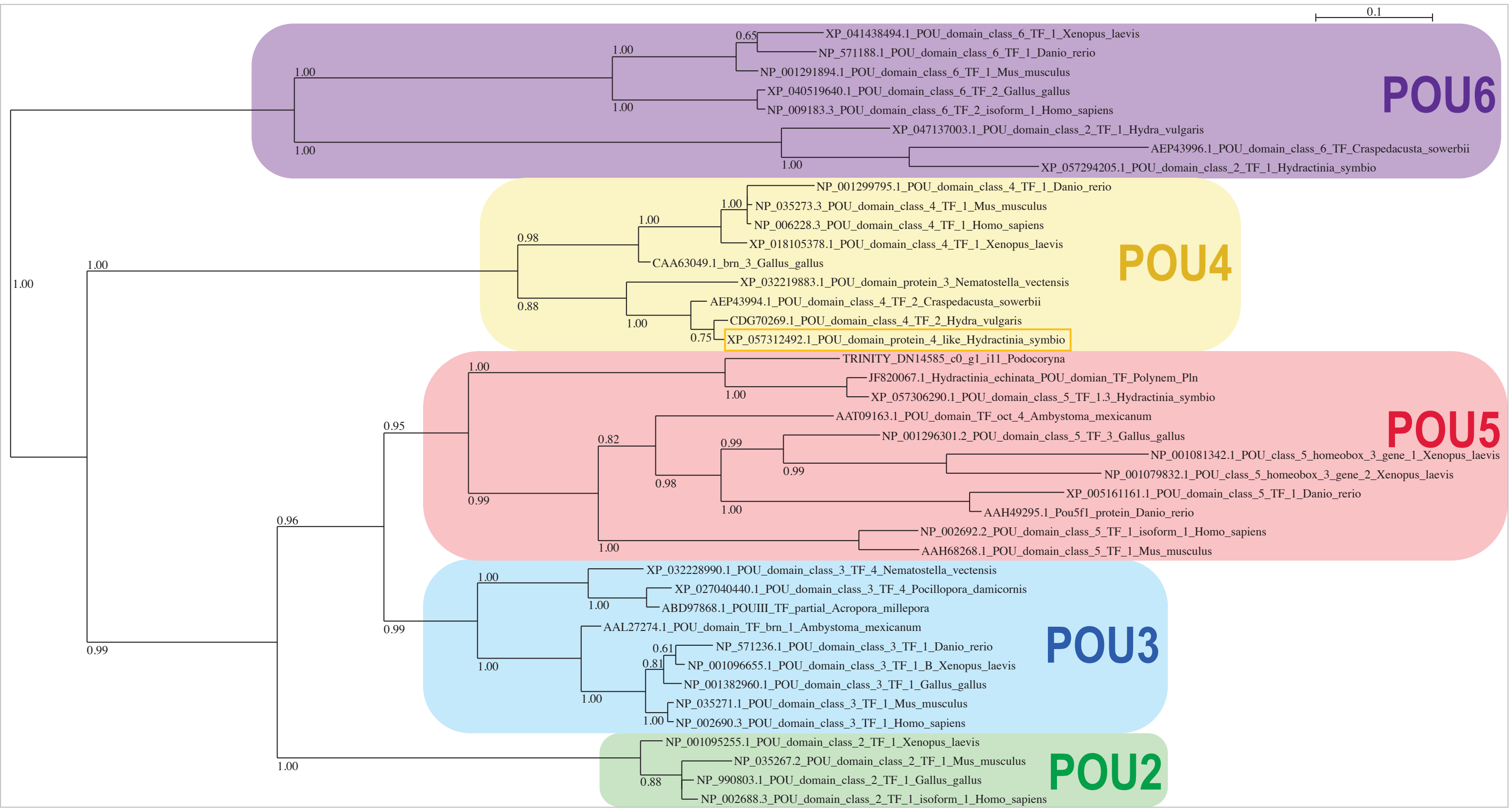

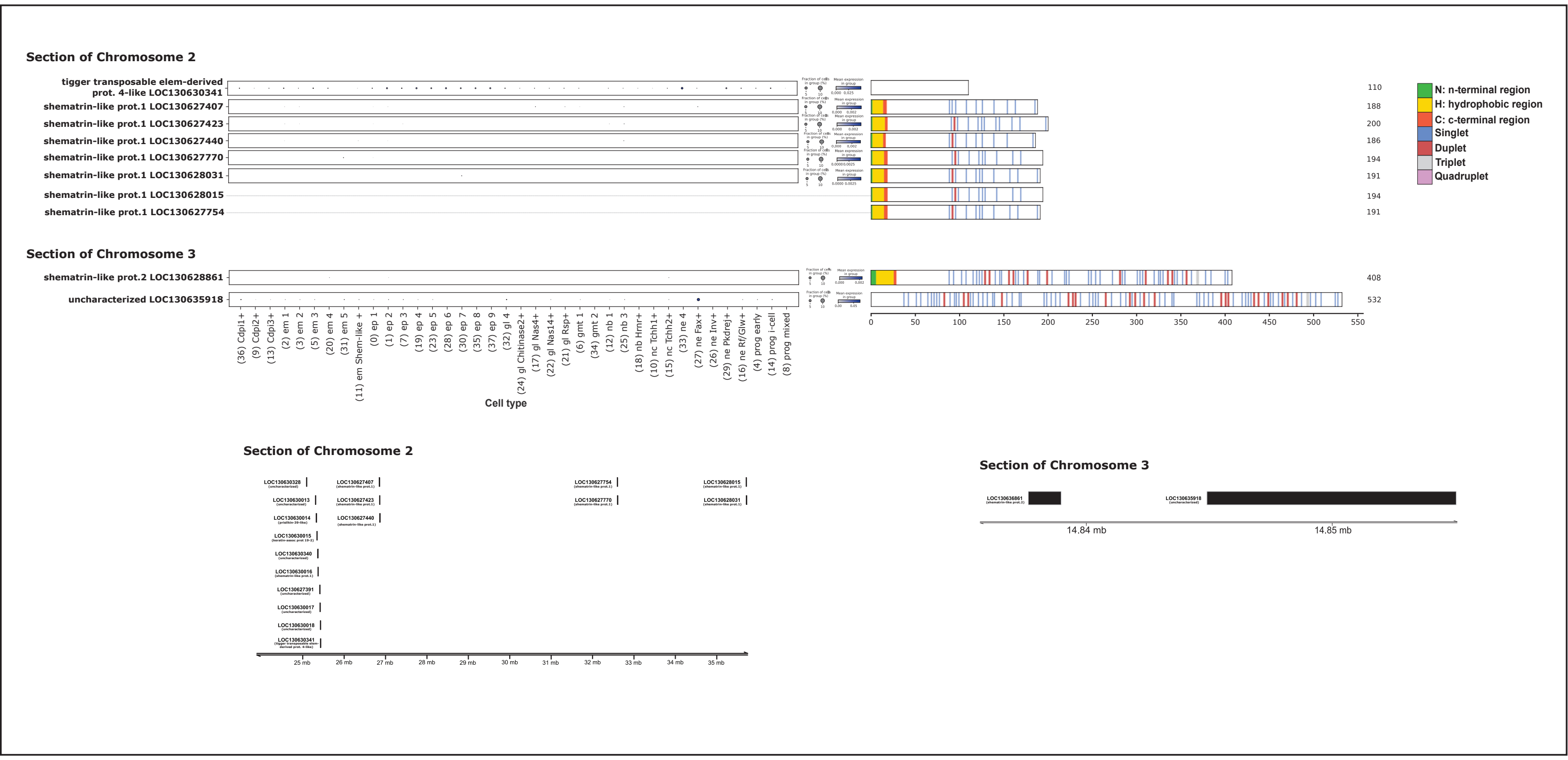

**Shematrin**

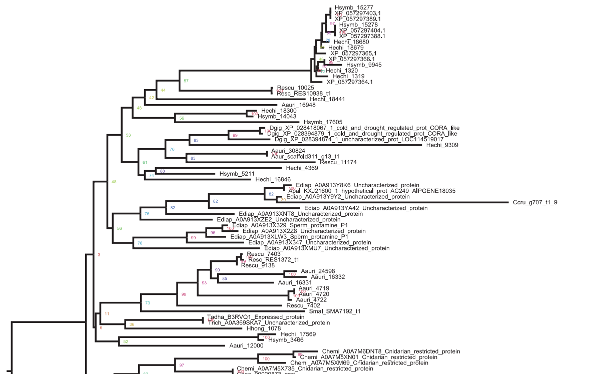

**Alpha Carbonic anhydrase**

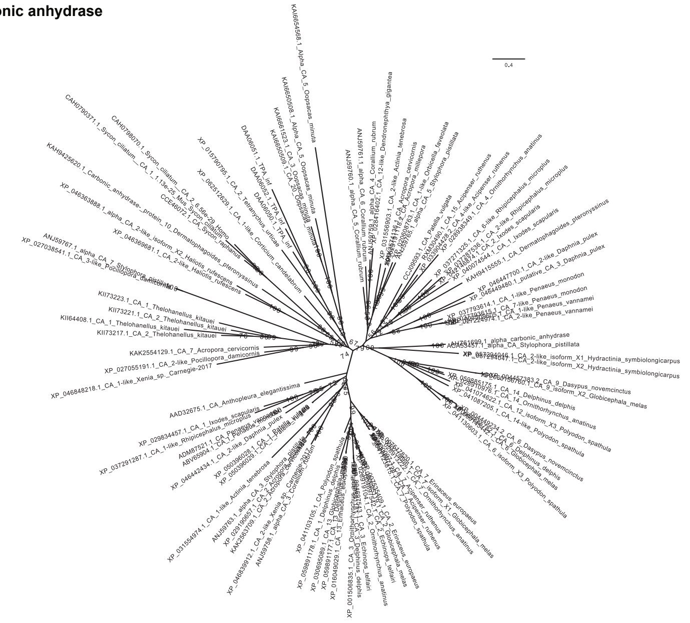

**Protocadherin**

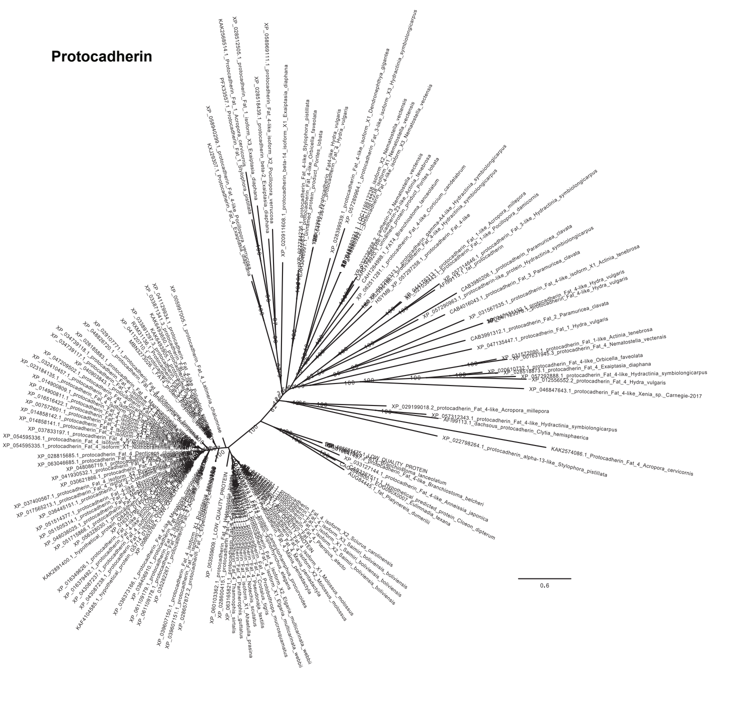

**Chitin Synthase**

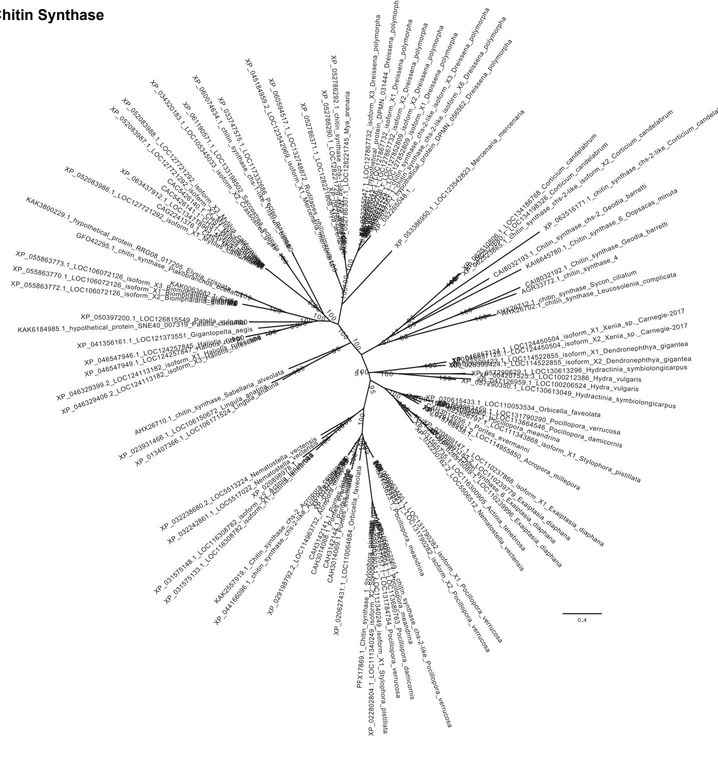
